# Novel insights into the specialized metabolism of the extremophile black yeast *Hortaea werneckii* and its modulation, unlocking its potential for compound discovery

**DOI:** 10.1101/2025.07.21.665953

**Authors:** Elise Gerometta, Bede Ffinian Rowe Davies, Rafia Ahmed Tuli, Bastien Cochereau, Annie Lebreton-Nicaise, Laurence Meslet-Cladière, Monika Kos, Nina Gunde-Cimerman, Catherine Roullier

**Author notes:** Luxembourg Institute of Science and Technology, 41 rue du Brill, L-4422 Belvaux, Luxembourg (Bastien Cochereau).

## Abstract

*Hortaea werneckii* is a halotolerant yeast, found in various habitats and which specialized metabolism remains largely unexplored. Moreover, its chemical response to high salinities has not been thoroughly investigated. To address this gap, a large-scale metabolomic study based on HPLC-HRMS/MS was conducted on 64 strains, collected from different habitats worldwide, and cultivated both on saline and non-saline media. The culture media salinity significantly modulated the strains metabolomes, suggesting the yeast exhibits a specific chemical response to high salt concentrations, potentially linked to halotolerance mechanisms. Additionally, the metabolomes were influenced by the ecological origin of the strains, with opportunistic pathogenic isolates producing distinct metabolites. Molecular networking revealed that *H. werneckii* synthesizes various chemical classes, including potential cytotoxic compounds, making this yeast a potentially interesting source of bioactive compounds. Hortein was detected as dominant and ubiquitous, emerging as a potential chemical marker for the species. This study highlights several compounds of interest, related to the chemical ecology and pathogenicity of *H. werneckii*. As most of them remain unidentified, future research should prioritize their isolation and identification, to improve our understanding of this extremophile organism and harness its potential in natural product discovery.

## 1. Introduction

*Hortaea werneckii* (Horta) Nishim. & Miyaji, 1984 (Ascomycota, Dothideomycetes) is a filamentous and extremophile black yeast, present primarily in hypersaline waters (Marchetta et al., 2018; Zalar et al., 2019). It is described as one of the most salt-tolerant eukaryotic organisms, able to grow from 0 to 30% NaCl (Marchetta et al., 2018). This species is globally distributed and has been isolated from both marine and hypersaline and osmotically stressed terrestrial environments, from the Atacama Desert to deep sea water and hydrothermal vents (Burgaud et al., 2010; De Leo et al., 2019; Gostinčar et al., 2022; Gunde-Cimerman and Plemenitaš, 2006; Zalar et al., 2019). Salty human skin is another ecological niche of *H. werneckii* (Marchetta et al., 2018). This yeast is the causative agent of *tinea nigra*, a superficial skin infection of hands and soles occurring mostly in tropical and subtropical areas (De Leo et al., 2019; Marchetta et al., 2018; Zalar et al., 2019).

So far, this species was mainly studied as a biological model to understand the halotolerance mechanisms in eukaryotic cells (Gunde-Cimerman et al., 2018; Gunde-Cimerman and Zalar, 2014; Marchetta et al., 2018; Zalar et al., 2019), and has been subject to several studies performed at the genetic, molecular and ecological levels (De Leo et al., 2019; Gostinčar et al., 2022; Gunde-Cimerman et al., 2018; Gunde-Cimerman and Plemenitaš, 2006; Gunde-Cimerman and Zalar, 2014; Marchetta et al., 2018; Zalar et al., 2019). Regarding its chemistry, very few investigations have been carried out. In fact, the yeast has been mainly studied for the production of melanin pigment (Elsayis et al., 2022), and only three other specialized metabolites have been isolated: hortacerebosides A and B (Chen et al., 2022) and hortein (Brauers et al., 2001). Nevertheless, *H*. *werneckii* seems to be an interesting candidate for the discovery of new metabolites and bioactive molecules, as 27 biosynthetic gene clusters have been identified in its genome (Blin et al., 2021; Grigoriev et al., 2013; Lenassi et al., 2013), and enzymes involved in sulphated and halogenated compounds synthesis have been purified from this yeast (Cochereau et al., 2023; Graziano et al., 2024). Also, while it has been reported that saline conditions induced an extremophilic ecotype in *H. werneckii*, characterized amongst others by enhanced production of melanin pigment, meristematic growth and synthesis of compatible solutes such as glycerol (Gunde-Cimerman et al., 2018; Gunde-Cimerman and Plemenitaš, 2006; Zalar et al., 2019), the effect of salt concentration on its overall specialized metabolome has not been investigated.

As numerous *H. werneckii* strains have been previously collected from various environments around the world and submitted to genetic and molecular analysis (Gostinčar et al., 2022, 2018; Zalar et al., 2019), this gave the opportunity to conduct a large-scale study on the species, to explore its specialized metabolism potential and the impact of several factors on its metabolite production. Therefore, here is reported an unprecedented work conducted through high resolution mass spectrometry (HRMS)-based metabolomics, in order to provide new insights on the chemodiversity and chemical ecology of the yeast, and its potential for compound discovery.

## 2. Experimental Procedures

### 2.1. Chemicals

Ethyl acetate (EtOAc) used for the extraction was of analytical grade quality and purchased from Carlo Erba (Val de Reuil, France). It was distilled before use. Acetonitrile, methanol and water used for sample preparation and HRMS/MS analyses were of LC/MS grade quality and purchased from Fisher Chemical.

### 2.2. Fungal strains and culture

Sixty-four strains (NCBI Bioproject Accession: PRJNA428320 ID: 428320) were obtained from the Ex Culture Collection of the Department of Biology, Biotechnical Faculty, University of Ljubljana, Slovenia (Infrastructural Centre Mycosmo, Network of Research and Infrastructural Centres University of Ljubljana, Slovenia) (Gostinčar et al., 2022, 2018). Each strain is described in **Table 1** (accession number, sampling habitat, sampling location, ploidy level, vHPO genes). Strains were revived and cultivated on Malt Extract Agar (MEA) medium (malt extract 20 g/L, peptone 1 g/L, glucose 20 g/L, agar 20 g/L) at 25 °C for 7 days. Subsequently, each strain was inoculated in triplicate onto 7 cm MEA plates, with and without 17% (w/v) NaCl (170 g/L), using a 10 µL loop. The plates were incubated at 25 °C for 14 days, yielding a total of 384 fungal samples (64*3*2). After incubation, fungal biomass was carefully harvested from the plates and immediately stored at –80 °C. Sterile media (with and without 17% salt) served as blank controls, with fourteen blanks prepared and processed under the same conditions as the biomass samples (seven for each medium type). All fungal and blank samples were freeze-dried at –60 °C for 24 hours at 0.5 mbar, followed by an additional 2 hours at 0.09 mbar (Christ Alpha 1-4 LSCbasic, Martin Christ Gefriertrocknungsanlagen GmbH, Germany).

**Table 1.**
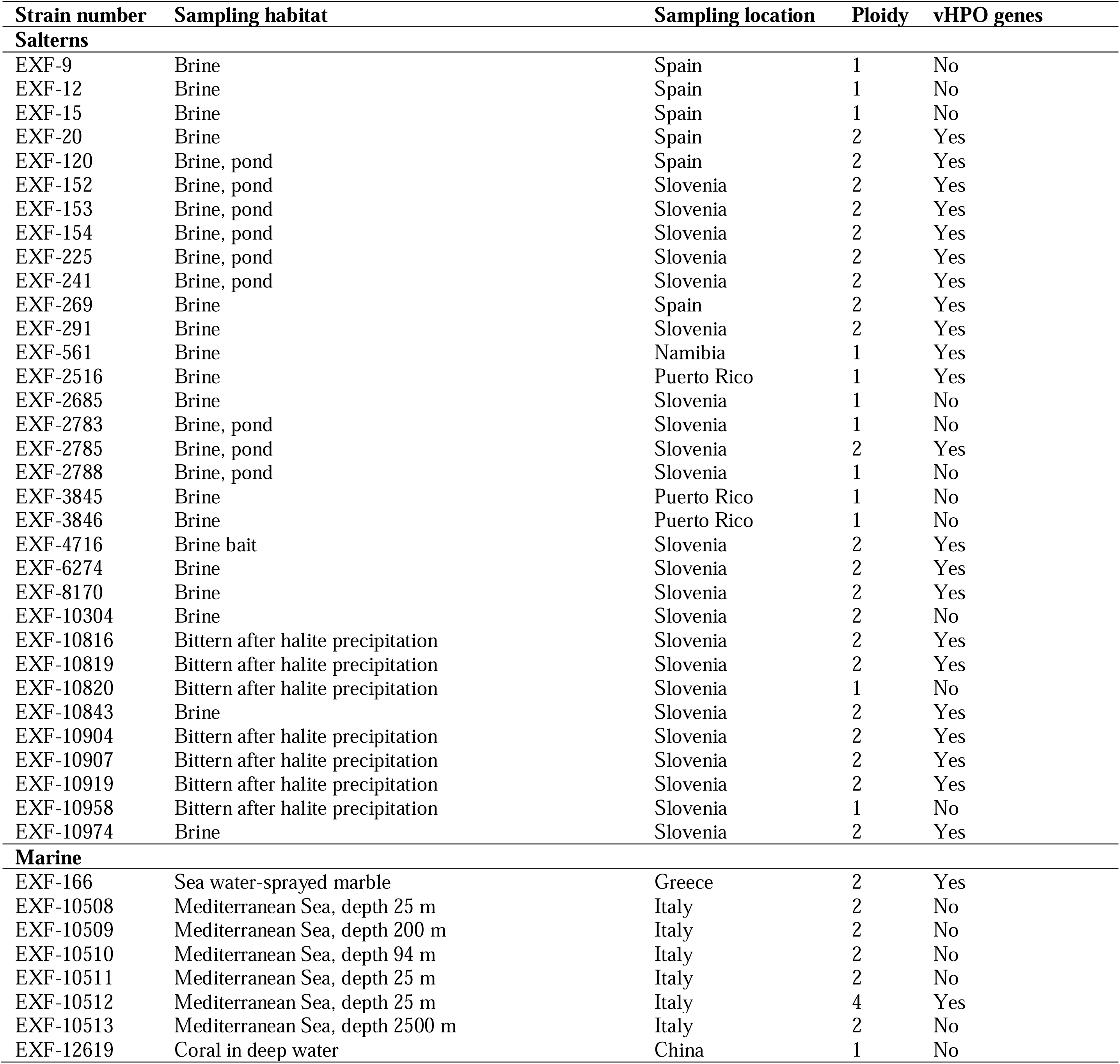
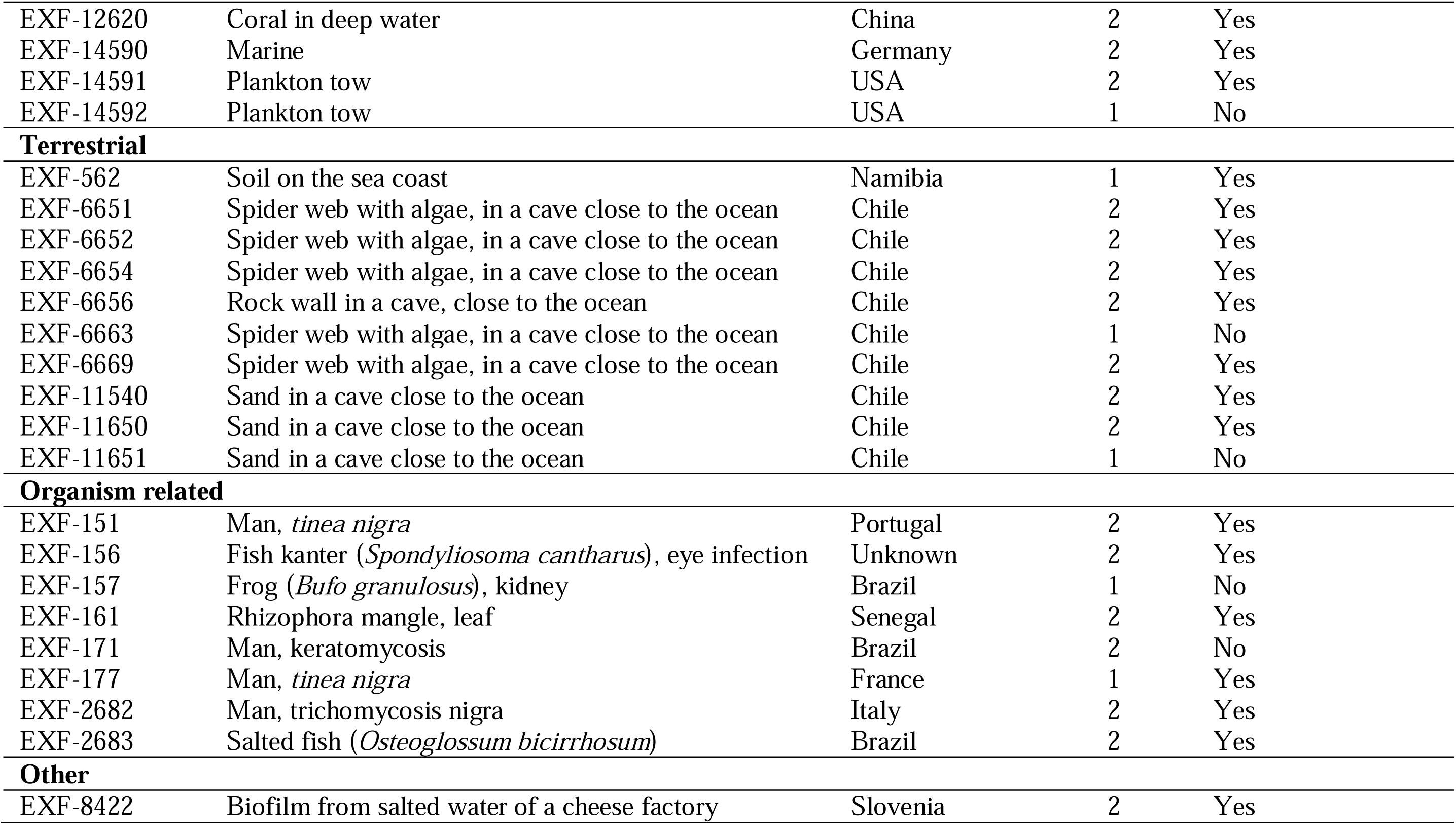
The 64 *H. werneckii* strains subjected to the large-scale metabolomic study.

### 2.3. Extraction

The 398 samples (384 fungal samples and 14 blanks) were extracted following the same procedure: 2 mL of EtOAc were added and the samples were sonicated for 30 minutes (Elmasonic S300, Elma, Singen, Germany),1.5 mL of supernatant was collected and evaporated under nitrogen flow at 40 °C. These steps were repeated two more times, to perform three successive extractions. 398 crude extracts, ranging from 0.4 to 6.5 mg, were thus obtained and stored at −20°C.

### 2.4. HPLC-HRMS/MS profiling

The extracts were profiled on a Shimadzu UFLC-HRMS/MS-IT-TOF instrument, equipped with two LC-20ADxr pumps, a SIL-20ACxr autosampler, a CTO-20AC column oven, an SPD-M20A PDA detector, a CBM-20A system controller, an ESI ion source and an IT-TOF mass spectrometer (Shimadzu, Kyoto, Japan). The analysis was performed in a mass range from *m/z* 100 to 1550, using negative mode (-) for the ESI source. The positive mode (+) was also explored but a poor ionization efficiency was observed, therefore the data could not be properly interpreted. The following parameters were used: heat block and curved desolvation line temperature 200 °C; nebulising nitrogen gas flow 1.5 L/min; interface voltage (-) – 3.5 kV; detector voltage of the TOF analyser 1.6 kV. High performance liquid chromatography analyses were performed on a 100 × 2.1 mm 2.6 µm RP-C_18_ column (Kinetex, Phenomenex, Torrance, CA, USA), heated in an oven at 40 °C, and using an elution gradient of H_2_O-CH_3_CN with 0.1% HCO_2_H (85:15 *v/v* over 2 min, 85:15 *v/v* to 0:100 *v/v* over 23 min, 0:100 *v/v* over 5 min) at a flow rate of 0.3 mL/min. The samples were prepared in 100% MeOH, at a concentration of 0.5 mg/mL, and filtered through a 0.45 µm filter (Chromafil AO 45/15 MS, Macherey-Nagel). The injection volume was 3 µL. Samples were analysed in two randomised batches.

### 2.5. HRMS/MS data processing and Feature-Based Molecular Networking

Raw data were converted into the open format .mzXML, calibrated and processed using software Mzmine Version 4.5.0 (Schmid et al., 2023). The data processing steps were as follows: .mzXML raw data import, mass detection, chromatogram builder, smoothing, local minimum feature resolver, ^13^C isotope filter, isotopic peaks finder, feature list rows filter, join aligner, feature finder (gap filling), feature list blank subtraction and duplicate peak filter. Setting parameters were as follows: negative ionization mode, centroid detection, MS1 peak detection limit: 1.0E4, MS2 peak detection limit: 1.0E4; minimum consecutive scans:3, minimum intensity for consecutive scans: 2.0E4, minimum absolute height: 5.0E4, *m/z* tolerance: 50.0 ppm; smoothing algorithm: Savitzky-Golay; MS1 to MS2 precursor tolerance: 50.0 ppm, chromatographic threshold: 80%, minimum search range RT/mobility: 0.070, minimum absolute height: 5.0E4, minimum ratio of peak top/edge: 1.70, peak duration range: 0.05-1.00 min, minimum scans: 3; representative isotope: most intense, *m/z* tolerance 10.0 ppm, retention time tolerance: 0.05 min, maximum charge: 3; chemical elements: H, C, N, O, S, Cl, Br, *m/z* tolerance: 10.0 ppm, search in scans: single most intense; filtering: retention time: 0.08-30.00 min; aligner: *m/z* tolerance: 50.0 ppm, weight for *m/z*: 75, weight for RT: 25, retention time tolerance: 0.50 min, mobility weight: 1; gap-filling: intensity tolerance: 15%, *m/z* tolerance: 10.0 ppm, retention time tolerance: 0.30 min, minimum scans: 3; blank/control raw data files: 21, minimum of detection in blanks: 1, quantification: area, ratio type: average, fold change increase: 300.0%, only keep features above fold change; duplicate peak: *m/z* tolerance 50.0 ppm, RT tolerance: 0.50 min. After data processing, a .csv file was exported and used as a matrix for statistical analysis. A metadata table was created with the following attributes: culture media salinity, ploidy, ecological niche and vHPO (vanadium HaloPerOxidase enzyme genes). To create the feature-based molecular network (FBMN), an additional step was included in data processing: feature list rows filter: keep only feature with MS2 scan. Processed files, including an .mgf and .csv file, as well as the metadata table, were uploaded to the GNPS platform (Wang et al., 2016). The FBMN was developed using the Advanced Analysis Tools – Feature Networking workflow (Nothias et al., 2020), and is available *via* the following link: https://gnps.ucsd.edu/ProteoSAFe/status.jsp?task=108a1228673741e98650d5b621ecb8c4. Parameters were set as follows: precursor ion mass and fragment ion mass tolerance: 0.02 Da, min pairs cos: 0.4, min matched fragment ions: 3, network topK: 1000. The FBMN was then exported and visualized using software Cytoscape Version 3.10.3.*2.6*.

Dereplication of the features of interest was performed with the Dictionary of Natural Products (DNP 33.2, 2024 – Type of organisms: Z.G.30000-Ascomycetes), the Natural Products Atlas (NPAtlas 3.0.2, 2024 – Fungi, Phylum: Ascomycota), the LOTUS Natural Products Online (Rutz et al., 2022), with GNPS spectral databases (score threshold: 0.7 and search min matched peaks: 3), and using ChromAnnot tool (Bertrand et al., 2017) and software SIRIUS Version 6.1.1 (Dührkop et al., 2019).

### 2.6. Multivariate statistical analysis

#### 2.6.1. Metabolome analysis

Permutational multivariate analysis of variance (PERMANOVA) (Anderson et al., 2008; Clarke and Gorley, 2015) was used from the “vegan” package in R (Oksanen et al., 2025; R Core Team, 2024) to test differences between ecological niche, culture media salinity, ploidy level and presence of vHPO genes for feature area data. The feature area was modelled as a function of the interaction between Ecological Niche and Culture media salinity, and Ploidy level and Presence of vHPO genes (Ecological Niche, 4 levels: Marine, Organism, Terrestrial and Saltern; Culture Media, 2 levels: Saline and Non-Saline; ploidy 3 levels: 1, 2 and 4; Presence of vHPO genes, 2 levels: Yes and No). Multivariate analyses were carried out on the basis of a Bray–Curtis dissimilarity matrix, calculated from fourth root transformed area data (normalized data). The statistical significance of the variance components was tested using 9,999 permutations under a reduced model (Anderson et al., 2008). Visualization of multivariate data was carried out by a non-metric multidimensional scaling (nMDS) ordination performed on the Bray-Curtis dissimilarity matrix.

#### 2.6.2. Features of interest

A subset of features was selected by *Two Different styles of Witchcraft* and analyzed by a multivariate General Linear Model (GLM) to assess how influential features change depending on the ecological niche of the strains and culture media salinity. The subset of features was modelled as a function of the interaction of ecological niche (*E*) and culture media salinity (*S*), where the response variable was a multivariate matrix of the centered and scaled area of each feature (M_i_) (Equation 1).

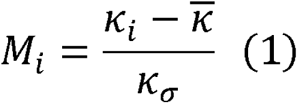

Where K_i_, 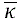 and K_a_ were the raw, mean and standard deviation of the area for each feature respectively (Equation 1). Model parameters were estimated within a bayesian framework using the ‘brms’ and ‘RStan’ packages in R to leverage the Stan language

(Bürkner, 2021; Carpenter et al., 2017; Stan Development Team, 2018). The multivariate response variable matrix was modelled assuming Gaussian distributions, with weakly informative priors. Model parameters were estimated using Markov Chain Monte Carlo (MCMC) sampling, with 4 chains of 10,000 iterations and a warm-up of 500.

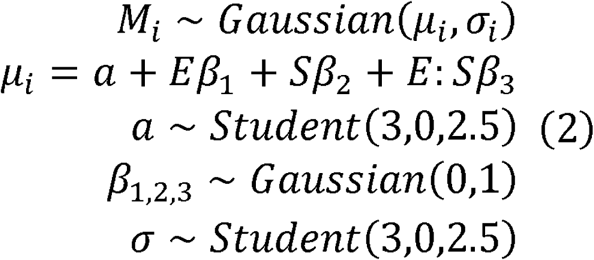

### 2.7. Univariate statistical analysis

For all features, Shannon’s Index (H Equation 3), Number of Features (R), Gini’s Coefficient (G Equation 4) and Pielou’s Index (D_pie_ Equation 5) were calculated to assess how the diversity, dominance and evenness of features changed with ecological niche and culture media salinity using GLMs.

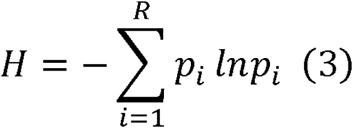

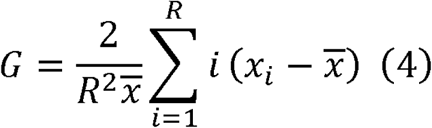

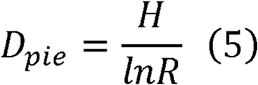

For each sample, p_i_ is the area of a specific feature proportional to the total area across all features within the sample, R is the total number of different features present in the sample and x_i_ is the area of a specific feature. Univariate indices were modelled as a function of the interaction between Ecological Niche and Culture media salinity. Shannon’s index was modelled assuming a Gaussian distribution (Equation 6), Number of Features a Poisson distribution (Equation 7), while both Gini’s Coefficient and Pielou’s Index were assumed to follow Beta distributions (Equation 8 and Equation 9). As above, model parameters were estimated within a bayesian framework using the ‘brms’ and ‘RStan’ packages in R to leverage the Stan language (Bürkner, 2021; Carpenter et al., 2017; Stan Development Team, 2018) using Markov Chain Monte Carlo (MCMC) sampling, with 4 chains of 10,000 iterations and a warm-up of 500.

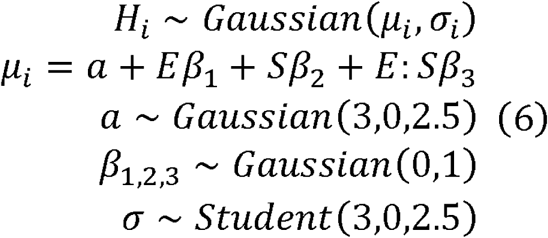

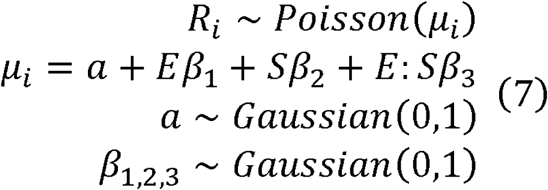

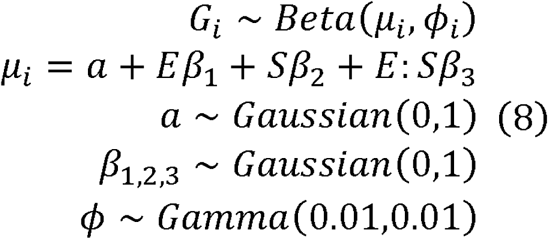

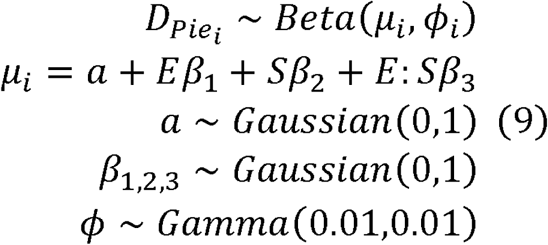

### 2.8. Specialized metabolism related genes

Specialized metabolism associated genes, including PolyKetide synthase of type I (PKS I), Non-Ribosomal Peptide Synthetase (NRPS), Terpene Synthase (TS), DiMethylAllyl Tryptophan Synthase (DMATS), NRPS-PKS or NRPS-TPS Hybrid and post-translationally modified peptides (RiPPs), were searched in each genome sequences of the 64 *H. werneckii* strains. Gene cluster associated with specialized metabolites were searched with antiSMASH v.4 fungal version (FungiSMASH) (Blin, 2021). To complement these analyses, and because some genomes were too fragmented, the presence or absence of genes encoding enzymes involved in biosynthetic pathways was also manually investigated.

### 2.9. Molecular phylogenetic analysis of PKSI and Indole sequences

Protein sequences were aligned using MAFFT v.7.525. Ambiguous regions (containing gaps and poorly aligned) were removed with TRIMAL v.1.4.1. MODELTEST-NG v.0.1.7 was then used to identify the best protein substitution model for each gene family. Gene family trees were reconstructed based on these trimmed alignments with RAXML-NG v.1.2.2 using the WAG+I model for indole family and LG+I+G4+ for PKS family, 100 bootstrap replicates were performed. (Capella-Gutiérrez et al., 2009; Darriba et al., 2020; Kozlov et al., 2019; Kuraku et al., 2013)

## 3. Results and Discussion

The study was performed on 64 sequenced strains (**Table 1**), isolated from different habitats around the world (**Figure 1**), with the aim of exploring the overall chemical diversity of *H. werneckii* and evaluating the potential impact of salinity, as well as other factors such as the ecological niche of the strains, on its metabolite production. A genome mining of the 64 strains was performed, yielding to the summary of predicted BGC counts and classes presented in **Figure 2** and **Appendix A**, and highlighting the presence of gene clusters such as hybrid PKS and NRPS-like clusters. The total number of predicted BGCs is consistent with previous work (Rokas et al., 2018). Even though this number is not very high compared to filamentous fungi, this specialized metabolism of *H. werneckii* remained worthy of interest as it had never been investigated before. Moreover, this is the first large-scale metabolomics study performed on 64 strains of the same species. The 64 strains were thus grown in triplicate on MEA culture media, without salt and with addition of 17% salt (NaCl). This is a high salinity which is not tolerated by other fungi, and is the highest salt concentration *H. werneckii* tolerates without affecting its growth rate (Petrovič et al., 2002). Regarding the limited amounts of fungal cultures, they were only extracted with ethyl acetate (EtOAc), commonly used for metabolomics studies, as it allows to extract a fairly wide range of mid-polar to apolar metabolites. The obtained extracts (from 0.4 to 6.5 mg) were subjected to HPLC-HRMS/MS analysis.

**Figure 1.**
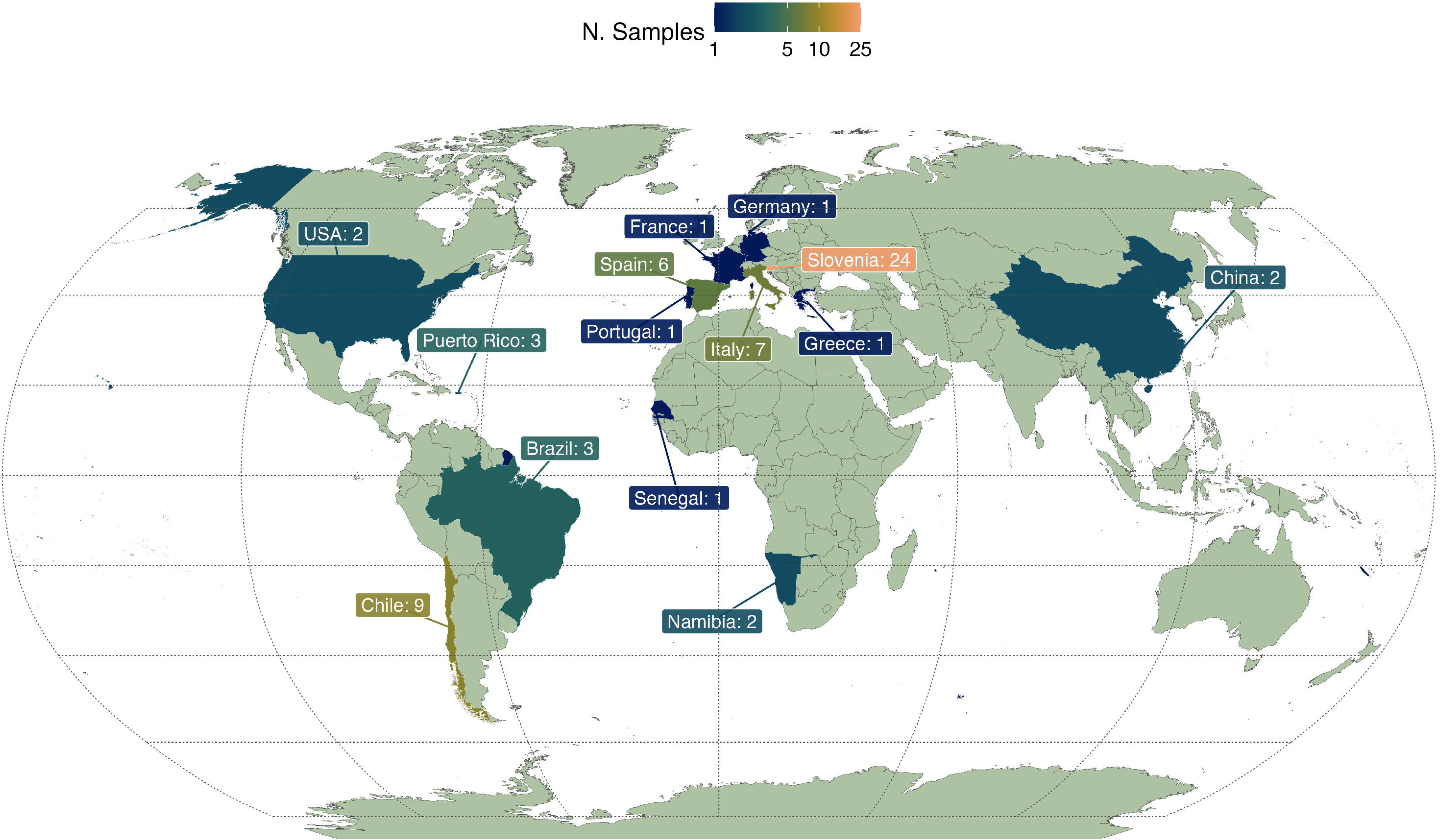
Origin of all samples studied. Color shows the number of samples per country on a log scale.

**Figure 2.**
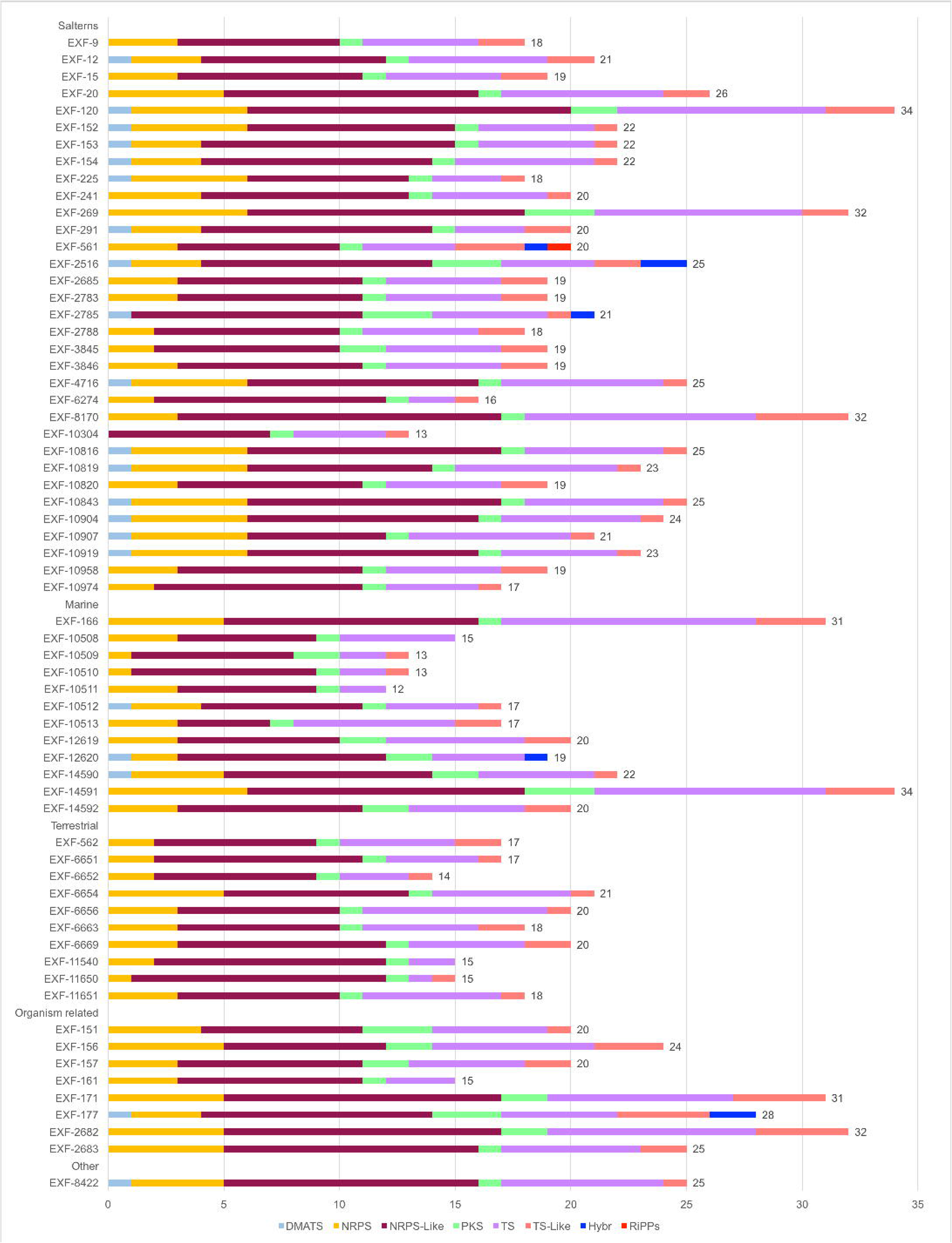
Distribution of biosynthetic gene clusters (BGCs) across the genomes of the 64 *H. werneckii* strains. The numbers indicate the total BGCs count for each strain. DMATS: DiMethylAllyl Tryptophan Synthase, NRPS: Non-Ribosomal Peptide Synthetase, PKS: Polyketide Synthase, TS: Terpene Synthase, TS-like: polyprenyl synthase synthesizing precursor substrates for PTS, Hybr: NRPS-PKS or NRPS-TS Hybrid, RiPPS: Post-translationally modified peptides.

### 3.1. Modulation of the overall metabolite production

The first factor examined in this study was the salinity of the culture media, in order to determine whether the extremophilic ecotype of the yeast could be associated with a distinct specialized metabolism. The strains have been collected from 4 distinct ecological environments, namely salterns, marine, terrestrial and organism-related environments (**Table 1**). Given the diversity of sampling environments, particular attention was paid to assess how the ecological niche influences the metabolomic profiles of the strains. Finally, as information about the genome of the strains is available, the effect of the ploidy level of the strains (haploid, diploid or tetraploid) and the presence/absence of vanadium HaloPerOxidase (vHPO) genes was assessed. vHPO are enzymes able to halogenate various substrates via the formation of reactive species such as BrOH or ClOH. (Cochereau et al., 2023), and the transcription of a vHPO gene by *H. werneckii* was demonstrated when cultivated on different media including the Wickerham medium (Cochereau et al., 2023), which is close to MEA. The presence/absence of genes coding for these enzymes could induce a significant change in the metabolite production by the yeast. By evaluating the impact of these 4 factors, this investigation allowed us to better understand *H. werneckii* specialized metabolism and its adaptation to its environment.

One hundred and thirty-two samples, corresponding to 128 fungal extracts (one replicate per strain per medium) and 4 blank extracts, were analyzed through HPLC-HRMS/MS. Processing of the HRMS data resulted in a matrix of 164 features, which was then subjected to statistical analyses. While this number of features may appear quite limited, it is not so surprising considering the fact that we are dealing with a unicellular yeast, usually known to lack or contain very few BGCs (Rokas et al., 2018). As mentioned above, while *H. werneckii* may be one of the most productive among yeasts with quite a high number of BGCs, it is still quite low in comparison with filamentous fungi. Moreover, in another study performed an ethyl acetate extract of a *H. werneckii* strain cultivated in large scale (personal data), we could observe that most of the extract was composed of fat, reinforcing the fact that primary metabolism is still preponderant in *H. werneckii*. Nevertheless, even with a limited number of features, the specialized metabolism of *H. werneckii* was considered to be interesting to dig in.

A Permutational Multivariate Analysis of Variance (PERMANOVA) was first performed to assess differences or similarities in metabolomes between culture media salinity, ecological niche, ploidy level and presence of vHPO genes. According to this analysis, it appeared that samples (*i.e.,* the metabolome of each strain) varied significantly with culture media salinity and ecological niche (**Figure 3** and **Appendix B**). The main differences between the samples were related to the salinity of the culture media, showing two clear groupings (0% vs 17% NaCl) (**Figure 3a**). Regarding the ecological niche, less obvious groupings were observed, with the samples from the terrestrial environment only distributed within the bottom half of the scaling plot (**Figure 3b**). No clear patterns were observed with the ploidy level and the presence/absence of vHPO genes (**Figure 3c** and **3d**), indicating these two factors have no effect on the metabolite production in the culture conditions used here. While vHPO genes presence/absence may have an impact on the metabolome of the yeast, this is something that has never been described in fungi. So far, only three bacterial vHPO could be associated to the biosynthesis of metabolites, and they are the only examples in Nature (Baumgartner et al., 2024). This is why this investigation of the impact of the presence/absence of vHPO genes on the metabolome of the yeast was purely exploratory. Their expression could be implied in many other diverse physiological processes as well in modification of exogenous signals (Baumgartner et al., 2024). Moreover, the absence of halogenated metabolites could also be explained by the fact that vHPO could act as an activating enzyme. Indeed, the halogenation of some molecules would only be transient, allowing a molecule to be activated to react with other molecules and thus form chemical complexes.

**Figure 3.**
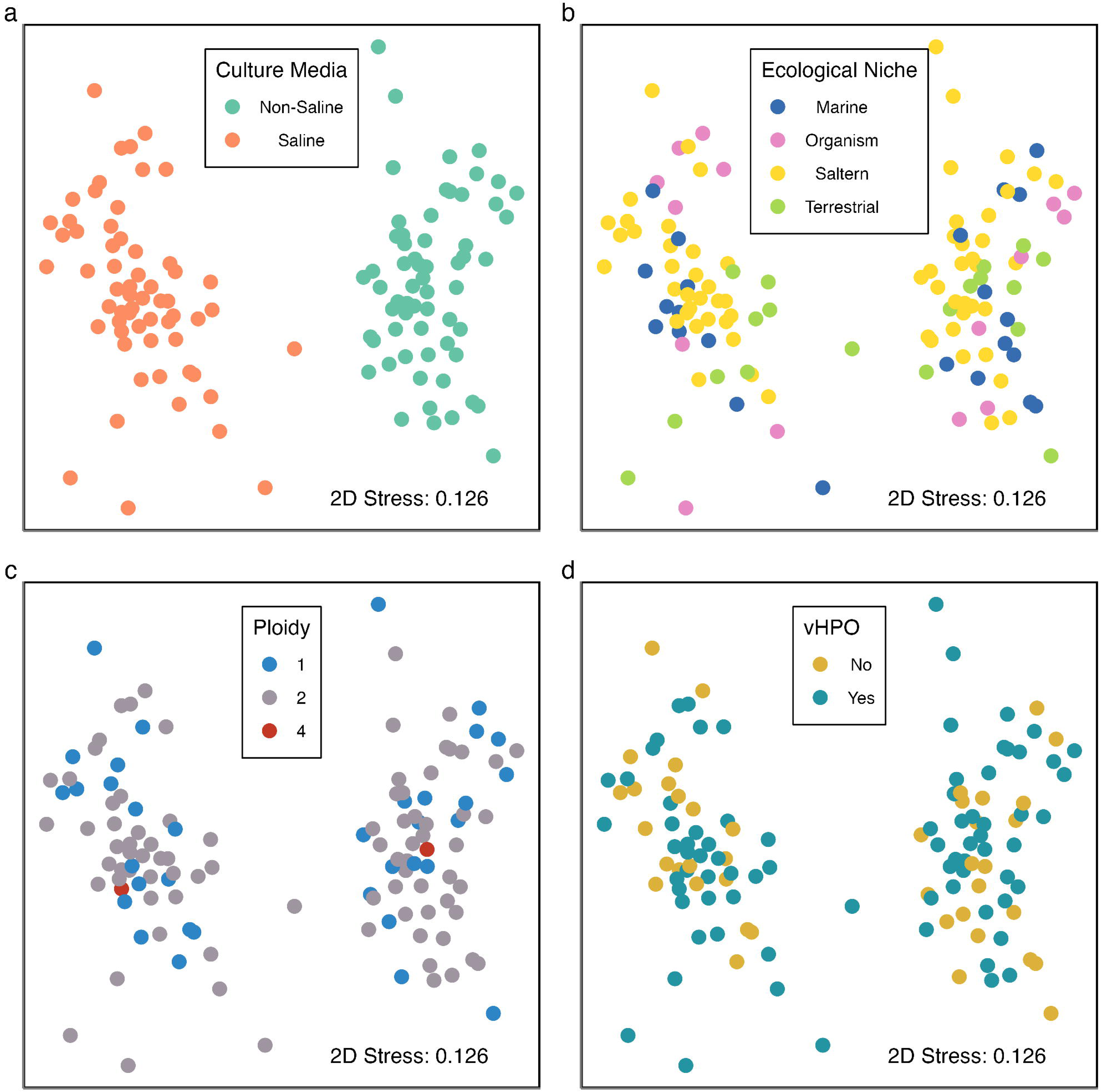
Non-metric Multidimensional Scaling plot of metabolome dissimilarity (Bray-Curtis of fourth root transformed data). Each circle represents one sample. Colors denote Culture Media Salinity (a), Ecological Niche (b), Ploidy Level (c) and Presence of vHPO Genes (d). 2D stress for this ordination was 0.126.

Consequently, in this work the absence/presence of NaCl in the culture media is the most discriminating factor in the metabolite production, regardless of the origin of the strains, indicating *H. werneckii* has a specific chemical response to the presence of salt in its environment. The slight grouping of samples observed with the ecological niche factor implied that regarding their native habitat, the strains are able to produce some specific compounds, and the latter could be part of their adaptation to this particular habitat.

A univariate General Linear Model (GLM) analysis was then carried out to describe the diversity, evenness and dominance of the features produced by the strains sampled from different ecological niches and grown on saline and non-saline media (**Figure 4**). The diversity (Shannon’s Index) is the combination between the number of features present in a sample with their evenness among the sample. The evenness (Pielou’s index) is reflected by the similarity between the areas of all the features in one sample, regardless of the number of features. By contrast, the dominance depicts the dominance of some features compared to others within a sample. Dominant features correspond to features having a greater area than others. As mass spectrometry is not a universal detection technique, the molecules associated to these features are not necessarily produced in higher amounts by the strains, but could be molecules that get ionized particularly well. Therefore, the conclusions drawn from the following results should be treated with caution as they may not fully reflect actual molecule production. To have actual quantitative information about the metabolite production, NMR metabolomics study could be conducted in addition. When considering the metabolome diversity weighted by the distribution of areas (Shannon’s Index), non-saline media samples generally had greater diversity than saline media samples, except for marine samples, where there seemed to be little difference between non-saline and saline culture media (**Figure 4a**). Conversely, the other ecological niches – salterns, terrestrial and organism-related environments – showed the same hierarchies of diversity within their culture media groupings. When assessing the number of features present in each sample, regardless of feature area, a very different pattern was found (**Figure 4b**). There seemed to be no clear difference between culture media salinity nor ecological niche alone, yet there was a large range in the number of features present, ranging from around 69 in non-saline media samples from salterns down to 55 in saline media samples from organism-related environments. Non-saline media samples from organism-related environments, alongside saline media samples from marine and terrestrial environments averaged around 65 features per sample, while non-saline media samples from marine and terrestrial environments with saline media samples from salterns were similarly diverse with 61 features per sample. For the evenness of areas found across features within the samples (Pielou’s Index), non-saline media samples generally had a greater evenness than saline media samples, but the evenness remained generally low for all the samples, with a value between 0.55 and 0.60 (**Figure 4c**). This trend was consistent with the results observed for dominance (Gini’s Coefficient), with samples from non-saline media showing a lower dominance, except for marine samples (**Figure 4d**). For the other ecological niches, saline media samples were generally greater than non-saline media samples, with organism-related samples showing the greatest dominance, followed by terrestrial and then saltern samples. Assuming dominant features correspond to molecules produced at high levels, these results showed that saline culture media generally induced the production of distinct metabolites in higher amounts, although all the samples showed a high dominance, with values above > 0.9. Regarding the potential of the strains for compound discovery, these results showed that strains grown on non-saline media are more interesting than strains grown on saline media, as they presented a higher diversity. More precisely, strains sampled from salterns and grown on non-saline media could be good candidates for the isolation of metabolites, as they showed the highest number of features and a high evenness. In particular, the strain EXF-291 (isolated from salterns in Slovenia, **Table 1**) produced the largest number of features when grown on non-saline medium (82 features). It is reported that most *H. werneckii* strains have a broad salinity optimum, from 5-10% NaCl, and an increased level of salt does not represent a stressful condition, but rather their optimal condition (Marchetta et al., 2018). Thus, the results observed here regarding the diversity could be explained by the fact that the non-saline culture medium represents a rare condition for *H. werneckii* strains isolated from salterns, as they normally grow at hypersaline conditions, thus triggering a wider production of specialized metabolites, potentially involved in stress response.

**Figure 4.**
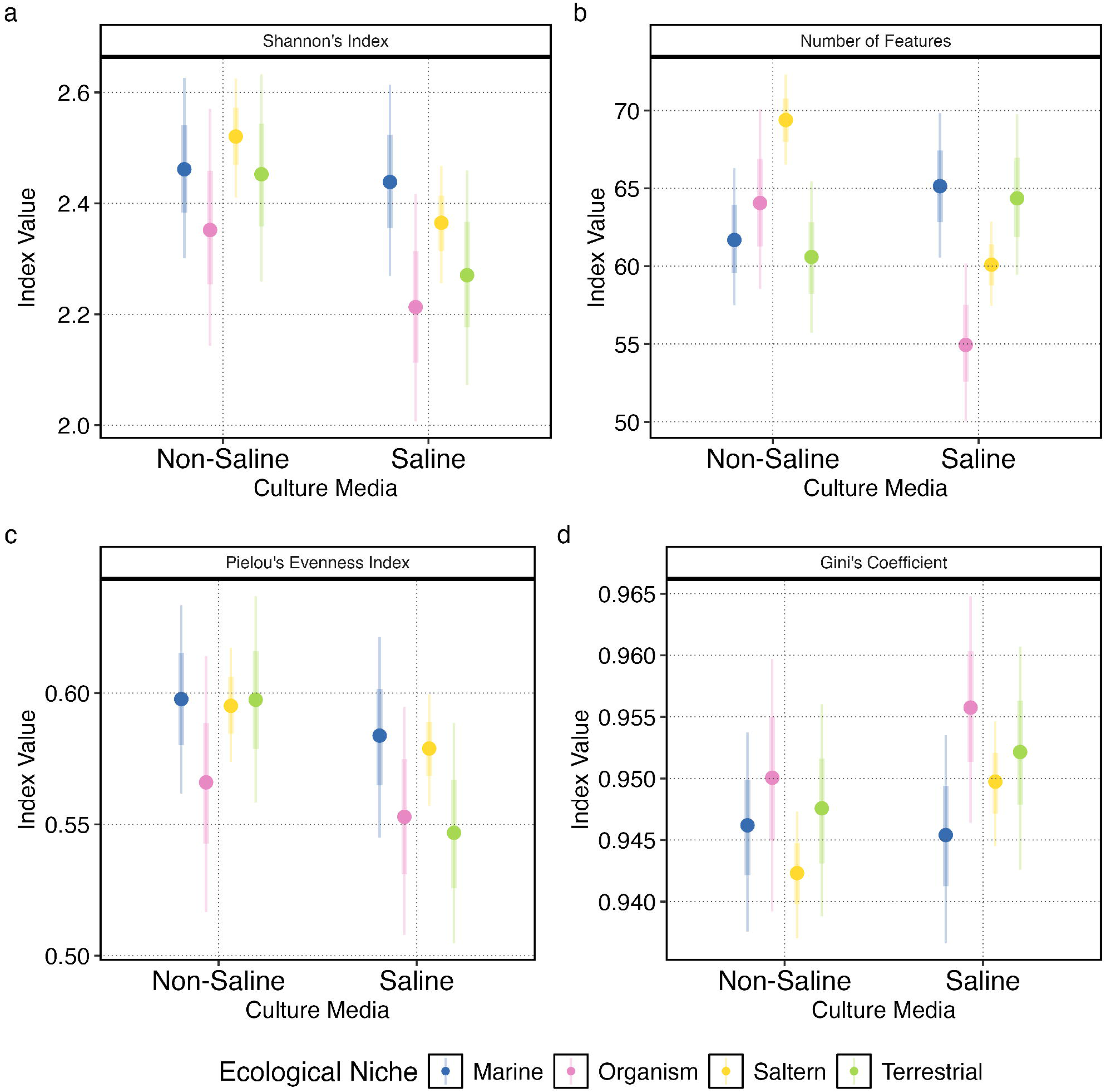
Features diversity, dominance and evenness across ecological niche and culture media salinity. Panels show Shannon’s Index (a), Number of Features (b), Pielou’s Index (c) and Gini’s Coefficient (d). Points show medians, while bars show 66 and 95 % quantiles of the posterior distributions. Colors denote ecological niche, while the x axis shows culture media salinity.

In order to highlight the most discriminating features between the samples and to find out which features were dominant within the samples, a heat map was generated (**Figure 5**). Eight features appeared to be particularly dominant: 181.0715_0.87, 347.0568_10.90, 355.1594_21.89, 381.1752_22.27, 383.1904_23.79, 405.2865_19.04, 429.2867_18.08 and 431.3021_19.60. Four of them were also discriminating, as they were only produced by strains grown on non-saline or saline culture media: 181.0715_0.87, 405.2865_19.04, 429.2867_18.08 and 431.3021_19.60. The feature 235.0622_6.23 could also be considered as a discriminating feature, as it was exclusively produced by organism-related strains (**Figure 5**).

**Figure 5.**
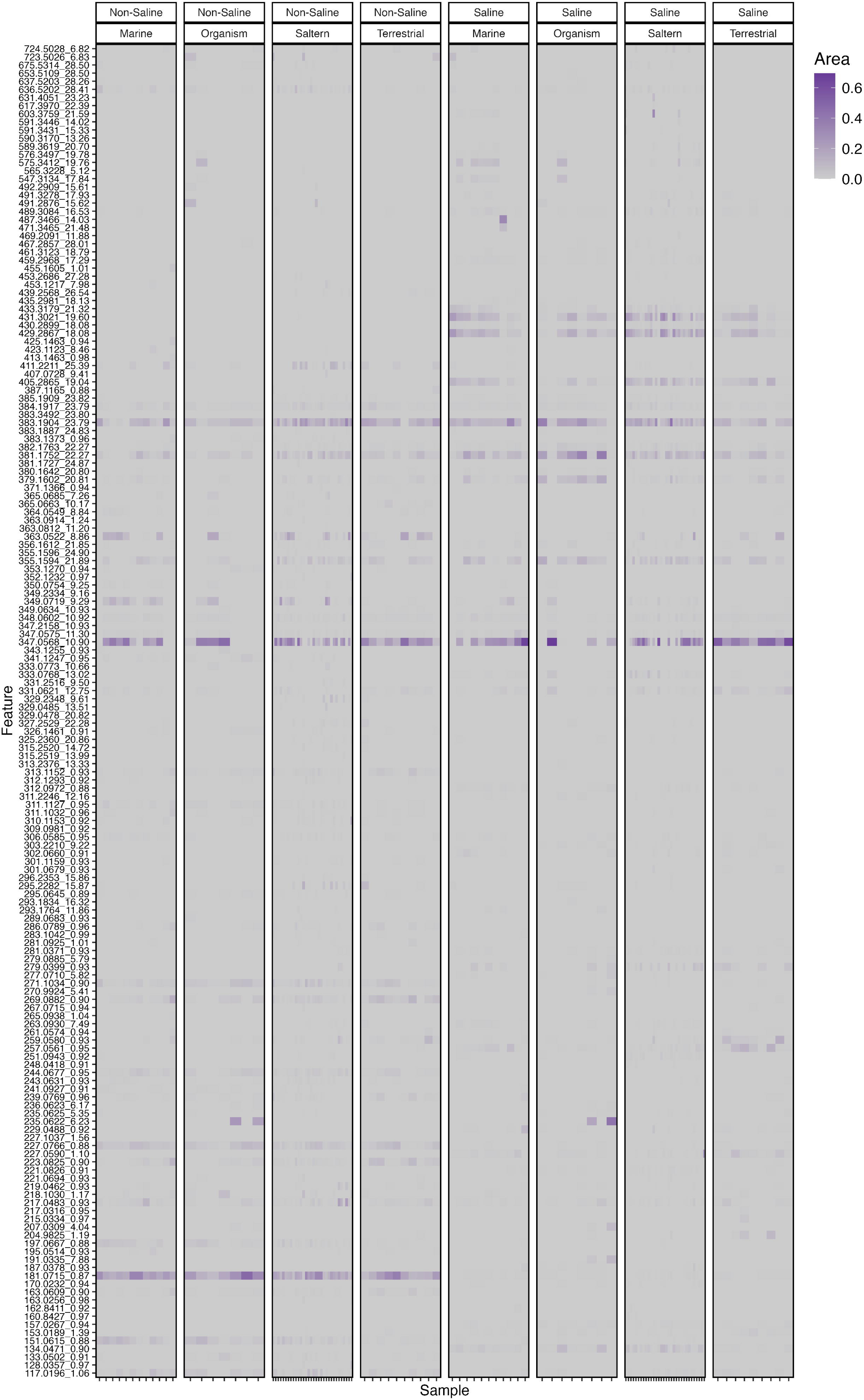
Heatmap of Fourth Root transformed areas for each feature within each sample. Panels show the culture media salinity and ecological niche.

### 3.2. Chemical diversity

To obtain information about the overall chemical diversity of the compounds produced by these 64 strains and to identify potential molecules of interest, i.e., dominant and discriminating features, a feature-based molecular network (FBMN) containing 101 nodes and three clusters (at least 3 nodes, C1 to C3) was generated (**Figure 6**). The rather small number of nodes observed here could be explained by the fact that a small number of specialized metabolites are produced in sufficient amounts in the culture conditions used here and maybe only a portion of the produced metabolites has been extracted by the solvent (EtOAc) used in this study. Moreover, some metabolites may not ionize properly in MS, or such in low intensity that they could not be fragmented. As only three clusters were observed, it seemed the metabolites didn’t share an important structural similarity and thus could belong to a wide range of chemical classes. The limited number of clusters can also be explained by the low rate of ion fragmentation observed during the HPLC-HRMS/MS analysis.

**Figure 6.**
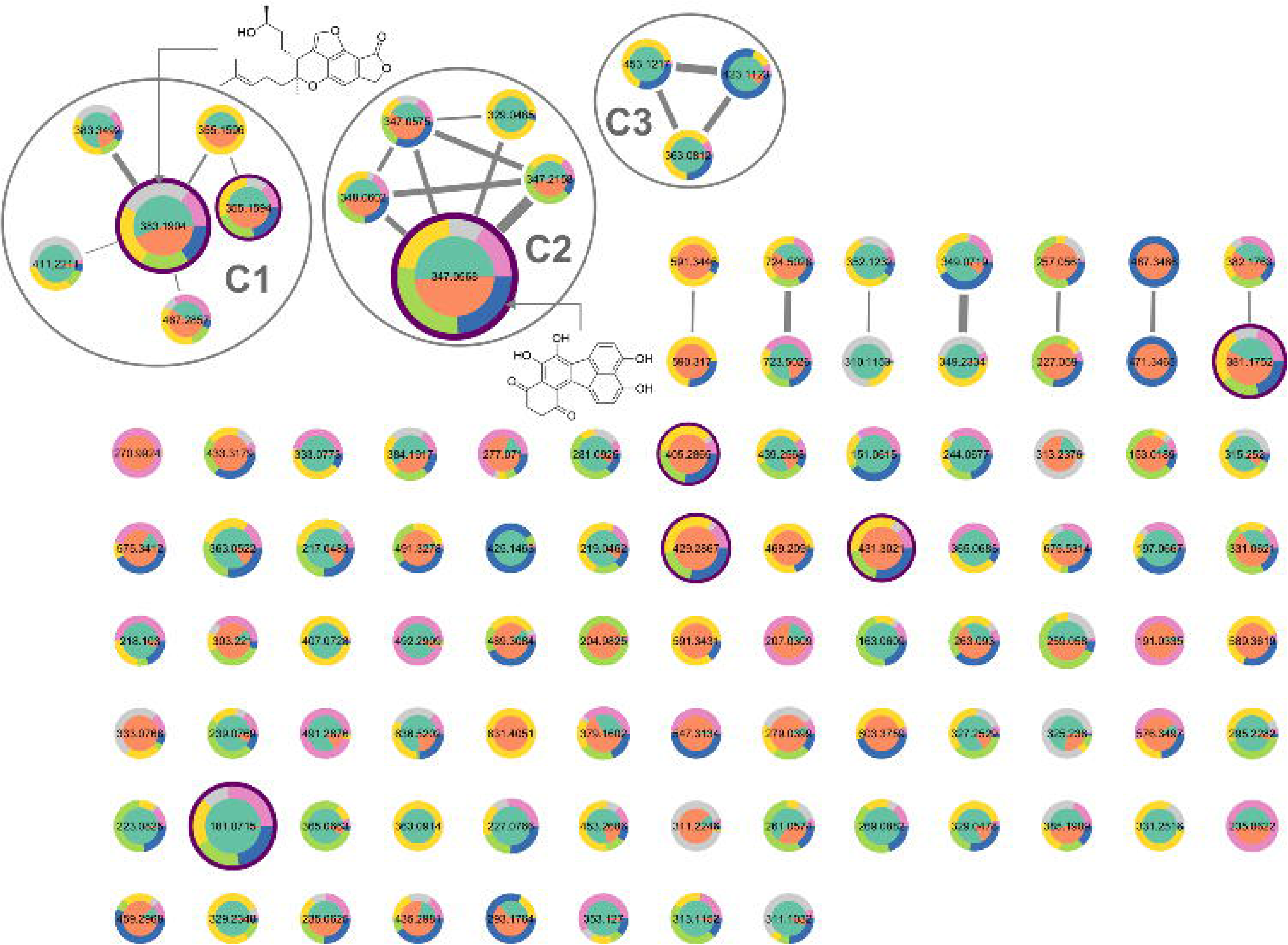
Feature-Based Molecular Network (FBMN) of the 64 samples. C1, C2 and C3 correspond to Cluster 1, 2 and 3. Each node corresponds to one feature. Node fill colors denote Culture Media Salinity (orange: saline, green: non-saline), Node border colors denote Ecological Niche (yellow: salterns, blue: marine, green: terrestrial, pink: organism-related, grey: others). Nodes circled in violet correspond to the eight dominant features identified on the Heat Map. The size of the nodes represents the ion (feature) intensity. The thickness of the edge represents the cosine score between two features (0.4 to 1).

The size of the nodes indicated four features were particularly intense: 383.1904_23.79, 381.1752_22.27, 347.0568_10.90 and 181.0715_0.87 (**Figure 6**), and these four features were previously identified as dominant features (**Figure 5**). The first three were detected independently of the ecological niche or the salinity of the culture media and thus could be considered as chemical markers of the species. The most intense feature, 347.0568_10.90, was annotated as hortein (**Table 2**), a phenolic compound isolated for the first time from *H. werneckii* (Brauers et al., 2001). In a previous study, hortein was obtained from a deep-sea hydrothermal vent strain (UBOCC-A-208029) (Burgaud et al., 2010), so comparison of its *m/z* and retention time with the feature 347.0568_10.90 allowed us to confirm this annotation (personal communication). As this feature was part of a cluster (cluster C2), this indicated molecules sharing structural similarities with hortein were produced by the strains.

**Table 2.**
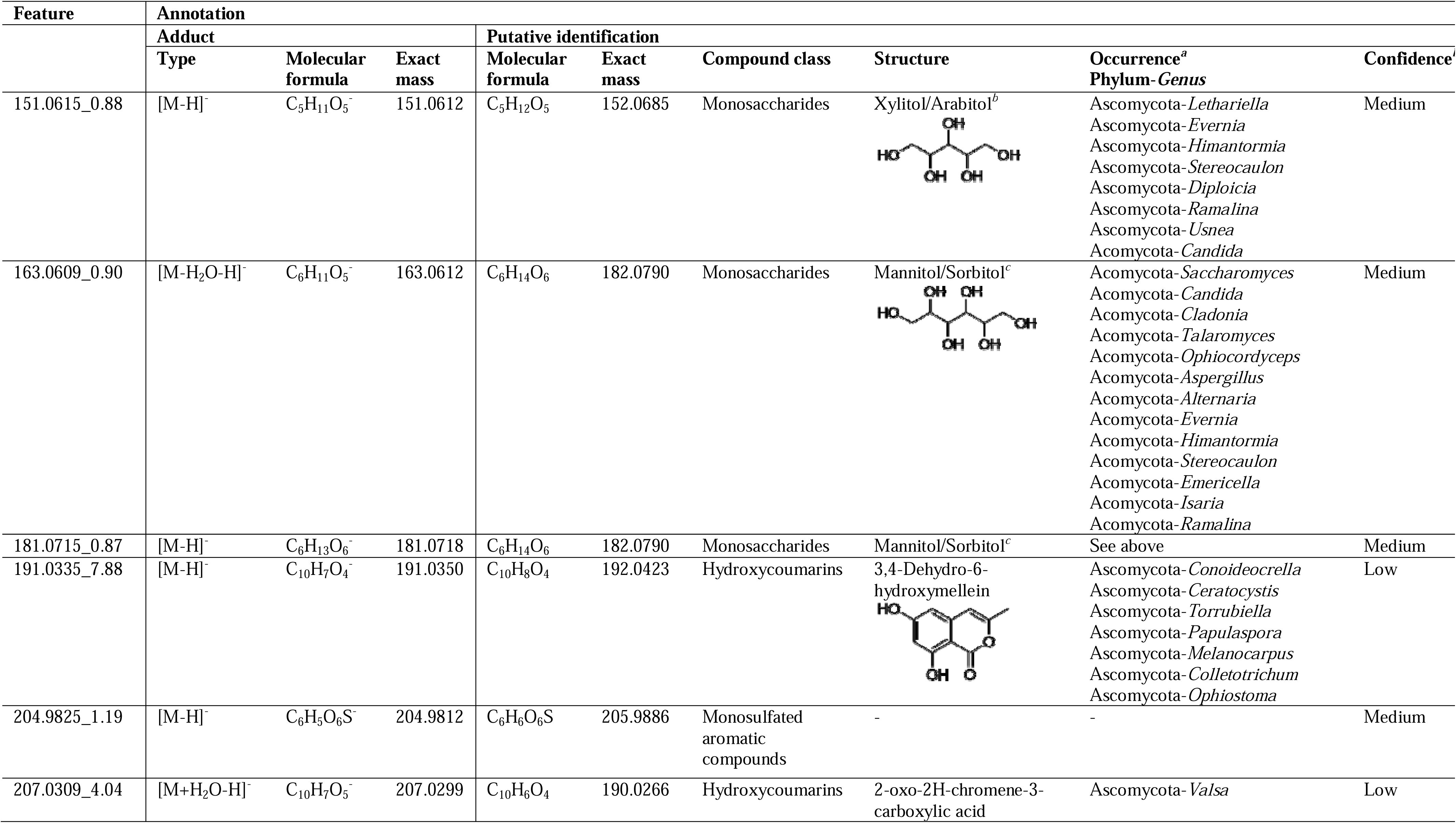
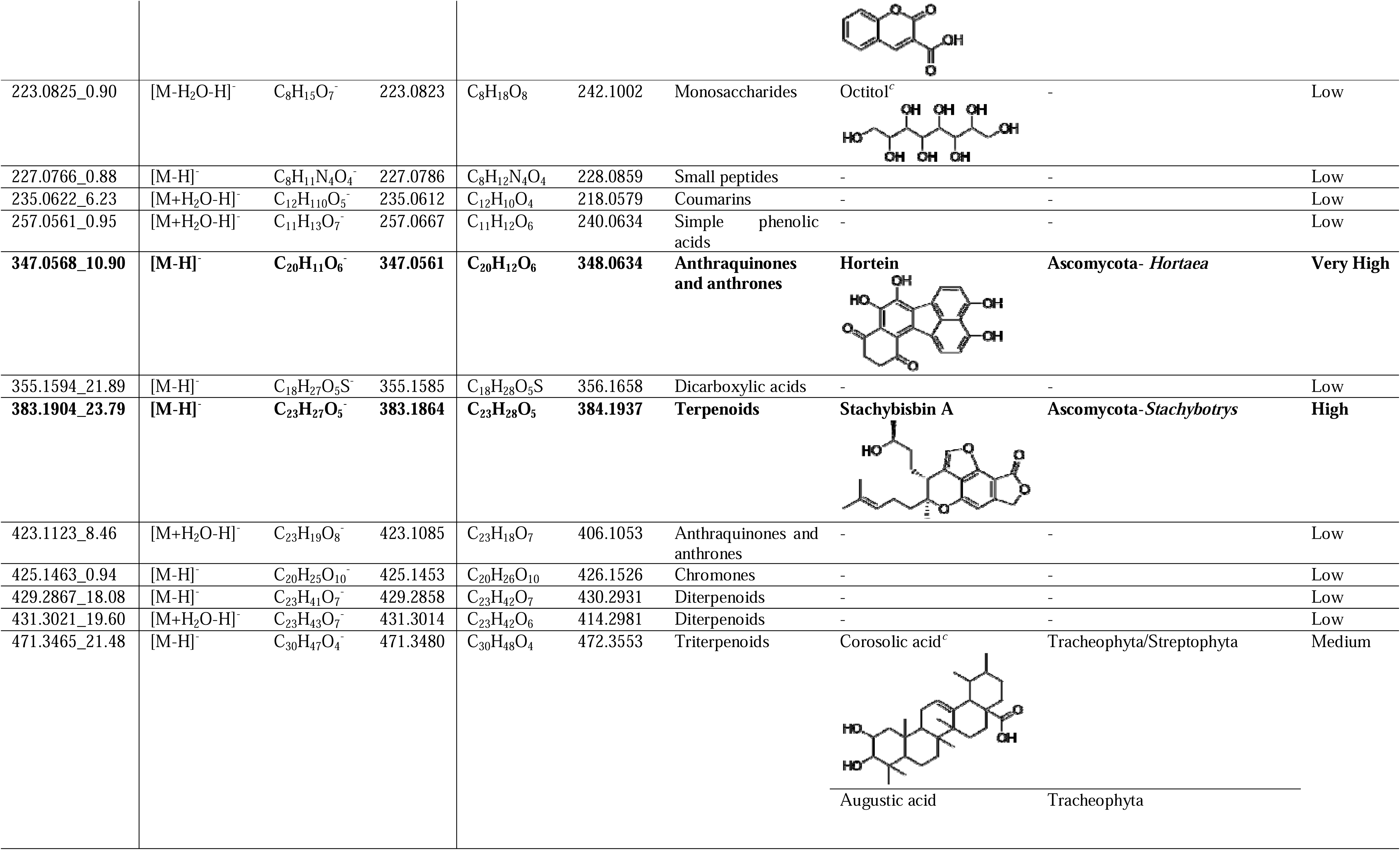
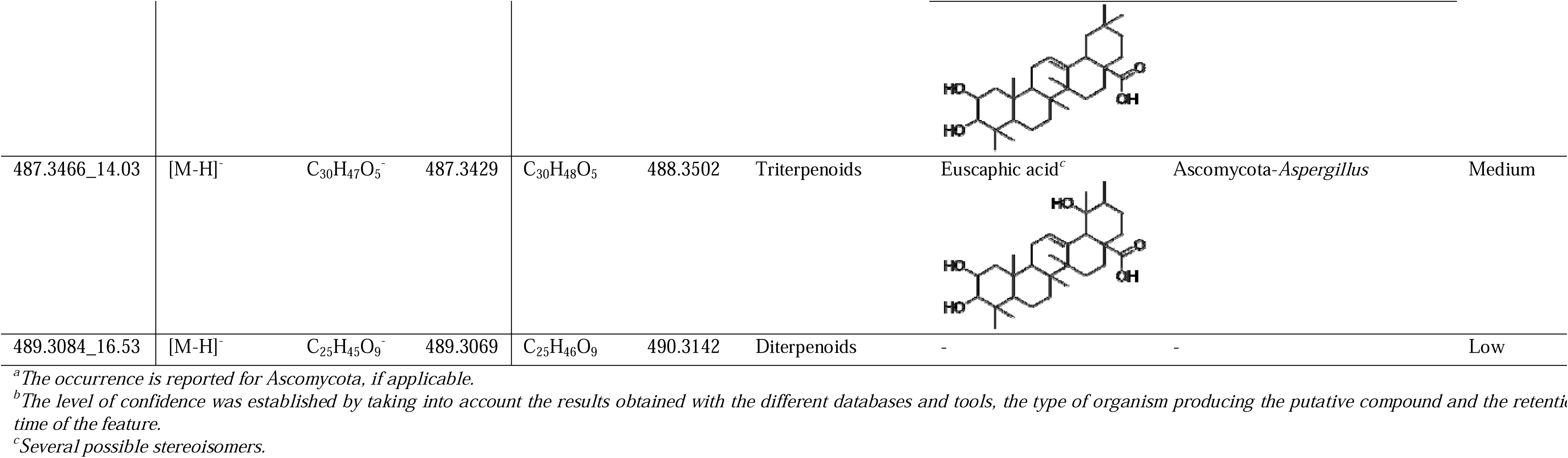
Putative annotation of the features of interest.

BGCs related to hortein biosynthesis have never been described so far. Regarding the polyketide nature of hortein, we hypothesized that the enzyme responsible for its biosynthesis could be a type I polyketide synthase (PKSI). The analysis of the 64 genomes was then performed and allowed us to reveal the presence of at least one gene coding for a PKSI in all 64 strains. We could relate for the first time these PKSI gene cluster to the biosynthesis of hortein, as a phylogenetic study showed that all these protein sequences belong to the same cluster (they branch together) (**Appendix C**), and the predicted structure of the molecule synthesized by the PKSI shared similarities with hortein (Zhang et al., 2017).

The second most intense feature, 383.1904_23.79, was putatively identified as stachybisbin A (**Table 2**), a meroterpenoid isolated from the fungus *Stachybotrys bisbyi* (Ascomycota, Sordariomycetes) (Bao et al., 2015). Five nodes were connected to this feature (cluster C1), indicating the strains probably produced meroterpenoid-like compounds. In the BGC analysis (**Figure 2** and **Appendix A**), several terpene synthases were identified. It is therefore not inconsistent to find numerous terpenoids in the strains. No annotation was obtained for the feature 381.1752_22.27. In contrast to the others, the feature 181.0715_0.87 was only detected for strains grown on non-saline medium (green node-filled color), regardless of the ecological niche. Its putative identification showed this feature probably corresponded to a monosaccharide, which was consistent with its very low retention time (0.87 min), related to xylitol or arabitol (**Table 2**). Assuming this annotation was correct, this observation was rather unexpected. Previous studies have reported that the production of polyols – primarily glycerol during the exponential phase and additional polyols such as arabitol during the stationary phase – is triggered in *H. werneckii* during saline growth conditions as a mechanism for osmoregulation (Gunde-Cimerman et al., 2018; Gunde-Cimerman and Plemenitaš, 2006). One possible hypothesis for the absence of polyols in samples from saline media is that these compounds are retained intracellularly to maintain osmotic balance, whereas at non-saline conditions, they may be excreted. Although sonication was employed during the extraction process, it is possible that the frequency and exposure time specified in the protocol were not optimal, resulting in only a partial release of intracellular compounds. Also, we performed a bath-type sonication, and it has been reported horn-type sonication was more effective in disrupting yeast cells (Zhao et al., 2024).

Even though cluster C3 contained nodes corresponding to features mainly detected in non-saline culture media (green-filled nodes) (**Figure 6**), no cluster specific of culture media salinity nor ecological niche was observed. However, several single nodes and pair of nodes associated to culture media salinity were spotted in the network, indicating the chemical response of the yeast to the absence/presence of salt in its environment does not imply a particular chemical class. Nineteen features appeared to be restricted to saline culture media (orange-filled nodes), including the three features observed previously (405.2865_19.04, 429.2867_18.08 and 431.3021_19.60, **Figure 5**), and 19 features appeared to be restricted to non-saline culture media (green-filled nodes). Although the ecological niche of the strains only induced a slight grouping among the samples (**Figure 3**), 5 features appeared to be specific to strains collected from salterns (yellow-bordered nodes), and 5 to strains from organism-related environments (pink-bordered nodes).

The search for specialized metabolite biosynthetic pathways in the genomes available made it also possible to highlight particularities in some strains, that could explain the diversity of the produced molecules. For example, a feature (631.4051_23.23) was identified exclusively in the strain EXF-561. After analysis of its genome, a hybrid TPS precursor/RiPPs cluster (Dunbar et al., 2012) and a gene encoding an enzyme that could synthesize a lassopeptide were exclusively found in this strain. Genome analysis also showed the presence of genes encoding DMATS, responsible for indole formation, in 21 *H. werneckii* strains. A phylogenic study revealed the presence of three different enzymes. Seventeen protein sequences showed 100% identity, four others clustered together (found in EXF-177, EXF-2516, EXF-2785 and EXF-12620) and one was alone (found in EXF-2515) (**Appendix D)**. They would be responsible for the synthesis of alkaloid-type molecules. However, none of the features putatively identified based on hits obtained in natural product databases (**Table 2**) matched with indole-containing structures. Further studies would then be needed to more specifically investigate the potential presence of indole alkaloids in *H. werneckii*.

### 3.3. Compounds of interest

Features identified as dominant and discriminating were defined as potential molecules of interest, as they could be involved in the ecology of the species and represent potential targets for compound discovery. In this way, a list of 58 important features was established (**Appendix E**), according to the molecular network (**Figure 6**) and the Heat Map (**Figure 5**), and these features were subjected to a multivariate GLM analysis to describe in more detail the effects of culture media salinity and ecological niche on their production. Features for which an important effect was observed are presented here. It appeared that 3 of the 4 features previously detected as dominant and ubiquitous were influenced by the culture media salinity and/or the ecological niche (**Figure 7**). Two of them were particularly influenced by the ecological niche, with the feature 347.0568_10.90 (hortein) being highly produced by terrestrial strains on saline culture media, and the feature 381.1752_22.27 (no annotation obtained) being highly produced by organism-related strains on saline culture media. For feature 355.1594_21.89 (no annotation obtained), it seemed its production was slightly enhanced on saline culture media, regardless of the ecological niche. No particular pattern was observed for feature 383.1904_23.79 (meroterpenoid).

**Figure 7.**
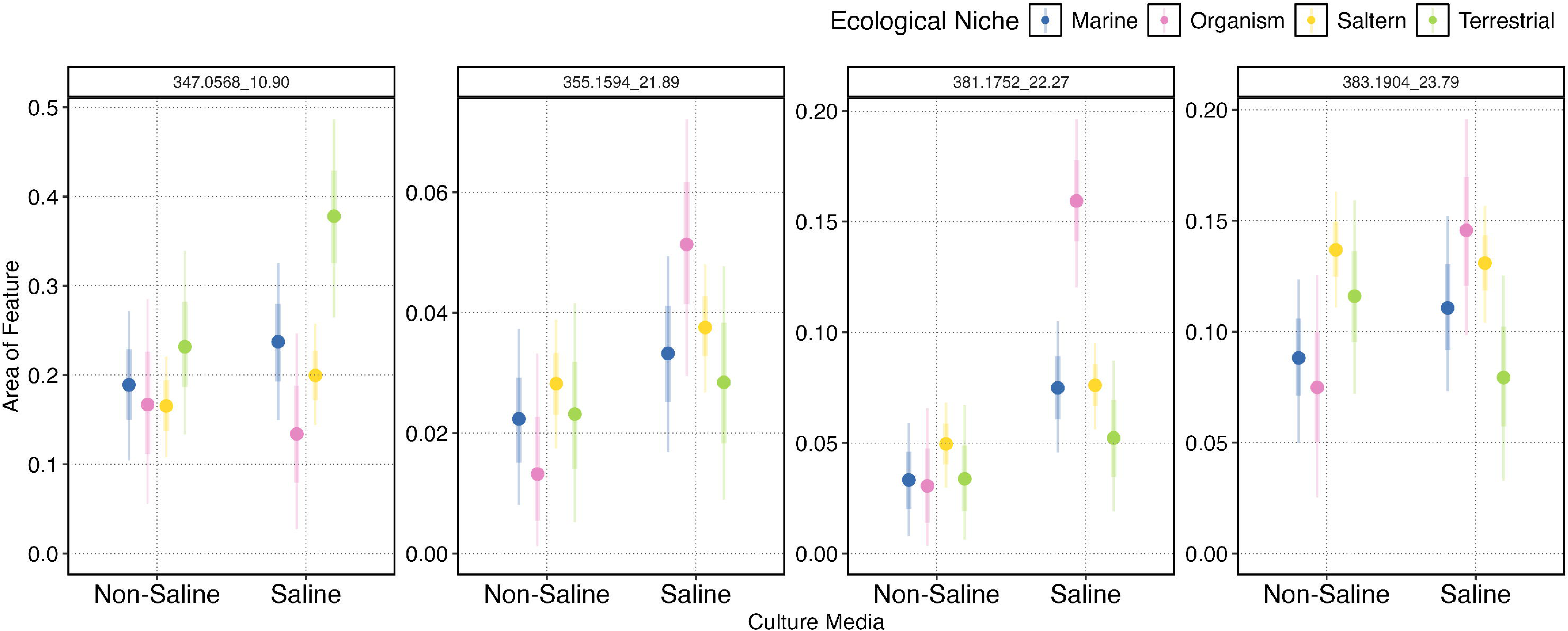
The change of ecological niche across culture media salinity for features seemingly dominant and ubiquitous across culture media salinity and ecological niche. Points show medians, while bars show 66 and 95% quantiles of the posterior distributions. Colors denote ecological niche, while panels show different features.

Eight features were particularly influenced by the culture media salinity, regardless of the ecological niche of the strains (**Figure 8**). Four features were specifically produced in samples from non-saline media, and they corresponded to polar structures (retention time < 1 min) of low molecular weight (*m/z* < 300). As mentioned above, the feature 181.0715_0.87 was putatively identify as a monosaccharide (**Table 2**). The feature 227.0766_0.88 could correspond to a small peptide, but the level of confidence of the annotation was low (**Table 2**). No annotation was obtained for the two other features. On the opposite, 4 features were specific to saline media, and corresponded mainly to apolar structures (retention time > 15 min) of medium molecular weight (*m/z* > 400). The features 429.2867_18.08 and 431.3021_19.60 could correspond to diterpenoids (**Table 2**). As they were only produced in saline conditions, these molecules could be involved in some halotolerance mechanisms. Besides polyols, it is reported that *H. werneckii* produces two mycosporines in response to salt concentration: mycosporine glutaminol glucoside is produced at high salinities, whereas mycosporine glutamicol glucoside is produced at low salinities (Gunde-Cimerman and Plemenitaš, 2006). No features corresponding to these mycosporines were detected in our analysis, which can likely be attributed to the use of EtOAc for extraction. Mycosporines are typically extracted only with polar solvents, such as water or short-chain alcohols, like methanol or ethanol (Vaz et al., 2025; Volkmann and Gorbushina, 2006).

**Figure 8.**
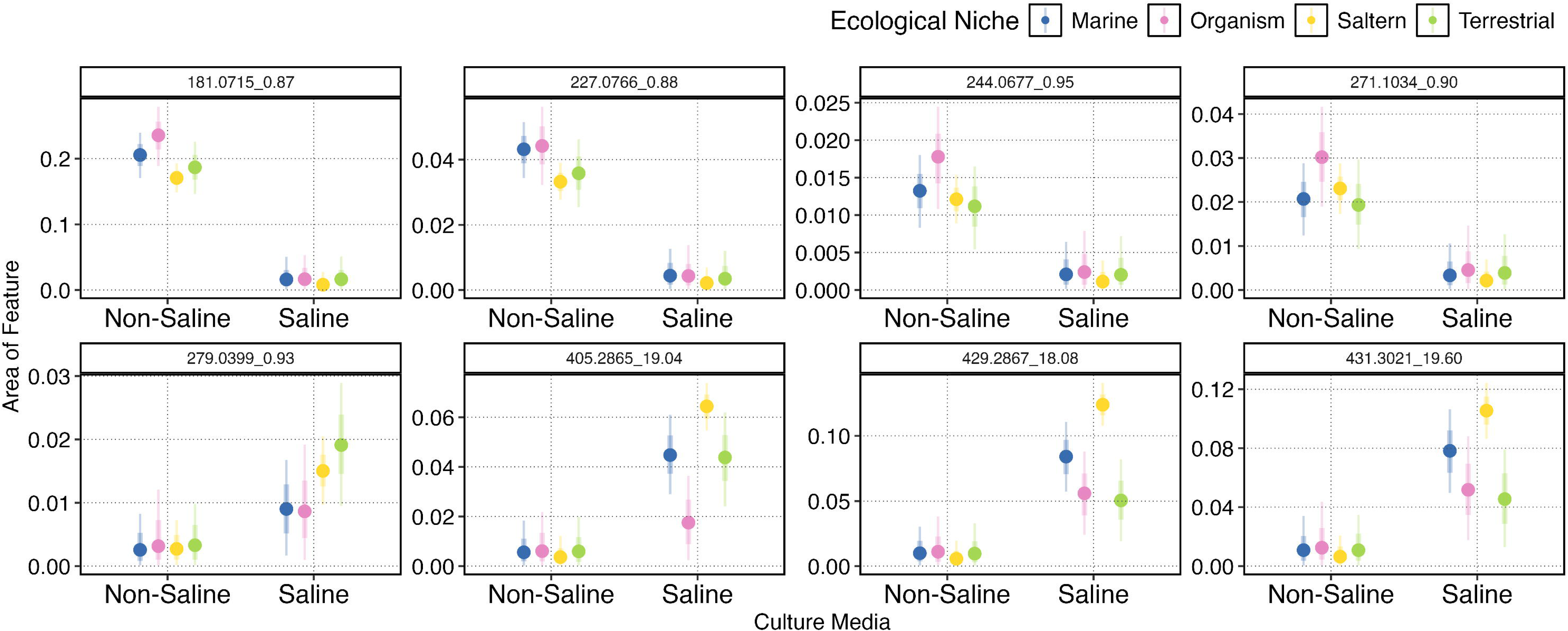
The change of ecological niche across culture media salinity for features most heavily influenced by culture media salinity. Points show medians, while bars show 66 and 95 % quantiles of the posterior distributions. Colors denote ecological niche, while panels show different features.

Twenty-four features were particularly influenced by the ecological niche of the strains (**Figure 9**), thus corroborating the above-formulated hypothesis about the production of specific compounds potentially involved in the adaptation of the strains to their habitat. A particularly important effect was observed for 7 features, which were highly produced in strains collected from organisms. We could thus hypothesize the metabolites corresponding to these features are involved in the chemical communication between the yeast and its host organism. Among these features, 235.0622_6.23 was specifically produced by organisms-related strains, regardless of the salinity of the culture media. It was detected in particularly high intensities (≥ 1E5) in the strains EXF-177 and EXF-2683 (**Appendix F**). According to the annotation, this feature could correspond to a coumarin (**Table 2**). Given that *H. werneckii* is known to cause *tinea nigra* (Zalar et al., 2019), these results suggested some compounds, including the 7 features detected in this work, could be linked to the pathogenic potential of the species. An important effect was also observed for the feature 204.9825_1.19, almost exclusively produced by terrestrial strains cultivated on saline medium (**Figure 6** and **Figure 9**). This feature was annotated as a monosulfated compound (**Table 2**). The presence of sulfotransferase genes in the genome of *H. werneckii* (Graziano et al., 2024), and more particularly in the genomes of the strains producing this feature, supports this annotation.

**Figure 9.**
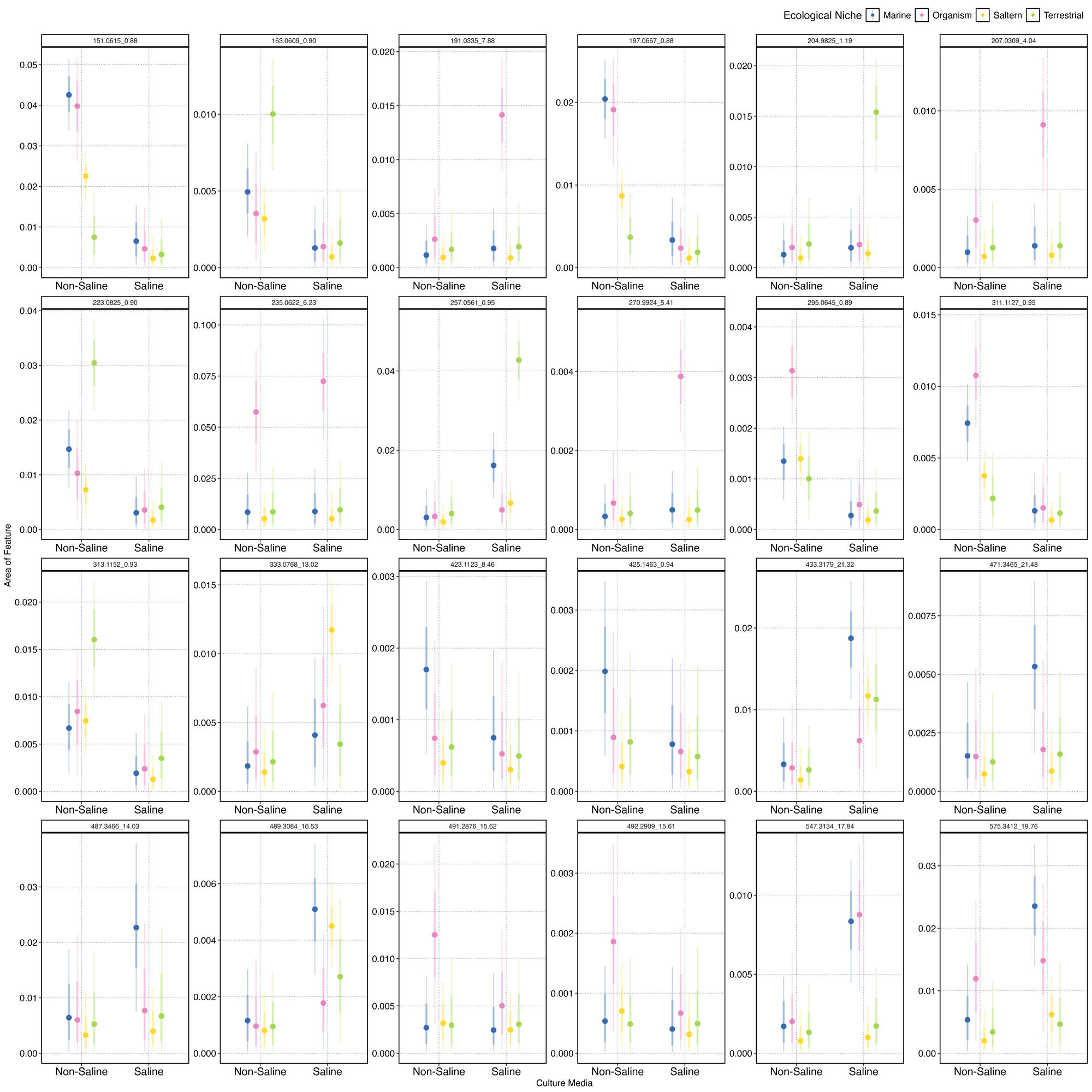
The change of ecological niche across culture media salinity for features most heavily influenced by ecological niche. Points show medians, while bars show 66 and 95 % quantiles of the posterior distributions. Colors denote ecological niche, while panels show different features.

## 4. Conclusion

A large-scale metabolomic study was conducted on the extremophile black yeast *Hortaea werneckii*, to explore its overall chemodiversity and chemical ecology, which remained poorly investigated so far, and to assess its potential for natural product discovery. The analysis of 64 strains revealed that while the number of detected metabolites was relatively limited, these belonged to diverse chemical classes, including terpenoids, coumarins, monosaccharides and hortein-like compounds. This diversity makes *H. werneckii* a species of interest for bioactive natural product isolation. Among the metabolites, hortein emerged as the most dominant feature, and was ubiquitously produced across the strains, along with three additional metabolites. This study also allowed to highlight the putative PKSI gene cluster responsible for its biosynthesis. Although the bioactivity of hortein itself has not been thoroughly investigated, several close analogues – such as viridistratin A and B, and truncatone A – have demonstrated notable cytotoxicity against cancer cell lines (Becker et al., 2020). Given that *H. werneckii* is a significant producer of hortein and structurally related compounds, this yeast should be further investigated as a potential source of new lead compounds with cytotoxic activity. Such compounds could have an ecological role and be involved in the general chemical defense of the yeast, explaining their biosynthesis.

The strains isolated from salterns and cultivated on non-saline media produced the most chemically diverse metabolomes, making them particularly interesting for future compound isolation efforts. This increased chemical diversity may be due to the absence of salt, constituting a stress condition for strains naturally adapted to hypersaline environments, thereby activating specialized metabolic pathways.

While the physiological, cellular and molecular mechanisms underlying *H. werneckii*’s adaptation to high salinities has been extensively studied (Gunde-Cimerman et al., 2018; Gunde-Cimerman and Plemenitaš, 2006), this work provides novel insights into how salinity and other ecological factors influence its specialized metabolism. Among the different factors studied, culture media salinity had the strongest effect on metabolite production, suggesting that biosynthetic pathways are up– or down-regulated in response to environmental salt concentrations. Compounds produced exclusively under saline conditions may be involved in the yeast’s halotolerance mechanisms. Interestingly, these metabolites did not share high structural similarities, indicating the salinity-dependant biosynthesis is structurally diverse. Nonetheless, polar and low molecular weight compounds tended to be associated with non-saline conditions, whereas apolar and medium molecular weight compounds were more frequently detected in saline conditions.

Although less pronounced, the ecological origin of the strains also influenced their specialized metabolomes. Some metabolites were uniquely produced by strains isolated from other organisms, suggesting that host-associated ecotypes may possess metabolic traits associated to pathogenicity. This was especially evident when individual compounds were examined, as several were exclusively found in strains of presumed pathogenic potential.

Since some *H. werneckii* strains can grow at 37 °C, and the addition of salt to culture media expands the growth range at this temperature (Zalar et al., 2019), further investigation into the combined effects of salinity and temperature on metabolite production could yield important insights, particularly for compounds potentially associated with pathogenicity.

In conclusion, this study advances our understanding of the chemical diversity of an extremophile yeast, *H. werneckii,* and its response to environmental factors. Several compounds of interest – including both dominant/ubiquitous molecules and metabolites produced under specific conditions – have been detected, but remained largely structurally uncharacterized. Therefore, future studies should focus on the isolation and structural elucidation of these metabolites, as they hold promise for elucidating the chemical ecology and the pathogenic mechanisms of *H. werneckii*, and they may serve as valuable sources of novel bioactive compounds. On the other hand, future works could also be performed according to the OSMAC strategy, in order to evaluate the impact of modulating other factors (*e.g*., type of culture medium, incubation temperature) on the biosynthetic pathways in the yeast, especially as some interesting BGCs highlighted in this work could not be associated to their metabolites.

## Supporting information

Supp data with sequences for Indole/DMATS

Supp data with sequences for PKSI

## Author Contribution

**Elise Gerometta:** Formal analysis, Investigation, Validation, Visualization, Writing – original draft, Writing – review & editing. **Bede Ffinian Rowe Davies:** Formal analysis, Resources, Software, Validation, Visualization, Writing – original draft, Writing – review & editing. **Rafia Ahmed Tuli:** Formal analysis, Investigation, Writing – review & editing. **Bastien Cochereau:** Investigation, Writing – review & editing. **Annie Lebreton-Nicaise:** Formal analysis, Investigation. **Laurence Meslet-Cladière:** Resources, Investigation, Writing – review & editing. **Monika Kos:** Investigation, Resources, Writing – review & editing. **Nina Gunde-Cimerman:** Conceptualization, Funding acquisition, Methodology, Resources, Supervision, Writing – review & editing**. Catherine Roullier:** Conceptualization, Funding acquisition, Methodology, Project administration, Supervision, Validation, Visualization, Writing – review & editing.

## Acknowledgments

The authors acknowledge the following funding sources: the French National Research Agency (ANR-21-CE44-0003 HALO-CAT) and the Slovenian Research Agency (MRIC UL, I0-0022), programs P4-0432 and P1-0198.

## Data Availability Statement

Data related to the molecular network have been submitted to a public database and the accession link has been mentioned in the manuscript. Data related to the strains are freely available on the National Library of Medicine (ncbi.nlm.nih.gov).

---

**Appendix A.** All BGC genes found in the 64 H. werneckii strains. DMATS: DiMethylAllyl Tryptophan Synthase, NRPS: Non-Ribosomal Peptide Synthetase, PKS: PolyKetide Synthase, TS: Terpene Synthase, TS-like: polyprenyl synthase synthesizing precursor substrates for PTS, Hybr: NRPS-PKS or NRPS-TS Hybrid, RiPPS: post-translationally modified peptides.

| Strain | DMATS | NRPS | NRPS-Like | PKS | TS | TS-like | Hybr | RiPPs |
| --- | --- | --- | --- | --- | --- | --- | --- | --- |
| **Salterns** |  |  |  |  |  |  |  |  |
| EXF-9 | 0 | 3 | 7 | 1 | 5 | 2 | 0 | 0 |
| EXF-12 | 1 | 3 | 8 | 1 | 6 | 2 | 0 | 0 |
| EXF-15 | 0 | 3 | 8 | 1 | 5 | 2 | 0 | 0 |
| EXF-20 | 0 | 5 | 11 | 1 | 7 | 2 | 0 | 0 |
| EXF-120 | 1 | 5 | 14 | 2 | 9 | 3 | 0 | 0 |
| EXF-152 | 1 | 5 | 9 | 1 | 5 | 1 | 0 | 0 |
| EXF-153 | 1 | 3 | 11 | 1 | 5 | 1 | 0 | 0 |
| EXF-154 | 1 | 3 | 10 | 1 | 6 | 1 | 0 | 0 |
| EXF-225 | 1 | 5 | 7 | 1 | 3 | 1 | 0 | 0 |
| EXF-241 | 0 | 4 | 9 | 1 | 5 | 1 | 0 | 0 |
| EXF-269 | 0 | 6 | 12 | 3 | 9 | 2 | 0 | 0 |
| EXF-291 | 1 | 3 | 10 | 1 | 3 | 2 | 0 | 0 |
| EXF-561 | 0 | 3 | 7 | 1 | 4 | 3* | 1 | 1 |
| EXF-2516 | 1 | 3 | 10* | 3* | 4 | 2 | 2 | 0 |
| EXF-2685 | 0 | 3 | 8 | 1 | 5 | 2 | 0 | 0 |
| EXF-2783 | 0 | 3 | 8 | 1 | 5 | 2 | 0 | 0 |
| EXF-2785 | 1 | 0 | 10* | 3* | 5 | 1 | 1 | 0 |
| EXF-2788 | 0 | 2 | 8 | 1 | 5 | 2 | 0 | 0 |
| EXF-3845 | 0 | 2 | 8 | 2 | 5 | 2 | 0 | 0 |
| EXF-3846 | 0 | 3 | 8 | 1 | 5 | 2 | 0 | 0 |
| EXF-4716 | 1 | 5 | 10 | 1 | 7 | 1 | 0 | 0 |
| EXF-6274 | 0 | 2 | 10 | 1 | 2 | 1 | 0 | 0 |
| EXF-8170 | 0 | 3 | 14 | 1 | 10 | 4 | 0 | 0 |
| EXF-10304 | 0 | 0 | 7 | 1 | 4 | 1 | 0 | 0 |
| EXF-10816 | 1 | 5 | 11 | 1 | 6 | 1 | 0 | 0 |
| EXF-10819 | 1 | 5 | 8 | 1 | 7 | 1 | 0 | 0 |
| EXF-10820 | 0 | 3 | 8 | 1 | 5 | 2 | 0 | 0 |
| EXF-10843 | 1 | 5 | 11 | 1 | 6 | 1 | 0 | 0 |
| EXF-10904 | 1 | 5 | 10 | 1 | 6 | 1 | 0 | 0 |
| EXF-10907 | 1 | 5 | 6 | 1 | 7 | 1 | 0 | 0 |
| EXF-10919 | 1 | 5 | 10 | 1 | 5 | 1 | 0 | 0 |
| EXF-10958 | 0 | 3 | 8 | 1 | 5 | 2 | 0 | 0 |
| EXF-10974 | 0 | 2 | 9 | 1 | 4 | 1 | 0 | 0 |
| **Marine** |  |  |  |  |  |  |  |  |
| EXF-166 | 0 | 5 | 11 | 1 | 11 | 3 | 0 | 0 |
| EXF-10508 | 0 | 3 | 6 | 1 | 5 | 0 | 0 | 0 |
| EXF-10509 | 0 | 1 | 7 | 2 | 2 | 1 | 0 | 0 |
| EXF-10510 | 0 | 1 | 8 | 1 | 2 | 1 | 0 | 0 |
| EXF-10511 | 0 | 3 | 6 | 1 | 2 | 0 | 0 | 0 |
| EXF-10512 | 1 | 3 | 7 | 1 | 4 | 1 | 0 | 0 |
| EXF-10513 | 0 | 3 | 4 | 1 | 7 | 2 | 0 | 0 |
| EXF-12619 | 0 | 3 | 7 | 2 | 6 | 2 | 0 | 0 |
| EXF-12620 | 1 | 2 | 9* | 2* | 4 | 0 | 1 | 0 |
| EXF-14590 | 1 | 4 | 9 | 2 | 5 | 1 | 0 | 0 |
| EXF-14591 | 0 | 6 | 12 | 3 | 10 | 3 | 0 | 0 |
| EXF-14592 | 0 | 3 | 8 | 2 | 5 | 2 | 0 | 0 |
| **Terrestrial** |  |  |  |  |  |  |  |  |
| EXF-562 | 0 | 2 | 7 | 1 | 5 | 2 | 0 | 0 |
| EXF-6651 | 0 | 2 | 9 | 1 | 4 | 1 | 0 | 0 |
| EXF-6652 | 0 | 2 | 7 | 1 | 3 | 1 | 0 | 0 |
| EXF-6654 | 0 | 5 | 8 | 1 | 6 | 1 | 0 | 0 |
| EXF-6656 | 0 | 3 | 7 | 1 | 8 | 1 | 0 | 0 |
| EXF-6663 | 0 | 3 | 7 | 1 | 5 | 2 | 0 | 0 |
| EXF-6669 | 0 | 3 | 9 | 1 | 5 | 2 | 0 | 0 |
| EXF-11540 | 0 | 2 | 10 | 1 | 2 | 0 | 0 | 0 |
| EXF-11650 | 0 | 1 | 11 | 1 | 1 | 1 | 0 | 0 |
| EXF-11651 | 0 | 3 | 7 | 1 | 6 | 1 | 0 | 0 |
| **Organism related** |  |  |  |  |  |  |  |  |
| EXF-151 | 0 | 4 | 7 | 3 | 5 | 1 | 0 | 0 |
| EXF-156 | 0 | 5 | 7 | 2 | 7 | 3 | 0 | 0 |
| EXF-157 | 0 | 3 | 8 | 2 | 5 | 2 | 0 | 0 |
| EXF-161 | 0 | 3 | 8 | 1 | 3 | 0 | 0 | 0 |
| EXF-171 | 0 | 5 | 12 | 2 | 8 | 4 | 0 | 0 |
| EXF-177 | 1 | 3 | 10* | 3* | 5 | 4 | 2 | 0 |
| EXF-2682 | 0 | 5 | 12 | 2 | 9 | 4 | 0 | 0 |
| EXF-2683 | 0 | 5 | 11 | 1 | 6 | 2 | 0 | 0 |
| **Other** |  |  |  |  |  |  |  |  |
| EXF-8422 | 1 | 4 | 11 | 1 | 7 | 1 | 0 | 0 |

\*one of the identified gene belongs to the hybrid cluster

**Appendix B.**
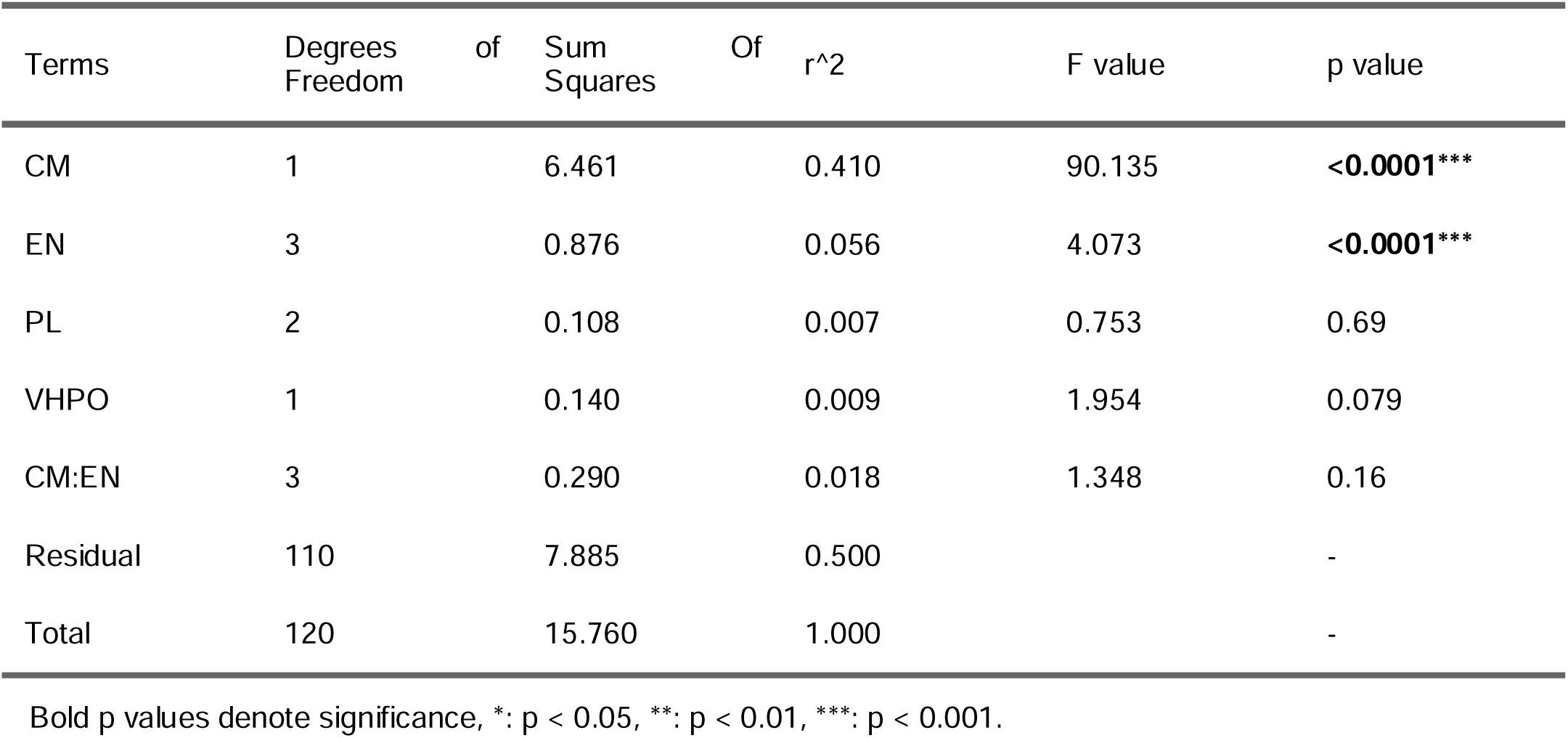
PERMANOVA analysis on Bray-Curtis dissimilarity assessing metabolomes change with Ecological Niche and Culture Media Salinity as interactions as well as Ploidy level and Presence of vHPO genes. *Culture Media Salinity, Ecological Niche, Ploidy level and Presence of vHPO genes are abbreviated to CM, EN, PL and VHPO, respectively*.

**Appendix C.**
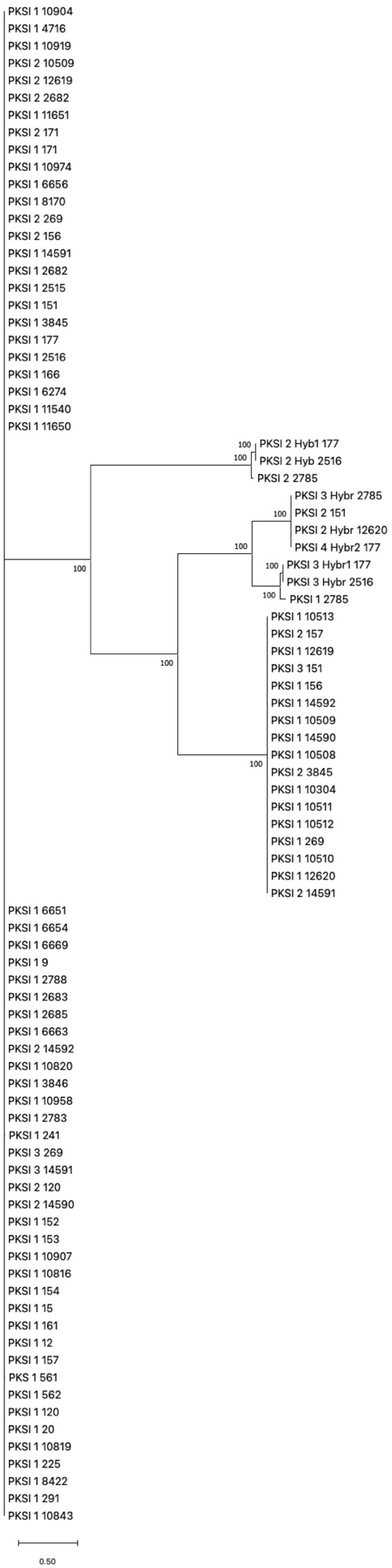
Unrooted phylogenetic tree of PKS I found in *H. werneckii* strains. *The phylogenetic tree presented here was constructed using the maximum likelihood approach. Numbers indicate the bootstrap values in the maximum likelihood analysis. The full listing of the aligned proteins is reported in Supplemental Data Set 1. The number is to reference of the number of each strain*.

**Appendix D.**
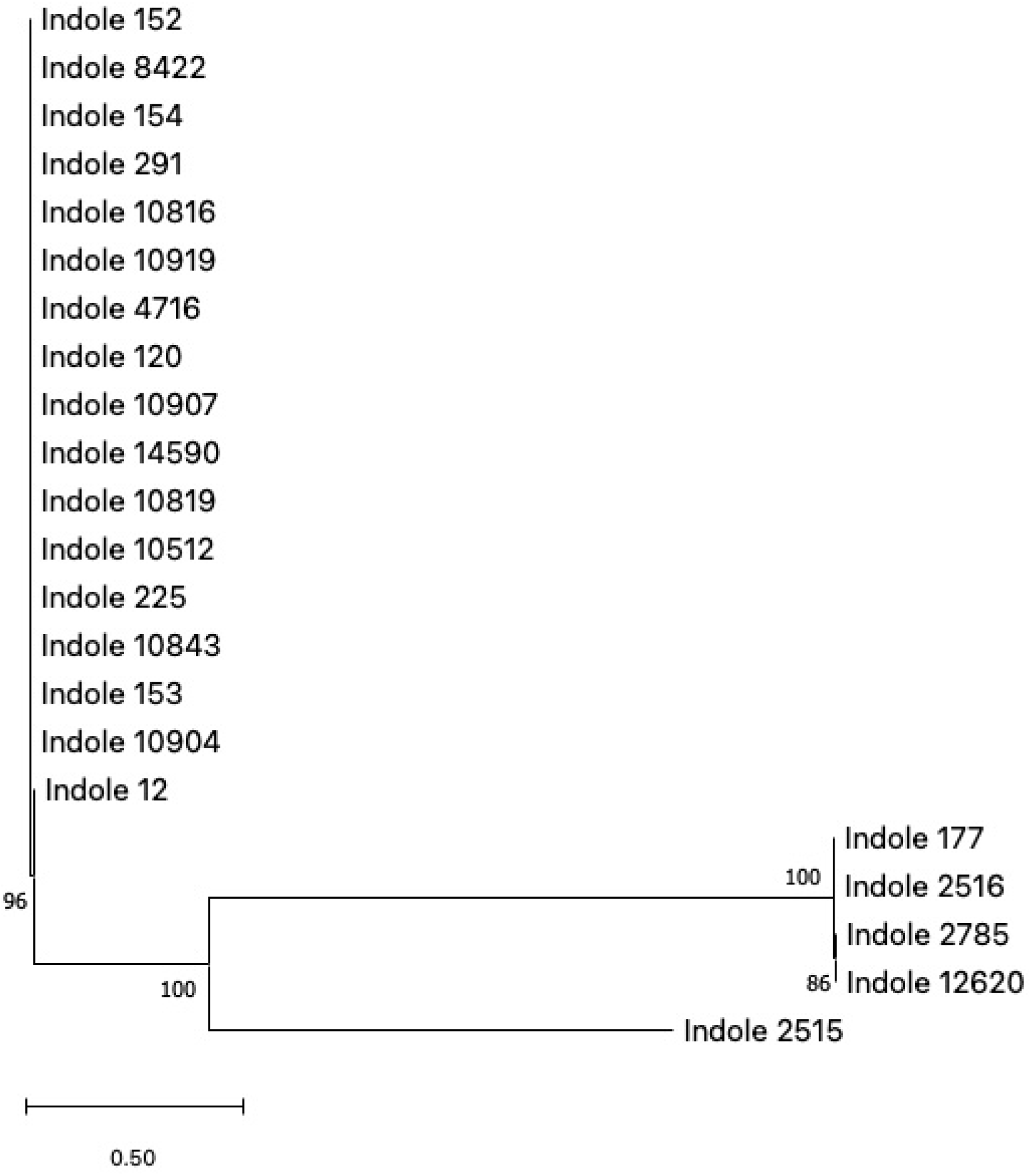
Unrooted phylogenetic tree of Indole (DMATS) found in *H. werneckii* strains. *The phylogenetic tree presented here was constructed using the maximum likelihood approach. Numbers indicate the bootstrap values in the maximum likelihood analysis. The full listing of the aligned proteins is reported in Supplemental Data Set 2. The number is to reference of the number of each strain*.

**Appendix E.** The 58 features of interest (discriminating, most intense and ubiquitous features).

| Feature | Category | Discriminating factor |
| --- | --- | --- |
| 151.0615_0.88 | Discriminating | Culture media salinity |
| 163.0609_0.90 | Discriminating | Culture media salinity |
| 181.0715_0.87 | Most intense/Discriminating | Culture media salinity |
| 191.0335_7.88 | Discriminating | Ecological niche |
| 197.0667_0.88 | Discriminating | Culture media salinity |
| 204.9825_1.19 | Discriminating | Culture media salinity/Ecological niche |
| 207.0309_4.04 | Discriminating | Ecological niche |
| 223.0825_0.90 | Discriminating | Culture media salinity |
| 227.059_1.10 | Discriminating | Culture media salinity |
| 227.0766_0.88 | Discriminating | Culture media salinity |
| 235.0622_6.23 | Discriminating | Ecological niche |
| 244.0677_0.95 | Discriminating | Culture media salinity |
| 257.0561_0.95 | Discriminating | Culture media salinity |
| 270.9924_5.41 | Discriminating | Culture media salinity/Ecological niche |
| 271.1034_0.90 | Discriminating | Culture media salinity |
| 279.0399_0.93 | Discriminating | Culture media salinity |
| 295.0645_0.89 | Discriminating | Culture media salinity |
| 310.1153_0.92 | Discriminating | Culture media salinity |
| 311.1127_0.95 | Discriminating | Culture media salinity |
| 313.1152_0.93 | Discriminating | Culture media salinity |
| 331.2516_9.50 | Discriminating | Culture media salinity |
| 333.0768_13.02 | Discriminating | Culture media salinity |
| 333.0773_10.66 | Discriminating | Culture media salinity |
| 347.0568_10.90 | Most intense/ubiquitous | - |
| 349.0719_9.29 | Discriminating | Culture media salinity |
| 349.2334_9.16 | Discriminating | Culture media salinity |
| 352.1232_0.97 | Discriminating | Culture media salinity |
| 355.1594_21.89 | Most intense/ubiquitous | - |
| 355.1596_24.90 | Discriminating | Ecological niche |
| 363.0522_8.86 | Discriminating | Culture media salinity |
| 363.0812_11.20 | Discriminating | Culture media salinity |
| 363.0914_1.24 | Discriminating | Culture media salinity/Ecological niche |
| 365.0685_7.26 | Discriminating | Culture media salinity |
| 381.1752_22.27 | Most intense/ubiquitous | - |
| 383.1904_23.79 | Most intense/ubiquitous | - |
| 405.2865_19.04 | Most intense/Discriminating | Culture media salinity |
| 407.0728_9.41 | Discriminating | Culture media salinity |
| 423.1123_8.46 | Discriminating | Culture media salinity |
| 425.1463_0.94 | Discriminating | Culture media salinity/Ecological niche |
| 429.2867_18.08 | Most intense/Discriminating | Culture media salinity |
| 431.3021_19.60 | Most intense/Discriminating | Culture media salinity |
| 433.3179_21.32 | Discriminating | Culture media salinity |
| 453.1217_7.98 | Discriminating | Culture media salinity |
| 469.2091_11.88 | Discriminating | Culture media salinity |
| 471.3465_21.48 | Discriminating | Culture media salinity/Ecological niche |
| 487.3466_14.03 | Discriminating | Culture media salinity/Ecological niche |
| 489.3084_16.53 | Discriminating | Culture media salinity |
| 491.2876_15.62 | Discriminating | Ecological niche |
| 492.2909_15.61 | Discriminating | Culture media salinity |
| 547.3134_17.84 | Discriminating | Culture media salinity |
| 575.3412_19.76 | Discriminating | Culture media salinity |
| 576.3497_19.78 | Discriminating | Culture media salinity |
| 589.3619_20.71 | Discriminating | Culture media salinity |
| 590.317_13.26 | Discriminating | Culture media salinity |
| 591.3431_15.33 | Discriminating | Culture media salinity |
| 591.3446_14.02 | Discriminating | Culture media salinity |
| 603.3759_21.59 | Discriminating | Culture media salinity |
| 631.4051_23.23 | Discriminating | Culture media salinity/Ecological niche |

**Appendix F.**
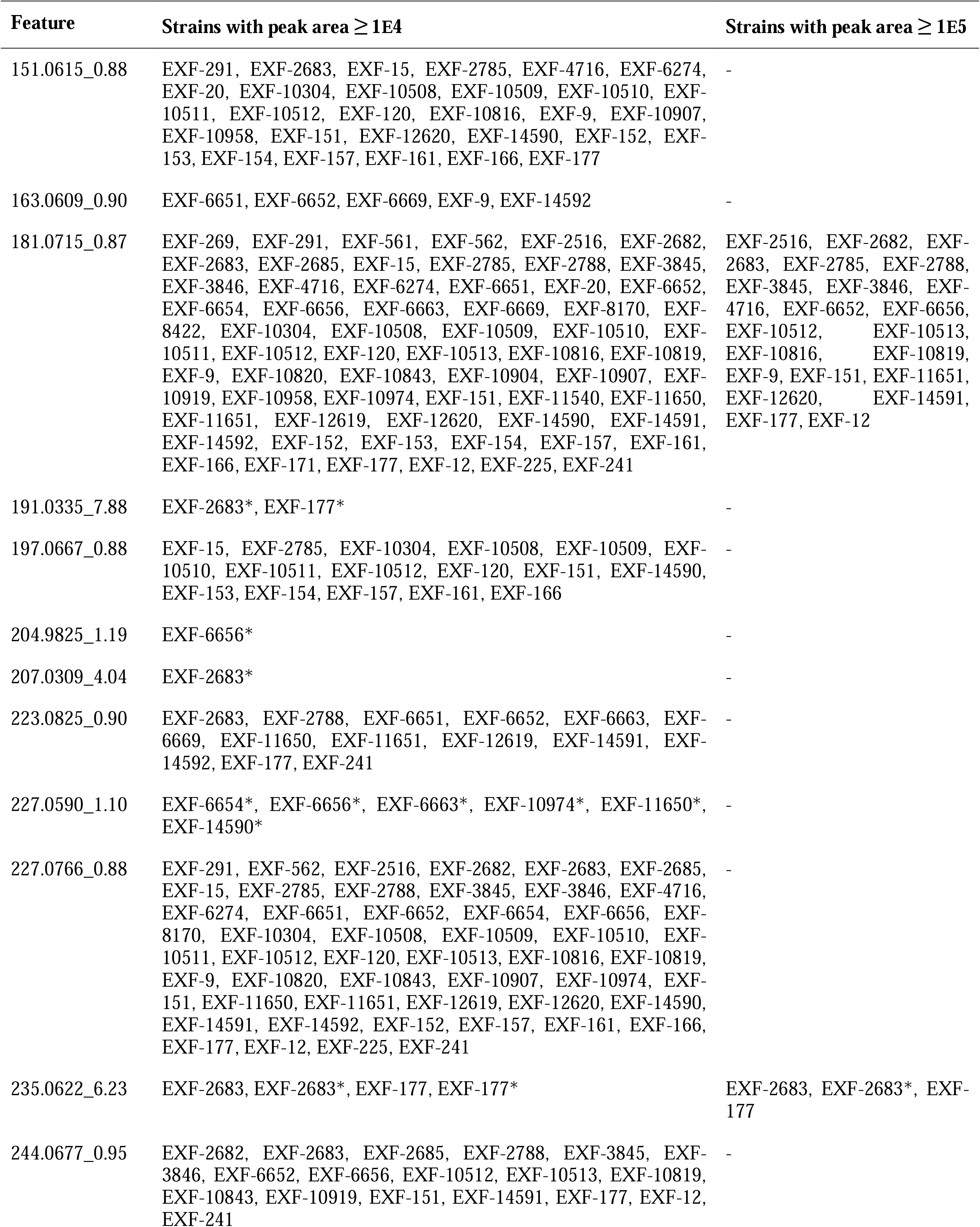
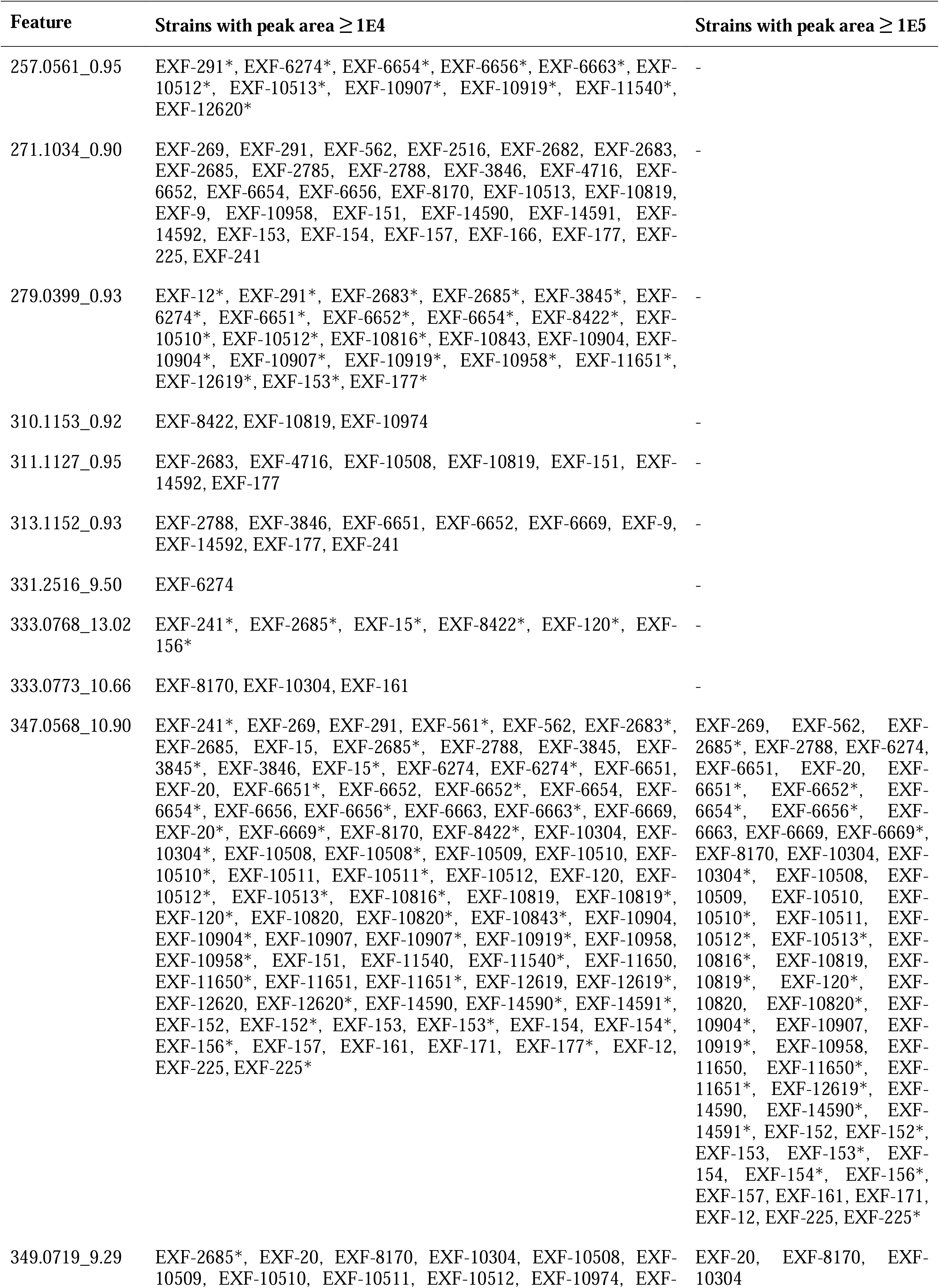
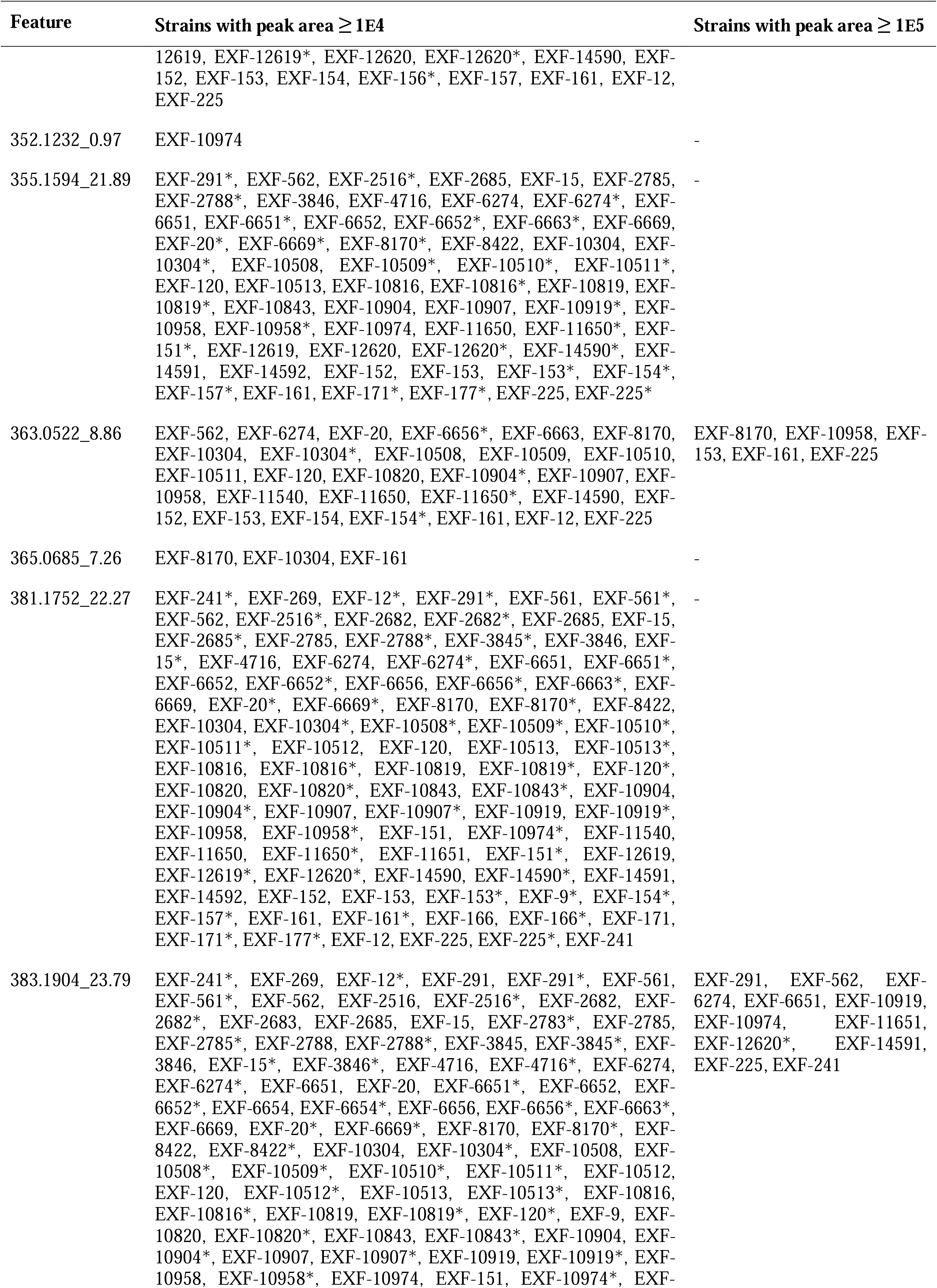
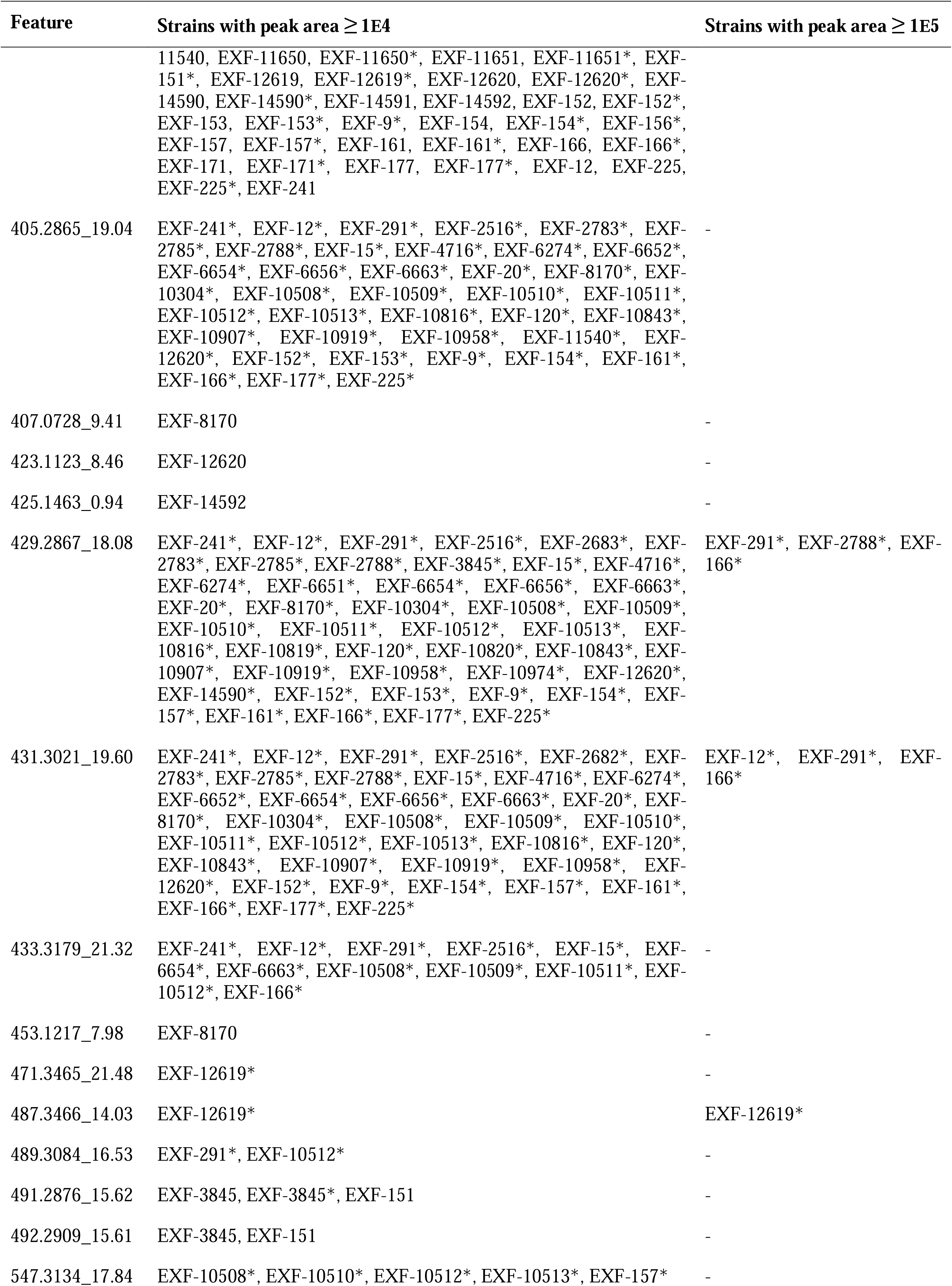
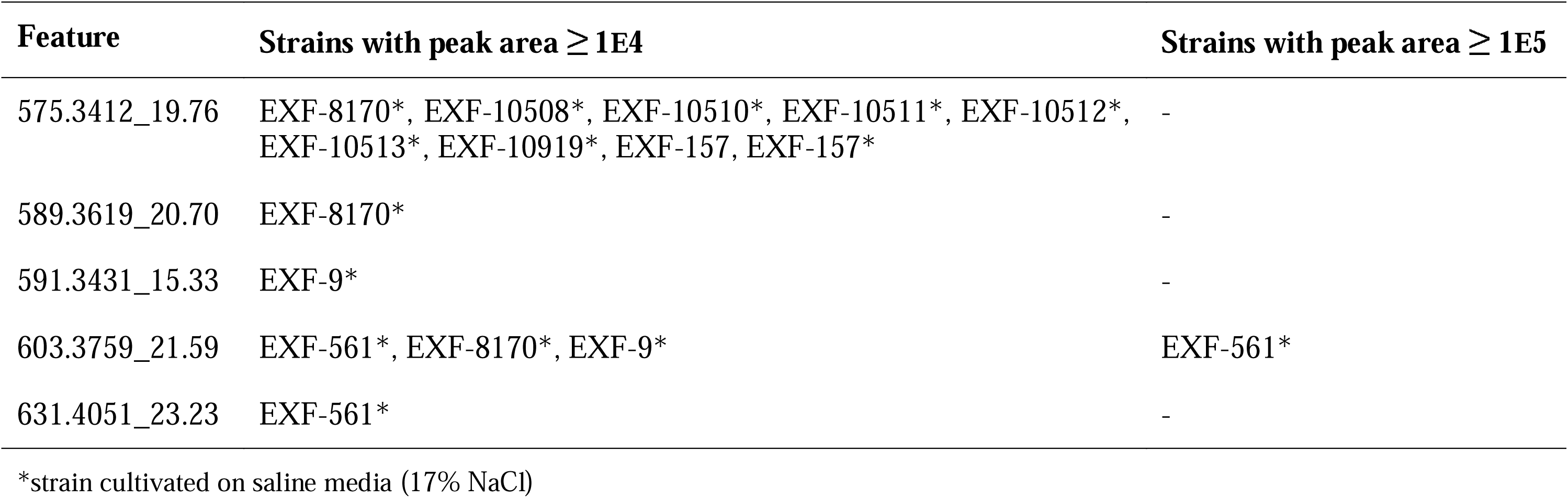
The top 36 features of interest and the strains in which they were detected in high intensity (peak area ≥ 1E4).

## References

1. Anderson, M.J., Gorley, R.N., Clarke, K.R., 2008. PERMANOVA+ for primer: Guide to software and statistical methods. PRIMER-E Ltd, Plymouth.

2. Bao, Y.-R., Chen, G.-D., Wu, Y.-H., Li, X.-X., Hu, D., Liu, X.-Z., Li, Y., Yao, X.-S., Gao, H., 2015. Stachybisbins A and B, the first cases of seco-bisabosquals from *Stachybotrys bisbyi*. Fitoterapia 105, 151–155. 10.1016/j.fitote.2015.06.022

3. Baumgartner, J.T., McCaughey, C.S., Fleming, H.S., Lentz, A.R., Sanchez, L.M., McKinnie, S.M.K., 2024. Vanadium-dependent haloperoxidases from diverse microbes halogenate exogenous alkyl quinolone quorum sensing signals. bioRxiv 2024.07.31.606109 preprint. 10.1101/2024.07.31.606109

4. Becker, K., Wessel, A.-C., Luangsa-ard, J.J., Stadler, M., 2020. Viridistratins A-C, Antimicrobial and cytotoxic benzo[j]fluoranthenes from stromata of Annulohypoxylon viridistratum (Hypoxylaceae, Ascomycota). Biomolecules 10, 1–11. 10.3390/biom10050805

5. Bertrand, S., Guitton, Y., Roullier, C., 2017. Successes and pitfalls in automated dereplication strategy using liquid chromatography coupled to mass spectrometry data: A CASMI 2016 experience. Phytochemistry Letters 21, 297–305. 10.1016/j.phytol.2016.12.025

6. Blin, K., Shaw, S., Kloosterman, A.M., Charlop-Powers, Z., van Wezel, P., Medema, M.H., Weber, T., 2021. antiSMASH 6.0: improving cluster detection and comparison capabilities. Nucleic Acids Research 49, W29–W35. 10.1093/nar/gkab335

7. Blin, K., Shaw, S., Vader, L., Szenei, J., Reitz, Z.L., Augustijn, H.E., Cediel-Becerra, J.D.D., de Crécy-Lagard, V., Koetsier, R.A., Williams, S.E., Cruz-Morales, P., Wongwas, S., Segurado Luchsinger, A.E., Biermann, F., Korenskaia, A., Zdouc, M.M., Meijer, D., Terlouw, B.R., van der Hooft, J.J.J., Ziemert, N., Helfrich, E.J.N., Masschelein, J., Corre, C., Chevrette, M.G., van Wezel, G.P., Medema, M.H., Weber, T., 2025. antiSMASH 8.0: extended gene cluster detection capabilities and analyses of chemistry, enzymology, and regulation. Nucleic Acids Research 53, W32–W38. 10.1093/nar/gkaf334

8. Brauers, G., Ebel, R., Edrada, R., Wray, V., Berg, A., Gräfe, U., Proksch, P., 2001. Hortein, a new natural product from the fungus *Hortaea werneckii* associated with the sponge *Aplysina aerophoba*. J. Nat. Prod. 64, 651–652. 10.1021/np000542u

9. Burgaud, G., Arzur, D., Durand, L., Cambon-Bonavita, M.-A., Barbier, G., 2010. Marine culturable yeasts in deep-sea hydrothermal vents: species richness and association with fauna. FEMS Microbiology Ecology 73, 121–133. 10.1111/j.1574-6941.2010.00881.x

10. Bürkner, P.-C., 2021. Bayesian Item Response Modeling in *R* with *brms* and *Stan*. J. Stat. Soft. 100. 10.18637/jss.v100.i05

11. Capella-Gutiérrez, S., Silla-Martínez, J.M., Gabaldón, T., 2009. trimAl: a tool for automated alignment trimming in large-scale phylogenetic analyses. Bioinformatics 25, 1972–1973. 10.1093/bioinformatics/btp348.

12. Carpenter, B., Gelman, A., Hoffman, M.D., Lee, D., Goodrich, B., Betancourt, M., Brubaker, M., Guo, J., Li, P., Riddell, A., 2017. *Stan*: A probabilistic programming language. J. Stat. Soft. 76. 10.18637/jss.v076.i01

13. Chen, Y., Wu, X., Xu, L., El-Shazly, M., Ma, C., Yuan, S., Wang, P., Luo, L., 2022. Two new cerebroside metabolites from the marine fungus *Hortaea werneckii*. Chemistry & Biodiversity 19. 10.1002/cbdv.202200008

14. Clarke, K.R., Gorley, R.N., 2015. PRIMER v7: User Manual / Tutorial. PRIMER-E Ltd, Plymouth.

15. Cochereau, B., Le Strat, Y., Ji, Q., Pawtowski, A., Delage, L., Weill, A., Mazéas, L., Hervé, C., Burgaud, G., Gunde-Cimerman, N., Pouchus, Y.F., Demont-Caulet, N., Roullier, C., Meslet-Cladiere, L., 2023. Heterologous expression and biochemical characterization of a new chloroperoxidase isolated from the deep-sea hydrothermal vent black yeast *Hortaea werneckii* UBOCC-A-208029. Mar Biotechnol 25, 519–536. 10.1007/s10126-023-10222-7

16. Darriba, D., Posada, D., Kozlov, A.M., Stamatakis, A., Morel, B., Flouri, T., 2020. ModelTest-NG: a new and scalable tool for the selection of DNA and protein evolutionary models. Mol Biol Evol 37, 291–294. 10.1093/molbev/msz189.

17. De Leo, F., Lo Giudice, A., Alaimo, C., De Carlo, G., Rappazzo, A.C., Graziano, M., De Domenico, E., Urzì, C., 2019. Occurrence of the black yeast H*ortaea werneckii* in the Mediterranean Sea. Extremophiles 23, 9–17. 10.1007/s00792-018-1056-1

18. Dührkop, K., Fleischauer, M., Ludwig, M., Aksenov, A.A., Melnik, A.V., Meusel, M., Dorrestein, P.C., Rousu, J., Böcker, S., 2019. SIRIUS 4: a rapid tool for turning tandem mass spectra into metabolite structure information. Nature Methods 16, 299–302.

19. Dunbar, K.L., Melby, J.O., Mitchell, D.A., 2012. YcaO domains use ATP to activate amide backbones during peptide cyclodehydrations. Nat Chem Biol 8(6), 569–75. 10.1038/nchembio.944

20. Elsayis, A., Hassan, S.W.M., Ghanem, K.M., Khairy, H., 2022. Suggested sustainable medical and environmental uses of melanin pigment from halotolerant black yeast *Hortaea werneckii* AS1. Front. Microbiol. 13, 871394. 10.3389/fmicb.2022.871394

21. Gostinčar, C., Stajich, J.E., Zupančič, J., Zalar, P., Gunde-Cimerman, N., 2018. Genomic evidence for intraspecific hybridization in a clonal and extremely halotolerant yeast. BMC Genomics 19, 364. 10.1186/s12864-018-4751-5

22. Gostinčar, C., Sun, X., Černoša, A., Fang, C., Gunde-Cimerman, N., Song, Z., 2022. Clonality, inbreeding, and hybridization in two extremotolerant black yeasts. GigaScience 11, giac095. 10.1093/gigascience/giac095

23. Graziano, N., Arce-López, B., Barbeyron, T., Delage, L., Gerometta, E., Roullier, C., Burgaud, G., Poirier, E., Martinelli, L., Jany, J.-L., Hymery, N., Meslet-Cladiere, L., 2024. Identification and characterization of two aryl sulfotransferases from deep-sea marine fungi and their implications in the sulfation of secondary metabolites. Mar. Drugs 22, 572. 10.3390/md22120572

24. Grigoriev, I.V., Nikitin, R., Haridas, S., Kuo, A., Ohm, R., Otillar, R., Riley, R., Salamov, A., Zhao, X., Korzeniewski, F., Smirnova, T., Nordberg, H., Dubchak, I., Shabalov, I., 2013. MycoCosm portal: gearing up for 1000 fungal genomes. Nucleic Acids Research 42, D699–D704. 10.1093/nar/gkt1183

25. Gunde-Cimerman, N., Plemenitaš, A., 2006. Ecology and molecular adaptations of the halophilic black yeast *Hortaea werneckii*. Rev Environ Sci Biotechnol 5, 323–331. 10.1007/s11157-006-9105-0

26. Gunde-Cimerman, N., Plemenitaš, A., Oren, A., 2018. Strategies of adaptation of microorganisms of the three domains of life to high salt concentrations. FEMS Microbiology Reviews 42, 353–375. 10.1093/femsre/fuy009

27. Gunde-Cimerman, N., Zalar, P., 2014. Extremely halotolerant and halophilic fungi inhabit brine in solar salterns around the globe. Food Technol. Biotechnol. 52, 170–179.

28. Kozlov, A.M., Darriba, D., Flouri, T., Morel, B., Stamatakis, A., 2019. RAxML-NG: a fast, scalable and user-friendly tool for maximum likelihood phylogenetic inference. Bioinformatics 35, 4453–4455. 10.1093/bioinformatics/btz305.

29. Kuraku, S., Zmasek, C.M., Nishimura, O., Katoh, K., 2013. aLeaves facilitates on-demand exploration of metazoan gene family trees on MAFFT sequence alignment server with enhanced interactivity. Nucleic Acids Res 41, W22–W28. 10.1093/nar/gkt389.

30. Lenassi, M., Gostinčar, C., Jackman, S., Turk, M., Sadowski, I., Nislow, C., Jones, S., Birol, I., Cimerman, N.G., Plemenitaš, A., 2013. Whole genome duplication and enrichment of metal cation transporters revealed by de novo genome sequencing of extremely halotolerant black yeast Hortaea werneckii. PLoS ONE 8, e71328. 10.1371/journal.pone.0071328

31. Marchetta, A., Gerrits van den Ende, B., Al-Hatmi, A., Hagen, F., Zalar, P., Sudhadham, M., Gunde-Cimerman, N., Urzì, C., de Hoog, S., De Leo, F., 2018. Global molecular diversity of the halotolerant fungus *Hortaea werneckii*. Life 8, 31. 10.3390/life8030031

32. Nothias, L.F., Petras, D., Schmid, R., Dührkop, K., Rainer, J., Sarvepalli, A., Protsyuk, I., Ernst, M., Tsugawa, H., Fleischauer, M., Aicheler, F., Aksenov, A., Alka, O., Allard, P.-M., Barsch, A., Cachet, X., Caraballo, M., Da Silva, R.R., Dang, T., Garg, N., Gauglitz, J.M., Gurevich, A., Isaac, G., Jarmusch, A.K., Kameník, Z., Kang, K.B., Kessler, N., Koester, I., Korf, A., Gouellec, A.L., Ludwig, M., Christian, M.H., McCall, L.-I., McSayles, J., Meyer, S.W., Mohimani, H., Morsy, M., Moyne, O., Neumann, S., Neuweger, H., Nguyen, N.H., Nothias-Esposito, M., Paolini, J., Phelan, V.V., Pluskal, T., Quinn, R.A., Rogers, S., Shrestha, B., Tripathi, A., van der Hooft, J.J.J., Vargas, F., Weldon, K.C., Witting, M., Yang, H., Zhang, Z., Zubeil, F., Kohlbacher, O., Böcker, S., Alexandrov, T., Bandeira, N., Wang, M., Dorrestein, P.C., 2020. Feature-based molecular networking in the GNPS analysis environment. Nature Methods 17, 905–908. 10.1038/s41592-020-0933-6

33. Oksanen, J., Simpson, G.L., Blanchet, F.G., Kindt, R., Legendre, P., Minchin, P.R., O’Hara, R.B., Solymos, P., Stevens, M.H.H., Szoecs, E., Wagner, H., Barbour, M., Bedward, M., Bolker, B., Borcard, D., Carvalho, G., Chirico, M., De Caceres, M., Durand, S., Evangelista, H.B.A., FitzJohn, R., Friendly, M., Furneaux, B., Hannigan, G., Hill, M.O., Lahti, L., McGlinn, D., Ouelette, M.-H., Ribeiro Cunha, E., Smith, T., Stier, A., Ter Braak, C.J.F., Weedon, J., Borman, T., 2025. Vegan: community ecology package.

34. Petrovič, U., Gunde-Cimerman, N., Plemenitaš, A., 2002. Cellular responses to environmental salinity in the halophilic black yeast *Hortaea werneckii*. Molecular Microbiology 45, 665–672. 10.1046/j.1365-2958.2002.03021.x

35. R Core Team, 2024. R: A language and environment for statistical computing. R Foundation for Statistical Computing, Vienna, Austria.

36. Rokas, A., Wisecaver, J.H., Lind, A.L., 2018. The birth, evolution and death of metabolic gene clusters in fungi. Nat Rev Microbiol 16, 731–744. 10.1038/s41579-018-0075-3

37. Rutz, A., Sorokina, M., Galgonek, J., Mietchen, D., Willighagen, E., Gaudry, A., Graham, J.G., Stephan, R., Page, R., Vondrášek, J., Steinbeck, C., Pauli, G.F., Wolfender, J.-L., Bisson, J., Allard, P.-M., 2022. The LOTUS initiative for open knowledge management in natural products research. eLife 11, e70780. 10.7554/eLife.70780

38. Schmid, R., Heuckeroth, S., Smirnov, A., Myers, O., Dyrlund, T.S., Bushuiev, R., Murray, K.J., Hoffman, N., Lu, M., Sarvepalli, A., Zhang, Z., Fleischauer, M., Dührkop, K., Wesner, M., Hoogstra, S.J., Rudt, E., Mokshyna, O., Brungs, C., Ponomarov, K., Mutabdzija, L., Damiani, T., Pudney, C.J., Earll, M., Helmer, P.O., Fallon, T.R., Schulze, T., Rivas-Ubach, A., Bilbao, A., Richter, H., Nothias, L.-F., Wang, M., Orešič, M., Weng, J.-K., Böcker, S., Jeibmann, A., Hayen, H., Karst, U., Dorrestein, P.C., Petras, D., Du, X., Pluskal, T., 2023. Integrative analysis of multimodal mass spectrometry data in MZmine 3. Nature Biotechnology 41, 447–449. 10.1038/s41587-023-01690-2

39. Stan Development Team, 2018. RStan: The R interface to Stan.

40. Vaz, B.M.C., C.T.S. Leite, M.S., Contieri, L.S., De Souza Mesquita, L.M., Conde, A., Oliveira, J.P., Pinto, D.C.G.A., Ventura, S.P.M., 2025. Efficient extraction and purification of mycosporines-like amino acids (MAAs) following a multiproduct biorefinery approach. Separation and Purification Technology 363, 132200. 10.1016/j.seppur.2025.132200

41. Volkmann, M., Gorbushina, A.A., 2006. A broadly applicable method for extraction and characterization of mycosporines and mycosporine-like amino acids of terrestrial, marine and freshwater origin. FEMS Microbiology Letters 255, 286–295. 10.1111/j.1574-6968.2006.00088.x

42. Wang, M., Carver, J.J., Phelan, V.V., Sanchez, L.M., Garg, N., Peng, Y., Nguyen, D.D., Watrous, J., Kapono, C.A., Luzzatto-Knaan, T., Porto, C., Bouslimani, A., Melnik, A.V., Meehan, M.J., Liu, W.-T., Crüsemann, M., Boudreau, P.D., Esquenazi, E., Sandoval-Calderón, M., Kersten, R.D., Pace, L.A., Quinn, R.A., Duncan, K.R., Hsu, C.-C., Floros, D.J., Gavilan, R.G., Kleigrewe, K., Northen, T., Dutton, R.J., Parrot, D., Carlson, E.E., Aigle, B., Michelsen, C.F., Jelsbak, L., Sohlenkamp, C., Pevzner, P., Edlund, A., McLean, J., Piel, J., Murphy, B.T., Gerwick, L., Liaw, C.-C., Yang, Y.-L., Humpf, H.-U., Maansson, M., Keyzers, R.A., Sims, A.C., Johnson, A.R., Sidebottom, A.M., Sedio, B.E., Klitgaard, A., Larson, C.B., Boya P, C.A., Torres-Mendoza, D., Gonzalez, D.J., Silva, D.B., Marques, L.M., Demarque, D.P., Pociute, E., O’Neill, E.C., Briand, E., Helfrich, E.J.N., Granatosky, E.A., Glukhov, E., Ryffel, F., Houson, H., Mohimani, H., Kharbush, J.J., Zeng, Y., Vorholt, J.A., Kurita, K.L., Charusanti, P., McPhail, K.L., Nielsen, K.F., Vuong, L., Elfeki, M., Traxler, M.F., Engene, N., Koyama, N., Vining, O.B., Baric, R., Silva, R.R., Mascuch, S.J., Tomasi, S., Jenkins, S., Macherla, V., Hoffman, T., Agarwal, V., Williams, P.G., Dai, J., Neupane, R., Gurr, J., Rodríguez, A.M.C., Lamsa, A., Zhang, C., Dorrestein, K., Duggan, B.M., Almaliti, J., Allard, P.-M., Phapale, P., Nothias, L.-F., Alexandrov, T., Litaudon, M., Wolfender, J.-L., Kyle, J.E., Metz, T.O., Peryea, T., Nguyen, D.-T., VanLeer, D., Shinn, P., Jadhav, A., Müller, R., Waters, K.M., Shi, W., Liu, X., Zhang, L., Knight, R., Jensen, P.R., Palsson, B.Ø., Pogliano, K., Linington, R.G., Gutiérrez, M., Lopes, N.P., Gerwick, W.H., Moore, B.S., Dorrestein, P.C., Bandeira, N., 2016. Sharing and community curation of mass spectrometry data with Global Natural Products Social Molecular Networking. Nat Biotechnol 34, 828–837. 10.1038/nbt.3597

43. Zalar, P., Zupančič, J., Gostinčar, C., Zajc, J., De Hoog, G.S., De Leo, F., Azua-Bustos, A., Gunde-Cimerman, N., 2019. The extremely halotolerant black yeast *Hortaea werneckii* – a model for intraspecific hybridization in clonal fungi. IMA Fungus 10, 10. 10.1186/s43008-019-0007-5

44. Zhang, P., Wang, X., Fan, A., Zheng, Y., Liu, X., Wang, S., Zou, H., Oakley, B.R., Keller, N.P., Yin, W.-B., 2017. A cryptic pigment biosynthetic pathway uncovered by heterologous expression is essential for conidial development in *Pestalotiopsis fici*. Molecular Microbiology 105, 469–483. 10.1111/mmi.13711

45. Zhao, F., Wang, Z., Huang, H., 2024. Physical cell disruption technologies for intracellular compound extraction from microorganisms. Processes 12, 2059. 10.3390/pr12102059

