## Supplementary material for "Novel insights into the specialized metabolism of the extremophile black yeast *Hortaea werneckii* and its modulation, unlocking its potential for compound discovery": Supp data with sequences for Indole/DMATS

**Sequences for Indole/DMATS from the 64 strains of *Hortaea werneckii* genomes**

>Indole_120

MAKGDLDTTAAWQSLNLYLPTRTHDEDYWWQKSGPQLAALVEGAEYPLAKQYEALLFHYHWMVPYMGPSPLPEGAARQWKSLLQPDGTPIECSWKWNTSRSPPDIRYDIEPIGPLAGTKADPLNQHALREMLHRLAGQVPNVDLTWCDHFLSTLFDHDLSKYVAESAAGKRPTTSGVIAAEFLESGTRFKTYFQPRKLGYTGIIPMKMWDEALEPIDPQRAARSMVKDFPESTAAGQTLTCFSIAVDVVKLEKSRLKWYFNTPSTAFSIVREVMTLGGRLSSPH

>Indole_225

MAKGDLDTTAAWQSLNLYLPTRTHDEDYWWQKSGPQLAALVEGAEYPLAKQYEALLFHYHWMVPYMGPSPLPEGAARQWKSLLQPDGTPIECSWKWNTSRSPPDIRYDIEPIGPLAGTKADPLNQHALREMLHRLAGQVPNVDLTWCDHFLSTLFDHDLSKYVAESAAGKRPTTSGVIAAEFLESGTRFKTYFQPRKLGYTGIIPMKMWDEALEPIDPQRAARSMVKDFPESTAAGQTLTCFSIAVDVVKLEKSRLKWYFNTPSTAFSIVREVMTLGGRLSSPH

>Indole_14590

MAKGDLDTTAAWQSLNLYLPTRTHDEDYWWQKSGPQLAALVEGAEYPLAKQYEALLFHYHWMVPYMGPSPLPEGAARQWKSLLQPDGTPIECSWKWNTSRSPPDIRYDIEPIGPLAGTKADPLNQHALREMLHRLAGQVPNVDLTWCDHFLSTLFDHDLSKYVAESAAGKRPTTSGVIAAEFLESGTRFKTYFQPRKLGYTGIIPMKMWDEALEPIDPQRAARSMVKDFPESTAAGQTLTCFSIAVDVVKLEKSRLKWYFNTPSTAFSIVREVMTLGGRLSSPH

>Indole_10919

MAKGDLDTTAAWQSLNLYLPTRTHDEDYWWQKSGPQLAALVEGAEYPLAKQYEALLFHYHWMVPYMGPSPLPEGAARQWKSLLQPDGTPIECSWKWNTSRSPPDIRYDIEPIGPLAGTKADPLNQHALREMLHRLAGQVPNVDLTWCDHFLSTLFDHDLSKYVAESAAGKRPTTSGVIAAEFLESGTRFKTYFQPRKLGYTGIIPMKMWDEALEPIDPQRAARSMVKDFPESTAAGQTLTCFSIAVDVVKLEKSRLKWYFNTPSTAFSIVREVMTLGGRLSSPH

>Indole_10907

MAKGDLDTTAAWQSLNLYLPTRTHDEDYWWQKSGPQLAALVEGAEYPLAKQYEALLFHYHWMVPYMGPSPLPEGAARQWKSLLQPDGTPIECSWKWNTSRSPPDIRYDIEPIGPLAGTKADPLNQHALREMLHRLAGQVPNVDLTWCDHFLSTLFDHDLSKYVAESAAGKRPTTSGVIAAEFLESGTRFKTYFQPRKLGYTGIIPMKMWDEALEPIDPQRAARSMVKDFPESTAAGQTLTCFSIAVDVVKLEKSRLKWYFNTPSTAFSIVREVMTLGGRLSSPH

>Indole_10904

MAKGDLDTTAAWQSLNLYLPTRTHDEDYWWQKSGPQLAALVEGAEYPLAKQYEALLFHYHWMVPYMGPSPLPEGAARQWKSLLQPDGTPIECSWKWNTSRSPPDIRYDIEPIGPLAGTKADPLNQHALREMLHRLAGQVPNVDLTWCDHFLSTLFDHDLSKYVAESAAGKRPTTSGVIAAEFLESGTRFKTYFQPRKLGYTGIIPMKMWDEALEPIDPQRAARSMVKDFPESTAAGQTLTCFSIAVDVVKLEKSRLKWYFNTPSTAFSIVREVMTLGGRLSSPH

>Indole_10843

MAKGDLDTTAAWQSLNLYLPTRTHDEDYWWQKSGPQLAALVEGAEYPLAKQYEALLFHYHWMVPYMGPSPLPEGAARQWKSLLQPDGTPIECSWKWNTSRSPPDIRYDIEPIGPLAGTKADPLNQHALREMLHRLAGQVPNVDLTWCDHFLSTLFDHDLSKYVAESAAGKRPTTSGVIAAEFLESGTRFKTYFQPRKLGYTGIIPMKMWDEALEPIDPQRAARSMVKDFPESTAAGQTLTCFSIAVDVVKLEKSRLKWYFNTPSTAFSIVREVMTLGGRLSSPH

>Indole_10819

MAKGDLDTTAAWQSLNLYLPTRTHDEDYWWQKSGPQLAALVEGAEYPLAKQYEALLFHYHWMVPYMGPSPLPEGAARQWKSLLQPDGTPIECSWKWNTSRSPPDIRYDIEPIGPLAGTKADPLNQHALREMLHRLAGQVPNVDLTWCDHFLSTLFDHDLSKYVAESAAGKRPTTSGVIAAEFLESGTRFKTYFQPRKLGYTGIIPMKMWDEALEPIDPQRAARSMVKDFPESTAAGQTLTCFSIAVDVVKLEKSRLKWYFNTPSTAFSIVREVMTLGGRLSSPH

>Indole_10816

MAKGDLDTTAAWQSLNLYLPTRTHDEDYWWQKSGPQLAALVEGAEYPLAKQYEALLFHYHWMVPYMGPSPLPEGAARQWKSLLQPDGTPIECSWKWNTSRSPPDIRYDIEPIGPLAGTKADPLNQHALREMLHRLAGQVPNVDLTWCDHFLSTLFDHDLSKYVAESAAGKRPTTSGVIAAEFLESGTRFKTYFQPRKLGYTGIIPMKMWDEALEPIDPQRAARSMVKDFPESTAAGQTLTCFSIAVDVVKLEKSRLKWYFNTPSTAFSIVREVMTLGGRLSSPH

>Indole_8422

MAKGDLDTTAAWQSLNLYLPTRTHDEDYWWQKSGPQLAALVEGAEYPLAKQYEALLFHYHWMVPYMGPSPLPEGAARQWKSLLQPDGTPIECSWKWNTSRSPPDIRYDIEPIGPLAGTKADPLNQHALREMLHRLAGQVPNVDLTWCDHFLSTLFDHDLSKYVAESAAGKRPTTSGVIAAEFLESGTRFKTYFQPRKLGYTGIIPMKMWDEALEPIDPQRAARSMVKDFPESTAAGQTLTCFSIAVDVVKLEKSRLKWYFNTPSTAFSIVREVMTLGGRLSSPH

>Indole_4716

MAKGDLDTTAAWQSLNLYLPTRTHDEDYWWQKSGPQLAALVEGAEYPLAKQYEALLFHYHWMVPYMGPSPLPEGAARQWKSLLQPDGTPIECSWKWNTSRSPPDIRYDIEPIGPLAGTKADPLNQHALREMLHRLAGQVPNVDLTWCDHFLSTLFDHDLSKYVAESAAGKRPTTSGVIAAEFLESGTRFKTYFQPRKLGYTGIIPMKMWDEALEPIDPQRAARSMVKDFPESTAAGQTLTCFSIAVDVVKLEKSRLKWYFNTPSTAFSIVREVMTLGGRLSSPH

>Indole_2516

MSSITTSDHVSYEAESLPMLAQINREVRPDDEDQAFWWNCLSETLASLLQANQYSNDMQLHYLRWFYKWIPQALGPRPIDGEPYYGSWITHDLSPLEYSLNWKEKSSKQTVRFTIEAVTKQAGTAKDPINQLGAKEFLDAASKDIEGLDLTRFNQFLEATNIPNDSAEETAAKHPAHFPRSRVWLAFDLEHSGTLMVKSYFLPHWRAIQSGISANTFISDTVRACNGPDGSKYDGALDAIHSYLLSLQEHPPQIGLLSNDCVAETAESRLKIYFRSEADTLNKARSMYNLGGRLKGTNIDASLEGISDIWHHLFRLDRSDPGSADKVCLGEHKCIFVYEMRPTQDNTPNIDVKFHIPMWQLAKTDNELSELMASWFASRGHADFAARYKSDLNKAFPRSMTEKSLGTHTYLSITYTPKTGHYLTMYLSPKLPETFF

>Indole_2515

MTTKQRPILTIGAGLAGLSLAHVLTNHVISNIVFEAATQDRSQGYAISLKRWAYDPLLASLGHLQLKQLTKVVAPDRARGGKGWLDLALRDNASGDILVAPDPGQRDGVVRANREALLDWIANGGDKNLDVRHGHKFRACRRPSGVVTAVFENGAEHTGSLVVAVDGVFSAVRSHLLPHITPEVLPVVHTGSSFRQEFLTFKRIDPTPLSWSGVDAITSCSINTMAYAAGWSKVPSNMSLIYEDRKFEAGQLVGHHVALMLHEADYHPARQVELLLFHRFTLLPFLGPRPTSSTPWFCSRVAPGAGDGTPIGYSWCWSSSGDRPTIRHYLEPLGPLTGTAADPLNDTAPKQLLLQLAKTFPDTQLDALWKFAAHLRPKARSSDARRFNGSSLLLGLEMGAENDRVDVTAGLITKVPEQVDALLRKIIPAAMRDAYGADVSLDALEAARAFIDADPHLVLNGTMAVDCISPTVSRFKFYVMNTSASFDHIAAVMTFGGRKEVDLEHLKELRQLWYRLKGLPDDFPDSAEPPLPATNGVSNGAAVAGNVTGLSFYFDLHPKHVLPDVKMQVDLSRHATTDLAAAEAVASFLESHGQPHYAAAYMNVLHAFVSREELAAERGVHAYFSFSLKPDGLEIKSKLVYPHYSTSEDLSSGATRVLGGNPGQLRLQGTNTYIIGTGSSRILIDNGSGDQQWIDTIADLLQNRHIELQYILLTHWHGDYTGGVQSLVAHDSKLADRIHKHMPDGDQQDISDGQIFAVEGATVRAVFTPGHAVDHMCFLLEEENALFTGDNVLGHGFSVVEDLGVYMRSIDTMAGLQCSTGYPGHGARIDKLPDKMKEYIVHKEFRARQVMSALSRTNRNGMSLTEVARSIYGEVPQEMIEKVLLPFLTQMMWNLAEDRIVGFAPGDPRMRKWFARPPIRRLHTV

>Indole_291

MAKGDLDTTAAWQSLNLYLPTRTHDEDYWWQKSGPQLAALVEGAEYPLAKQYEALLFHYHWMVPYMGPSPLPEGAARQWKSLLQPDGTPIECSWKWNTSRSPPDIRYDIEPIGPLAGTKADPLNQHALREMLHRLAGQVPNVDLTWCDHFLSTLFDHDLSKYVAESAAGKRPTTSGVIAAEFLESGTRFKTYFQPRKLGYTGIIPMKMWDEALEPIDPQRAARSMVKDFPESTAAGQTLTCFSIAVDVVKLEKSRLKWYFNTPSTAFSIVREVMTLGGRLSSPH

>Indole_10512

MAKGDLDTTAAWQSLNLYLPTRTHDEDYWWQKSGPQLAALVEGAEYPLAKQYEALLFHYHWMVPYMGPSPLPEGAARQWKSLLQPDGTPIECSWKWNTSRSPPDIRYDIEPIGPLAGTKADPLNQHALREMLHRLAGQVPNVDLTWCDHFLSTLFDHDLSKYVAESAAGKRPTTSGVIAAEFLESGTRFKTYFQPRKLGYTGIIPMKMWDEALEPIDPQRAARSMVKDFPESTAAGQTLTCFSIAVDVVKLEKSRLKWYFNTPSTAFSIVREVMTLGGRLSSPH

>Indole_177

MSSITTSDHVSYEAESLPMLAQINREVRPDDEDQAFWWNCLSETLASLLQANQYSNDMQLHYLRWFYKWIPQALGPRPIDGEPYYGSWITHDLSPLEYSLNWKEKSSKQTVRFTIEAVTKQAGTAKDPINQLGAKEFLDAASKDIEGLDLTRFNQFLEATNIPNDSAEETAAKHPAHFPRSRVWLAFDLEHSGTLMVKSYFLPHWRAIQSGISANTFISDTVRACNGPDGSKYDGALDAIHSYLLSLQEHPPQIGLLSNDCVAETAESRLKIYFRSEADTLNKARSMYNLGGRLKGTNIDASLEGISDIWHHLFRLDRSDPGSADKVCLGEHKCIFVYEMRPTQDNTPNIDVKFHIPMWQLAKTDNELSELMASWFASRGHADFAARYKSDLNKAFPRSMTEKSLGTHTYLSITYTPKTGHYLTMYLSPKLPETFF

>Indole_154

MAKGDLDTTAAWQSLNLYLPTRTHDEDYWWQKSGPQLAALVEGAEYPLAKQYEALLFHYHWMVPYMGPSPLPEGAARQWKSLLQPDGTPIECSWKWNTSRSPPDIRYDIEPIGPLAGTKADPLNQHALREMLHRLAGQVPNVDLTWCDHFLSTLFDHDLSKYVAESAAGKRPTTSGVIAAEFLESGTRFKTYFQPRKLGYTGIIPMKMWDEALEPIDPQRAARSMVKDFPESTAAGQTLTCFSIAVDVVKLEKSRLKWYFNTPSTAFSIVREVMTLGGRLSSPH

>Indole_152

MAKGDLDTTAAWQSLNLYLPTRTHDEDYWWQKSGPQLAALVEGAEYPLAKQYEALLFHYHWMVPYMGPSPLPEGAARQWKSLLQPDGTPIECSWKWNTSRSPPDIRYDIEPIGPLAGTKADPLNQHALREMLHRLAGQVPNVDLTWCDHFLSTLFDHDLSKYVAESAAGKRPTTSGVIAAEFLESGTRFKTYFQPRKLGYTGIIPMKMWDEALEPIDPQRAARSMVKDFPESTAAGQTLTCFSIAVDVVKLEKSRLKWYFNTPSTAFSIVREVMTLGGRLSSPH

>Indole_12

MAKGDLDTTAAWQSLNLYLPTRTHDEDYWWQKSGPQLAALVEGAEYPLAKQYEALLFHYHWMVPYMGPSPLPEGAARQWKSLLQPDGTPIEYSWKWNTSRSPPDIRYDIEPIGPLAGTKADPLNQHALREMLHRLADQVPNVDLTWCDHFLSTLFDHDLSKYVAESAAGKRPTTSGVIAAEFLESGTRFKTYFQPRKLGYTGIIPMKMWDEALEPIDPQRAARSMVKDFPESTAAGQTLTCFSIAVDVVKLEKSRLKWYFNTPSTAFSIVREVMTLGGRLSSPH

>Indole_12620

MSSITTSDHVSYEAESLPMLAQINREVRPDDEDQAFWWNCLSETLASLLQANQYSNDMQLHYLRWFYKWIPQALGPRPIDGKPYYGSWITHDLSPLEYSLNWKEKSSKQTVRFTIEAVTKQAGTAKDPINQLGAKEFLDAASKDIEGLDLTRFNQFLEATNIPNDSAEETAAKHPAHFPRSRVWLAFDLEHSGTLMVKSYFLPHWRAIQSGISANTFISDTVRACNGPDGSKYDGALDAIHSYLLSLQEHPPQIGLLSNDCVAETAESRLKIYFRSEADTLNKARSMYNLGGRLKGTNIDASLEGISDIWHHLFRLDRSDPGSADKVCLGEHKCIFVYEMRPTQDNTPNIDVKFHIPMWQLAKTDNELSELMASWFASRGHADFAARYKSDLNKAFPQSMTEKSLGTHTYLSITYTPKTGHYLTMYLSPKLPETFF

>Indole_153

MAKGDLDTTAAWQSLNLYLPTRTHDEDYWWQKSGPQLAALVEGAEYPLAKQYEALLFHYHWMVPYMGPSPLPEGAARQWKSLLQPDGTPIECSWKWNTSRSPPDIRYDIEPIGPLAGTKADPLNQHALREMLHRLAGQVPNVDLTWCDHFLSTLFDHDLSKYVAESAAGKRPTTSGVIAAEFLESGTRFKTYFQPRKLGYTGIIPMKMWDEALEPIDPQRAARSMVKDFPESTAAGQTLTCFSIAVDVVKLEKSRLKWYFNTPSTAFSIVREVMTLGGRLSSPH

>Indole_2785

MSSITTSDHVSYEAESLPMLAQINREVRPDDEDQAFWWNCLSETLASLLQANQYSNDMQLHYLRWFYKWIPQALGPRPIDGKPYYGSWITHDLSPLEYSLNWKEKSSKQTVRFTIEAVTKQAGTAKDPINQLGAKEFLDAASKDIEGLDLTRFNQFLEATNIPNDSAEETAAKHPAHFPRSRVWLAFDLEHSGTLMVKSYFLPHWRAIQSGISANTFISDTVRACNGPDGSKYDGALDAIHSYLLSLQEHPPQIGLLSNDCVAETAESRLKIYFRSEADTLNKARSMYNLGGRLKGTNIDASLEGISDIWHHLFRLDRSDPGSADKVCLGEHKCIFVYEMRPTQDNTPNIDVKFHIPMWQLAKTDNELSELMASWFASRGHADFAARYKSDLNKAFPQSMTEKSLGTHTYLSITYTPKTGHYLTMYLSPKLPETFF
