## Supplementary material for "Novel insights into the specialized metabolism of the extremophile black yeast *Hortaea werneckii* and its modulation, unlocking its potential for compound discovery": Supp data with sequences for PKSI

**Sequences for PKSI from the 64 strains of *Hortaea werneckii* genomes**

>PKSI_1_6656

```
MNKTFPADLKDRKVNISQKAVDWPRNGKEKRKVFLNNFSAAGGNTALLLEDGPAYEAPTATDPRGTVPVT
VTARSISALKRNIANLQKYVSENPSTTLTSMSYTLTARRIQHNYRVSFPLDQINKFSDALQAQVKESYSP
VPNAPTRVAFCFTGQGSQYTGLGQKLYNDLKSFRDDIDQLDHLARVQGLPSFLEIVQGADVQTLSPVKVQ
LGMACIQVALARMWAAWGITPAAVIGHSLGEYAALHVAGVISASDMVLLVGRRAELLVRDCTPHTHGMLA
VKGGAEAIRNTLGNKMTEIACINGPEETVLCGSGDVVGAANETLAAKGFKATKLNVPFAFHSAQVDPILE
QFKKIAASVTYNKPAVPVLSPLEGDIIREAGKINPEYLARHARETVKFWTALTAGQKEKVFDEKTAWLEV
GAHPVCSGMVKASIGATTTAPSLRRGEDAWKTISNSMCTLFTAGVNFNFDEFHKEFNDAQEMYTLPTYSF
DNKKYWLDYHNDWTLRKGEPAQTKEVIVEKPVASASAPAVEMPAKRLSTSCQRVIAENFSGNNGSVTVQS
SLADPKLYPVVCGHMVNNAALCPSSLYADMALTISDYIWKQMRPGTETPGYNVCNMEVPKPLIAQIPQPA
EGQHIQLEANADLDSGIVKLNFRSVMPDGQKLQDHAHCIVRLEDKAAWEDEWSRYNYMVQAQMELLQHKT
LNGGAHKVQRGMAYKLFKALVNYDEKYRAMAEVVLASGQTEASAMLDFPTKPEDGDFYCPPYHIDGSCHI
SGFIVNASDLLDSEQNVYVSHGWGAMKFSRPLTAGMKLRNYVRMQPQPNNVSKGDVYIMEGDQIVAVCEG
IKFQQIPRRVLNTFLPPNKGSGPASAAKPAAAPVAAARPAPAAAPIKTAPAPAKAAPAPAPAAPKAAPKP
KKAAAPKKPAGGLTAKVMKILAKETEVDEGELVDEAQFENLGVDSLLSLTISAVFREELDMDISSTLFTD
YPTVGDMKKYFAQFDNGSSTSSSAEEEDSDEDSIPPTDAATPMDVLSTPASSVPSSAPSDAGKPDSPTRE
TLEDVGDVSLAKHIVAQEMGVDIAEVTDDADLAEMGMDSLMSLTILGELREKTGIDLPSTFLTTNPTMKD
IDNALGMRPKPKAAPKPAAPKAAAPSSSKKTDMNEVSARLSALNNNTDISRYPNATS

>PKSI_1_6669
MRRSTDPAAGYAYCAPAVTIARSVLAVEPLSPGWRYAFHTYWSQ
                     GRLVVYPVSSIPLLLSHQSSTAVGSLQTSHLLQHPLPFLSSVSIDSFACSPKVLSSQE
                     PLDFLHSLAPESYPLHEYPLCLANIVDPEGTHEPHLTDPETIVATHELNTDDYIQKQF
                     YIKMSNVLLFGDQTAEQYPLLNKIVLRKENALVITFVERCAKALREETNALPRSQRNA
                     VPDFLTVNDLKEAYHQKGVKVPMVESALVTIAQIGHYIGYFSEHSAEQPSATNTRALG
                     LCTGLLAAAAVVASKTVEELVLVGVEFVRLSFRSGAAVDAARTALCQTGDDNAPWSTI
                     VTGTTEASAKEALAKFHEEKGIPQTSHAYISAVSVMAITVSGPPTTVKRFFEESPALS
                     KNHRVPIPVYGPYHAEHLFGETEINKIASASILEGLKQHQPVSLVHSAATGKALVAEN
                     AAELAKLVLAEMLQLPVRWDHLLEEAVSQITSKKAPAKIWAMGVSNVANSLVSALKAG
                     GQTDVSTVDQSTWTENEPDTHGRTQNDKVAIVGMAGRFPNSADHEALWELLMKGLDVH
                     RRIPKDRFDADTHVDPSGKGKNKSHTPFGCFIDEPGFFDPRFFNMSPREAAQTDPMGR
                     LALVTAYEALEMSGYVPNRTPSTKLHRIGTFYGQTSDDWREINAAENVDTYFITGGVR
                     AFAPGRINYYFKFSGPSYSVDTACSSSLAAIQLACTSLWAGDCDTACAGGLNVLTNPD
                     IFSGLSKGQFLSKTGSCKTYDNNADGYCRGDAVGTVILKRYEDAIADKDNILGCILGA
                     ATNHSAEAVSITHPHAGAQEFLYKRVLANAGVDAHEISYVEMHGTGTQAGDGIEMTSV
                     TNVFAPRHRQRRDDQPVYLGAIKANVGHAEAASGINSLAKVLLMMKHNKIPANVGIKG
                     EMNKTFPADLKDRKVNISQKAVDWPRNGKEKRKVFLNNFSAAGGNTALLLEDGPAYEA
                     PTATDPRGTVPVTVTARSISALKRNIANLQKYVSENPSTTLTSMSYTLTARRIQHNYR
                     VAFPLDQINKFSDALQAQVKESYSPVPNAPTRVAFCFTGQGSQYTGLGQKLYNDLKSF
                     RDDIDQLDHLARVQGLPSFLEIVQGADVQTLSPVKVQLGMACIQVALARMWAAWGITP
                     AAVIGHSLGEYAALHVAGVISASDMVLLVGRRAELLVRDCTPHTHGMLAVKGGAEAIR
                     NTLGNKMTEIACINGPEETVLCGSGDVVGAANETLAAKGFKATKLNVPFAFHSAQVDP
                     ILEQFKKIAASVTYNKPAVPVLSPLEGDIIREAGKINPEYLARHARETVNFWTALTAG
                     QKEKVFDEKTAWLEVGAHPVCSGMVKASIGATTTAPSLRRGEDAWKTISNSMCTLFTA
                     GVNFNFDEFHKEFNDAQEMYTLPTYSFDNKKYWLDYHNDWTLRKGEPAQTKEVIVEKP
                     VASASAPAVEMPAKRLSTSCQRVISENFSGNNGSVTVQSSLADPKLYPVVCGHMVNNA
                     ALCPSSLYADMALTISDYIWKQMRPGTETPGYNVCNMEVPKPLIAQIPQPAEGQHIQL
                     EANADLDSGIVKLNFRSVKPDGQKLQDHAHCIVRLEDKAAWEDEWSRYNYMVQAQMEL
                     LQHKTLNGGAHKVQRGMAYKLFKALVNYDEKYRAMAEVVLASGQTEASAMLDFPTKPE
                     DGDFYCPPYHIDGSCHISGFIVNASDLLDSEQNVYVSHGWGAMKFSRPLTAGMKLRNY
                     VRMQPQPNNVSKGDVYIMEGDQIVAVCEGIKFQQIPRRVLNTFLPPNKGSGPASAAKP
                     AAAPVAAARPAPAAAPIKSAPAPAKAAPAPAPAAPKAAPKPKKAAAPKKPAGGLTAKV
                     MKILAKETEVDEGELVDEAQFENLGVDSLLSLTISAVFREELDMDISSTLFTDYPTVG
                     DMKKYFAQFDNGSSTSSSAEEEDSDEDSIPPTDAATPMDDLSTPASSVPSSAPSDAGK
                     PDSPTRETLDDVGDVSLAKHIVAQEMGVDIAEVTDDADLAEMGMDSLMSLTILGELRE
                     KTGIDLPSTFLTTNPTMKDIDNALGMRPKPKAAPKPAAPKAAAPSSSKKTDMNEVSAR
                     LSALNNNTDISRYPNATSVLLQGNPKQATKKIFFLPDGSGSATSYVSIPNLGPDVCAY
                     GLNCPFMKNPEQWQCGIEISALVYLAEIKRRQPQGPYIIGGWSAGGVIAYSVAQALLA
                     ANEGVEKLLLLDSPCPVNLAPLPARLHNFFNQIGLLGTGDPAKTPKWLLPHFSAAIRS
                     LSDYDPKPSLRPIPTYAIWCREGVAGNPGDPRPPPAEEEDPAPMTWLLEHRTNFKDNG
                     WAQLCGDSMKFGVMGGHHFSMMKPPHADDLGNLIREGLDWQP


>PKSI_1_6654
MSNVLLFGDQTAEQYQLLNKIVLRKENALVITFVERCAKALREETNALPRSQRNAVPDFLTVNDLKEAYY
QKGVKVPMVESALVTIAQIGHYIGYFSEHSAEQPSATNTRALGLCTGLLAAAAVVASKTVEELVLVGVEF
VRLSFRSGAAVDAARTALCQTGDDNAPWSTIVTGTTEASAKEALAKFHEEKGIPQTSHAYISAVSVMAIT
VSGPPTTVKRFFEESPALSKNHRVPIPVYGPYHAEHLFGETEINKIASAPILEGLKQHQPVSLVHSAATG
KALVAENAAELAKLVLAEMLQLPVRWDHLLEEAVSQITSKKAPAKIWAMGVSNVANSLVSALKAGGQTDV
STVDQSTWTENEPDTHGRTQNDKVAIVGMAGRFPNSADHEALWELLMKGLDVHRRIPKDRFDADTHVDPS
GKGKNKSHTPFGCFIDEPGFFDPRFFNMSPREAAQTDPMGRLALVTAYEALEMSGYVPNRTPSTKLHRIG
TFYGQTSDDWREINAAENVDTYFITGGVRAFAPGRINYYFKFSGPSYSVDTACSSSLAAIQLACTSLWAG
DCDTACAGGLNVLTNPDIFSGLSKGQFLSKTGSCKTYDNNADGYCRGDAVGTVILKRYEDAIADKDNILG
CILGAATNHSAEAVSITHPHAGAQEFLYKRVLANAGVDAHEISYVEMHGTGTQAGDGIEMTSVTNVFAPR
HRQRRDDQPVYLGAIKANVGHAEAASGINSLAKVLLMMKHNKIPANVGIKGEMNKTFPADLKDRKVYISQ
KAVDWPRNGKEKRKVFLNNFSAAGGNTALLLEDGPAYEAPTATDPRGTVPVTVTARSISALKRNIANLQK
YVSENPSTTLTSMSYTLTARRIQHNYRVAFPLDQINKFSDALQAQVKESYSPVPNAPTRVAFCFTGQGSQYTGLGQKLYNDLKSFRDDIDQLDHLARVQGLPSFLEIVQGADVQTLSPVKV
QLGMACIQVALARMWAAWGITPAAVIGHSLGEYAALHVAGVISASDMVLLVGRRAELLVRDCTPHTHGML
AVKGGAEAIRNTLGNKMTEIACINGPEETVLCGSGDVVGAANETLAAKGFKATKLNVPFAFHSAQVDPIL
EQFKKIAASVTYNKPAVPVLSPLEGDIIREAGKINPEYLARHARETVKFWTALTAGQKEKVFDEKTAWLE
VGAHPVCSGMVKASIGATTTAPSLRRGEDAWKTISNSMCTLFTAGVNFNFDEFHKEFNDAQEMYTLPTYS
FDNKKYWLDYHNDWTLRKGEPAQTKEVIVEKPVASASAPAVEMPAKRLSTSCQRVIAENFSGNNGSVTVQ
SSLADPKLYPVVCGHMVNNAALCPSSLYADMALTISDYIWKQMRPGTETPGYNVCNMEVPKPLIAQIPQP
AEGQHIQLEANADLDSGIVKLNFRSVKPDGQKLQDHAHCIVRLEDKAAWEDEWSRYNYMVQAQMELLQHK
TLNGGAHKVQRGMAYKLFKALVNYDEKYRAMAEVVLASGQTEASAMLDFPTKPEDGDFYCPPYHIDGSCH
ISGFIVNASDLLDSEQNVYVSHGWGAMKFSRPLTAGMKLRNYVRMQPQPNNVSKGDVYIMEGDQIVAVCE
GIKFQQIPRRVLNTFLPPNKGSGPASAAKPAAAPVAAARPAPAAAPIKTAPAPAKAAPAPAPAAPKAAPK
PKKAAAPKKPAGGLTAKVMKILAKETEVDEGELVDEAQFENLGVDSLLSLTISAVFREELDMDISSTLFT
DYPTVGDMKKYFAQFDNGSSTSSSAEEEDSDEDSIPPTDAATPMDVLSTPASSVPSSAPSDAGKPDSPTR
ETLEDVGDVSLAKHIVAQEMGVDIAEVTDDADLAEMGMDSLMSLTILGELREKTGIDLPSTFLTTNPTMK
DIDNALGMRPKPKAAPKPAAPKAAAPSSSKKTDMNEVSARLSALNNNTDISRYPNATSVLLQGNPKQATK
KIFFLPDGSGSATSYVSIPNLGPDVCAYGLNCPFMKNPEQWQCGIEISALVYLAEIKRRQPQGPYIIGGW
SAGGVIAYSVAQALLAANEGVEKLLLLDSPCPVNLAPLPARLHNFFNQIGLLGTGDPAKTPKWLLPHFSA
AIRSLSDYDPKPSLRPIPTYAIWCREGVAGNPGDPRPPPAEEEDPAPMTWLLEHRTNFKDNGWAQLCGDS
MKFGVMGGHHFSMMKPPHADDLGNLIREGLDWQP


>PKSI_1_6651
MSNVLLFGDQTAEQYQLLNKIVLRKENALVITFVERCAKALREETNALPRSQRNAVPDFLTVNDLKEAYH
QKGVKVPMVESALVTIAQIGHYIGYFSEHSAEQPSATNTRALGLCTGLLAAAAAVASKTVEELVLVGVEF
VRLSFRSGAAVDAARTALCQTGDDNAPWSTIVTGTTEASAKEALAKFHEEKGIPQTSHAYISAVSVMAIT
VSGPPTTVKRFFEESPALSKNHRVPIPVYGPYHAEHLFGETEINKIASASILEGLKQHQPVSLVHSAATG
KALVAENAAELAKLVLAEMLQLPVRWDHLLEEAVSQITSKKAPAKIWAMGVSNVANSLVSALKAGGQTDV
STVDQSTWTENEPDTHGRTQNDKVAIVGMAGRFPNSADHEALWELLMKGLDVHRRIPKDRFDADTHVDPS
GKGKNKSHTPFGCFIDEPGFFDPRFFNMSPREAAQTDPMGRLALVTAYEALEMSGYVPNRTPSTKLHRIG
TFYGQTSDDWREINAAENVDTYFITGGVRAFAPGRINYYFKFSGPSYSVDTACSSSLAAIQLACTSLWAG
DCDTACAGGLNVLTNPDIFSGLSKGQFLSKTGSCKTYDNNADGYCRGDAVGTVILKRYEDAIADKDNILG
CILGAATNHSAEAVSITHPHAGAQEFLYKRVLANAGVDAHEISYVEMHGTGTQAGDGIEMTSVTNVFAPR
HRQRRDDQPVYLGAIKANVGHAEAASGINSLAKVLLMMKHNKIPANVGIKGEMNKTFPADLKDRKVNISQ
KAVDWPRNGKEKRKVFLNNFSAAGGNTALLLEDGPAYEAPTATDPRGTVPVTVTARSISALKRNIANLQK
YVSENPSTTLTSMSYTLTARRIQHNYRVAFPLDQINKFSDALQAQVKESYSPVPNAPTRVAFCFTGQGSQ
YTGLGQKLYNDLKSFRDDIDQLDHLARVQGLPSFLEIVQGADVQTLSPVKVQLGMACIQVALARMWAAWG
ITPAAVIGHSLGEYAALHVAGVISASDMVLLVGRRAELLVRDCTPHTHGMLAVKGGAEAIRNTLGNKMTE
IACINGPEETVLCGSGDVVGAANETLAAKGFKATKLNVPFAFHSAQVDPILEQFKKIAASVTYNKPAAPV
LSPLEGDIIREAGKINPEYLARHARETVNFWTALTAGQKEKVFDEKTAWLEVGAHPVCSGMVKASIGATT
TAPSLRRGEDAWKTISNSMCTLFTAGVNFNFDEFHKEFNDAQEMYTLPTYSFDNKKYWLDYHNDWTLRKG
EPAQTKEVIVEKPVASASAPAVEMPAKRLSTSCQRVIAENFSGNNGSVTVQSSLADPKLYPVVCGHMVNN
AALCPSSLYADMALTISDYIWKQMRPGTETPGYNVCNMEVPKPLIAQIPQPAEGQHIQLEANADLDSGIV
KLNFRSVKPDGQKLQDHAHCIVRLEDKAAWEDEWSRYNYMVQAQMELLQHKTLNGGAHKVQRGMAYKLFK
ALVNYDEKYRAMAEVVLASGQTEASAMLDFPTKPEDGDFYCPPYHIDGSCHISGFIVNASDLLDSEQNVY
VSHGWGAMKFSRPLTAGMKLRNYVRMQPQPNNVSKGDVYIMEGDQIVAVCEGIKFQQIPRRVLNTFLPPN
KGSGPASAAKPAAAPVAAARPAPAAAPIKTAPAPAKAAPAPAPAAPKAAPKPKKAAAPKKPAGGLTAKVM
KILAKETEVDEGELVDEAQFENLGVDSLLSLTISAVFREELDMDISSTLFTDYPTVGDMKKYFAQFDNGS
STSSSAEEEDSDEDSIPPTDAATPMDDLSTPASSVPSSAPSDAGKPDSPTRETLEDVGDVSLAKHIVAQE
MGVDIAEVTDDADLAEMGMDSLMSLTILGELREKTGIDLPSTFLTTNPTMKDIDNALGMRPKPKAAPKSA
APKAAAPSSSKKTDMNEVSARLSALNNNTDISRYPNATSVLLQGNPKQATKKIFFLPDGSGSATSYV
SIPNLGPDVCAYGLNCPFMKNPEQWQCGIEISALVYLAEIKRRQPQGPYIIGGWSAGGVIAYSVAQALLA
ANEGVEKLLLLDSPCPVNLAPLPARLHNFFNQIGLLGTGDPAKTPKWLLPHFSAAIRSLSDYDPKPSLRP
IPTYAIWCREGVAGNPGDPRPPPAEEEDPAPMTWLLEHRTNFKDNGWAQLCGDSMKFGVMGGHHFSMMKP
PHADDLGNLIREGLDWQP


>PKSI_1_225

MSNVLLFGDQTAEQYPLLNKIVLRKENALVITFIERCAKALREETNALPRSQRNAVPDFLTVNDLKEAYH
QKGVKVPMVESALVTIAQIGHYIGYFSEHSAEQPSATNTRALGLCTGLLAAAAVVASKTVEELVLVGVEF
VRLSFRSGAAVDAARTALCQTGDDNSPWSTIVTGTTEAAAKEALAKFHEEKSIPQTSHAYISAVSVMAIT
VSGPPTTVKRFFEESSALSKNHRVPIPVYGPYHAEHLFGETEINKIASASILEGLKQHQPVSLVHSAATG
KALVAENAAELAKLVLAEMLQHPVRWDHLLEEAVSQITSKKAPAKIWAMGVSNVANSLVSALKAGGQTDV
STIDQSTWTENEPDTHGRTQNDKVAIVGMAGRFPNSADHEALWELLMKGLDVHRRIPKDRFDADTHVDPS
GKGKNKSHTPFGCFIDEPGFFDPRFFNMSPREAAQTDPMGRLALVTAYEALEMSGYVPNRTPSTKLHRIG
TFYGQTSDDWREINAAENVDTYFITGGVRAFAPGRINYYFKFSGPSYSVDTACSSSLAAIQLACTSLWAG
DCDTACAGGLNVLTNPDIFSGLSKGQFLSKTGSCKTYDNNADGYCRGDAVGTVILKRYEDAIADKDNILG
CILGAATNHSAEAVSITHPHAGAQEFLYKRVLANAGVDAHEISYVEMHGTGTQAGDGIEMTSVTNVFAPR
HRQRRDDQPVYLGAIKANVGHAEAASGINSLAKVLLMMKHNKIPANVGIKGEMNKTFPADLKDRKVNISQ
KAVDWPRNGKEKRKVFLNNFSAAGGNTALLLEDGPAYEAPTATDPRGTVPVTVTARSISALKRNIANLQK
YVSENPSTTLTSMSYTLTARRIQHNYRVAFPLDQINKFSDALQAQVKESYSPVPNAPTRVAFCFTGQGSQ
YTGLGQKLYNDLKSFRDDIDQLDHLARVQGLPSFLEIVQGADVQTLSPVKVQLGMACIQVALARMWAAWG
ITPAAVIGHSLGEYAALHVAGVISASDMVLLVGRRAELLVRDCTPHTHGMLAVKGGAEAIRNTLGSKMTE
IACINGPEETVLCGSGDVVGAANETLAAKGFKATKLNVPFAFHSAQVDPILEQFKKIAASVTYNKPAVPV
LSPLEGDIIREAGKINPEYLARHARETVNFWTALTAGQKEKVFDEKTAWLEVGAHPVCSGMVKASIGATT
TAPSLRRGEDAWKTISNSMCTLFTAGVNFNFDEFHKEFNDAQEMYTLPTYSFDNKKYWLDYHNDWTLRKG
EPAQTKEVIVEKPVASASAPAVEMPAKRLSTSCQRVIAENFSSNNGSVTVQSSLADPKLYPVVCGHMVNN
AALCPSSLYADMALTISDYIWKQMRPGTETPGYNVCNMEVPKPLIAQIPQPAEGQHIQLEANADLDSGIVKLNFRSVKPDGQKLQDHAHCIVRLEDRAAWEDEWSRYNYMVQAQMELLQHKTLNGGAHKVQRGMAYKLFKALVNYDEKYRAMAEVVLASGQTEASAVLDFPTKPEDGDFYCPPYHIDGSCHISGFIVNASDLLDSEQNVYVSHGWGAMKFS
RPLTAGMKLRNYVRMQPQPNNVSKGDVYIMEGDQIVAVCEGIKFQQIPRRVLNTFLPPNKGSGPASAAKP
AAAPVAAARPAPAAAPIKTAPAPAKAAPAPAPAAPKAAPKPKKAAAPKKAAGGLTAKVMKILAKETEVDE
GELVDEAQFENLGVDSLLSLTISAVFREELDMDISSTLFTDYPTVGDMKKYFAQFDNGSSTSSSAEEEDS
DEDSIPPTDAATPMDDLSTPASSVPSSAPSDAGKPDSPTRETLEDVGDVSLAKHIVAQEMGVDIAEVTDD
ADLAEMGMDSLMSLTILGELREKTGIDLPSTFLTTNPTMKDIDNALGMRPKPKAAPKPAAPKAAAPSSSK
KTDMNEVSARLSALNNNTDISRYPNATSVLLQGNPKQATKKIFFLPDGSGSATSYVSIPNLGPDVCAYGVNCP
FMKNPEQWQCGIEISALVYLAEIKRRQPQGPYIIGGWSAGGVIAYSVAQALLAANEGVEKLLLLDSPCPV
NLAPLPARLHNFFNEIGLLGTGDPAKTPKWLLPHFSAAIRSLSDYDPKPSLRPIPTYAIWCREGVAGNPG
DPRPPPAEEEDPAPMTWLLEHRTNFKDNGWAQLCGDSMKFGVMGGHHFSMMKPPHADDLGNLIREGLDWQ
P


>PKSI_1_11650
MSNVLLFGDQTAEQYQLLNKIVLRKENALVITFVERCAKALREETNALPRSQRNAVPDFLTVNDLKEAYH
QKGVKVPMVESALVTIAQIGHYIGYFSEHSAEQPSATNTRALGLCTGLLAAAAAVASKTVEELVLVGVEF
VRLSFRSGAAVDAARTALCQTGDDNAPWSTIVTGTTEASAKEALAKFHEEKGIPQTSHAYISAVSVMAIT
VSGPPTTVKRFFEESPALSKNHRVPIPVYGPYHAEHLFGETEINKIASASILEGLKQHQPVSLVHSAATG
KALVAENAAELAKLVLAEMLQLPVRWDHLLEEAVSQITSKKAPAKIWAMGVSNVANSLVSALKAGGQTDV
STVDQSTWTENEPDTHGRTQNDKVAIVGMAGRFPNSADHEALWELLMKGLDVHRRIPKDRFDADTHVDPS
G
NKSHTPFGCFIDEPGFFDPRFFNMSPREAAQTDPMGRLALVTAYEALEMSGYVPNRTPSTKLHRVGTFYG
QTSDDWREINAAENVDTYFITGGVRAFAPGRINYYFKFSGPSYSVDTACSSSLAAIQLACTSLWAGDCDT
ACAGGLNVLTNPDIFSGLSKGQFLSKTGSCKTYDNNADGYCRGDAVGTVILKRYEDAIADKDNILGCILG
AATNHSAEAVSITHPHAGAQEFLYKRVLANAGVDAHEISYVEMHGTGTQAGDGIEMTSVTNVFAPRHRQR
RDDQPVYLGAIKANVGHAEAASGINSLAKVLLMMKHNKIPANVGIKGEMNKTFPADLKDRKVNISQKAVD
WPRNGKEKRKVFLNNFSAAGGNTALLLEDGPAYEAPTATDPRGTVPVTVTARSISALKRNIANLQKYVSE
NPSTTLTSMSYTLTARRIQHNYRVAFPLDQINKFSDALQAQVKESYSPVPNAPTRVAFCFTGQGSQYTGL
GQKLYNDLKSFRDDIDQLDHLARVQGLPSFLEIVQGADVQTLSPVKVQLGMACIQVALARMWAAWGITPA
AVIGHSLGEYAALHVAGVISASDMVLLVGRRAELLVRDCTPHTHGMLAVKGGAEAIRNTLGNKMTEIACI
NGPEETVLCGSGDVVGAANETLAAKGFKATKLNVPFAFHSAQVDPILEQFKKIAASVTYNKPAVPVLSPL
EGDIIREAGKINPEYLARHARETVNFWTALTAGQKEKVFDEKTAWLEVGAHPVCSGMVKASIGATTTAPS
LRRGEDAWKTISNSMCTLFTAGVNFNFDEFHKEFNDAQEMYTLPTYSFDNKKYWLDYHNDWTLRKGEPAQ
TKEVIVEKPVASASAPAVEMPAKRLSTSCQRVISENFSGNNGSVTVQSSLADPKLYPVVCGHMVNNAALC
PSSLYADMALTISDYIWKQMRPGTETPGYNVCNMEVPKPLIAQIPQPAEGQHIQLEANADLDSGIVKLNF
RSVKPDGQKLQDHAHCIVRLEDKAAWEDEWSRYNYMVQAQMELLQHKTLNGGAHKVQRGMAYKLFKALVN
YDEKYRAMAEVVLASGQTEASAMLDFPTKPEDGDFYCPPYHIDGSCHISGFIVNASDLLDSEQNVYVSHG
WGAMKFSRPLTAGMKLRNYVRMQPQPNNVSKGDVYIMEGDQI VAVCEGIKFQQIPRRVLNTFLPPNKGSGPASAAKPAAAPVAAARPAPAAAPIKTAPAPAKAAPAPAPAAPKAAPKPKKAAAPKKPAGGLTAKVMKILAKETEVDEGELVDEAQFENLGVDSLLSLTISAVFREELDMDISSTLFTDYPTVGDMKKYFAQFDNGSSTSSSAEEEDSDEDSIPPTDAATPMDDLSTPASSVPSSAPSDAGKPDSPTRETLEDVGDVSLAKHIVAQEMGVDIAEVTDDADLAEMGMDSLMSLTILGELREKTGIDLPSTFLTTNPTMKDIDNALGMRPKPKAAPKSAAPKAAAPSSSKKTDMNEVSARLSALNNNTDISRYPNATSVLLQGNPKQATKKIFFLPDGSGSATSYVSIPNLGP
MKNPEQWQCGIEISALVYLAEIKRRQPQGPYIIGGWSAGGVIAYSVAQALLAANEGVEKLLLLDSPCPVN
LAPLPARLHNFFNQIGLLGTGDPAKTPKWLLPHFSAAIRSLSDYDPKPSLRPIPTYAIWCREGVAGNPGD
PRPPPAEEEDPAPMTWLLEHRTNFKDNGWAQLCGDSMKFGVMGGHHFSMMKPPHADDLGNLIREGLDWQP

>PKSI_1_10919
MSNVLLFGDQTAEQYPLLNKIVLRKENALVITFIERCAKALREETNALPRSQRNAVPDFLTVNDLKEAYH
QKGVKVPMVESALVTIAQIGHYIGYFSEHSAEQPSATNTRALGLCTGLLAAAAVVASKTVEELVLVGVEF
VRLSFRSGAAVDAARTALCQTGDDNSPWSTIVTGTTEAAAKEALAKFHEEKSIPQTSHAYISAVSVMAIT
VSGPPTTVKRFFEESSALSKNHRVPIPVYGPYHAEHLFGETEINKIASASILEGLKQHQPVSLVHSAATG
KALVAENAAELAKLVLAEMLQHPVRWDHLLEEAVSQITSKKAPAKIWAMGVSNVANSLVSALKAGGQTDV
STIDQSTWTENEPDTHGRTQNDKVAIVGMAGRFPNSADHEALWELLMKGLDVHRRIPKDRFDADTHVDPS
GKGKNKSHTPFGCFIDEPGFFDPRFFNMSPREAAQTDPMGRLALVTAYEALEMSGYVPNRTPSTKLHRIG
TFYGQTSDDWREINAAENVDTYFITGGVRAFAPGRINYYFKFSGPSYSVDTACSSSLAAIQLACTSLWAG
DCDTACAGGLNVLTNPDIFSGLSKGQFLSKTGSCKTYDNNADGYCRGDAVGTVILKRYEDAIADKDNILG
CILGAATNHSAEAVSITHPHAGAQEFLYKRVLANAGVDAHEISYVEMHGTGTQAGDGIEMTSVTNVFAPR
HRQRRDDQPVYLGAIKANVGHAEAASGINSLAKVLLMMKHNKIPANVGIKGEMNKTFPADLKDRKVNISQ
KAVDWPRNGKEKRKVFLNNFSAAGGNTALLLEDGPAYEAPTATDPRGTVPVTVTARSISALKRNIANLQK
YVSENPSTTLTSMSYTLTARRIQHNYRVAFPLDQINKFSDALQAQVKESYSPVPNAPTRVAFCFTGQGSQ
YTGLGQKLYNDLKSFRDDIDQLDHLARVQGLPSFLEIVQGADVQTLSPVKVQLGMACIQVALARMWAAWG
ITPAAVIGHSLGEYAALHVAGVISASDMVLLVGRRAELLVRDCTPHTHGMLAVKGGAEAIRNTLGSKMTE
IACINGPEETVLCGSGDVVGAANETLAAKGFKATKLNVPFAFHSAQVDPILEQFKKIAASVTYNKPAVPV
LSPLEGDIIREAGKINPEYLARHARETVNFWTALTAGQKEKVFDEKTAWLEVGAHPVCSGMVKASIGATT
TAPSLRRGEDAWKTISNSMCTLFTAGVNFNFDEFHKEFNDAQEMYTLPTYSFDNKKYWLDYHNDWTLRKG
EPAQTKEVIVEKPVASASAPAVEMPAKRLSTSCQRVIAENFSSNNGSVTVQSSLADPKLYPVVCGHMVNN
AALCPSSLYADMALTISDYIWKQMRPGTETPGYNVCNM EVPKPLIAQIPQPAEGQHIQLEANADLDSGIVKLNFRSVKPDG
QKLQDHAHCIVRLEDRAAWEDEWSRYNYMVQAQMELLQHKTLNGGAHKVQRGMAYKLFKALVNYDEKYRA
MAEVVLASGQTEASAVLDFPTKPEDGDFYCPPYHIDGSCHISGFIVNASDLLDSEQNVYVSHGWGAMKFS
RPLTAGMKLRNYVRMQPQPNNVSKGDVYIMEGDQIVAVCEGIKFQQIPRRVLNTFLPPNKGSGPASAAKP
AAAPVAAARPAPAAAPIKTAPAPAKAAPAPAPAAPKAAPKPKKAAAPKKAAGGLTAKVMKILAKETEVDE
GELVDEAQFENLGVDSLLSLTISAVFREELDMDISSTLFTDYPTVGDMKKYFAQFDNGSSTSSSTEEEDS
DEDSIPPTDAATPMDDLSTPASSVGSSAPSDAGKPDSPTRETLEDVGDVSLAKHIVAQEMGVDIAEVTDD
ADLAEMGMDSLMSLTILGELREKTGIDLPSTFLTTNPTMKDIDNALGMRPKPKAVPKPAAPKAAAPSSSK
KTD MNEVSARLSALNNNTDISRYPNATSVLLQGNPKQATKKIFFLPDGSGSATSYVSIPNLGPDVCAYGLNCP
FMKNPEQWQCGIEISALVYLAEIKRRQPQGPYIIGGWSAGGVIAYSVAQALLAANEGVEKLLLLDSPCPV
NLAPLPARLHNFFNEIGLLGTGDPAKTPKWLLPHFSAAIRSLSDYDPKPSLRPIPTYAIWCREGVAGNPG
DPRPPPAEEEDPAPMTWLLEHRTNFKDNGWAQLCGDSMKFGVMGGHHFSMMKPPHADDLGNLIREGLDWQ
P

>PKSI_1_10843
MSNVLLFGDQTAEQYPLLNKIVLRKENALVITFIERCAKALREETNALPRSQRNAVPDFLTVNDLKEAYH
QKGVKVPMVESALVTIAQIGHYIGYFSEHSAEQPSATNTRALGLCTGLLAAAAVVASKTVEELVLVGVEF
VRLSFRSGAAVDAARTALCQTGDDNSPWSTIVTGTTEAAAKEALAKFHEEKSIPQTSHAYISAVSVMAIT
VSGPPTTVKRFFEESSALSKNHRVPIPVYGPYHAEHLFGETEINKIASASILEGLKQHQPVSLVHSAATG
KALVAENAAELAKLVLAEMLQHPVRWDHLLEEAVSQITSKKAPAKIWAMGVSNVANSLVSALKAGGQTDV
STIDQSTWTENEPDTHGRTQNDKVAIVGMAGRFPNSADHEALWELLMKGLDVHRRIPKDRFDADTHVDPS
GKGKNKSHTPFGCFIDEPGFFDPRFFNMSPREAAQTDPMGRLALVTAYEALEMSGYVPNRTPSTKLHRIG
TFYGQTSDDWREINAAENVDTYFITGGVRAFAPGRINYYFKFSGPSYSVDTACSSSLAAIQLACTSLWAG
DCDTACAGGLNVLTNPDIFSGLSKGQFLSKTGSCKTYDNNADGYCRGDAVGTVILKRYEDAIADKDNILG
CILGAATNHSAEAVSITHPHAGAQEFLYKRVLANAGVDAHEISYVEMHGTGTQAGDGIEMTSVTNVFAPR
HRQRRDDQPVYLGAIKANVGHAEAASGINSLAKVLLMMKHNKIPANVGIKGEMNKTFPADLKDRKVNISQ
KAVDWPRNGKEKRKVFLNNFSAAGGNTALLLEDGPAYEAPTATDPRGTVPVTVTARSISALKRNIANLQK
YVSENPSTTLTSMSYTLTARRIQHNYRVAFPLDQINKFSDALQAQVKESYSPVPNAPTRVAFCFTGQGSQ
YTGLGQKLYNDLKSFRDDIDQLDHLARVQGLPSFLEIVQGADVQTLSPVKVQLGMACIQVALARMWAAWG
ITPAAVIGHSLGEYAALHVAGVISASDMVLLVGRRAELLVRDCTPHTHGMLAVKGGAEAIRNTLGSKMTE
IACINGPEETVLCGSGDVVGAANETLAAKGFKATKLNVPFAFHSAQVDPILEQFKKIAASVTYNKPAVPV
LSPLEGDIIREAGKINPEYLARHARETVNFWTALTAGQKEKVFDEKTAWLEVGAHPVCSGMVKASIGATT
TAPSLRRGEDAWKTISNSMCTLFTAGVNFNFDEFHKEFNDAQEMYTLPTYSFDNKKYWLDYHNDWTLRKG
EPAQTKEVIVEKPVASASAPAVEMPAKRLSTSCQRVIAENFSSNNGSVTVQSSLADPKLYPVVCGHMVNN
AALCPSSLYADMALTISDYIWKQMRPGTETPGYNV CNMEVPKPLIAQIPQPAEGQHIQLEANADLDSGIVKLNFRSVKPDG
QKLQDHAHCIVRLEDRAAWEDEWSRYNYMVQAQMELLQHKTLNGGAHKVQRGMAYKLFKALVNYDEKYRA
MAEVVLASGQTEASAVLDFPTKPEDGDFYCPPYHIDGSCHISGFIVNASDLLDSEQNVYVSHGWGAMKFS
RPLTAGMKLRNYVRMQPQPNNVSKGDVYIMEGDQIVAVCEGIKFQQIPRRVLNTFLPPNKGSGPASAAKP
AAAPVAAARPAPAAAPIKTAPAPAKAAPAPAPAAPKAAPKPKKAAAPKKAAGGLTAKVMKILAKETEVDE
GELVDEAQFENLGVDSLLSLTISAVFREELDMDISSTLFTDYPTVGDMKKYFAQFDNGSSTSSSAEEEDS
DEDSIPPTDAATPMDDLSTPASSVPSSAPSDAGKPDSPTRETLEDVGDVSLAKHIVAQEMGVDIAEVTDD
ADLAEMGMDSLMSLTILGELREKTGIDLPSTFLTTNPTMKDIDNALGMRPKPKAAPKPAAPKAAAPSSSK
KTDMNEVSARLSALNNNTDISRYPNATSVLLQGNPKQATKKIFFLPDGSGSATSYVSIPNLGPDVCAYGLNCP
FMKNPEQWQCGIEISALVYLAEIKRRQPQGPYIIGGWSAGGVIAYSVAQALLAANEGVEKLLLLDSPCPV
NLAPLPARLHNFFNEIGLLGTGDPAKTPKWLLPHFSAAIRSLSDYDPKPSL RPIPTYAIWCREGVAGNPGDPRPPPAEEEDPAPMT
WLLEHRTNFKDNGWAQLCGDSMKFGVMGGHHFSMMKPPHADDLGNLIREGLDWQP

>PKSI_1_11540
MSNVLLFGDQTAEQYQLLNKIVLRKENALVITFVERCAKALREETNALPRSQRNAVPDFLTVNDLKEAYH
QKGVKVPMVESALVTIAQIGHYIGYFSEHSAEQPSATNTRALGLCTGLLAAAAAVASKTVEELVLVGVEF
VRLSFRSGAAVDAARTALCQTGDDNAPWSTIVTGTTEASAKEALAKFHEEKGIPQTSHAYISAVSVMAIT
VSGPPTTVKRFFEESPALSKNHRVPIPVYGPYHAEHLFGETEINKIASASILEGLKQHQPVSLVHSAATG
KALVAENAAELAKLVLAEMLQLPVRWDHLLEEAVSQITSKKAPAKIWAMGVSNVANSLVSALKAGGQTDV
STVDQSTWTENEPDTHGRTQNDKVAIVGMAGRFPNSADHEALWELLMKGLDVHRRIPKDRF DADTHVDPSGKGKNKSHTPFGCFIDEPGFFDPRFFNMSPR
EAAQTDPMGRLALVTAYEALEMSGYVPNRTPSTKLHRIGTFYGQTSDDWREINAAENVDTYFITGGVRAF
APGRINYYFKFSGPSYSVDTACSSSLAAIQLACTSLWAGDCDTACAGGLNVLTNPDIFSGLSKGQFLSKT
GSCKTYDNNADGYCRGDAVGTVILKRYEDAIADKDNILGCILGAATNHSAEAVSITHPHAGAQEFLYKRV
LANAGVDAHEISYVEMHGTGTQAGDGIEMTSVTNVFAPRHRQRRDDQPVYLGAIKANVGHAEAASGINSL
AKVLLMMKHNKIPANVGIKGEMNKTFPADLKDRKVNISQKAVDWPRNGKEKRKVFLNNFSAAGGNTALLL
EDGPAYEAPTATDPRGTVPVTVTARSISALKRNIANLQKYVSENPSTTLTSMSYTLTARRIQHNYRVAFP
LDQINKFSDALQAQVKESYSPVPNAPTRVAFCFTGQGSQYTGLGQKLYNDLKSFRDDIDQLDHLARVQGL
PSFLEIVQGADVQTLSPVKVQLGMACIQVALARMWAAWGITPAAVIGHSLGEYAALHVAGVISASDMVLL
VGRRAELLVRDCTPHTHGMLAVKGGAEAIRNTLGNKMTEIACINGPEETVLCGSGDVVGAANETLAAKGF
KATKLNVPFAFHSAQVDPILEQFKKIAASVTYNKPAVPVLSPLEGDIIREAGKINPEYLARHARETVNFW
TALTAGQKEKVFDEKTAWLEVGAHPVCSGMVKASIGATTTAPSLRRGEDAWKTISNSMCTLFTAGVNFNF
DEFHKEFNDAQEMYTLPTYSFDNKKYWLDYHNDWTLRKGEPAQTKEVIVEKPVASASAPAVEMPAKRLST
SCQRVISENFSGNNGSVTVQSSLADPKLYPVVCGHMVNNAALCPSSLYADMALTISDYIWKQMRPGTETP
GYNVCNMEVPKPLIAQIPQPAEGQHIQLEANADLDSGIVKLNFRSVKPDGQKLQDHAHCIVRLEDKAAWE
DEWSRYNYMVQAQMELLQHKTLNGGAHKVQRGMAYKLFKALVNYDEKYRAMAEVVLASGQTEASAMLDFP
TKPEDGDFYCPPYHIDGSCHISGFIVNASDLLDSEQNVYVSHGWGAMKFSRPLTAGMKLRNYVRMQ PQPNNVSKGDVYIMEGDQIVAVCEGIKF
QQIPRRVLNTFLPPNKGSGPASAAKPAAAPVAAARPAPAAAPIKTAPAPAKAAPAPAPAAPKAAPKPKKA
AAPKKPAGGLTAKVMKILAKETEVDEGELVDEAQFENLGVDSLLSLTISAVFREELDMDISSTLFTDYPT
VGDMKKYFAQFDNGSSTSSSAEEEDSDEDSIPPTDAATPMDDLSTPASSVPSSAPSDAGKPDSPTRETLE
DVGDVSLAKHIVAQEMGVDIAEVTDDADLAEMGMDSLMSLTILGELREKTGIDLPSTFLTTNPTMKDIDN
ALGMRPKPKAAPKSAAPKAAAPSSSKKTDMNEVSARLSALNNNTDISRYPNATSVLLQGNPKQATKKIFF
LPDGSGSATSYVSIPNLGPDVCAYGLNCPF MKNPEQWQCGIEISALVYLAEIKRRQPQGPYIIGGWSAGGVIAYSVAQALLAANEGVEKLLLLDSPCPVN
LAPLPARLHNFFNQIGLLGTGDPAKTPKWLLPHFSAAIRSLSDYDPKPSLRPIPTYAIWCREGVAGNPGD
PRPPPAEEEDPAPMTWLLEHRTNFKDNGWAQLCGDSMKFGVMGGHHFSMMKPPHADDLGNLIREGLDWQP


>PKSI_1_10819
MSNVLLFGDQTAEQYPLLNKIVLRKENALVITFIERCAKALREETNALPRSQRNAVPDFLTVNDLKEAYH
QKGVKVPMVESALVTIAQIGHYIGYFSEHSAEQPSATNTRALGLCTGLLAAAAVVASKTVEELVLVGVEF
VRLSFRSGAAVDAARTALCQTGDDNSPWSTIVTGTTEAAAKEALAKFHEEKSIPQTSHAYISAVSVMAIT
VSGPPTTVKRFFEESSALSKNHRVPIPVYGPYHAEHLFGETEINKIASASILEGLKQHQPVSLVHSAATG
KALVAENAAELAKLVLAEMLQHPVRWDHLLEEAVSQITSKKAPAKIWAMGVSNVANSLVSALKAGGQTDV
STIDQSTWTENEPDTHGRTQNDKVAIVGMAGRFPNSADHEALWELLMKGLDVHRRIPKDRFDADTHVDPS
GKGKNKSHTPFGCFIDEPGFFDPRFFNMSPREAAQTDPMGRLALVTAYEALEMSGYVPNRTPSTKLHRIG
TFYGQTSDDWREINAAENVDTYFITGGVRAFAPGRINYYFKFSGPSYSVDTACSSSLAAIQLACTSLWAG
DCDTACAGGLNVLTNPDIFSGLSKGQFLSKTGSCKTYDNNADGYCRGDAVGTVILKRYEDAIADKDNILG
CILGAATNHSAEAVSITHPHAGAQEFLYKRVLANAGVDAHEISYVEMHGTGTQAGDGIEMTSVTNVFAPR
HRQRRDDQPVYLGAIKANVGHAEAASGINSLAKVLLMMKHNKIPANVGIKGEMNKTFPADLKDRKVNISQ
KAVDWPRNGKEKRKVFLNNFSAAGGNTALLLEDGPAYEAPTATDPRGTVPVTVTARSISALKRNIANLQK
YVSENPSTTLTSMSYTLTARRIQHNYRVAFPLDQINKFSDALQAQVKESYSPVPNAPTRVAFCFTGQGSQ
YTGLGQKLYNDLKSFRDDIDQLDHLARVQGLPSFLEIVQGADVQTLSPVKVQLGMACIQVALARMWAAWG
ITPAAVIGHSLGEYAALHVAGVISASDMVLLVGRRAELLVRDCTPHTHGMLAVKGGAEAIRNTLGSKMTE
IACINGPEETVLCGSGDVVGAANETLAAKGFKATKLNVPFAFHSAQVDPILEQFKKIAASVTYNKPAVPV
LSPLEGDIIREAGKINPEYLARHARETVNFWTALTAGQKEKVFDEKTAWLEVGAHPVCSGMVKASIGATT
TAPSLRRGEDAWKTISNSMCTLFTAGVNFNFDEFHKEFNDAQEMYTLPTYSFDNKKYWLDYHNDWTLRKG
EPAQTKEVIVEKPVASASAPAVEMPAKRLSTSCQRVIAENFSSNNGSVTVQSSLADPKLYPVVCGHMVNN
AALCPSSLYADMALTISDYIWKQMRPGTETPGYNVCNMEVPKPLIAQIPQPAEGQHIQLEANADLDSGIVKLNFRSVKPDG
QKLQDHAHCIVRLEDRAAWEDEWSRYNYMVQAQMELLQHKTLNGGAHKVQRGMAYKLFKALVNYDEKYRA
MAEVVLASGQTEASAVLDFPTKPEDGDFYCPPYHIDGSCHISGFIVNASDLLDSEQNVYVSHGWGAMKFS
RPLTAGMKLRNYVRMQPQPNNVSKGDVYIMEGDQIVAVCEGIKFQQIPRRVLNTFLPPNKGSGPASAAKP
AAAPVAAARPAPAAAPIKTAPAPAKAAPAPAPAAPKAAPKPKKAAAPKKAAGGLTAKVMKILAKETEVDE
GELVDEAQFENLGVDSLLSLTISAVFREELDMDISSTLFTDYPTVGDMKKYFAQFDNGSSTSSSAEEEDS
DEDSIPPTDAATPMDDLSTPASSVPSSAPSDAGKPDSPTRETLEDVGDVSLAKHIVAQEMGVDIAEVTDD
ADLAEMGMDSLMSLTILGELREKTGIDLPSTFLTTNPTMKDIDNALGMRPKPKAAPKPAAPKAAAPSSSK
KTDMNEVSARLSALNNNTDISRYPNATSVLLQGNPKQATKKIFFLPDGSGSATSYVSIPNLG
PDVCAYGLNCPFMKNPEQWQCGIEISALVYLAEIKRRQPQGPYIIGGWSAGGVIAYSVAQALLAANEGVE
KLLLLDSPCPVNLAPLPARLHNFFNEIGLLGTGDPAKTPKWLLPHFSAAIRSLSDYDPKPSLRPIPTYAI
WCREGVAGNPGDPRPPPAEEEDPAPMTWLLEHRTNFKDNGWAQLCGDSMKFGVMGGHHFSMMKPPHADDL
GNLIREGLDWQP


>PKSI_1_8422
MSNVLLFGDQTAEQYPLLNKIVLRKENALVITFIERCAKALREETNALPRSQRNAVPDFLTVNDLKEAYH
QKGVKVPMVESALVTIAQIGHYIGYFSEHSAEQPSATNTRALGLCTGLLAAAAVVASKTVEELVLVGVEF
VRLSFRSGAAVDAARTALCQTGDDNSPWSTIVTGTTEAAAKEALAKFHEEKSIPQTSHAYISAVSVMAIT
VSGPPTTVKRFFEESSALSKNHRVPIPVYGPYHAEHLFGETEINKIASASILEGLKQHQPVSLVHSAATG
KALVAENAAELAKLVLAEMLQHPVRWDHLLEEAVSQITSKKAPAKIWAMGVSNVANSLVSALKAGGQTDV
STIDQSTWTENEPDTHGRTQNDKVAIVGMAGRFPNSADHEALWELLMKGLDVHRRIPKDRFDADTHVDPS
GKGKNKSHTPFGCFIDEPGFFDPRFFNMSPREAAQTDPMGRLALVTAYEALEMSGYVPNRTPSTKLHRIG
TFYGQTSDDWREINAAENVDTYFITGGVRAFAPGRINYYFKFSGPSYSVDTACSSSLAAIQLACTSLWAG
DCDTACAGGLNVLTNPDIFSGLSKGQFLSKTGSCKTYDNNADGYCRGDAVGTVILKRYEDAIADKDNILG
CILGAATNHSAEAVSITHPHAGAQEFLYKRVLANAGVDAHEISYVEMHGTGTQAGDGIEMTSVTNVFAPR
HRQRRDDQPVYLGAIKANVGHAEAASGINSLAKVLLMMKHNKIPANVGIKGEMNKTFPADLKDRKVNISQ
KAVDWPRNGKEKRKVFLNNFSAAGGNTALLLEDGPAYEAPTATDPRGTVPVTVTARSISALKRNIANLQK
YVSENPSTTLTSMSYTLTARRIQHNYRVAFPLDQINKFSDALQAQVKESYSPVPNAPTRVAFCFTGQGSQ
YTGLGQKLYNDLKSFRDDIDQLDHLARVQGLPSFLEIVQGADVQTLSPVKVQLGMACIQVALARMWAAWG
ITPAAVIGHSLGEYAALHVAGVISASDMVLLVGRRAELLVRDCTPHTHGMLAVKGGAEAIRNTLGSKMTE
IACINGPEETVLCGSGDVVGAANETLAAKGFKATKLNVPFAFHSAQVDPILEQFKKIAASVTYNKPAVPV
LSPLEGDIIREAGKINPEYLARHARETVNFWTALTAGQKEKVFDEKTAWLEVGAHPVCSGMVKASIGATT
TAPSLRRGEDAWKTISNSMCTLFTAGVNFNFDEFHKEFNDAQEMYTLPTYSFDNKKYWLDYHNDWTLRKG
EPAQTKEVIVEKPVASASAPAVEMPAKRLSTSCQRVIAENFSSNNGSVTVQSSLADPKLYPVVCGHMVNN
AALCPSSLYADMALTISDYIWKQMRPGTETPGYNVCNMEVPKPLIAQIPQPAEGQHIQLEANADLDSGIVKLNFRSVKPDG
QKLQDHAHCIVRLEDKATWEDEWSRYNYMVQAQMELLQHKTLNGGAHKVQRGMAYKLFKALVNYDEKYRA
MAEVVLASGQTEASAVLDFPTKPEDGDFYCPPYHIDGSCHISGFIVNASDLLDSEQNVYVSHGWGAMKFS
RPLTAGMKLRNYVRMQPQPNNVSKGDVYIMEGDQIVAVCEGIKFQQIPRRVLNTFLPPNKGSGPASAAKP
AAAPVAAARPAPAAAPIKTAPAPAKAAPAPAPAAPKAAPKPKKAAAPKKAAGGLTAKVMKILAKETEVDE
GELVDEAQFENLGVDSLLSLTISAVFREELDMDISSTLFTDYPTVGDMKKYFAQFDNGSSTSSSAEEEDS
DEDSIPPTDAATPMDDLSTPASSVPSSAPSDAGKPDSPTRETLEDVGDVSLAKHIVAQEMGVDIAEVTDD
ADLAEMGMDSLMSLTILGELREKTGIDLPSTFLTTNPTMKDIDNALGMRPKPKAAPKPAAPKAAAPSSSK
KTDMNEVSARLSALNNNTDISRYPNATSVLLQGNPKQATKKIFFLPDGSGSATSYVSIPNLGPDVCAYGLNCP
FMKNPEQWQCGIEISALVYLAEIKRRQPQGPYIIGGWSAGGVIAYSVAQALLAANEGVEKLLLLDSPCPV
NLAPLPARLHNFFNEIGLLGTGDPAKTPKWLLPHFSAAIRSLSDYDPKPSLRPIPTYAIWCREGVAGNPG
DPRPPPAEEEDPAPMTWLLEHRTNFKDNGWAQLCGDSMKFGVMGGHHFSMMKPPHADDLGNLIREGLDWQ
P

>PKSI_1_4716
MSNVLLFGDQTAEQYPLLNKIVLRKENALVITFIERCAKALREETNALPRSQRNAVPDFLTVNDLKEAYH
QKGVKVPMVESALVTIAQIGHYIGYFSEHSAEQPSATNTRALGLCTGLLAAAAVVASKTVEELVLVGVEF
VRLSFRSGAAVDAARTALCQTGDDNSPWSTIVTGTTEAAAKEALAKFHEEKSIPQTSHAYISAVSVMAIT
VSGPPTTVKRFFEESSALSKNHRVPIPVYGPYHAEHLFGETEINKIASASILEGLKQHQPVSLVHSAATG
KALVAENAAELAKLVLAEMLQHPVRWDHLLEEAVSQITSKKAPAKIWAMGVSNVANSLVSALKAGGQTDV
STIDQSTWTENEPDTHGRTQNDKVAIVGMAGRFPNSADHEALWELLMKGLDVHRRIPKDRFDADTHVDPS
GKGKNKSHTPFGCFIDEPGFFDPRFFNMSPREAAQTDPMGRLALVTAYEALEMSGYVPNRTPSTKLHRIG
TFYGQTSDDWREINAAENVDTYFITGGVRAFAPGRINYYFKFSGPSYSVDTACSSSLAAIQLACTSLWAG
DCDTACAGGLNVLTNPDIFSGLSKGQFLSKTGSCKTYDNNADGYCRGDAVGTVILKRYEDAIADKDNILG
CILGAATNHSAEAVSITHPHAGAQEFLYKRVLANAGVDAHEISYVEMHGTGTQAGDGIEMTSVTNVFAPR
HRQRRDDQPVYLGAIKANVGHAEAASGINSLAKVLLMMKHNKIPANVGIKGEMNKTFPADLKDRKVNISQ
KAVDWPRNGKEKRKVFLNNFSAAGGNTALLLEDGPAYEAPTATDPRGTVPVTVTARSISALKRNIANLQK
YVSENPSTTLTSMSYTLTARRIQHNYRVAFPLDQINKFSDALQAQVKESYSPVPNAPTRVAFCFTGQGSQ
YTGLGQKLYNDLKSFRDDIDQLDHLARVQGLPSFLEIVQGADVQTLSPVKVQLGMACIQVALARMWAAWG
ITPAAVIGHSLGEYAALHVAGVISASDMVLLVGRRAELLVRDCTPHTHGMLAVKGGAEAIRNTLGSKMTE
IACINGPEETVLCGSGDVVGAANETLAAKGFKATKLNVPFAFHSAQVDPILEQFKKIAASVTYNKPAVPV
LSPLEGDIIREAGKINPEYLARHARETVNFWTALTAGQKEKVFDEKTAWLEVGAHPVCSGMVKASIGATT
TAPSLRRGEDAWKTISNSMCTLFTAGVNFNFDEFHKEFNDAQEMYTLPTYSFDNKKYWLDYHNDWTLRKG
EPAQTKEVIVEKPVASASAPAVEMPAKRLSTSCQRVIAENFSSNNGSVTVQSSLADPKLYPVVCGHMVNN
AALCPSSLYADMALTISDYIWKQMRPGTETPGYNVCNMEVPKPLIAQIPQPAEGQHIQLEANADLDSGIVKLNFRSVKPDG
QKLQDHAHCIVRLEDKATWEDEWSRYNYMVQAQMELLQHKTLNGGAHKVQRGMAYKLFKALVNYDEKYRA
MAEVVLASGQTEASAVLDFPTKPEDGDFYCPPYHIDGSCHISGFIVNASDLLDSEQNVYVSHGWGAMKFS
RPLTAGMKLRNYVRMQPQPNNVSKGDVYIMEGDQIVAVCEGIKFQQIPRRVLNTFLPPNKGSGPASAAKP
AAAPVAAARPAPAAAPIKTAPAPAKAAPAPAPAAPKAAPKPKKAAAPKKAAGGLTAKVMKILAKETEVDE
GELVDEAQFENLGVDSLLSLTISAVFREELDMDISSTLFTDYPTVGDMKKYFAQFDNGSSTSSSTEEEDS
DEDSIPPTDAATPMDDLSTPASSVGSSAPSDAGKPDSPTRETLEDVGDVSLAKHIVAQEMGVDIAEVTDD
ADLAEMGMDSLMSLTILGELREKTGIDLPSTFLTTNPTMKDIDNALGMRPKPKAVPKPAAPKAAAPSSSK
KTDMNEVSARLSALNNNTDISRYPNATSVLLQGNPKQATKKIFFLPDGSGSATSYVSIPNLG
PDVCAYGLNCPFMKNPEQWQCGIEISALVYLAEIKRRQPQGPYIIGGWSAGGVIAYSVAQALLAANEGVE
KLLLLDSPCPVNLAPLPARLHNFFNEIGLLGTGDPAKTPKWLLPHFSAAIRSLSDYDPKPSLRPIPTYAI
WCREGVAGNPGDPRPPPAEEEDPAPMTWLLEHRTNFKDNGWAQLCGDSMKFGVMGGHHFSMMKPPHADDL
GNLIREGLDWQP

>PKSI_1_291
MSNVLLFGDQTAEQYPLLNKIVLRKENALVITFIERCAKALREETNALPRSQRNAVPDFLTVNDLKEAYH
QKGVKVPMVESALVTIAQIGHYIGYFSEHSAEQPSATNTRALGLCTGLLAAAAVVASKTVEELVLVGVEF
VRLSFRSGAAVDAARTALCQTGDDNSPWSTIVTGTTEAAAKEALAKFHEEKSIPQTSHAYISAVSVMAIT
VSGPPTTVKRFFEESSALSKNHRVPIPVYGPYHAEHLFGETEINKIASASILEGLKQHQPVSLVHSAATG
KALVAENAAELAKLVLAEMLQHPVRWDHLLEEAVSQITSKKAPAKIWAMGVSNVANSLVSALKAGGQTDV
STIDQSTWTENEPDTHGRTQNDKVAIVGMAGRFPNSADHEALWELLMKGLDVHRRIPKDRFDADTHVDPS
GKGKNKSHTPFGCFIDEPGFFDPRFFNMSPREAAQTDPMGRLALVTAYEALEMSGYVPNRTPSTKLHRIG
TFYGQTSDDWREINAAENVDTYFITGGVRAFAPGRINYYFKFSGPSYSVDTACSSSLAAIQLACTSLWAG
DCDTACAGGLNVLTNPDIFSGLSKGQFLSKTGSCKTYDNNADGYCRGDAVGTVILKRYEDAIADKDNILG
CILGAATNHSAEAVSITHPHAGAQEFLYKRVLANAGVDAHEISYVEMHGTGTQAGDGIEMTSVTNVFAPR
HRQRRDDQPVYLGAIKANVGHAEAASGINSLAKVLLMMKHNKIPANVGIKGEMNKTFPADLKDRKVNISQ
KAVDWPRNGKEKRKVFLNNFSAAGGNTALLLEDGPAYEAPTATDPRGTVPVTVTARSISALKRNIANLQK
YVSENPSTTLTSMSYTLTARRIQHNYRVAFPLDQINKFSDALQAQVKESYSPVPNAPTRVAFCFTGQGSQ
YTGLGQKLYNDLKSFRDDIDQLDHLARVQGLPSFLEIVQGADVQTLSPVKVQLGMACIQVALARMWAAWG
ITPAAVIGHSLGEYAALHVAGVISASDMVLLVGRRAELLVRDCTPHTHGMLAVKGGAEAIRNTLGSKMTE
IACINGPEETVLCGSGDVVGAANETLAAKGFKATKLNVPFAFHSAQVDPILEQFKKIAASVTYNKPAVPV
LSPLEGDIIREAGKINPEYLARHARETVNFWTALTAGQKEKVFDEKTAWLEVGAHPVCSGMVKASIGATT
TAPSLRRGEDAWKTISNSMCTLFTAGVNFNFDEFHKEFNDAQEMYTLPTYSFDNKKYWLDYHNDWTLRKG
EPAQTKEVIVEKPVASASAPAVEMPAKRLSTSCQRVIAENFSSNNGSVTVQSSLADPKLYPVVCGHMVNN
AALCPSSLYADMALTISDYIWKQMRPGTETPGYNVCNMEVPKPLIAQIPQPAEGQHIQLEANADLDSGIVKLNFRSVKPDG
QKLQDHAHCIVRLEDRAAWEDEWSRYNYMVQAQMELLQHKTLNGGAHKVQRGMAYKLFKALVNYDEKYRA
MAEVVLASGQTEASAVLDFPTKPEDGDFYCPPYHIDGSCHISGFIVNASDLLDSEQNVYVSHGWGAMKFS
RPLTAGMKLRNYVRMQPQPNNVSKGDVYIMEGDQIVAVCEGIKFQQIPRRVLNTFLPPNKGSGPASAAKP
AAAPVAAARPAPAAAPIKTAPAPAKAAPAPAPAAPKAAPKPKKAAAPKKAAGGLTAKVMKILAKETEVDE
GELVDEAQFENLGVDSLLSLTISAVFREELDMDISSTLFTDYPTVGDMKKYFAQFDNGSSTSSSAEEEDS
DEDSIPPTDAATPMDDLSTPASSVPSSAPSDAGKPDSPTRETLEDVGDVSLAKHIVAQEMGVDIAEVTDD
ADLAEMGMDSLMSLTILGELREKTGIDLPSTFLTTNPTMKDIDNALGMRPKPKAAPKPAAPKAAAPSSSK
KTDMNEVSARLSALNNNTDISRYPNATSVLLQGNPKQATKKIFFLPDGSGSATSYVSIPNLGPDVCAYGLNCP
FMKNPEQWQCGIEISALVYLAEIKRRQPQGPYIIGGWSAGGVIAYSVAQALLAANEGVEKLLLLDSPCPV
NLAPLPARLHNFFNEIGLLGTGDPAKTPKWLLPHFSAAIRSLSDYDPKPSLRPIPTYAIWCREGVAGNPGDPRPPPAEEEDPAPMT
WLLEHRTNFKDNGWAQLCGDSMKFGVMGGHHFSMMKPPHADDLGNLIREGLDWQP

>PKSI_1_166
MSNVLLFGDQTAEQYQLLNKIVLRKENALVITFVERCAKALREETNALPRSQRNAVPDFLTVNDLKEAYH
QKGVKVPMVESALVTIAQIGHYIGYFSEHSAEQPSATNTRALGLCTGLLAAAAVVASKTVEELVLVGVEF
VRLSFRSGAAVDAARTALCQTGDGNAPWSTIVTGTTEASAKEALAKFHEEKGIPQTSHAYISAVSVMAIT
VSGPPTTVKRFFEESPALSKNHRVPIPVYGPYHAEHLFGETEINKIASASILEGLKQHQPVSLVHSAATG
KALVAENAAELAKLVLAEMLQLPVRWDHLLEEAVSQITSKKAPAKIWAMGVSNVANSLVSALKAGGQTDV
STVDQGTWTENEPDTHGRTQNDKVAIVGMAGRFPNSADHEALWELLMKGLDVHRRIPKDRFDADTHVDPS
GKGKNKSHTPFGCFIDEPGFFDPRFFNMSPREAAQTDPMGRLALVTAYEALEMSGYVPNRTPSTKLHRIG
TFYGQTSDDWREINAAENVDTYFITGGVRAFAPGRINYYFKFSGPSYSVDTACSSSLAAIQLACTSLWAG
DCDTACAGGLNVLTNPDIFSGLSKGQFLSKTGSCKTYDNNADGYCRGDAVGTVILKRYEDAIADKDNILG
CILGAATNHSAEAVSITHPHAGAQEFLYKRVLANAGVDAHEISYVEMHGTGTQAGDGIEMTSVTNVFAPR
HRQRRDDQPVYLGAIKANVGHAEAASGINSLAKVLLMMKHNKIPANVGIKGEMNKTFPADLKDRKVNISQ
KAVEWPRNGKEKRKVFLNNFSAAGGNTALLLEDGPAYEAPTATDPRGTVPVTVTARSISALKRNIANLQK
YVSENPSTTLTSMSYTLTARRIQHNYRVAFPLDQINKFSDALQAQVKESYSPVPNVPTRVAFCFTGQGSQ
YTGLGQKLYNDLKSFRDDIDQLDHLARVQGLPSFLEIVQGADVQTLSPVKVQLGMACIQVALARMWAAWG
ITPAAVIGHSLGEYAALHVAGVISASDMVLLVGRRAELLVRDCTPHTHGMLAVKGGAEAIRNTLGNKMTE
IACINGPEETVLCGSGDVVGAANETLAAKGFKATKLNVPFAFHSAQVDPILEQFKKIAASVTYNKPAVPV
LSPLEGDIIREAGKINPEYLARHARETVNFWTALTAGQKEKVFDEKTAWLEVGAHPVCSGMVKASIGATTTAPSLRRGEDAWKTISNSMCTLFTAGVNFNFDEFHKEFNDAQEMYTLPTYSF
DNKKYWLDYHNDWTLRKGEPAQTKEVIVEKPVASASAPAVEMPAKRLSTSCQRVIAENFSGNNGSVTVQS
SLADPKLYPVVCGHMVNNAALCPSSLYADMALTISDYIWKQMRPGTETPGYNVCNMEVPKPLIAQIPQPA
EGQHIQLEANADLDSGIVKLNFRSVKPDGQKLQDHAHCIVRLEDKAAWEDEWSRYNYMVQAQMELLQHKT
LNGGAHKVQRGMAYKLFKALVNYDEKYRAMAEVVLASGQTEASAMLDFPTKPEDGDFYCPPYHIDGSCHI
SGFIVNASDLLDSEQNVYVSHGWGAMKFSRPLTAGMKLRNYVRMQPQPNNVSKGDVYIMEGDQIVAVCEG
IKFQQIPRRVLNTFLPPNKGSGPASAAKPAAAPVAAARPAPAAAPIKSAPAPAKAAPAPAPAAPKAAPKP
KKAAAPKKPAGGLTAKVMKILAKETEVDEGELVDEAQFENLGVDSLLSLTISAVFREELDMDISSTLFTD
YPTVGDMKKYFAQFDNGSSTSSSAEEEDSDEDSIPPTDAATPMDDLSTPASSVPSSAPSDAGKPDSPTRE
TLDDVGDVSLAKHIVAQEMGVDIAEVTDDADLAEMGMDSLMSLTILGELREKTGIDLPSTFLTTNPTMKD
IDNALGMRPKPKAAPKPAAPKAAAPSSSKKTDMNEVSARLSALNNNTDISRYPNATSVLLQGNPKQATKK
IFFLPDGSGSATSYVSIPNLGPDVCAYGLNCPFMKNPEQWQCGIEISALVYLAEIKRRQPQGPYIIGGWS
AGGVIAYSVAQALLAANEGVEKLLLLDSPCPVNLAPLPARLHNFFNQIGLLGTGDPAKTPKWLLPHFSAA
IRSLSDYDPKPSLRPIPTYAIWCREGVAGNPGDPRPPPAEEEDPAPMTWLLEHRTNFKDNGWAQLCGDSM
KFGVMGGHHFSMMKPPHADDLGNLIREGLDWQP

>PKSI_1_10974
MSNVLLFGDQTAEQYPLLNKIVLRKENALVITFVERCAKALREETNALPRSQRNAVPDFLTVIDLKEAYH
QKGVKVPMVESALVTIAQIGHYIGYFSEHSAEQPSATNTRALGLCTGLLAAAAVVASKTVEELVLVGVEF
VRLSFRSGAAVDAARTALCQTGDDNAPWSTIVTGTTEASAKEALAKFHEEKGIPQTSHAYISAVSVMAIT
VSGPPTTVKRFFEESPALSKNHRVPIPVYGPYHAEHLFGETEINKIASASILEGLKQHQPVSLVHSAATG
KALVAENAAELAKLVLAEMLQLPVRWDHLLEEAVSQITSKKAPAKVWAMGVSNVANSLVSALKAGGQTDV
STVDQGTWTENEPDTHGRTQNDKVAIVGMAGRFPNSADHEALWELLMKGLDVHRRIPKDRFDADTHVDPS
GKGKNKSHTPFGCFIDEPGFFDPRFFNMSPREAAQTDPMGRLALVTAYEALEMSGYVPNRTPSTKLHRIG
TFYGQTSDDWREINAAENVDTYFITGGVRAFAPGRINYYFKFSGPSYSVDTACSSSLAAIQLACTSLWAG
DCDTACAGGLNVLTNPDIFSGLSKGQFLSKTGSCKTYDNNADGYCRGDAVGTVILKRYEDAIADKDNILG
CILGAATNHSAEAVSITHPHAGAQEFLYKRVLANAGVDAHEISYVEMHGTGTQAGDGIEMTSVTNVFAPR
HRQRRDDQPVYLGAIKANVGHAEAASGINSLAKVLLMMKHNKIPANVGIKGEMNKTFPADLKDRKVNISQKAVDWPRNGKEKRKVFLNNFSAAGGNTALLLEDGPAYEAPTATDPRGTVPVT
VTARSISALKRNIANLQKYVSENPSTTLTSMSYTLTARRIQHNYRVAFPLDQINKFSDALQAQVKESYSP
VPNAPTRVAFCFTGQGSQYTGLGQKLYNDLKSFRDDIDQLDHLARVQGLPSFLEIVQGADVQTLSPVKVQ
LGMACIQVALARMWAAWGITPAAVIGHSL
DMVLLVGRRAELLVRDCTPHTHGMLAVKGGAEAIRNTLGNKMTEIACINGPEETVLCGSGDVVGAANETL
AAKGFKATKLNVPFAFHSAQVDPILEQFKKIAASVTYNKPAVPVLSPLEGDIIREAGKINPEYLARHARE
TVKFWTALTAGQKEKVFDEKTAWLEVGAHPVCSGMVKASIGATTTAPSLRRGEDAWKTISNSMCTLFTAG
VNFNFDEFHKEFNDAQEMYTLPTYSFDNKKYWLDYHNDWTLRKGEPAQTKEVIVEKPVASASAPAVEMPA
KRLSTSCQRVIAENFSGNNGSVTVQSSLADPKLYPVVCGHMVNNAALCPSSLYADMALTISDYIWKQMRP
GTETPGYNVCNMEVPKPLIAQIPQPAEGQHIQLEANADLDSGIVKLNFRSVMPDGQKLQDHAHCIVRLED
KAAWEDEWSRYNYMVQAQMELLQHKTLNGGAHKVQRGMAYKLFKALVNYDEKYRAMAEVVLASGQTEASA
MLDFPTKPEDGDFYCPPYHIDGSCHISGFIVNASDLLDSEQNVYVSHGWGAMKFSRPLTAGMKLRNYVRMQPQPNNVSKGDVYIMEGDQIVAVCEGIKFQQIPRRVLNIFLPPNKGSGPASAAKPAAAPVA
AARPAPAAAPIKSAPAPAKAAPAPAPAAPKAAPKPKKAAAPKKPAGGLTAKVMKILAKETEVDEGELVDE
AQFENLGVDSLLSLTISAVFREELDMDISSTLFTDYPTVGDMKKYFAQFDNGSSTSSSAEEEDSDEDSIP
PTDAATPMDDLSTPASSVPSSAPSDAGKPDSPTRETLEDVGDVSLAKHIVAQEMGVDIAEVTDDADLAEMGMDSLMSLTIL
GELREKTGIDLPSTFLTTNPTMKDIDNALGMRPKPKAAPKPAAPKAAAPSSSKKTDMNEVSARLSALNNN
TDISRYPNATSVLLQGNPKQATKKIFFLPDGSGSATSYVSIPNLGPDVCAYGLNCPFMKNPEQWQCGIEI
SALVYLAEIKRRQPQGPYIIGGWSAGGVIAYSVAQALLAANEGVEKLLLLDSPCPVN
LAPLPARLHNFFNQIGLLGTGDPAKTPKWLLPHFSAAIRSLSDYDPKPSLRPIPTYAIWCREGVAGNPGD
PRPPPAEEEDPAPMTWLLEHRTNFKDNGWAQLCGDSMKFGVMGGHHFSMMKPPHADDLGNLIREGLDWQP

>PKSI_1_161
MSNVLLFGDQTAEQYPLLNKIVLRKENALVITFVERCAKALREETNALPRSQRNAVPDFLTVNDLKEAYH
QKGVKVPMVESALVTIAQIGHYIGYFSEHSAEQPSATNTRALGLCTGLLAAAAVVASKTVEELVLVGVDF
VRLSFRSGAAVDAARTALCQTGDDNAPWSTIVTGTTEASAKEALAKFHEEKGIPQTSHAYISAVSVMAIT
VSGPPTTVKRFFEESPALSKNHRVPIPVYGPYHAEHLFGETEINKIASDSILEGLKQHQPVSLVHSAATG
KALVAENAAELAKLVLAEMLQHSVRWDHLLEEAVSQITSKKAPAKIWAMGVSNVANSLVSALKAGGQSDV
STVDQSSWTENEPDTHGRTQNDKVAIVGMAGRFPNSADHEALWDLLMKGLDVHRRIPKDRFDADTHVDPS
GKGKNKSHTPFGCFIDEPGFFDPRFFNMSPREAAQTDPMGRLALVTAYEALEMSGYVPNRTPSTKLHRIG
TFYGQTSDDWREINAAENVDTYFITGGVRAFAPGRINYYFKFSGPSYSVDTACSSSLAAIQLACTSLWAG
DCDTACAGGLNVLTNPDIFSGLSKGQFLSKTGSCKTYDNNADGYCRGDAVGTVILKRYEDAIADKDNILG
CILGAATNHSAEAVSITHPHAGAQEFLYKRVLANAGVDAHEISYVEMHGTGTQAGDGIEMTSVTNVFAPR
HRQRRDDQPVYLGAIKANVGHAEAASGINSLAKVLMMMKHNKIPANVGIKGEMNKTFPADLKDRKVNISQ
KAVEWPRNGKEKRKVFLNNFSAAGGNTALLLEDGPAYEAPTATDPRGTVPVTVTARSISALKRNIANLQK
YVSENPSTTLTSMSYTLTARRIQHNYRVAFPLDQIDKFSDALQAQVKESYSPVPNVPTRVAFCFTGQGSQ
YTGLGQKLYNDLKSFRDDIDQLDHLARVQGLPSFLEIVQGADVQTLSPVKVQLGMACIQVALARMWAAWG
ITPAAVIGHSLGEYAALHVAGVISASDMVLLVGRRAELLVRDCTPHTHGMLAVKGGAEAIRNTLGNKMTE
IACINGPEETVLCGSGDVVGAANETLAAKGFKATKLNVPFAFHSAQVDPILEQFKKIAASVTYNKPAVPV
LSPLEGDIIREAGKINPEYLARHARETVNFWTALTAGQKEKVFDEKTAWLEVGAHPVCSGMVKASIGATT
TAPSLRRGEDAWKTISNSMCTLFTAGVNFNFDEFHKEFNDAQEMYTLPTYSFDNKKYWLDYHNDWTLRKG
EPAQTKEVIVEKPVASASAPAVEMPAKRLSTSCQRVIAENFSGNNGSVTVQSSLADPKLYPVVCGHMVNN
AALCPSSLYADMALTISDYIWKQMRPGTETPGYNVCNMEVPKPLIAQIPQPAEGQHIQLEANADLDSGIVKLNFRSVKPDG
QKLQDHAHCIVRLEDKATWEDEWSRYNYMVQAQMELLQHKTLNGGAHKVQRGMAYKLFKALVNYDEKYRA
MAEVVLASGQTEASAVLDFPTKPEDGDFYCPPYHIDGSCHISGFIVNASDLLDSEQNVYVSHGWGAMKFS
RPLTAGMKLRNYVRMQPQPNNVSKGDVYIMEGDQIVAVCEGIKFQQIPRRVLNTFLPPNKGSGPASAAKP
AAAPVAAARPAPAAAPIKTAPAPAKAAPAPAPAAPKAAPKPKKAAAPKKAAGGLTAKVMKILAKETEVDE
GELVDEAQFENLGVDSLLSLTISAVFREELDMDISSTLFTDYPTVGDMKKYFAQFDNGSSTSSSAEEEDS
DEDSIPPTDAATPMDDLSTPASSVPSSAPSDAGKPDSPTRETLEDVGDVSLAKHIVAQEMGVDIAEVTDD
ADLAEMGMDSLMSLTILGELREKTGIDLPSTFLTTNPTMKDIDNALGMRPKPKAAPKPAAPKAAAPSSSK
KTDMNEVSARLSALNNNTDISRYPNATSVLLQGNPKQATKKIFFLPDGSGSATSYVSI
PNLGPDVCAYGLNCPFMKNPEQWQCGIEISALVYLAEIKRRQPQGPYIIGGWSAGGVIAYSVAQALLAAN
EGVEKLLLLDSPCPVNLAPLPARLHNFFNEIGLLGTGDPAKTPKWLLPHFSAAIRSLSDYDPKPSLRPIP
TYAIWCREGVAGNPGDPRPPPAEEEDPAPMTWLLEHRTNFKDNGWAQLCGDSMKFGVMGGHHFSMMKPPH
ADDLGNLIREGLDWQP

>PKSI_1_6274
MSNVLLFGDQTAEQYQLLNKIVLRKENALVITFVERCAKALREETNALPRSQRNAVPDFLTVNDLKEAYH
QKGVKVPMVESALVTIAQIGHYIGYFSEHSAEQPSATNTRALGLCTGLLAAAAAVASKTVEELVLVGVEF
VRLSFRSGAAVDAARTALCQTGDDNAPWSTIVTGTTEASAKEALAKFHEEKGIPQTSHAYISAVSVMAIT
VSGPPTTVKRFFEESPALSKNHRVPIPVYGPYHAEHLFGETEINKIASASILEGLKQHQPVSLVHSAATG
KALVAENAAELAKLVLAEMLQLPVRWDHLLEEAVSQITSKKAPAKVWAMGVSNVANSLVSALKAGGQTDV
STVDQGTWTENEPDTHGRTQNDKVAIVGMAGRFPNSADHEALWELLMKGLDVHRRIPKDRFDADTHVDPS
GKGKNKSHTPFGCFIDEPGFFDPRFFNMSPREAAQTDPMGRLALVTAYEALEMSGYVPNRTPSTKLHRVG
TFYGQTSDDWREINAAENVDTYFITGGVRAFAPGRINYYFKFSGPSYSVDTACSSSLAAIQLACTSLWAG
DCDTACAGGLNVLTNPDIFSGLSKGQFLSKTGSCKTYDNNADGYCRGDAVGTVILKRYEDAIADKDNILG
CILGAATNHSAEAVSITHPHAGAQEFLYKRVLANAGVDAHEISYVEMHGTGTQAGDGIEMTSVTNVFAPR
HRQRRDDQPVYLGAIKANVGHAEAASGINSLAKVLLMMKHNKIPANVGIKGEMNKTFPADLKDRKVNISQ
KAVDWPRNGKEKRKVFLNNFSAAGGNTALLLEDGPAYEAPTATDPRGTVPVTVTARSISALKRNIANLQK
YVSENPSTTLTSMSYTLTARRIQHNYRVAFPLDQINKFSDALQAQVKESYSPVPNAPTRVAFCFTGQGSQ
YTGLGQKLYNDLKSFRDDIDQLDHLARV
NMSPREAAQTDPMGRLALVTAYEALEMSGYVPNRTPSTKLHRIGTFYGQTSDDWREINAAENVDTYFITG
GVRAFAPGRINYYFKFSGPSYSVDTACSSSLAAIQLACTSLWAGDCDTACAGGLNVLTNPDIFSGLSKGQ
FLSKTGSCKTYDNNADGYCRGDAVGTVILKRYEDAIADKDNILGCILGAATNHSAEAVSITHPHAGAQEF
LYKRVLANAGVDAHEISYVEMHGTGTQAGDGIEMTSVTNVFAPRHRQRRDDQPVYLGAIKANVGHAEAAS
GINSLAKVLLMMKHNKIPANVGIKGEMNKTFPADLKDRKVNISQKAVDWPRNGKEKRKVFLNNFSAAGGN
TALLLEDGPAYEAPTATDPRGTVPVTVTARSISALKRNIANLQKYVSENPSTTLTSMSYTLTARRIQHNY
RVAFPLDQINKFSDALQAQVKESYSPVPNAPTRVAFCFTGQGSQYTGLGQKLYNDLKSFRDDIDQLDHLA
RVQGLPSFLEIVQGADVQTLSPVKVQLGMACIQVALARMWAAWGITPAAVIGHSLGEYAALHVAGVISAS
DMVLLVGRRAELLVRDCTPHTHGMLAVKGGAEAIRNTLGNKMTEIACINGPEETVLCGSGDVVGAANETL
AAKGFKATKLNVPFAFHSAQVDPILEQFKKIAASVTYNKPAVPVLSPLEGDIIREAGKINPEYLARHARE
TVNFWTALTAGQKEKVFDEKTA

IREAGKINPEYLARHARETVNFWTALTAGQKEKVFDEKTAWLEVGAHPVCSGMVKASIGATTTAPSLRRG
EDAWKTISNSMCTLFTAGVNFNFDEFHKEFNDAQEMYTLPTYSFDNKKYWLDYHNDWTLRKGEPAQTKEV
IVEKPVASASAPAVEMPAKRLSTSCQRVIAENFSGNNGSVTVQSSLADPKLYPVVCGHMVNNAALCPSSL
YADMALTISDYIWKQMRPGTETPGYNVCNMEVPKPLIAQIPQPAEGQHIQLEANADLDSGIVKLNFRSVK
PDGQKLQDHAHCIVRLEDKAAWEDEWSRYNYMVQAQMELLQHKTLNGGAHKVQRGMAYKLFKA

MVQAQMELLQHKTLNGGAHKVQRGMAYKLFKALVNYDEKYRAMAEVVLASGQTEASAMLDFPTKPEDGDF
YCPPYHIDGSCHISGFIVNASDLLDSEQNVYVSHGWGAMKFSRPLTAGMKLRNYVRMQPQPNNVSKGDVY
IMEGDQIVAVCEGIKFQQIPRRVLNTFLPPNKGSGPASAAKPAAAPVAAARPAPAAAPIKTAPAPAKAAP
APAPAAPKAAPKPKKAAAPKKPAGGLTAKVMKILAKETEVDEGELVDEAQFENLGVDSLLSLTISAVFRE
ELDMDISSTLFTDYPTVGDMKKYFAQFDNGSSTSSSAEEEDSDEDSIPPTDAATPMDDLSTPASSVPSSA
PSDAGKPDSPTRETLEDVGDVSLAKHIVAQEMGVDIAEVTDDADLAEMGMDSLMSLTILGELREKTGIDL
PSTFLTTNPTMKDIDNALGMRPKPKAAPKSAAPKAAAPSSSKKTDMNEVSARLSALNNNTDISRYPNAT

ATKKIFFLPDGSGSATSYVSIPNLGPDVCAYGLNCPFMKNPEQWQCGIEISALVYLAEIKRRQPQGPYII
GGWSAGGVIAYSVAQALLAANEGVEKLLLLDSPCPVNLAPLPARLHNFFNQIGLLGTGDPAKTPKWLLPH
FSAAIRSLSDYDPKPSLRPIPTYAIWCREGVAGNPGDPRPPPAEEEDPAPMTWLLEHRTNFKDNGWAQLC
GDSMKFGVMGGHHFSMMKPPHADDLGNLIREGLDWQP
```

>PKSI_1_151

MSNVLLFGDQTAEQYQLLNKIVLRKENALVITFVERCAKALREETNALPRSQRNAVPDFLTVNDLKEAYHQKGVKVPMVESALVTIAQIGHYIGYFSEHSAEQPSATNTRALGLCTGLLAAAAVVASKTVEELVLVGVEFVRLSFRSGAAVDAARTALCQTGDDNAPWSTIVTGTTEASAKEALAKFHEEKGIPQTSHAYISAVSVMAITVSGPPTTVKRFFEESPALSKNHRVPIPVYGPYHAEHLFGETEINKIASASILEGLKQHQPVSLVHSAATGKALVAENAAELAKLVLAEMLQLPVRWDHLLEEAVSQITSKKAPAKIWAMGVSNVANSLVSALKAGGQTDVSTVDQSTWTENEPDTHGRTQNDKVAIVGMAGRFPNSADHEALWELLMKGLDVHRRIPKDRFDADTHVDPSGKGKNKSHTPFGCFIDEPGFFDPRFFNMSPREAAQTDPMGRLALVTAYEALEMSGYVPNRTPSTKLHRIGTFYGQTSDDWREINAAENVDTYFITGGVRAFAPGRINYYFKFSGPSYSVDTACSSSLAAIQLACTSLWAGDCDTACAGGLNVLTNPDIFSGLSKGQFLSKTGSCKTYDNNADGYCRGDAVGTVILKRYEDAIADKDNILGCILGAATNHSAEAVSITHPHAGAQEFLYKRVLANAGVDAHEISYVEMHGTGTQAGDGIEMTSVTNVFAPRHRQRRDDQPVYLGAIKANVGHAEAASGINSLAKVLLMMKHNKIPANVGIKGEMNKTFPADLKDRKVNISQKAVDWPRNGKEKRKVFLNNFSAAGGNTALLLEDGPAYEAPTATDPRGTVPVTVTARSISALKRNIANLQKYVSENPSTTLTSMSYTLTARRIQHNYRVAFPLDQINKFSDALQAQVKESYSPVPNAPTRVAFCFTGQGSQYTGLGQKLYNDLKSFRDDIDQLDHLARVQGLPSFLEIVQGADVQTLSPVKVQLGMACIQVALARMWAAWGITPAAVIGHSLGEYAALHVAGVISASDMVLLVGRRAELLVRDCTPHTHGMLAVKGGAEAIRNTLGNKMTEIACINGPEETVLCGSGDVVGAANETLAAKGFKATKLNVPFAFHSAQVDPILEQFKKIAASVTYNKPAVPVLSPLEGDIIREAGKINPEYLARHARETVNFWTALTAGQKEKVFDEKTAWLEVGAHPVCSGMVKASIGATTTAPSLRRGEDAWKTISNSMCTLFTAGVNFNFDEFHKEFNDAQEMYTLPTYSFDNKKYWLDYHNDWTLRKGEPAQTKEVIVEKPVASASAPAVEMPAKRLSTSCQRVIAENFSGNNGNVTVQSSLADPKLYPVVCGHMVNNAALCPSSLYADMALTISDYIWKQMRPGTETPGYNVCNMEVPKPLIAQIPQPAEGQHIQLEANADLDSGIVKLNFRSVKPDGQKLQDHAHCIVRLEDKAAWEDEWSRYNYMVQAQMELLQHKTLNGGAHKVQRGMAYKLFKALVNYDEKYRAMAEVVLASGQTEASAMLDFPTKPEDGDFYCPPYHIDGSCHISGFIVNASDLLDSEQNVYVSHGWGAMKFSRPLTAGMKLRNYVRMQPQPNNVSKGDVYIMEGDQIVAVCEGIKFQQIPRRVLNTFLPPNKGSGPASAAKPAAAPVAAARPAPAAAPIKTALAPAKAAPAPAPAAPKAAPKPKKAAAPKKPAGGLTAKVMKILAKETEVDEGELVDEAQFENLGVDSLLSLTISAVFREELDMDISSTLFTDYPTVGDMKKYFAQFDNGSSTSSSAEEEDSDEDSIPPTDAATPMDDLSTPASSVPSSAPSDAGKPDSPTRETLEDVGDVSLAKHIVAQEMGVDIAEVTDDADLAEMGMDSLMSLTILGELREKTGIDLPSTFLTTNPTMKDIDNALGMRPKPKAAPKPAAPKASAPSSSKKTDMNEVSARLSALNNNTDISRYPNATSVLLQGNPKQATKKIFFLPDGSGSATSYVSIPNLGPDVCAYGLNCPFMKNPEQWQCGIEISALVYLAEIKRRQPQGPYIIGGWSAGGVIAYSVAQALLAANEGVEKLLLLDSPCPVNLAPLPARLHNFFNQIGLLGTGDPAKTPKWLLPHFSAAIRSLSDYDPKPSLRPIPTYAIWCREGVAGNPGDPRPPPAEEEDPAPMTWLLEHRTNFKDNGWAQLCGDSMKFGVMGGHHFSMMKPPHADDLGNLIREGLDWQP

>PKSI_2_151

MAKVERKSWRPEDVAVIGLACRFSGSASNEAKLWELLEKCESAHSKVPGERYNVEAFHQQGSTNSNNLAADGGHYLEQDVKSFDAPFFNIITKEAKAMDPQARMLLESSYEALENAGLSLESVRGSDTGCYVGCFNRDYYELLMADAEDSPEYSVTGAGFSLLANRLSWYYDLRGPSKSEDTACSSSLVALDSAYKSLLRGESKMAMVCGANLMLSPNIGLWLSKLNMLSSEGLSRSFAEGVSGYGRGEGIATVILKPLADALRDGDTIRAVINATGVNQDGHTKGITVPNSQAQSTLIESTYRRAGLDFADTGYFEAHGTGTAVGDPLEIEALERVIKHAKRTSPLHVGSIKSSIGHLEGAAGLAGLIKCILMLEKGVILPNLHFERPNRKISFENIVVPTTAVPWPEGVKRRASVNLFGYGGTNAHVIVEAFVVPCPQSRDDDLRNAQDDGLERPSPQRLFVLTGREQSTVRKMRLRYASYIQTMKDTASPSVKFDDLSYTLGKRRSRLDWAEAHVASDFKELEEKLSAPEVAATRSSNKTRLGFVFTGQGAQWPRMGLELMRYTAFRDSVEAADEYLTQKLDCSWSVIEELEKHGDESRVTSSELGQPLCTIIQVAMVDLLGSWNVRPTAVVGHSSGEIAAAYCTGVMTKQAAWQIAFHRGRECAKLKGKAPELEGSMLAVGLDVESIRPYFNNLQSGRINVACVNSPNSITISGDASEIRKLQAMLVADSVFARELPVENAYHSHHMELVAESYLHSISDVDIQHSVALSDITMVSSVTGQSIEPSELIPEYWVRNLVSPVLFADAVTAMLRGSRRRFRRGAKAEPAVDFLLELGPHATLQTPFSDIVKAQAQEDVKYASMLLRGENAVDSAMTTAGKLYCHGCPVDVTAVNDIRHECRVLVDLPAYPWNRSTKYWGVSRLMQGYLHRTHGYHSLLGARLIGSDALNPAWRHFLNLDDSPWIEEHVVHGAVVYPGAGFLSMAIEAALQLAQPGREIANVRLQNVRVLKALVIKEGEDDPEVITRFRQADSVSDEASSMRWAFEISCAKGHDERDNRATGQDEFERHATGQITLDYQPEHPYLSPLSEQIHDVRRGEYARLADTCVDTMKQDGFYEASKDVGLAYGHDFQGIATMARGPNSCCWDLRVTERSTSLPGAYESKHLIHPTTLDAIVHSLFGAMNGGKKFQNAALPVAFDSIMISPATLTTSGTKLSGFTVIREAKEREIVADIHVSSEDWSQPLVQIAGLRCTQMPSPESELHDQDARPSPVGTITYRPDIALFDENGLMNYLNERQKPESCAKVHPDLYSERLRNAVAQVVELAMFKDPGLSVLQIGGYDRGVTDSLLFTLKAESAGQALSSKIVLLDPSQETLVEIRQQYDAESSIVDALHFAVDKPLPSEVSREYGFDVVLVAIDHGLDDMSKQNVLAEAQNVLKAGGIFVVFDTLRAITERTRDVARILQELLRSHDVEVTVCAWPPNAADVQNKSVISLLDLEQPFLCNLDASDFYTIRQIALKTTRLLWISPNDGPHTAVATGWLRVLQSENPNRQYQHLALGEATERSPFDLALAIAKVAVPLHLAAGQVKVELRFVDITHQDLVEPDMRALREASGVIKAVGSDVSLLRPGDNVCLSFVGHLSTSVNVDEALCQRIPPGVNMAEAACIPITLATALRALVGVAGVKPQHNVLVQAGGTKMGRAAILIASAANAVVYTTARDAEEVESLLALGISKQNIVPEGDPLLPTVTKILTGNRGWDVIVRTTKIVAETFILPECVADFGVVLDVFPSSGTGCTQETTISVMGIGSLLPEDPVLMQKTVSRIMDYLPQVSTLANSFDVFPSSAIPAALDRHGAQNQHRGVMLSFDQEDLVRVSPSATNTMKLYRDATYVMAGGLGGLGRSIARLLVDNGARNLVFLSRSGPNTTAATTMMSKMAGLGVTVKTLKCDVGDEKSVAAALDECSSMPPVRGVIQAAADIQDAIFDTYTFEQWQANLRPKVQGSWNLHCQLPEDMDFFVMLSSISGLIGHEGQAGYAAGNTFQDSLALFRHSRGLPAVTIDLGAMLDVGTIAEGSTTATFRSSDAVLMKAIDLHEIMTMCISNEINGYAIPAQVCTGLPSGGMLQVEQQEIPSYFHKPLFAALKCLGTSAVSAVNVAAPVEGVIDFAAQLTTVGSLDEADCVIANILRAHIAKAVQRAGKRSYNPCDDP

>PKSI_3_151

MPDNVSFMDESQDLRHIRETSRTSHTTSILNVEGDYRNNDMPREPGECNRSTNGTVDHERMTPSADGGIPIAICGIGLRLPGGIRNDRDLYDSLYNKKDARGVIPEDRFSIDSFHSAHGKTGTIITKHGYFLQDIDLTKFDVNMFNMTPAEVERLDPHQRILLETVRETLESAGEASFRGKKVGTYVGNFTDDWLDLQNVDTVDFATYQLHGKMDFSLANRISYEYDLRGPSMTIKTACSSSALAIHEAVYSIRNGECDAAIVSGSNLNLAPRLWVGMSSQGAISPDGSSKTFDESANGYARGDGIAALFIKRLDDAVRDGNPVRAVIRSTASNADGRTPGMTMPSTEAQEALIRRAYDAANLPLSETAMVECHGTGTAVGDPMEANAVARCFGDQGMLIGSVKPNLGHSEGASAITSVVKAVLSLENRTILPNIKFHRPNPAIPWSEAKLTVPVEPLAWPKDRQERISVNSFGIGGSNVHVVLDSAASMGFRPRSLAPSKDDRPGRLLLFSGGHRASVEQSSSQHQDYVTKYPNRLSDVAYTLAKRREHLHLRSFCVTRGAAPFHTTAPVKCPGLTRSTFVFTGQGAQWLHMGKELLHESPVFAKSIGRMDSVIHSLKHAPQWTLEGIINDPENPSALTNAEISQPLCTAVQIGLVDLLKSWAIYPHAVLGHSSGEIGAAYASGVVDRAEAILLAFYRGYVCRFAQKAGGMAAVGLEKSQVIKYLQPGVCVACENSGSSVTLSGDLETLEEVLQSIRAENLNAFARKLQVGIAYHSDHMKALGGLYHQYITEHLDPKDPQVPFFSSVSGRALHSKDDFGATYWQDNLENPVLFHTAVLKSLEHTGDKQVHLEVGPHGALNGPLRQIYAETGSKARYVALQKRGANCFDTFLEGIGQLYCNGVPLQYPESADDRTLIDLPPYPWHYDHSYWSETRVMKNWRFRRQLPHDLLGLRTLDCSDAEPMWRNILRITDLPWLRDHCVGKDVVFPASGYMCMAGEAVFQETGCRDYTLREVDISTAMVLSSDHSTELLTTMKKRRLNAFLDSRWYEFLIMSYDGASWTQHCSGLVTNGPSTSHPKALLQTYDRPVSTNRWYTAMSKIGLNYGPRFTGLQNITTHVQEKKASMTIMDKQEDYESPYALHPSTLDLILQSWTVASVRGEYRRFTQLFLPTFVDEFYIGNSASKLIHLNTTAIGPDGSARGEAIGKDNDGQISFNLKGFKGSKLDNVGVDQPQEMQTIMKQQWKRDFDFADTAQLMRPAFDSTSELSLLERMFVLAAIEVHVRTSGMEGKLPHHQRYKLWIDAQIRRFGEPGYPMVEDSMELLRLDSRERQRQLCRLLEHSRKTTAHPVAEAIWRALDRIEDVFDGRIEYLDLLFNDGLMPKFYDWSNSLSDVSRLFRLLSHKKPQLKILEVGAGTGGSTARLLQYLQSDFGERQYHSYTYTDVSSGFFVQAQERFKDYEGMKYRVLDISQDPFEQGFGADEFDLICASNVLHATPRLTETLRNCRKMLRPDGILFLQELCPRQQFMGFIMGLFEGWWLGAEDGRADTPLLLPPAWDRRLRDVGFEGVEAFSFDNNPPYFMAANMTARPTVTAKSKGSITLLTFNDFLDDVAGALKQAIRAAGFEIDHCVWGEHVPLDQSLISLVDLERDKPLLQDIGDDDLRIFLDLIDAVLQTTVIWLNKPAQVSSADPNAAQMLGLARTLRAELAMHFATVEMADPTLNGMSAVVHLMCMLQRGSALPENSLDQDMEYVWAHDAMHVSRFHFQPVDEALMEMSPKFDVKTLVPLQRGMLSSLKWIGTRMLPLAEAEVQIRMSAVGMNFHDMMIAMNMFDSPLTLGSGYNSIGMEGVGYVTRKSPEVDHVQVGDRVIVIGSNSSGFATDVHRPADYCIKCPSSLTDVEAAGMSFAYMTVLWSFLDKGGLRKGQSVLIHSAAGGVGIAALHVSRWLGLEAFVTVGNEEKVRFIMNNFGLPRNRIFNSHSATFLDDVMDATAGRGVDAALSAAAGELLHNTWSCIAPGGVMLEIGKRDLINRGRLSLAPFEENRSYVGIDFSRLTIVNKPAVVRLLRQTMRLVEQGHIHPIHPTTCFDAEHAEDAFRFMQTGQHIGRIVVKIPRDTSTIPLASRPPAPEFEGQKTYLLVGGMGGLGRSVASWMVSAGARNLIFMSRSAGKSEQDQNFARELELSGCKVHCCAIDITDTDAVREALERTQASVAGVLQMAMVLRDVGIMNMDKANWDAAVAPKVQGTWNLHHALPNVDFFVMFGSNSGTLGSYGQANYAAANAFLDSFVQYRQNLGQAASVIDIGAVGDVGYVAETQVAAENMESMAGRLISEQDFLNCLQLAIARSTPTERRNKGPSTEADGYIDLKQVILLNSSILSMADPSNQIFWRKDPRMGIYRNVQRTSAESPTTDSNSLRRLVATLKADSSIADQAELAQTLARELSKQVATVLMLGEDEIDVDRSLTDVGMDSLVAIEMRNWWKQNLGVDVSVLELKDGRSILRLGELAATRLKERLIDFALKEEAIIISPDYRLLPESNGSDILADVGSFWEWLQDSFVRLSADWHVQPDLDKILCAGQSSGGRLAVQSALLFHPISNIKALISISAPLHGDVPHFTIPYPKTIMGFQPLPSRDAEARVRSYVKAITPGTIVSSRDGADLWELLMCIIQQGWLPRLMGGRRDHRLDLMALLDEVQDMPAIWIIHGANDTVASSAFHSPINTHSFISGQAAWQNRLPSLRSRSAQSQAFFASFSPQSSATSSLPPGLKYNPLHLEPKSFSELSHRFLDYDQHIKVNRDFEEALRHVLSHFRPPICYAFAYGSGVFGQGKCDGGDELSPHPHPPRAVEEWQKNGAKIIDFIFGVSHTQHWHSVNLGDHPDHYSGLKHLPYSSGVISHIQDRFGAGVYFNPFITVNGIMIKYGVVNIDTLRSDLSAWDTLYLAGRLQKPVKILRDDPRICLANQINLKAALGTSLLMLPEIFSERQLYECIAGLSYVGDPRMNSYIASENPNKVSHIVGAQLPAFRQLYVPLIQDIPNVMFIHARKPDDTACGTGHAFRHTDVNPELNTFLSGGEGVSLSQDLDPVQRGEIVRHLPKAFKRKLYLAYLKKTRFPGSNFAGTPPEIEDEDPENDSLRSSQGGEIARRIAAHEDLPEMG

>PKSI_1_2682

MSNVLLFGDQTAEQYPLLNKIVLRKENALVITFVERCAKALREETNALPRSQRNAVPDFLTVNDLKEAYHQKGVKVPMVESALVTIAQIGHYIGYFSEHSAEQPSATNTRALGLCTGLLAAAAVVASKTVEELVLVGVEFVRLSFRSGAAVDAARTALCQTGDDNAPWSTIVTGTTEASAKEALAKFHEEKGIPQTSHAYISAVSVMAITVSGPPTTVKRFFEESPALSKNHRVPIPVYGPYHAEHLFGETEINKIASASILEGLKQHQPVSLVHSAATGKALVAENAAELAKLVLAEMLQLPVRWDHLLEEAVSQITSKKAPAKIWAMGVSNVANSLVSALKAGGQTDVSTVDQGTWTENEPDTHGRTQNDKVAIVGMAGRFPNSADHEALWELLMKGLDVHRRIPKDRFDADTHVDPSGKGKNKSHTPFGCFIDEPGFFDPRFFNMSPREAAQTDPMGRLALVTAYEALEMSGYVPNRTPSTKLHRIGTFYGQTSDDWREINAAENVDTYFITGGVRAFAPGRINYYFKFSGPSYSVDTACSSSLAAIQLACTSLWAGDCDTACAGGLNVLTNPDIFSGLSKGQFLSKTGSCKTYDNNADGYCRGDAVGTVILKRYEDAIADKDNILGCILGAATNHSAEAVSITHPHAGAQEFLYKRVLANAGVDAHEISYVEMHGTGTQAGDGIEMTSVTNVFAPRHRQRRDDQPVYLGAIKANVGHAEAASGINSLAKVLLMMKHNKIPANVGIKGEMNKTFPADLKDRKVNISQKAVDWPRNGKEKRKVFLNNFSAAGGNTALLLEDGPAYEAPTATDPRGTVPVTVTARSISALKRNIANLQKYVSENPSTTLTSMSYTLTARRIQHNYRVAFPLDQINKFSDALQAQVKESYSPVPNAPTRVAFCFTGQGSQYTGLGQKLYNDLKSFRDDIDQLDHLARVQGQPSFLEIVQGADVQTLSPVKVQLGMACIQVALARMWAAWGITPAAVIGHSLGEYAALHVAGVISASDMVLLVGRRAELLVRDCTPHTHGMLAVKGGAEAIRNTLGNKMTEIACINGPEETVLCGSGDVVGAANETLAAKGFKATKLNVPFAFHSAQVDPILEQFKKIAASVTYNKPAVPVLSPLEGDIIREAGKINPEYLARHARETVNFWTALTAGQKEKVFDEKTAWLEVGAHPVCSGMVKASIGATTTAPSLRRGEDAWKTISNSMCTLFTAGVNFNFDEFHKEFNDAQEMYTLPTYSFDNKKYWLDYHNDWTLRKGEPAQTKEVIVEKPVASASAPAVEMPAKRLSTSCQRVISENFSGNNGSVTVQSSLADPKLYPVVCGHMVNNAALCPSSLYADMALTISDYIWKQMRPGTETPGYNVCNMEVPKPLIAQIPQPAEGQHIQLEANADLDSGIVKLNFRSVKPDGQKLQDHAHCIVRLEDKAAWEDEWSRYNYMVQAQMELLQHKTLNGGAHKVQRGMAYKLFKALVNYDEKYRAMAEVVLASGQTEASAMLDFPTKPEDGDFYCPPYHIDGSCHISGFIVNASDLLDSEQNVYVSHGWGAMKFSRPLTAGMKLRNYVRMQPQPNNVSKGDVYIMEGDEIVAVCEGIKFQQIPRRVLNTFLPPNKGSGPASAAKPAAAPVAAARPAPAAAPIKTAPAPAKAAPAPAPAAPKAAPKPKKAAAPKKPAGGLTAKVMKILAKETEVDEGELVDEAQFENLGVDSLLSLTISAVFREELDMDISSTLFTDYPTVGDMKKYFAQFDNGSSTSSSAEEEDSDEDSIPPTDAATPMDDLSTPASSVPSSAPSDAGKPDSPTRETLEDVGDVSLAKHIVAQEMGVDIAEVTDDADLAEMGMDSLMSLTILGELREKTGIDLPSTFLTTNPTMKDIDNALGMRPKPKAAPKPAAPKAAAPSSSKKTDMNEVSARLSALNNNTDISRYPNATSVLLQGNPKQATKKIFFLPDGSGSATSYVSIPNLGPDVCAYGLNCPFMKNP

>PKSI_2_2682

MSNVLLFGDQTAEQYPLLNKIVLRKENALVITFVERCAKALREETNALPRSQRNAVPDFLTVNDLKEAYHQKGVKVPMVESALVTIAQIGHYIGYFSEHSAEQPSATNTRALGLCTGLLAAAAVVASKTVEELVLVGVEFVRLSFRSGAAVDAARTALCQTGDDNAPWSTIVTGTTEAAAKEALAKFHEEKGIPQTSHAYISAVSVMAITVSGPPTTVKRFFEESPALSKNHRVPIPVYGPYHAEHLFGETEINKIASASILEGLKQHQPVSLVHSAATGKALVAENAAELAKLVLAEMLQHPVRWDHLLEEAVSQITSKKAPAKIWAMGVSNVANSLVSALKAGGQTDVSTVDQSTWTENEPDTHGRTQNDKVAIVGMAGRFPNSADHEALWELLMKGLDVHRRIPKDRFDADTHVDPSGKGKNKSHTPFGCFIDEPGFFDPRFFNMSPREAAQTDPMGRLALVTAYEALEMSGYVPNRTPSTKLHRIGTFYGQTSDDWREINAAENVDTYFITGGVRAFAPGRINYYFKFSGPSYSVDTACSSSLAAIQLACTSLWAGDCDTACAGGLNVLTNPDIFSGLSKGQFLSKTGSCKTYDNNADGYCRGDAVGTVILKRYEDAIADKDNILGCILGAATNHSAEAVSITHPHAGAQEFLYKRVLANAGVDAHEISYVEMHGTGTQAGDGIEMTSVTNVFAPRHRQRRDDQPVYLGAIKANVGHAEAASGINSLAKVLLMMKHNKIPANVGIKGEMNKTFPADLKDRKVNISQKAVDWPRNGKEKRKVFLNNFSAAGGNTALLLEDGPAYEAPTATDPRGTVPVTVTARSISALKRNIANLQKYVSENPSTTLTSMSYTLTARRIQHNYRVAFPLDQINKFSDALQAQVKESYSPVPNALTRVAFCFTGQGSQYTGLGQKLYNDLKSFRDDIDQLDHLARVQGLPSFLEIVQGADVQTLSPVKVQLGMACIQVALARMWAAWGITPAAVIGHSLGEYAALHVAGVISASDMVLLVGRRAELLVRDCTPHTHGMLAVKGGAEAIRNTLGSKMTEIACINGPEETVLCGSGDVVGAANETLAAKGFKATKLNVPFAFHSAQVDPILEQFKKIAASVTYNKPAVPVLSPLEGDIIREAGKINPEYLARHARETVNFWTALTAGQKEKVFDEKTAWLEVGAHPVCSGMVKASIGATTTAPSLRRGEDAWKTISNSMCTLFTAGVNFNFDEFHKEFNDAQEMYTLPTYSFDNKKYWLDYHNDWTLRKGEPAQTKEVIVEKPVASASAPAVEMPAKRLSTSCQRVIAENFSGNNGSVTVQSSLADPKLYPVVCGHMVNNAALCPSSLYADMALTISDYIWKQMRPGTETPGYNVCNMEVPKPLIAQIPQPAEGQHIQLEANADLDSGIVKLNFRSVKPDGQKLQDHAHCIVRLEDRAAWEDEWSRYNYMVQAQMELLQHKTLNGGAHKVQRGMAYKLFKALVNYDEKYRAMAEVVLASGQTEASAVLDFPTKPEDGDFYCPPYHIDGSCHISGFIVNASDLLDSEQNVYVSHGWGAMKFSRPLTAGMKLRNYVRMQPQPNNVSKGAVYIMEGDQIVAVCEGIKFQQIPRRVLNTFLPPNKGSGPASAAKPAAAPVAAARPAPAAAPIKTAPAPAKAAPAPAPAAPKAAPKPKKAAAPKKAAGGLTAKVMKILAKETEVDEGELVDEAQFENLGVDSLLSLTISAVFREELDMDISSTLFTDYPTVGDMKKYFAQFDNGSSTSSSTEEEDSDEDSIPPTDAATPMDDLSTPASSVGSSAPSDAGKPDSPTRETLEDVGDVSLAKHIVAQEMGVDIAEVTDDADLAEMGMDSLMSLTILGELREKTGIDLPSTFLTTNPTMKDIDNALGMRPKPKAAPKPAAPKAAAPSSSKKTDMNEVSARLSALNNNTDISRYPNATSVLLQGNPKQATKKIFFLPDGSGSATSYVSIPNLGPDVCAYGLNCPFMKNPEQWQCGIEISAL

>PKSI_1_10513

MPDNVSFMDESQDLRHIRETSRTSHTTSILNVEGDYRNNDMPREPGECNRSTNGTVDHERMTPSADGGIPIAICGIGLRLPGGIRNDRDLYDSLYNKKDARGVIPEDRFSIDSFHSAHGKTGTIITKHGYFLQDIDLTKFDVNMFNMTPAEVERLDPHQRILLETVRETLESAGEASFRGKKVGTYVGNFTDDWLDLQNVDTVDFATYQLHGKMDFSLANRISYEYDLRGPSMTIKTACSSSALAIHEAVYSIRNGECDAAIVSGSNLNLAPRLWVGMSSQGAISPDGSSKTFDESANGYARGDGIAALFIKRLDDAVRDGNPVRAVIRSTASNADGRTPGMTMPSTEAQEALIRRAYDAANLPLSETAMVECHGTGTAVGDPMEANAVARCFGDQGMLIGSVKPNLGHSEGASAITSVVKAVLSLENRTILPNIKFHRPNPAIPWSEAKLTVPVEPLAWPKDRQERISVNSFGIGGSNVHVVLDSAASMGFRPRSLAPSKDDRPGRLLLFSGGHRASVEQSSSQHQDYVTKYPNRLSDVAYTLAKRREHLHLRSFYVTRGAAPFHTTAPVKCPGLTRSTFVFTGQGAQWLHMGKELLHESPVFAKSIGRMDSVIHSLKHAPQWTLEGIINDPENPSALTNAEISQPLCTAVQIGLVDLLKSWAIYPHAVLGHSSGEIGAAYASGVVDRAEAILLAFYRGYVCRFAQKAGGMAAVGLEKSQVIKYLQPGVCVACENSGSSVTLSGDLETLEEVLQSIRAENLNAFARKLQVGIAYHSDHMKALGGLYHQYITEHLDPKDPQVPFFSSVSGRALHSKDDFGATYWQDNLENPVLFHTAVLKSLEHTGDKQVHLEVGPHGALNGPLRQIYAETGSKARYVALQKRGANCFDTFLEGIGQLYCNGVPLQYPESADDRTLIDLPPYPWHYDHSYWSETRVMKNWRFRRQLPHDLLGLRTLDCSDAEPMWRNILRITDLPWLRDHCVGKDVVFPASGYMCMAGEAVFQETGCRDYTLREVDISTAMVLSSDHSTELLTTMKKRRLNAFLDSRWYEFLIMSYDGASWTQHCSGLVTNGPSTSHPKALLQTYDRPVSTNRWYTAMSKIGLNYGPRFTGLQNITTHVQEKKASMTIMDKQEDYESPYALHPSTLDLILQSWTVASVRGEYRRFTQLFLPTFVDEFYIGNSASKLIHLNTTAIGPDGSARGEAIGKDNDGQISFNLKGFKGSKLDNVGVDQPQEMQTIMKQQWKRDFDFADTAQLMRPAFDSTSELSLLERMFVLAAIEVHVRTSGMEGKLPHHQRYKLWIDAQIRRFGEPGYPMVEDSMELLRLDSRERQRQLCRLLEHSRKTTAHPVAEAIWRALDRIEDVFDGRIEYLDLLFNDGLMPKFYDWSNSLSDVSRLFRLLSHKKPQLKILEVGAGTGGSTARLLQYLQSDFGERQYHSYTYTDVSSGFFVQAQERFKDYEGMKYRVLDISQDPFEQGFGADEFDLICASNVLHATPRLTETLRNCRKMLRPDGILFLQELCPRQQFMGFIMGLFEGWWLGAEDGRADTPLLLPPAWDRRLRDVGFEGVEAFSFDNNPPYFMAANMTARPTVTAKSKGSITLLTFNDFLDDVAGALKQAIRAAGFEIDHCVWGEHVPLDQSLISLVDLERDKPLLQDIGDDDLRIFLDLIDAVLQTTVIWLNKPAQVSSADPNAAQMLGLARTLRAELAMHFATVEMADPTLNGMSAVVHLMCMLQRGSALPENSLDQDMEYVWAHDAMHVSRFHFQPVDEALMEMSPKFDVKTLVPLQRGMLSSLKWIGTRMLPLAEAEVQIRMSAVGMNFHDMMIAMNMFDSPLTLGSGYNSIGMEGVGYVTRKSPEVDHVQVGDRVIVIGSNSSGFATDVHRPADYCIKCPSSLTDVEAAGMSFAYMTVLWSFLDKGGLRKGQSVLIHSAAGGVGIAALHVSRWLGLEAFVTVGNEEKVRFIMNNFGLPRNRIFNSHSATFLDDVMDATAGRGVDAALSAAAGELLHNTWSCIAPGGVMLEIGKRDLINRGRLSLAPFEENRSYVGIDFSRLTIVNKPAVVRLLRQTMRLVEQGHIHPIHPTTCFDAEHAEDAFRFMQTGQHIGRIVVKIPRDTSTIPLASRPPAPEFEGQKTYLLVGGMGGLGRSVASWMVSAGARNLIFMSRSAGKSEQDQNFARELELSGCKVHCCAIDITDTDAVREALERTQASVAGVLQMAMVLRDVGIMNMDKANWDAAVAPKVQGTWNLHHALPNVDFFVMFGSNSGTLGSYGQANYAAANAFLDSFVQYRQNLGQAASVIDIGAVGDVGYVAETQVAAENMESMAGRLISEQDFLNCLQLAIARSTPTERRNKGPSTEADGYIDLKQVILLNSSILSMADPSNQIFWRKDPRMGIYRNVQRTSAESPTTDSNSLRRLVATLKADSSIADQAELAQTLARELSKQVATVLMLGEDEIDVDRSLTDVGMDSLVAIEMRNWWKQNLGVDVSVLELKDGRSILRLGELAATRLKERLIDFALKEEAIIISPDYRLLPESNGSDILADVGSFWEWLQDSFVRLSADWHVQPDLDKILCAGQSSGGRLAVQSALLFHPISNIKALISISAPLHGDVPHFTIPYPKTIMGFQPLPSRDAEARVRSYVKAITPGTIVSSRDGADLWELLMCIIQQGWLPRLMGGRRDHRLDLMALLDEVQDMPAIWIIHGANDTVASSAFHSPINTHSFISGQAAWQNRLPSLRSRSAQSQAFFASFSPQSSATSSLPPGLKYNPLHLEPKSFSELSHRFLDYDQHIKVNRDFEEALRHVLSHFRPPICYAFAYGSGVFGQGKCDGGDELSPHPHPPRAVEEWQKNGAKIIDFIFGVSHTQHWHSVNLGDHPDHYSGLKHLPYSSGVISHIQDRFGAGVYFNPFITVNGIMIKYGVVNIDTLRSDLSAWDTLYLAGRLQKPVKILRDDPRICLANQINLKAALGTSLLMLPEIFSERQLYECIAGLSYVGDPRMNSYIASENPNKVSHIVGAQLPAFRQLYVPLIQDIPNVMFIHARKPDDTACGTGHAFRHTDVNPELNTFLSGGEGVSLSQDLDPVQRGEIVRHLPKAFKRKLYLAYLKKTRFPGSNFAGTPPEIEDEDPENDSLRSSQGGEIARRIAAHEDLPEMG

>PKSI_1_171

MSNVLLFGDQTAEQYPLLNKIVLRKENALVITFVERCAKALREETNALPRSQRNAVPDFLTVNDLKEAYHQKGVKVPMVESALVTIAQIGHYIGYFSEHSAEQPSATNTRALGLCTGLLAAAAVVASKTVEELVLVGVEFVRLSFRSGAAVDAARTALCQTGDDNAPWSTIVTGTTEASAKEALAKFHEEKGIPQTSHAYISAVSVMAITVSGPPTTVKRFFEESPALSKNHRVPIPVYGPYHAEHLFGETEINKIASASILEGLKQHQPVSLVHSAATGKALVAENAAELAKLVLAEMLQHPVRWDHLLEEAVSQITSKKAPAKIWAMGVSNVANSLVSALKAGGQTDVSTVDQSSWTENEPDTHGRTQNDKVAIVGMAGRFPNSADHEALWELLMKGLDVHRRIPKDRFDADTHVDPSGKGKNKSHTPFGCFIDEPGFFDPRFFNMSPREAAQTDPMGRLALVTAYEALEMSGYVPNRTPSTKLHRIGTFYGQTSDDWREINAAENVDTYFITGGVRAFAPGRINYYFKFSGPSYSVDTACSSSLAAIQLACTSLWAGDCDTACAGGLNVLTNPDIFSGLSKGQFLSKTGSCKTYDNNADGYCRGDAVGTVILKRYEDAIADKDNILGCILGAATNHSAEAVSITHPHAGAQEFLYKRVLANAGVDAHEISYVEMHGTGTQAGDGIEMTSVTNVFAPRHRQRRDDQPVYLGAIKANVGHAEAASGINSLAKVLLMMKHNKIPANVGIKGEMNKTFPADLKDRKVNISQKAVDWPRNGKEKRKVFLNNFSAAGGNTALLLEDGPAYEAPTATDPRGTVPVTVTARSISALKRNIANLQKYVSENPSTTLTSMSYTLTARRIQHNYRVAFPLDQINKFSDALQAQVKESYSPVPNAPTRVAFCFTGQGSQYTGLGQKLYNDLKSFRDDIDQLDHLARVQGLPSFLEIVQGADVQTLSPVKVQLGMACIQVALARMWAAWGITPAAVIGHSLGEYAALHVAGVISASDMVLLVGRRAELLVRDCTPHTHGMLAVKGGAEAIRNTLGNKMTEIACINGPEETVLCGSGDVVGAANETLAAKGFKATKLNVPFAFHSAQVDPILEQFKKIAASVTYNKPAVPVLSPLEGDIIREAGKINPEYLARHARETVNFWTALTAGQKEKVFDEKTAWLEVGAHPVCSGMVKASIGATTTAPSLRRGEDAWKTISNSMCTLFTAGVNFNFDEFHKEFNDAQEMYTLPTYSFDNKKYWLDYHNDWTLRKGEPAQTKEVIVEKPVASASAPAVEMPAKRLSTSCQRVIAENFSGSNGSVTVQSSLADPKLYPVVCGHMVNNAALCPSSLYADMALTISDYIWKQMRPGTETPGYNVCNMEVPKPLIAQIPQPADGQHIQLEANADLDSGIVKLNFRSVKPDGQKLQDHAHCIVRLEDKAAWEDEWSRYNYMVQAQMELLQHKTLNGGAHKVQRGMAYKLFKALVNYDEKYRAMAEVVLASGQTEASAVLDFPTKPEDGDFYCPPYHIDGSCHISGFIVNASDLLDSEQNVYVSHGWGAMKFSRPLTAGMKLRNYVRMQPQPNNVSKGDVYIMEGDQIVAVCEGIKFQQIPRRVLNTFLPPNKGSGPASAAKPAAAPVAAARPAPAAAPIKTAPAPAKAAPAPAPAAPKAAPKPKKAAAPKKPAGGLTAKVMKILAKETEVDEGELVDEAQFENLGVDSLLSLTISAVFREELDMDISSTLFTDYPTVGDMKKYFARFDNGSSTSSSAEEEDSDEDSIPPTDAATPMDDLSTPASSVPSSAPSDAGKPDSPTRETLEDVGDVSLAKHIVAQEMGVDIAEVTDDADLAEMGMDSLMSLTILGELREKTGIDLPSTFLTTNPTMKDIDNALGMRPKPKAAPKPAAPKAAAPSSSKKTDMNEVSARLSALNNNTDISRYPNATSVLLQGNPKQATKKIFFLPDGSGSATSYVSIPNLGPDVCAYGLNCPFMKNPEQWQCGIEISALVYLAEIKRRQPQGPYIIGGWSAGGVIAYSVAQALLAANEGVEKLLLLDSPCPVNLAPLPARLHNFFNEIGLLGTGDPAKTPKWLLPHFSAAIRSLSDYDPKPSLRPIPTYAIWCREGVAGNPGDPRPPPAEEEDPAPMTWLLEHRTNFKDNGWAQLCGDSMKFGVMGGHHFSMMKPPHADDLGNLIREGLDWQP

>PKSI_2_171

MSNVLLFGDQTAEQYPLLNKIVLRKENALVITFVERCAKALREETNALPRSQRNAVPDFLTVNDLKEAYHQKGVKVPMVESALVTIAQIGHYIGYFSEHSAEQPSATNTRALGLCTGLLAAAAVVASKTVEELVLVGVEFVRLSFRSGAAVDAARTALCQTGDDNAPWSTIVTGTTEAAAKEALAKFHEEKGIPQTSHAYISAVSVMAITVSGPPTTVKRFFEESPALSKNHRVPIPVYGPYHAEHLFGATEINKIASASILEGLKQHQPVSLVHSAATGKALVAENAAELAKLVLAEMLQHPVRWDHLLEEAVSQITSKKAPAKIWAMGVSNVANSLVSALKAGGQTDVSTVDQSTWTENEPDTHGRTQNDKVAIVGMAGRFPNSADHEALWELLMKGLDVHRRIPKDRFDADTHVDPSGKGKNKSHTPFGCFIDEPGFFDPRFFNMSPREAAQTDPMGRLALVTAYEALEMSGYVPNRTPSTKLHRIGTFYGQTSDDWREINAAENVDTYFITGGVRAFAPGRINYYFKFSGPSYSVDTACSSSLAAIQLACTSLWAGDCDTACAGGLNVLTNPDIFSGLSKGQFLSKTGSCKTYDNNADGYCRGDAVGTVILKRYEDAIADKDNILGCILGAATNHSAEAVSITHPHAGAQEFLYKRVLANAGVDAHEISYVEMHGTGTQAGDGIEMTSVTNVFAPRHRQRRDDQPVYLGAIKANVGHAEAASGINSLAKVLLMMKHNKIPANVGIKGEMNKTFPADLKDRKVNISQKAVDWPRNGKEKRKVFLNNFSAAGGNTALLLEDGPAYEAPTATDPRGTVPVTVTARSISALKRNIANLQKYVSENPSTTLTSMSYTLTARRIQHNYRVAFPLDQINKFSDALQAQVKESYSPVPNAPTRVAFCFTGQGSQYTGLGQKLYNDLKSFRDDIDQLDHLARVQGLPSFLEIVQGADVQTLSPVKVQLGMACIQVALARMWAAWGITPAAVIGHSLGEYAALHVAGVISASDMVLLVGRRAELLVRDCTPHTHGMLAVKGGAEAIRNTLGSKMTEIACINGPEETVLCGSGDVVGAANETLAAKGFKATKLNVPFAFHSAQVDPILEQFKKIAASVTYNKPAVPVLSPLEGDIIREAGKINPEYLARHARETVNFWTALTAGQKEKVFDEKTAWLEVGAHPVCSGMVKASIGATTTAPSLRRGEDAWKTISNSMCTLFTAGVNFNFDEFHKEFNDAQEMYTLPTYSFDNKKYWLDYHNDWTLRKGEPAQTKEVIVEKPVASASAPAVEMPAKRLSTSCQRVIAENFTGNNGSVTVQSSLADPKLYPVVCGHMVNNAALCPSSLYADMALTISDYIWKQMRPGTETPGYNVCNMEVPKPLIAQIPQPAEGQHIQLEANADLDSGIVKLNFRSVKPDGQKLQDHAHCIVRLEDRAAWEDEWSRYNYMVRAQMELLQHKTLNGGAHKVQRGMAYKLFKALVNYDEKYRAMAEVVLASGQTEASAVLDFPTKPEDGDFYCPPYHIDGSCHISGFIVNASDLLDSEQNVYVSHGWGAMKFSRPLTAGMKLRNYVRMQPQPNNVSKGDVYIMEGDQIVAVCEGIKFQQIPRRVLNTFLPPNKGSGPASAAKPAAAPVAAARPAPAAAPIKTAPAPAKAAPAPAPAAPKAAPKPKKAAAPKKAAGGLTAKVMKILAKETEVDEGELVDEAQFENLGVDSLLSLTISAVFREELDMDISSTLFTDYPTVGDMKKYFAQFDNGSSTSSSTEEEDSDEDSIPPTDAATPMDDLSTPASSVGSSAPSDAGKPDSPTRETLEDVGDVSLAKHIVAQEMGVDIAEVTDDADLAEMGMDSLMSLTILGELREKTGIDLPSTFLTTNPTMKDIDNALGMRPKPKAAPKPAAPKAAAPSSSKKTDMNEVSARLSALNNNTDISRYPNATSVLLQGNPKQATKKIFFLPDGSGSATSYVSIPNLGPDVCAYGLNCPFMKNPEQWQCGIEISAL

>PKSI_1_2788

MSNVLLFGDQTAEQYPLLNKIVLRKENALVITFVERCAKALREETNALPRSQRNAVPDFLTVNDLKEAYHQKGVKVPMVESALVTIAQIGHYIGYFSEHSAEQPSATNTRALGLCTGLLAAAAVVASKTVEELVLVGVEFVRLSFRSGAAVDAARTALCQTGDDNAPWSTIVTGTTEASAKEALAKFHEEKGIPQTSHAYISAVSVMAITVSGPPTTVKRFFEESPALSKNHRVPIPVYGPYHAEHLFGETEINKIASASILEGLKQHQPVSLVHSAATGKALVAENAAELAKLVLAEMLQHPVRWDHLLEEAVSQITSKKAPAKIWAMGVSNVANSLVSALKAGGQSDVSTVDQSSWTENEPDTHGRTQNDKVAIVGMAGRFPNSADHEALWDLLMKGLDVHRRIPKDRFDADTHVDPSGKGKNKSHTPFGCFIDEPGFFDPRFFNMSPREAAQTDPMGRLALVTAYEALEMSGYVPNRTPSTKLHRIGTFYGQTSDDWREINAAENVDTYFITGGVRAFAPGRINYYFKFSGPSYSVDTACSSSLAAIQLACTSLWAGDCDTACAGGLNVLTNPDIFSGLSKGQFLSKTGSCKTYDNNADGYCRGDAVGTVILKRYEDAIADKDNILGCILGAATNHSAEAVSITHPHAGAQEFLYKRVLANAGVDAHEISYVEMHGTGTQAGDGIEMTSVTNVFAPRHRQRRDDQPVYLGAIKANVGHAEAASGINSLAKVLMMMKHNKIPANVGIKGEMNKTFPADLKDRKVNISQKAVEWPRNGKEKRKVFLNNFSAAGGNTALLLEDGPAYEAPTASDPRGTVPVTVTARSISALKRNIANLQKYVSENPSTTLTSMSYTLTARRIQHNYRVAFPLDQINKFSDALQAQVKESYSPVPNAPTRVAFCFTGQGSQYTGLGQKLYNDLKSFRDDIDQLDHLARVQGLPSFLEIVQGADVQTLSPVKVQLGMACIQVALARMWAAWGITPAAVIGHSLGEYAALHVAGVISASDMVLLVGRRAELLVRDCTPHTHGMLAVKGGAEAIRNTLGNKMTEIACINGPEETVLCGSGDVVGAANETLAAKGFKATKLNVPFAFHSAQVDPILEQFKKIAASVTYNKPAVPVLSPLEGDIIREAGKINPEYLARHARETVNFWTALTAGQKEKVFDEKTAWLEVGAHPVCSGMVKASIGATTTAPSLRRGEDAWKTISNSMCTLFTAGVNFNFDEFHKEFNDAQEMYTLPTYSFDNKKYWLDYHNDWTLRKGEPAQTKEVIVEKPVASASAPAVEIPAKRLSTSCQRVISENFSGNNGSVTVQSSLADPKLYPVVCGHMVNNAALCPSSLYADMALTISDYIWKQMRPGTETPGYNVCNMEVPKPLIAQIPQPAEGQHIQLEANADLDSGIVKLNFRSVKPDGQKLQDHAHCIVRLEDRAAWEDEWSRYNYMVQAQMELLQHKTLNGGAHKVQRGMAYKLFKALVNYDEKYRAMAEVVLASGQTEASAVLDFPTKPEDGDFYCPPYHIDGSCHISGFIVNASDLLDSEQNVYVSHGWGAMKFSRPLTAGMKLRNYVRMQPQPNNVSKGDVYIMEGDQIVAVCEGIKFQQIPRRVLNTFLPPNKGSGPASAAKPAAAPVAAARPAPAAAPIKTAPAPAKAAPAPAPAAPKAAPKPKKAAAPKKAAGGLTAKVMKILAKETEVDEGELVDEAQFENLGVDSLLSLTISAVFREELDMDISSTLFTDYPTVGDMKKYFAQFDNGSSTSSSAEEEDSDEDSIPPTDAATPMDDLSTPASSVPSSAPSDAGKPDSPTRETLEDVGDVSLAKHIVAQEMGVDIAEVTDDADLAEMGMDSLMSLTILGELREKTGIDLPSTFLTTNPTMKDIDNALGMRPKPKAAPKPAAPKAAAPSSSKKTDMNEVSARLSALNNNTDISRYPNATSVLLQGNPKQATKKIFFLPDGSGSATSYVSIPNLGPDVCAYGLNCPFMKNPEQWQCGIEISALVYLAEIKRRQPQGPYIIGGWSAGGVIAYSVAQALLAANEGVEKLLLLDSPCPVNLAPLPARLHNFFNEIGLLGTGDPAKTPKWLLPHFSAAIRSLSDYDPKPSLRPIPTYAIWCREGVAGNPGDPRPPPAEEEDPAPMTWLLEHRTDFKDNGWAQLCGDSMKFGVMGGHHFSMMKPPHADDLGNLIREGLDWQP

>PKSI_1_562

MSNVLLFGDQTAEQYPLLNKIVLRKENALVITFIERCAKALREETNALPRSQRNAVPDFLTVNDLKEAYHQKGVKVPMVESALVTIAQIGHYIGYFSEHSAEQPSATNTRALGLCTGLLAAAAVVASKTVEELVLVGVEFVRLSFRSGAAVDAARTALCQTGDDNSPWSTIVTGTTEAAAKEALAKFHEEKSIPQTSHAYISAVSVMAITVSGPPTTVKRFFEETSALSKNHRVPIPVYGPYHAEHLFGETEINKIASASILEGLKQHQPVSLVHSAATGKALVAENAAELAKLVLAEMLQHPVRWDHLLEEAVSQITSKKAPAKIWAMGVSNVANSLVSALKAGGQTDVSTVDQSTWTENEPDTHGRTQNDKVAIVGMAGRFPNSADHEALWELLMKGLDVHRRIPKDRFDADTHVDPSGKGKNKSHTPFGCFIDEPGFFDPRFFNMSPREAAQTDPMGRLALVTAYEALEMSGYVPNRTPSTKLHRIGTFYGQTSDDWREINAAENVDTYFITGGVRAFAPGRINYYFKFSGPSYSVDTACSSSLAAIQLACTSLWAGDCDTACAGGLNVLTNPDIFSGLSKGQFLSKTGSCKTYDNNADGYCRGDAVGTVILKRYEDAIADKDNILGCILGAATNHSAEAVSITHPHAGAQEFLYKRVLANAGVDAHEISYVEMHGTGTQAGDGIEMTSVTNVFAPRHRQRRDDQPVYLGAIKANVGHAEAASGINSLAKVLLMMKHNKIPANVGIKGEMNKTFPADLKDRKVNISQKAVDWPRNGKEKRKVFLNNFSAAGGNTALLLEDGPAYEAPTATDPRGTVPVTVTARSISALKRNIANLQKYVSENPSTTLTSMSYTLTARRIQHNYRVAFPLDQINKFSDALQAQVKESYSPVPNAPTRVAFCFTGQGSQYTGLGQKLYNDLKSFRDDIDQLDHLARVQGLPSFLEIVQGADVQTLSPVKVQLGMACIQIALARMWAAWGITPAAVIGHSLGEYAALHVAGVISASDMVLLVGRRAELLVRDCTPHTHGMLAVKGGAEAIRNTLGSKMTEIACINGPEETVLCGSGDVVGAANETLAAKGFKATKLNVPFAFHSAQVDPILEQFKKIAASVTYNKPAVPVLSPLEGDIIREAGKINPEYLARHARETVNFWTALTAGQKEKVFDEKTAWLEVGAHPVCSGMVKASIGATTTAPSLRRGEDAWKTISNSMCTLFTAGVNFNFDEFHKEFNDAQEMYTLPTYSFDNKKYWLDYHNDWTLRKGEPAQTKEVIVEKPVASASAPAVEMPAKRLSTSCQRVIAENFSSNNGSVTVQSSLADPKLYPVVCGHMVNNAALCPSSLYADMALTISDYIWKQMRPGTETPGYNVCNMEVPKPLIAQIPQPAEGQHIQLEANADLDSGIVKLNFRSVKPDGQKLQDHAHCIVRLEDKATWEDEWSRYNYMVQAQMELLQHKTLNGGAHKVQRGMAYKLFKALVNYDEKYRAMAEVVLASGQTEASAVLDFPTKPEDGDFYCPPYHIDGSCHISGFIVNASDLLDSEQNVYVSHGWGAMKFSRPLTAGMKLRNYVRMQPQPNNVSKGDVYIMEGDQIVAVCEGIKFQQIPRRVLNTFLPPNKGSGPASAAKPAAAPVAAARPAPAAAPIKTAPAPAKAAPAPAPAAPKAAPKPKKAAAPKKAAGGLTAKVMKILAKETEVDEGELVDEAQFENLGVDSLLSLTISAVFREELDMDISSTLFTDYPTVGDMKKYFAQFDNGSSTSSSTEEEDSDEDSIPPTDAATPMDDLSTPASSVGSSAPSDAGKPDSPTRETLEDVGDVSLAKHIVAQEMGVDIAEVTDDADLAEMGMDSLMSLTILGELREKTGIDLPSTFLTTNPTMKDIDNALGMRPKPKAVPKPAAPKAAAPSSSKKTDMNEVSARLSALNNNTDISRYPNATSVLLQGNPKQATKKIFFLPDGSGSATSYVSIPNLGPDVCAYGLNCPFMKNPEQWQCGIEISALVYLAEIKRRQPQGPYIIGGWSAGGVIAYSVAQALLAANEGVEKLLLLDSPCPVNLAPLPARLHNFFNEIGLLGTGDPAKTPKWLLPHFSAAIRSLSDYDPKPSLRPIPTYAIWCREGVAGNPGDPRPPPAEEEDPAPMTWLLEHRTNFKDNGWAQLCGDSMKFGVMGGHHFSMVSHNLTLDSFSSNTMLTPVQMKPPHADDLGNLIREGLDWQP

>PKSI_1_120

MSNVLLFGDQTAEQYPLLNKIVLRKENALVITFIERCAKALREETNALPRSQRNAVPDFLTVNDLKEAYHQKGVKVPMVESALVTIAQIGHYIGYFSEHSAEQPSATNTRALGLCTGLLAAAAVVASKTVEELVLVGVEFVRLSFRSGAAVDAARTALCQTGDDNSPWSTIVTGTTEAAAKEALAKFHEEKSIPQTSHAYISAVSVMAITVSGPPTTVKRFFEESSALSKNHRVPIPVYGPYHAEHLFGETEINKIASASILEGLKQHQPVSLVHSAATGKALVAENAAELAKLVLAEMLQHPVRWDHLLEEAVSQITSKKAPAKIWAMGVSNVANSLVSALKAGGQTDVSTIDQSTWTENEPDTHGRTQNDKVAIVGMAGRFPNSADHEALWELLMKGLDVHRRIPKDRFDADTHVDPSGKGKNKSHTPFGCFIDEPGFFDPRFFNMSPREAAQTDPMGRLALVTAYEALEMSGYVPNRTPSTKLHRIGTFYGQTSDDWREINAAENVDTYFITGGVRAFAPGRINYYFKFSGPSYSVDTACSSSLAAIQLACTSLWAGDCDTACAGGLNVLTNPDIFSGLSKGQFLSKTGSCKTYDNNADGYCRGDAVGTVILKRYEDAIADKDNILGCILGAATNHSAEAVSITHPHAGAQEFLYKRVLANAGVDAHEISYVEMHGTGTQAGDGIEMTSVTNVFAPRHRQRRDDQPVYLGAIKANVGHAEAASGINSLAKVLLMMKHNKIPANVGIKGEMNKTFPADLKDRKVNISQKAVDWPRNGKEKRKVFLNNFSAAGGNTALLLEDGPAYEAPTATDPRGTVPVTVTARSISALKRNIANLQKYVSENPSTTLTSMSYTLTARRIQHNYRVAFPLDQINKFSDALQAQVKESYSPVPNAPTRVAFCFTGQGSQYTGLGQKLYNDLKSFRDDIDQLDHLARVQGLPSFLEIVQGADVQTLSPVKVQLGMACIQVALARMWAAWGITPAAVIGHSLGEYAALHVAGVISASDMVLLVGRRAELLVRDCTPHTHGMLAVKGGAEAIRNTLGSKMTEIACINGPEETVLCGSGDVVGAANETLAAKGFKATKLNVPFAFHSAQVDPILEQFKKIAASVTYNKPAVPVLSPLEGDIIREAGKINPEYLARHARETVNFWTALTAGQKEKVFDEKTAWLEVGAHPVCSGMVKASIGATTTAPSLRRGEDAWKTISNSMCTLFTAGVNFNFDEFHKEFNDAQEMYTLPTYSFDNKKYWLDYHNDWTLRKGEPAQTKEVIVEKPVASASAPAVEMPAKRLSTSCQRVIAENFSSNNGSVTVQSSLADPKLYPVVCGHMVNNAALCPSSLYADMALTISDYIWKQMRPGTETPGYNVCNMEVPKPLIAQIPQPAEGQHIQL

>PKSI_2_120

MSNVLLFGDQTAEQYPLLNKIVLRKENALVITFVERCAKALREETNALPRSQRNAVPDFLTVNDLKEAYHQKGVKVPMVESALVTIAQIGHYIGYFSEHSAEQPSATNTRALGLCTGLLAAAAVVASKTVEELVLVGVDFVRLSFRSGAAVDAARTALCQTGDDNAPWSTIVTGTTEASAKEALAKFHEEKGIPQTSHAYISAVSVMAITVSGPPTTVKRFFEESPALSKNHRVPIPVYGPYHAEHLFGETEINKIASDSILEGLKQHQPVSLVHSAATGKALVAENAAELAKLVLAEMLQHSVRWDHLLEEAVSQITSKKAPAKIWAMGVSNVANSLVSALKAGGQSDVSTVDQSSWTENEPDTHGRTQNDKVAIVGMAGRFPNSADHEALWDLLMKGLDVHRRIPKDRFDADTHVDPSGKGKNKSHTPFGCFIDEPGFFDPRFFNMSPREAAQTDPMGRLALVTAYEALEMSGYVPNRTPSTKLHRIGTFYGQTSDDWREINAAENVDTYFITGGVRAFAPGRINYYFKFSGPSYSVDTACSSSLAAIQLACTSLWAGDCDTACAGGLNVLTNPDIFSGLSKGQFLSKTGSCKTYDNNADGYCRGDAVGTVILKRYEDAIADKDNILGCILGAATNHSAEAVSITHPHAGAQEFLYKRVLANAGVDAHEISYVEMHGTGTQAGDGIEMTSVTNVFAPRHRQRRDDQPVYLGAIKANVGHAEAASGINSLAKVLMMMKHNKIPANVGIKGEMNKTFPADLKDRKVNISQKAVEWPRNGKEKRKVFLNNFSAAGGNTALLLEDGPAYEAPTATDPRGTVPVTVTARSISALKRNIANLQKYVSENPSTTLTSMSYTLTARRIQHNYRVAFPLDQIDKFSDALQAQVKESYSPVPNVPTRVAFCFTGQGSQYTGLGQKLYNDLKSFRDDIDQLDHLARVQGLPSFLEIVQGADVQTLSPVKVQLGMACIQVALARMWAAWGITPAAVIGHSLGEYAALHVAGVISASDMVLLVGRRAELLVRDCTPHTHGMLAVKGGAEAIRNTLGNKMTEIACINGPEETVLCGSGDVVGAANETLAAKGFKATKLNVPFAFHSAQVDPILEQFKKIAASVTYNKPAVPVLSPLEGDIIREAGKINPEYLARHARETVNFWTALTAGQKEKVFDEKTAWLEVGAHPVCSGMVKASIGATTTAPSLRRGEDAWKTISNSMCTLFTAGVNFNFDEFHKEFNDAQEMYTLPTYSFDNKKYWLDYHNDWTLRKGEPAQTKEVIVEKPVASASAPAVEMPAKRLSTSCQRVIAENFSGNNGSVTVQSSLADPKLYPVVCGHMVNNAALCPSSLYADMALTISDYIWKQMRPGTETPGYNVCNMEVPKPLIA

>PKSI_1_14592

MPDNVSFMDESQDLRHIRETSRTSHTTSILNVEGDYRNNDMPREPGECNRSTNGTVDHERMTPSADGGIPIAICGIGLRLPGGIRNDRDLYDSLYNKKDARGVIPEDRFSIDSFHSAHGKTGTIITKHGYFLQDIDLTKFDVNMFNMTPAEVERLDPHQRILLETVRETLESAGEASFRGKKVGTYVGNFTDDWLDLQNVDTVDFATYQLHGKMDFSLANRISYEYDLRGPSMTIKTACSSSALAIHEAVYSIRNGECDAAIVSGSNLNLAPRLWVGMSSQGAISPDGSSKTFDESANGYARGDGIAALFIKRLDDAVRDGNPVRAVIRSTASNADGRTPGMTMPSTEAQEALIRRAYDAANLPLSETAMVECHGTGTAVGDPMEANAVARCFGDQGMLIGSVKPNLGHSEGASAITSVVKAVLSLENRTILPNIKFHRPNPAIPWSEAKLTVPVEPLAWPKDRQERISVNSFGIGGSNVHVVLDSAASMGFRPRSLAPSKDDRPGRLLLFSGGHRASVEQSSSQHQDYVTKYPNRLSDVAYTLAKRPPVKCPGLTRSTFVFTGQGAQWLHMGKELLHESPVFAKSIGRMDSVIHSLKHAPQWTLEGIINDPENPSALTNAEISQPLCTAVQIGLVDLLKSWAIYPHAVLGHSSGEIGAAYASGVVDRAEAILLAFYRGYVCRFAQKAGGMAAVGLEKSQVIKYLQPGVCVACENSGSSVTLSGDLETLEEVLQSIRAENLNAFARKLQVGIAYHSDHMKALGGLYHQYITEHLDPKDPQVPFFSSVSGRALHSKDDFGATYWQDNLENPVLFHTAVLKSLEHTGDKQVHLEVGPHGALNGPLRQIYAETGSKARYVALQKRGANCFDTFLEGIGQLYCNGVPLQYPESADDRTLIDLPPYPWHYDHSYWSETRVMKNWRFRRQLPHDLLGLRTLDCSDAEPMWRNILRITDLPWLRDHCVGKDVVFPASGYMCMAGEAVFQETGCRDYTLREVDISTAMVLSSDHSTELLTTMKKRRLNAFLDSRWYEFLIMSYDGASWTQHCSGLVTNGPSTSHPKALLQTYDRPVSTNRWYTAMSKIGLNYGPRFTGLQNITTHVQEKKASMTIMDKQEDYESPYALHPSTLDLILQSWTVASVRGEYRRFTQLFLPTFVDEFYIGNSASKLIHLNTTAIGPDGSARGEAIGKDNDGQISFNLKGFKGSKLDNVGVDQPQEMQTIMKQQWKRDFDFADTAQLMRPAFDSTSELSLLERMFVLAAIEVHVRTSGMEGKLPHHQRYKLWIDAQIRRFGEPGYPMVEDSMELLRLDSRERQRQLCRLLEHSRKTIAHPVAEAIWRALDRIEDVFDGRIEYLDLLFNDGLMPKFYDWSNSLSDVSRLFRLLSHKKPQLKILEVGAGTGGSTARLLQYLQSDFGERQYHSYTYTDVSSGFFVQAQERFKDYEGMKYRVLDISQDPFEQGFGADEFDLICASNVLHATPRLTETLRNCRKMLRPDGILFLQELCPRQQFMGFIMGLFEGWWLGAEDGRADTPLLLPPAWDRRLRDVGFEGVEAFSFDNNPPYFMAANMTARPTVTAKSKGSITLLTFNDFLDDVAGALKQAIRAAGFEIDHCVWGEHVPLDQSLISLVDLERDKPLLQDIGDDDLRIFLDLIDAVLQTTVIWLNKPAQVSSADPNAAQMLGLARTLRAELAMHFATVEMADPTLNGMSAVVHLMCMLQRGSALPENSLDQDMEYVWAHDAMHVSRFHFQPVDEALMEMSPKFDVKTLVPLQRGMLSSLKWIGTRMLPLAEAEVQIRMSAVGMNFHDMMIAMNMFDSPLTLGSGYNSIGMEGVGYVTRKSPEVDHVQVGDRVIVIGSNSSGFATDVHRPADYCIKCPSSLTDVEAAGMSFAYMTVLWSFLDKGGLRKGQSVLIHSAAGGVGIAALHVSRWLGLEAFVTVGNEEKVRFIMNNFGLPRNRIFNSHSATFLDDVMDATAGRGVDAALSAAAGELLHNTWSCIAPGGVMLEIGKRDLINRGRLSLAPFEENRSYVGIDFSRLTIVNKPAVVRLLRQTMRLVEQGHIHPIHPTTCFDAEHAEDAFRFMQTGQHIGRIVVKIPRDTSTIPLASRPPAPEFEGQKTYLLVGGMGGLGRSVASWMVSAGARNLIFMSRSAGKSEQDQNFARELELSGCKVHCCAIDITDTDAVREALERTQASVAGVLQMAMVLRDVGIMNMDKANWDAAVAPKVQGTWNLHHALPNVDFFVMFGSNSGTLGSYGQANYAAANAFLDSFVQYRQNLGQAASVIDIGAVGDVGYVAETQVAAENMESMAGRLISEQDFLNCLQLAIARSTPTERRNKGPSTEADGYIDLKQVILLNSSILSMADPSNQIFWRKDPRMGIYRNVQRTSAESPTTDSNSLRRLVATLKADSSIADQAELAQTLARELSKQVATVLMLGEDEIDVDRSLTDVGMDSLVAIEMRNWWKQNLGVDVSVLELKDGRSILRLGELAATRLKERYSRDS

>PKSI_2_14592

MSNVLLFGDQTAEQYPLLNKIVLRKENALVITFVERCAKALREETNALPRSQRNAVPDFLTVNDLKEAYHQKGVKVPMVESALVTIAQIGHYIGYFSEHSAEQPSATNTRALGLCTGLLAAAAVVASKTVEELVLVGVEFVRLSFRSGAAVDAARTALCQTGDDNAPWSTIVTGTTEASAKEALAKFHEEKGIPQTSHAYISAVSVMAITVSGPPTTVKRFFEESPALSKNHRVPIPVYGPYHAEHLFGETEINKIADASILEGLKQHQPVSLVHSAATGKALVAENAAELAKLVLAEMLQHPVRWDHLLEEAVSQITSKKAPAKIWAMGVSNVANSLVSALKAGGQSDVSTVDQSSWTENEPDTHGRTQNDKVAIVGMAGRFPNSADHEALWDLLMKGLDVHRRIPKDRFDADTHVDPSGKGKNKSHTPFGCFIDEPGFFDPRFFNMSPREAAQTDPMGRLALVTAYEALEMSGYVPNRTPSTKLHRIGTFYGQTSDDWREINAAENVDTYFITGGVRAFAPGRINYYFKFSGPSYSVDTACSSSLAAIQLACTSLWAGDCDTACAGGLNVLTNPDIFSGLSKGQFLSKTGSCKTYDNNADGYCRGDAVGTVILKRYEDAIADKDNILGCILGAATNHSAEAVSITHPHAGAQEFLYKRVLANAGVDAHEISYVEMHGTGTQAGDGIEMTSVTNVFAPRHRQRRDDQPVYLGAIKANVGHAEAASGINSLAKVLMMMKHNKIPANVGIKGEMNKTFPADLKDRKVNISQKAVEWPRNGKEKRKVFLNNFSAAGGNTALLLEDGPAYEAPTASDPRGTVPVTVTARSISALKRNIANLQKYVSENPSTTLTSMSYTLTARRIQHNYRVAFPLDQINKFSDALQAQVKESYSPVPNAPTRVAFCFTGQGSQYTGLGQKLYNDLKSFRDDIDQLDHLARVQGLPSFLEIVQGADVQTLSPVKVQLGMACIQVALARMWAAWGITPAAVIGHSLGEYAALHVAGVISASDMVLLVGRRAELLVRDCTPHTHGMLAVKGGAEAIRNTLGNKMTEIACINGPEETVLCGSGDVVGAANETLAAKGFKATKLNVPFAFHSAQVDPILEQFKKIAASVTYNKPAVPVLSPLEGDIIREAGKINPEYLARHARETVNFWTALTAGQKEKVFDEKTAWLEVGAHPVCSGMVKASIGATTTAPSLRRGEDAWKTISNSMCTLFTAGVNFNFDEFHKEFNDAQEMYTLPTYSFDNKKYWLDYHNDWTLRKGEPAQTKEVIVEKPVASASAPAVEIPAKRLSTSCQRVISENFSGNNGSVTVQSSLADPKLYPVVCGHMVNNAALCPSSLYADMALTISDYIWKQMRPGTETPGYNVCNMEVPKPLIAQIPQPAEGQHIQLEANADLDSGIVKLNFRSVKPDGQKLQDHAHCIVRLEDRAAWEDEWSRYNYMVQAQMELLQHKTLNGGAHKVQRGMAYKLFKALVNYDEKYRAMAEVVLASGQTEASAVLDFPTKPEDGDFYCPPYHIDGSCHISGFIVNASDLLDSEQNVYVSHGWGAMKFSRPLTAGMKLRNYVRMQPQPNNVSKGDVYIMEGDQIVAVCEGIKFQQIPRRVLNTFLPPNKGSGPASAAKPAAAPVAAARPAPAAAPIKTAPAPAKAAPAPAPAAPKAAPKPKKAAAPKKAAGGLTAKVMKILAKETEVDEGELVDEAQFENLGVDSLLSLTISAVFREELDMDISSTLFTDYPTVGDMKKYFAQFDNGSSTSSSAEEEDSDEDSIPPTDAATPMDDLSTPASSVPSSAPSDAGKPDSPTRETLEDVGDVSLAKHIVAQEMGVDIAEVTDDADLAEMGMDSLMSLTILGELREKTGIDLPSTFLTTNPTMKDIDNALGMRPKPKAAPKPAAPKAAAPSSSKKTDMNEVSARLSALNNNTDISRYPNATSVLLQGNPKQATKKIFFLPDGSGSATSYVSIPNLGPDVCAYGLNCPFMKNPEQWQCGIEISALVYLAEIKRRQPQGPYIIGGWSAGGVIAYSVAQALLAANEGVEKLLLLDSPCPVNLAPLPARLHNFFNEIGLLGTGDPAKTPKWLLPHFSAAIRSLSDYDPKPSLRPIPTYAIWCREGVAGNPGDPRPPPAEEEDPAPMTWLLEHRTDFKDNGWAQLCGDSMKFGVMGGHHFSMMKPPHADDLGNLIREGLDWQP

>PKSI_1_14591

MSNVLLFGDQTAEQYPLLNKIVLRKENALVITFVERCAKALREETNALPRSQRNAVPDFLTVNDLKEAYHQKGVKVPMVESALVTIAQIGHYIGYFSEHSAEQPSATNTRALGLCTGLLAAAAVVASKTVEELVLVGVEFVRLSFRSGAAVDAARTALCQTGDDNAPWSTIVTGTTEASAKEALAKFHEEKGIPQTNHAYISAVSVMAITVSGPPTTVKRFFEESPALSKNHRVPIPVYGPYHAEHLFGETEINKIASASILEGLKQHQPVSLVHSAATGKALVAENAAELAKLVLAEMLQLPVRWDHLLEEAVSQITSKKAPAKIWAMGVSNVANSLVSALKAGGQTDVSTVDQGTWTENEPDTHGRTQNDKVAIVGMAGRFPNSADHEALWELLMKGLDVHRRIPKDRFDADTHVDPSGKGKNKSHTPFGCFIDEPGFFDPRFFNMSPREAAQTDPMGRLALVTAYEALEMSGYVPNRTPSTKLHRIGTFYGQTSDDWREINAAENVDTYFITGGVRAFAPGRINYYFKFSGPSYSVDTACSSSLAAIQLACTSLWAGDCDTACAGGLNVLTNPDIFSGLSKGQFLSKTGSCKTYDNNADGYCRGDAVGTVILKRYEDAIADKDNILGCILGAATNHSAEAVSITHPHAGAQEFLYKRVLANAGVDAHEISYVEMHGTGTQAGDGIEMTSVTNVFAPRHRQRRDDQPVYLGAIKANVGHAEAASGINSLAKVLLMMKHNKIPANVGIKGEMNKTFPADLKDRKVNISQKAVDWPRNGKEKRKVFLNNFSAAGGNTALLLEDGPAYEAPTATDPRGTVPVTVTARSISALKRNIANLQKYVSENPSTTLTSMSYTLTARRIQHNYRVAFPLDQINKFSDALQAQVKESYSPVPNAPTRVAFCFTGQGSQYTGLGQKLYNDLKSFRDDIDQLDHLARVQGLPSFLEIVQGADVQTLSPVKVQLGMACIQVALARMWAAWGITPAAVIGHSLGEYAALHVAGVISASDMVLLVGRRAELLVRDCTPHTHGMLAVKGGAEAIRNTLGNKMTEIACINGPEETVLCGSGDVVGAANETLAAKGFKATKLNVPFAFHSAQVDPILEQFKKIAASVTYNKPAVPVLSPLEGDIIREAGKINPEYLARHARETVNFWTALTAGQKEKVFDEKTAWLEVGAHPVCSGMVKASIGATTTAPSLRRGEDAWKTISNSMCTLFTAGVNFNFDEFHKEFNDAQEMYTLPTYSFDNKKYWLDYHNDWTLRKGEPAQTKEVIVEKPVASASAPAVEMPAKRLSTSCQRVIAENFSGNNGSVTVQSSLADPKLYPVVCGHMVNNAALCPSSLYADMALTISDYIWKQMRPGTETPGYNVCNMEVPKPLIAQIPQPAEGQHIQLEANADLDSGIVKLNFRSVKPDGQKLQDHAHCIVRLEDKAAWEDEWSRYNYMVQAQMELLQHKTLNGGAHKVQRGMAYKLFKALVNYDEKYRAMAEVVLASGQTEASAMLDFPTKPEDGDFYCPPYHIDGSCHISGFIVNASDLLDSEQNVYVSHGWGAMKFSRPLTAGMKLRNYVRMQPQPNNVSKGDVYIMEGDQIVAVCEGIKFQQIPRRVLNTFLPPNKGSGPASAAKPAAAPVAAARPAPAAAPIKTAPAPAKAAPAPAPAAPKAAPKPKKAAAPKKPAGGLTAKVMKILAKETEVDEGELVDEAQFENLGVDSLLSLTIS

>PKSI_2_14591

MPDNVSFMDESQDLRHIRETSRTSHTTSILNVEGDYRNNDMPREPGECNRSTNGTVDHERMTPSADGGIPIAICGIGLRLPGGIRNDRDLYDSLYNKKDARGVIPEDRFSIDSFHSAHGKTGTIITKHGYFLQDIDLTKFDVNMFNMTPAEVERLDPHQRILLETVRETLESAGEASFRGKKVGTYVGNFTDDWLDLQNVDTVDFATYQLHGKMDFSLANRISYEYDLRGPSMTIKTACSSSALAIHEAVYSIRNGECDAAIVSGSNLNLAPRLWVGMSSQGAISPDGSSKTFDESANGYARGDGIAALFIKRLDDAVRDGNPVRAVIRSTASNADGRTPGMTMPSTEAQEALIRRAYDAANLPLSETAMVECHGTGTAVGDPMEANAVARCFGDQGMLIGSVKPNLGHSEGASAITSVVKAVLSLENRTILPNIKFHRPNPAIPWSEAKLTVPVEPLAWPKDRQERISVNSFGIGGSNVHVVLDSAASMGFRPRSLAPSKDDRPGRLLLFSGGHRASVEQSSSQHQDYVTKYPNRLSDVAYTLAKRPPVKCPGLTRSTFVFTGQGAQWLHMGKELLHESPVFAKSIGRMDSVIHSLKHAPQWTLEGIINDPENPSALTNAEISQPLCTAVQIGLVDLLKSWAIYPHAVLGHSSGEIGAAYASGVVDRAEAILLAFYRGYVCRFAQKAGGMAAVGLEKSQVIKYLQPGVCVACENSGSSVTLSGDLETLEEVLQSIRAENLNAFARKLQVGIAYHSDHMKALGGLYHQYITEHLDPKDPQVPFFSSVSGRALHSKDDFGATYWQDNLENPVLFHTAVLKSLEHTGDKQVHLEVGPHGALNGPLRQIYAETGSKARYVALQKRGANCFDTFLEGIGQLYCNGVPLQYPESADDRTLIDLPPYPWHYDHSYWSETRVMKNWRFRRQLPHDLLGLRTLDCSDAEPMWRNILRITDLPWLRDHCVGKDVVFPASGYMCMAGEAVFQETGCRDYTLREVDISTAMVLSSDHSTELLTTMKKRRLNAFLDSRWYEFLIMSYDGASWTQHCSGLVTNGPSTSHPKALLQTYDRPVSTNRWYTAMSKIGLNYGPRFTGLQNITTHVQEKKASMTIMDKQEDYESPYALHPSTLDLILQSWTVASVRGEYRRFTQLFLPTFVDEFYIGNSASKLIHLNTTAIGPDGSARGEAIGKDNDGQISFNLKGFKGSKLDNVGVDQPQEMQTIMKQQWKRDFDFADTAQLMRPAFDSTSELSLLERMFVLAAIEVHVRTSGMEGKLPHHQRYKLWIDAQIRRFGEPGYPMVEDSMELLRLDSRERQRQLCRLLEHSRKTTAHPVAEAIWRALDRIEDVFDGRIEYLDLLFNDGLMPKFYDWSNSLSDVSRLFRLLSHKKPQLKILEVGAGTGGSTARLLQYLQSDFGERQYHSYTYTDVSSGFFVQAQERFKDYEGMKYRVLDISQDPFEQGFGADEFDLICASNVLHATPRLTETLRNCRKMLRPDGILFLQELCPRQQFMGFIMGLFEGWWLGAEDGRADTPLLLPPAWDRRLRDVGFEGVEAFSFDNNPPYFMAANMTARPTVTAKSKGSITLLTFNDFLDDVAGALKQAIRAAGFEIDHCVWGEHVPLDQSLISLVDLERDKPLLQDIGDDDLRIFLDLIDAVLQTTVIWLNKPAQVSSADPNAAQMLGLARTLRAELAMHFATVEMADPTLNGMSAVVHLMCMLQRGSALPENSLDQDMEYVWAHDAMHVSRFHFQPVDEALMEMSPKFDVKTLVPLQRGMLSSLKWIGTRMLPLAEAEVQIRMSAVGMNFHDMMIAMNMFDSPLTLGSGYNSIGMEGVGYVTRKSPEVDHVQVGDRVIVIGSNSSGFATDVHRPADYCIKCPSSLTDVEAAGMSFAYMTVLWSFLDKGGLRKGQSVLIHSAAGGVGIAALHVSRWLGLEAFVTVGNEEKVRFIMNNFGLPRNRIFNSHSATFLDDVMDATAGRGVDAALSAAAGELLHNTWSCIAPGGVMLEIGKRDLINRGRLSLAPFEENRSYVGIDFSRLTIVNKPAVVRLLRQTMRLVEQGHIHPIHPTTCFDAEHAEDAFRFMQTGQHIGRIVVKIPRDTSTIPLASRPPAPEFEGQKTYLLVGGMGGLGRSVASWMVSAGARNLIFMSRSAGKSEQDQNFARELELSGCKVHCCAIDITDTDAVREALERTQASVAGVLQMAMVLRDVGIMNMDKANWDAAVAPKVQGTWNLHHALPNVDFFVMFRSNSGTLGSYGQANYAAANAFLDSFVQYRQNLGQAASVIDIGAVGDVGYVAETQVAAENMESMAGRLISEQDFLNCLQLAIARSTPTERRNKGPSTEADGYIDLKQVILLNSSILSMADPSNQIFWRKDPRMGIYRNVQRTSAESPTTDSNSLRRLVATLKADSSIADQAELAQTLARELSKQVATVLMLGEDEIDVDRSLTDVGMDSLVAIEMRNWWKQNLGVDVSVLELKDGRSILRLGELAATRLKERYSRDS

>PKSI_3_14591

MSNVLLFGDQTAEQYPLLNKIVLRKENALVITFVERCAKALREETNALPRSQRNAVPDFLTVNDLKEAYHQKGVKVPMVESALVTIAQIGHYIGYFSEHSAEQPSATNTRALGLCTGLLAAAAVVASKTVEELVLVGVGFVRLSFRSGAAVDAARTALCQTGDDNAPWSTIVTGTTEASAKEALAKFHEEKGIPQTSHAYISAVSVMAITVSGPPTTVKRFFEESPALSKNHRVPIPVYGPYHAEHLFGETEINKIASDSILEGLKQHQPVSLVHSAATGKALVAENAAELAKLVLAEMLQHSVRWDHLLEEAVSQITSKKAPAKIWAMGVSNVANSLVSALKAGGQSDVSTVDQSSWTENEPDTHGRTQNDKVAIVGMAGRFPNSADHEALWDLLMKGLDVHRRIPKDRFDADTHVDPSGKGKNKSHTPFGCFIDEPGFFDPRFFNMSPREAAQTDPMGRLALVTAYEALEMSGYVPNRTPSTKLHRIGTFYGQTSDDWREINAAENVDTYFITGGVRAFAPGRINYYFKFSGPSYSVDTACSSSLAAIQLACTSLWAGDCDTACAGGLNVLTNPDIFSGLSKGQFLSKTGSCKTYDNNADGYCRGDAVGTVILKRYEDAIADKDNILGCILGAATNHSAEAVSITHPHAGAQEFLYKRVLANAGVDAHEISYVEMHGTGTQAGDGIEMTSVTNVFAPRHRQRRDDQPVYLGAIKANVGHAEAASGINSLAKVLMMMKHNKIPANVGIKGEMNKTFPADLKDRKVNISQKAVEWPRNGKEKRKVFLNNFSAAGGNTALLLEDGPAYEAPTATDPRGTVPVTVTARSISALKRNIANLQKYVSENPSTTLTSMSYTLTARRIQHNYRVAFPLDQIDKFSDALQAQVKESYSPVPNVPTRVAFCFTGQGSQYTGLGQKLYNDLKSFRDDIDQLDHLARVQGLPSFLEIVQGADVQTLSPVKVQLGMACIQVALARMWAAWGITPAAVIGHSLGEYAALHVAGVISASDMVLVVGRRAELLVRDCTPHTHGMLAVKGGAEAIRNTLGNKMTEIACINGPEETVLCGSGDVVGAANETLAAKGFKATKLNVPFAFHSAQVDPILEQFKKIAASVTYNKPAVPVLSPLEGDIIREAGKINPEYLARHARETVNFWTALTAGQKEKVFDEKTAWLEVGAHPVCSGMVKASIGATTTAPSLRRGEDAWKTISNSMCTLFTAGVNFNFDEFHKEFNDAQEMYTLPTYSFDNKKYWLDYHNDWTLRKGEPAQTKEVIVEKPVASASAPAVEMPAKRLSTSCQRVIAENFSGNNGSVTVQSSLADPKLYPVVCGHMVNNAALCPSSLYADMALTISDYIWKQMRPGTETPGYNVCNMEVPKPLIAQIPQPAEGQHIQLEANADLDSGIVKLNFRSVKPDGQKLQDHAHCIVRLEDRAAWEDEWSRYNYMVQAQMELLQHKTLNGGAHKVQRGMAYKLFKALVNYDEKYRAMAEVVLASGQTEASAVLDFPTKPEDGDFYCPPYHIDGSCHISGFIVNASDLLDSEQNVYVSHGWGAMKFSRPLTAGMKLRNYVRMQPQPNNVSKGDVYIMEGDQIVAVCEGIKFQQIPRRVLNTFLPPNKGSGPASAAKPAAAPVAAARPAPAAAPIKTAPAPAKAAPAPAPAAPKAAPKPKKAAAPKKAAGGLTAKVMKILAKETEVDEGELVDEAQFENLGVDSLLSLTISAVFREELDMDISSTLFTDYPTVGDMKKYFAQFDNGSSTSSSAEEEDSDEDSIPPTDAATPMDDLSTPASSVPSSAPSDAGKPDSPTRETLEDVGDVSLAKHIVAQEMGVDIAEVTDDADLAEMGMDSLMSLTILGELREKTGIDLPSTFLTTNPTMKDIDNALGMRPKPKAAPKPAAPKAAAPSSSKKTDMNEVSARLSALNNNTDISRYPNATSVLL

>PKSI_1_12619

MPDNVSFMDESQDLRHIRETSRTSHTTSILNVEGDYRNNDMPREPGECNRSTNGTVDHERMTPSADGGIPIAICGIGLRLPGGIRNDRDLYDSLYNKKDARGVIPEDRFSIDSFHSAHGKTGTIITKHGYFLQDIDLTKFDVNMFNMTPAEVERLDPHQRILLETVRETLESAGEASFRGKKVGTYVGNFTDDWLDLQNVDTVDFATYQLHGKMDFSLANRISYEYDLRGPSMTIKTACSSSALAIHEAVYSIRNGECDAAIVSGSNLNLAPRLWVGMSSQGAISPDGSSKTFDESANGYARGDGIAALFIKRLDDAVRDGNPVRAVIRSTASNADGRTPGMTMPSTEAQEALIRRAYDAANLPLSETAMVECHGTGTAVGDPMEANAVARCFGDQGMLIGSVKPNLGHSEGASAITSVVKAVLSLENRTILPNIKFHRPNPAIPWSEAKLTVPVEPLAWPKDRQERISVNSFGIGGSNVHVVLDSAASMGFRPRSLAPSKDDRPGRLLLFSGGHRASVEQSSSQHQDYVTKYPNRLSDVAYTLAKRPPVKCPGLTRSTFVFTGQGAQWLHMGKELLHESPVFAKSIGRMDSVIHSLKHAPQWTLEGIINDPENPSALTNAEISQPLCTAVQIGLVDLLKSWAIYPHAVLGHSSGEIGAAYASGVVDRAEAILLAFYRGYVCRFAQKAGGMAAVGLEKSQVIKYLQPGVCVACENSGSSVTLSGDLETLEEVLQSIRAENLNAFARKLQVGIAYHSDHMKALGGLYHQYITEHLDPKDPQVPFFSSVSGRALHSKDDFGATYWQDNLENPVLFHTAVLKSLEHTGDKQVHLEVGPHGALNGPLRQIYAETGSKARYVALQKRGANCFDTFLEGIGQLYCNGVPLQYPESADDRTLIDLPPYPWHYDHSYWSETRVMKNWRFRRQLPHDLLGLRTLDCSDAEPMWRNILRITDLPWLRDHCVGKDVVFPASGYMCMAGEAVFQETGCRDYTLREVDISTAMVWSSDHSTELLTTMKKRRLNAFLDSRWYEFLIMSYDGASWTQHCSGLVTNGPSTSHPKALLQTYDRPVSTNRWYTAMSKIGLNYGPRFTGLQNITTHVQEKKASMTIMDKQEDYESPYALHPSTLDLILQSWTVASVRGEYRRFTQLFLPTFVDEFYIGNSASKLIHLNTTAIGPDGSARGEAIGKDNDGQISFNLKGFKGSKLDNVGVDQPQEMQTIMKQQWKRDFDFADTAQLMRPAFDSTSELSLLERMFVLAAIEVHVRTSGMEGKLPHHQRYKLWIDAQIRRFGEPGYPMVEDSMELLRLDSRERQRQLCRLLEHSRKTTAHPVAEAIWRALDRIEDVFDGRIEYLDLLFNDGLMPKFYDWSNSLSDVSRLFRLLSHKKPQLKILEVGAGTGGSTARLLQYLQSDFGERQYHSYTYTDVSSGFFVQAQERFKDYEGMKYRVLDISQDPFEQGFGADEFDLICASNVLHATPRLTETLRNCRKMLRPDGILFLQELCPRQQFMGFIMGLFEGWWLGAEDGRADTPLLLPPAWDRRLRDVGFEGVEAFSFDNNPPYFMAANMTARPTVTAKSKGSITLLTFNDFLDDVAGALKQAIRAAGFEIDHCVWGEHVPLDQSLISLVDLERDKPLLQDIGDDDLRIFLDLIDAVLQTTVIWLNKPAQVSSADPNAAQMLGLARTLRAELAMHFATVEMADPTLNGMSAVVHLMCMLQRGSALPENSLDQDMEYVWAHDAMHVSRFHFQPVDEALMEMSPKFDVKTLVPLQRGMLSSLKWIGTRMLPLAEAEVQIRMSAVGMNFHDMMIAMNMFDSPLTLGSGYNSIGMEGVGYVTRKSPEVDHVQVGDRVIVIGSNSSGFATDVHRPADYCIKCPSSLTDVEAAGMSFAYMTVLWSFLDKGGLRKGQSVLIHSAAGGVGIAALHVSRWLGLEAFVTVGNEEKVRFIMNNFGLPRNRIFNSHSATFLDDVMDATAGRGVDAALSAAAGELLHNTWSCIAPGGVMLEIGKRDLINRGRLSLAPFEENRSYVGIDFSRLTIVNKPAVVRLLRQTMRLVEQGHIHPIHPTTCFDAEHAEDAFRFMQTGQHIGRIVVKIPRDTSTIPLASRPPAPEFEGQKTYLLVGGMGGLGRSVASWMVSAGARNLIFMSRSAGKSEQDQNFARELELSGCKVHCCAIDITDTDAVREALERTQASVAGVLQMAMVLRDVGIMNMDKANWDAAVAPKVQGTWNLHHALPNVDFFVMFGSNSGTLGSYGQANYAAANAFLDSFVQYRQNLGQAASVIDIGAVGDVGYVAETQVAAENMESMAGRLISEQDFLNCLQLAIARSTPTERRNKGPSTEADGYIDLKQVILLNSSILSMADPSNQIFWRKDPRMGIYRNVQRTSAESPTTDSNSLRRLVATLKADSSIADQAELAQTLARELSKQVATVLMLGEDEIDVDRSLTDVGMDSLVAIEMRNWWKQNLGVDVSVLELKDGRSILRLGELAATRLKERYSRDS

>PKSI_2_12619

MSNVLLFGDQTAEQYPLLNKIVLRKENALVITFVERCAKALREETNALPRSQRNAVPDFLTVNDLKEAYHQKGVKVPMVESALVTIAQIGHYIGYFSEHSAEQPSATNTRALGLCTGLLAAAAVVASKTVEELVLVGVEFVRLSFRSGAAVDAARTALCQTGDDNAPWSTIVTGTTEAAAKEALAKFHEEKGIPQTSHAYISAVSVMAITVSGPPTTVKRFFEESPALSKNHRVPIPVYGPYHAEHLFGETEINKIASASILEGLKQHQPVSLVHSAATGKALVAENAAELAKLVLAEMLQHPVRWDHLLEEAVSQITSKKAPAKIWAMGVSNVANSLVSALKAGGQTDVSTVDQSTWTENEPDTHGRTQNDKVAIVGMAGRFPNSADHEALWELLMKGLDVHRRIPKDRFDADTHVDPSGKGKNKSHTPFGCFIDEPGFFDPRFFNMSPREAAQTDPMGRLALVTAYEALEMSGYVPNRTPSTKLHRIGTFYGQTSDDWREINAAENVDTYFITGGVRAFAPGRINYYFKFSGPSYSVDTACSSSLAAIQLACTSLWAGDCDTACAGGLNVLTNPDIFSGLSKGQFLSKTGSCKTYDNNADGYCRGDAVGTVILKRYEDAIADKDNILGCILGAATNHSAEAVSITHPHAGAQEFLYKRVLANAGVDAHEISYVEMHGTGTQAGDGIEMTSVTNVFAPRHRQRRDDQPVYLGAIKANVGHAEAASGINSLAKVLLMMKHNKIPANVGIKGEMNKTFPADLKDRKVNISQKAVDWPRNGKEKRKVFLNNFSAAGGNTALLLEDGPAYEAPTATDPRGTVPVTVTARSISALKRNIANLQKYVSENPSTTLTSMSYTLTARRIQHNYRVAFPLDQINKFSDALQAQVKESYSPVPNAPTRVAFCFTGQGSQYTGLGQKLYNDLKSFRDDIDQLDHLARVQGLPSFLEIVQGADVQTLSPVKVQLGMACIQVALARMWAAWGITPAAVIGHSLGEYAALHVAGVISASDMVLLVGRRAELLVRDCTPHTHGMLAVKGGAEAIRNTLGSKMTEIACINGPEETVLCGSGDVVGAANETLAAKGFKATKLNVPFAFHSAQVDPILEQFKKIAASVTYNKPAVPVLSPLEGDIIREAGKINPEYLARHARETVNFWTALTAGQKEKVFDEKTAWLEVGAHPVCSGMVKASIGATTTAPSLRRGEDAWKTISNSMCTLFTAGVNFNFDEFHKEFNDAQEMYTLPTYSFDNKKYWLDYHNDWTLRKGEPAQTKEVIVEKPVASASAPAVEMPAKRLSTSCQRVIAENFSGNNGSVTVQSSLADPKLYPVVCGHMVNNAALCPSSLYADMALTISDYIWKQMRPGTETPGYNVCNMEVPKPLIAQIPQPAEGQHIQLEANADLDSGIVKLNFRSVKPDGQKLQDHAHCIVRLEDRAAWEDEWSRYNYMVQAQMELLQHKTLNGGAHKVQRGMAYKLFKALVNYDEKYRAMAEVVLASGQTEASAVLDFPTKPEDGDFYCPPYHIDGSCHISGFIVNASDLLDSEQNVYVSHGWGAMKFSRPLTAGMKLRNYVRMQPQPNNVSKGDVYIMEGDQIVAVCEGIKFQQIPRRVLNTFLPPNKGSGPASAAKPAAAPVAAARPAPAAAPIKTAPAPAKAAPAPAPAAPKAAPKPKKAAAPKKAAGGLTAKVMKILAKETEVDEGELVDEAQFENLGVDSLLSLTISAVFREELDMDISSTLFTDYPTVGDMKKYFAQFDNGSSTSSSTEEEDSDEDSIPPTDAATPMDDLSTPASSVGSSAPSDAGKPDSPTRETLEDVGDVSLAKHIVAQEMGVDIAEVTDDADLAEMGMDSLMSLTILGELREKTGIDLPSTFLTTNPTMKDIDNALGMRPKPKAAPKPAAPKAAAPSSSKKTDMNEVSARLSALNNNTDISRYPNATSVLLQGNPKQATKKIFFLPDGSGSATSYVSIPNLGPDVCAYGLNCPFMKNPEQWQCGIEISALVYLAEIKRRQPQGPYIIGGWSAGGVIAYSVAQALLAANEGVEKLLLLDSPCPVNLAPLPARLHNFFNEIGLLGTGDPAKTPKWLLPHFSAAIRSLSDYDPKPSLRPIPTYAIWCREGVAGNPGDPRPPPAEEEDPAPMTWLLEHRTNFKDNGWAQLCGDSMKFGVMGGHHFSMMKPPHADDLGNLIREGLDWQP

>PKSI_1_14590

MPDNVSFMDESQDLRHIRETSRTSHTTSILNVEGDYRNNDMPREPGECNRSTNGTVDHERMTPSADGGIPIAICGIGLRLPGGIRNDRDLYDSLYNKKDARGVIPEDRFSIDSFHSAHGKTGTIITKHGYFLQDIDLTKFDVNMFNMTPAEVERLDPHQRILLETVRETLESAGEASFRGKKVGTYVGNFTDDWLDLQNVDTVDFATYQLHGKMDFSLANRISYEYDLRGPSMTIKTACSSSALAIHEAVYSIRNGECDAAIVSGSNLNLAPRLWVGMSSQGAISPDGSSKTFDESANGYARGDGIAALFIKRLDDAVRDGNPVRAVIRSTASNADGRTPGMTMPSTEAQEALIRRAYDAANLPLSETAMVECHGTGTAVGDPMEANAVARCFGDQGMLIGSVKPNLGHSEGASAITSVVKAVLSLENRTILPNIKFHRPNPAIPWSEAKLTVPVEPLAWPKDRQERISVNSFGIGGSNVHVVLDSAASMGFRPRSLAPSKDDRPGRLLLFSGGHRASVEQSSSQHQDYVTKYPNRLSDVAYTLAKRPPVKCPGLTRSTFVFTGQGAQWLHMGKELLHESPVFAKSIGRMDSVIHSLKHAPQWTLEGIINDPENPSALTNAEISQPLCTAVQIGLVDLLKSWAIYPHAVLGHSSGEIGAAYASGVVDRAEAILLAFYRGYVCRFAQKAGGMAAVGLEKSQVIKYLQPGVCVACENSGSSVTLSGDLETLEEVLQSIRAENLNAFARKLQVGIAYHSDHMKALGGLYHQYITEHLDPKDPQVPFFSSVSGRALHSKDDFGATYWQDNLENPVLFHTAVLKSLEHTGDKQVHLEVGPHGALNGPLRQIYAETGSKARYVALQKRGANCFDTFLEGIGQLYCNGVPLQYPESADDRTLIDLPPYPWHYDHSYWSETRVMKNWRFRRQLPHDLLGLRTLDCSDAEPMWRNILRITDLPWLRDHCVGKDVVFPASGYMCMAGEAVFQETGCRDYTLREVDISTAMVLSSDHSTELLTTMKKRRLNAFLDSRWYEFLIMSYDGASWTQHCSGLVTNGPSTSHPKALLQTYDRPVSTNRWYTAMSKIGLNYGPRFTGLQNITTHVQEKKASMTIMDKQEDYESPYALHPSTLDLILQSWTVASVRGEYRRFTQLFLPTFVDEFYIGNSASKLIHLNTTAIGPDGSARGEAIGKDNDGQISFNLKGFKGSKLDNVGVDQPQEMQTIMKQQWKRDFDFADTAQLMRPAFDSTSELSLLERMFVLAAIEVHVRTSGMEGKLPHHQRYKLWIDAQIRRFGEPGYPMVEDSMELLRLDSRERQRQLCRLLEHSRKTTAHPVAEAIWRALDRIEDVFDGRIEYLDLLFNDGLMPKFYDWSNSLSDVSRLFRLLSHKKPQLKILEVGAGTGGSTARLLQYLQSDFGERQYHSYTYTDVSSGFFVQAQERFKDYEGMKYRVLDISQDPFEQGFGADEFDLICASNVLHATPRLTETLRNCRKMLRPDGILFLQELCPRQQFMGFIMGLFEGWWLGAEDGRADTPLLLPPAWDRRLRDVGFEGVEAFSFDNNPPYFMAANMTARPTVTAKSKGSITLLTFNDFLDDVAGALKQAIRAAGFEIDHCVWGEHVPLDQSLISLVDLERDKPLLQDIGDDDLRIFLDLIDAVLQTTVIWLNKPAQVSSADPNAAQMLGLARTLRAELAMHFATVEMADPTLNGMSAVVHLMCMLQRGSALPENSLDQDMEYVWAHDAMHVSRFHFQPVDEALMEMSPKFDVKTLVPLQRGMLSSLKWIGTRMLPLAEAEVQIRMSAVGMNFHDMMIAMNMFDSPLTLGSGYNSIGMEGVGYVTRKSPEVDHVQVGDRVIVIGSNSSGFATDVHRPADYCIKCPSSLTDVEAAGMSFAYMTVLWSFLDKGGLRKGQSVLIHSAAGGVGIAALHVSRWLGLEAFVTVGNEEKVRFIMNNFGLPRNRIFNSHSATFLDDVMDATAGRGVDAALSAAAGELLHNTWSCIAPGGVMLEIGKRDLINRGRLSLAPFEENRSYVGIDFSRLTIVNKPAVVRLLRQTMRLVEQGHIHPIHPTTCFDAEHAEDAFRFMQTGQHIGRIVVKIPRDTSTIPLASRPPAPEFEGQKTYLLVGGMGGLGRSVASWMVSAGARNLIFMSRSAGKSEQDQNFARELELSGCKVHCCAIDITDTDAVREALERTQASVAGVLQMAMVLRDVGIMNMDKANWDAAVAPKVQGTWNLHHALPNVDFFVMFRSNSGTLGSYGQANYAAANAFLDSFVQYRQNLGQAASVIDIGAVGDVGYVAETQVAAENMESMAGRLISEQDFLNCLQLAIARSTPTERRNKGPSTEADGYIDLKQVILLNSSILSMADPSNQIFWRKDPRMGIYRNVQRTSAESPTTDSNSLRRLVATLKADSSIADQAELAQTLARELSKQVATVLMLGEDEIDVDRSLTDVGMDSLVAIEMRNWWKQNLGVDVSVLELKDGRSILRLGELAATRLKERYSRDS

>PKSI_2_14590

MSNVLLFGDQTAEQYPLLNKIVLRKENALVITFVERCAKALREETNALPRSQRNAVPDFLTVNDLKEAYHQKGVKVPMVESALVTIAQIGHYIGYFSEHSAEQPSATNTRALGLCTGLLAAAAVVASKTVEELVLVGVDFVRLSFRSGAAVDAARTALCQTGDDNAPWSTIVTGTTEASAKEALAKFHEEKGIPQTSHAYISAVSVMAITVSGPPTTVKRFFEESPALSKNHRVPIPVYGPYHAEHLFGETEINKIASDSILEGLKQHQPVSLVHSAATGKALVAENAAELAKLVLAEMLQHSVRWDHLLEEAVSQITSKKAPAKIWAMGVSNVANSLVSALKAGGQSDVSTVDQSSWTENEPDTHGRTQNDKVAIVGMAGRFPNSADHEALWDLLMKGLDVHRRIPKDRFDADTHVDPSGKGKNKSHTPFGCFIDEPGFFDPRFFNMSPREAAQTDPMGRLALVTAYEALEMSGYVPNRTPSTKLHRIGTFYGQTSDDWREINAAENVDTYFITGGVRAFAPGRINYYFKFSGPSYSVDTACSSSLAAIQLACTSLWAGDCDTACAGGLNVLTNPDIFSGLSKGQFLSKTGSCKTYDNNADGYCRGDAVGTVILKRYEDAIADKDNILGCILGAATNHSAEAVSITHPHAGAQEFLYKRVLANAGVDAHEISYVEMHGTGTQAGDGIEMTSVTNVFAPRHRQRRDDQPVYLGAIKANVGHAEAASGINSLAKVLMMMKHNKIPANVGIKGEMNKTFPADLKDRKVNISQKAVEWPRNGKEKRKVFLNNFSAAGGNTALLLEDGPAYEAPTATDPRGTVPVTVTARSISALKRNIANLQKYVSENPSTTLTSMSYTLTARRIQHNYRVAFPLDQIDKFSDALQAQVKESYSPVPNVPTRVAFCFTGQGSQYTGLGQKLYNDLKSFRDDIDQLDHLARVQGLPSFLEIVQGADVQTLSPVKVQLGMACIQVALARMWAAWGITPAAVIGHSLGEYAALHVAGVISASDMVLLVGRRAELLVRDCTPHTHGMLAVKGGAEAIRNTLGNKMTEIACINGPEETVLCGSGDVVGAANETLAAKGFKATKLNVPFAFHSAQVDPILEQFKKIAASVTYNKPAVPVLSPLEGDIIREAGKINPEYLARHARETVNFWTALTAGQKEKVFDEKTAWLEVGAHPVCSGMVKASIGATTTAPSLRRGEDAWKTISNSMCTLFTAGVNFNFDEFHKEFNDAQEMYTLPTYSFDNKKYWLDYHNDWTLRKGEPAQTKEVIVEKPVASASAPAVEMPAKRLSTSCQRVIAENFSGNNGSVTVQSSLADPKLYPVVCGHMVNNAALCPSSLYADMALTISDYIWKQMRPGTETPGYNVCNMEVPKP

>PKSI_1_11651

MSNVLLFGDQTAEQYPLLNKIVLRKENALVITFVERCAKALREETNALPRSQRNAVPDFLTVNDLKEAYHQKGVKVPMVESALVTIAQIGHYIGYFSEHSAEQPSATNTRALGLCTGLLAAAAVVASKTVEELVLVGVEFVRLSFRSGAAVDAARTALCQTGDDNAPWSTIVTGTTEAAAKEALAKFHEEKGIPQTSHAYISAVSVMAITVSGPPTTVKRFFEESPALSKNHRVPIPVYGPYHAEHLFGATEINKIASASILEGLKQHQPVSLVHSAATGKALVAENAAELAKLVLAEMLQHPVRWDHLLEEAVSQITSKKAPAKILAMGVSNVANSLVSALKAGGQTDVSTVDQSTWTENEPDTHGRTQNDKVAIVGMAGRFPNSADHEALWELLMKGLDVHRRIPKDRFDADTHVDPSGKGKNKSHTPFGCFIDEPGFFDPRFFNMSPREAAQTDPMGRLALVTAYEALEMSGYVPNRTPSTKLHRIGTFYGQTSDDWREINAAENVDTYFITGGVRAFAPGRINYYFKFSGPSYSVDTACSSSLAAIQLACTSLWAGDCDTACAGGLNVLTNPDIFSGLSKGQFLSKTGSCKTYDNNADGYCRGDAVGTVILKRYEDAIADKDNILGCILGAATNHSAEAVSITHPHAGAQEFLYKRVLANAGVDAHEISYVEMHGTGTQAGDGIEMTSVTNVFAPRHRQRRDDQPVYLGAIKANVGHAEAASGINSLAKVLLMMKHNKIPANVGIKGEMNKTFPADLKDRKVNISQKAVDWPRNGKEKRKVFLNNFSAAGGNTALLLEDGPAYEAPTATDPRGTVPVTVTARSISALKRNIANLQKYVSENPSTTLTSMSYTLTARRIQHNYRVAFPLDQINKFSDALQAQVKESYSPVPNAPTRVAFCFTGQGSQYTGLGQKLYNDLKSFRDDIDQLDHLARVQGLPSFLEIVQGADVQTLSPVKVQLGMACIQVALARMWAAWGITPAAVIGHSLGEYAALHVAGVISASDMVLLVGRRAELLVRDCTPHTHGMLAVKGGAEAIRNTLGSKMTEIACINGPEETVLCGSGDVVGAANETLAAKGFKATKLNVPFAFHSAQVDPILEQFKKIAASVTYNKPAVPVLSPLEGDIIREAGKINPEYLARHARETVNFWTALTAGQKEKVFDEKTAWLEVGAHPVCSGMVKASIGATTTAPSLRRGEDAWKTISNSMCTLFTAGVNFNFDEFHKEFNDAQEMYTLPTYSFDNKKYWLDYHNDWTLRKGEPAQTKEVIVEKPVASASAPAVEMPAKRLSTSCQRVIAENFTGNNGSVTVQSSLADPKLYPVVCGHMVNNAALCPSSLYADMALTISDYIWKQMRPGTETPGYNVCNMEVPKPLIAQIPQPAEGQHIQLEANADLDSGIVKLNFRSVKPDGQKLQDHAHCIVRLEDRAAWEDEWSRYNYMVRAQMELLQHKTLNGGAHKVQRGMAYKLFKALVNYDEKYRAMAEVVLASGQTEASAVLDFPTKPEDGDFYCPPYHIDGSCHISGFIVNASDLLDSEQNVYVSHGWGAMKFSRPLTAGMKLRNYVRMQPQPNNVSKGDVYIMEGDQIVAVCEGIKFQQIPRRVLNTFLPPNKGSGPASAAKPAAAPVAAARPAPAAAPIKTAPAPAKAAPAPAPAAPKAAPKPKKAAAPKKAAGGLTAKVMKILAKETEVDEGELVDEAQFENLGVDSLLSLTISAVFREELDMDISSTLFTDYPTVGDMKKYFAQFDNGSSTSSSTEEEDSDEDSIPPTDAATPMDDLSTPASSVGSSAPSDAGKPDSPTRETLEDVGDVSLAKHIVAQEMGVDIAEVTDDADLAEMGMDSLMSLTILGELREKTGIDLPSTFLTTNPTMKDIDNALGMRPKPKAAPKPAAPKAAAPSSSKKTDMNEVSARLSALNNNTDISRYPNATSVLLQGNPKQATKKIFFLPDGSGSATSYVSIPNLGPDVCAYGLNCPFMKNPEQWQCGIEISALVYLAEIKRRQPQGPYIIGGWSAGGVIAYSVAQALLAANEGVEKLLLLDSPCPVNLAPLPARLHNFFNEIGLLGTGDPAKTPKWLLPHFSAAIRSLSDYDPKPSLRPIPTYAIWCREGVAGNPGDPRPPPAEEEDPAPMTWLLEHRTNFKDNGWAQLCGDSMKFGVMGGHHFSMMKPPHADDLGNLIREGLDWQP

>PKSI_1_10958

MSNVLLFGDQTAEQYPLLNKIVLRKENALVITFVERCAKALREETNALPRSQRNAVPDFLTVNDLKEAYHQKGVKVPMVESALVTIAQIGHYIGYFSEHSAEQPSATNTRALGLCTGLLAAAAVVASKTVEELVLVGVEFVRLSFRSGAAVDAARTALCQTGDDNAPWSTIVTGTTEASAKEALAKFHEEKGIPQTSHAYISAVSVMAITVSGPPTTVKRFFEESPALSKNHRVPIPVYGPYHAEHLFGETEINKIADASILEGLKQHQPVSLVHSAATGKALVAENAAELAKLVLAEMLQHPVRWDHLLEEAVSQITSKKAPAKIWAMGVSNVANSLVSALKAGGQSDVSTVDQSSWTENEPDTHGRTQNDKVAIVGMAGRFPNSADHEALWDLLMKGLDVHRRIPKDRFDADTHVDPSGKGKNKSHTPFGCFIDEPGFFDPRFFNMSPREAAQTDPMGRLALVTAYEALEMSGYVPNRTPSTKLHRIGTFYGQTSDDWREINAAENVDTYFITGGVRAFAPGRINYYFKFSGPSYSVDTACSSSLAAIQLACTSLWAGDCDTACAGGLNVLTNPDIFSGLSKGQFLSKTGSCKTYDNNADGYCRGDAVGTVILKRYEDAIADKDNILGCILGAATNHSAEAVSITHPHAGAQEFLYKRVLANAGVDAHEISYVEMHGTGTQAGDGIEMTSVTNVFAPRHRQRRDDQPVYLGAIKANVGHAEAASGINSLAKVLMMMKHNKIPANVGIKGEMNKTFPADLKDRKVNISQKAVEWPRNGKEKRKVFLNNFSAAGGNTALLLEDGPAYEAPTASDPRGTVPVTVTARSISALKRNIANLQKYVSENPSTTLTSMSYTLTARRIQHNYRVAFPLDQINKFSDALQAQVKESYSPVPNAPTRVAFCFTGQGSQYTGLGQKLYNDLKSFRDDIDQLDHLARVQGLPSFLEIVQGADVQTLSPVKVQLGMACIQVALARMWAAWGITPAAVIGHSLGEYAALHVAGVISASDMVLLVGRRAELLVRDCTPHTHGMLAVKGGAEAIRNTLGNKMTEIACINGPEETVLCGSGDVVGAANETLAAKGFKATKLNVPFAFHSAQVDPILEQFKKIAASVTYNKPAVPVLSPLEGDIIREAGKINPEYLARHARETVNFWTALTAGQKEKVFDEKTAWLEVGAHPVCSGMVKASIGATTTAPSLRRGEDAWKTISNSMCTLFTAGVNFNFDEFHKEFNDAQEMYTLPTYSFDNKKYWLDYHNDWTLRKGEPAQTKEVIVEKPVASASAPAVEMPAKRLSTSCQRVIAENFSGNNGSVTVQSSLADPKLYPVVCGHMVNNAALCPSSLYADMALTISDYIWKQMRPGTETPGYNVCNMEVPKPLIAQIPQPAEGQHIQLEANADLDSGIVKLNFRSVKPDGQKLQDHAHCIVRLEDRAAWEDEWSRYNYMVQAQMELLQHKTLNGGAHKVQRGMAYKLFKALVNYDEKYRAMAEVVLASGQTEASAVLDFPTKPEDGDFYCPPYHIDGSCHISGFIVNASDLLDSEQNVYVSHGWGAMKFSRPLTAGMKLRNYVRMQPQPNNVSKGDVYIMEGDQIVAVCEGIKFQQIPRRVLNTFLPPNKGSGPASAAKPAAAPVAAARPAPAAAPMKTAPAPAKAAPAPAPAAPKAAPKPKKAAAPKKAAGGLTAKVMKILAKETEVDEGELVDEAQFENLGVDSLLSLTISAVFREELDMDISSTLFTDYPTVGDMKKYFAQFDNGSSTSSSAEEEDSDEDSIPPTDAATPMDDLSTPASSVPSSAPSDAGKPDSPTRETLEDVGDVSLAKHIVAQEMGVDIAEVTDDADLAEMGMDSLMSLTILGELREKTGIDLPSTFLTTNPTMKDIDNALGMRPKPKAAPKPAAPKAAAPSSSKKTDMNEVSARLSALNNNTDISRYPNATSVLLQGNPKQATKKIFFLPDGSGSATSYVSIPNLGPDVCAYGLNCPFMKNPEQWQCGIEISALVYLAEIKRRQPQGPYIIGGWSAGGVIAYSVAQALLAANEGVEKLLLLDSPCPVNLAPLPARLHNFFNEIGLLGTGDPAKTPKWLLPHFSAAIRSLSDYDPKPSLRPIPTYAIWCREGVAGNPSDPRPPPAEEEDPAPMTWLLEHRTNFKDNGWAQLCGDSMKFGVMGGHHFSMMKPPHADDLGNLIREGLDWQP

>PKSI_1_10907

MSNVLLFGDQTAEQYPLLNKIVLRKENALVITFVERCAKALREETNALPRSQRNAVPDFLTVNDLKEAYHQKGVKVPMVESALVTIAQIGHYIGYFSEHSAEQPSATNTRALGLCTGLLAAAAVVASKTVEELVLVGVDFVRLSFRSGAAVDAARTALCQTGDDNAPWSTIVTGTTEASAKEALAKFHEEKGIPQTSHAYISAVSVMAITVSGPPTTVKRFFEESPALSKNHRVPIPVYGPYHAEHLFGETEINKIASDSILEGLKQHQPVSLVHSAATGKALVAENAAELAKLVLAEMLQHSVRWDHLLEEAVSQITSKKAPAKIWAMGVSNVANSLVSALKAGGQSDVSTVDQSSWTENEPDTHGRTQNDKVAIVGMAGRFPNSADHEALWDLLMKGLDVHRRIPKDRFDADTHVDPSGKGKNKSHTPFGCFIDEPGFFDPRFFNMSPREAAQTDPMGRLALVTAYEALEMSGYVPNRTPSTKLHRIGTFYGQTSDDWREINAAENVDTYFITGGVRAFAPGRINYYFKFSGPSYSVDTACSSSLAAIQLACTSLWAGDCDTACAGGLNVLTNPDIFSGLSKGQFLSKTGSCKTYDNNADGYCRGDAVGTVILKRYEDAIADKDNILGCILGAATNHSAEAVSITHPHAGAQEFLYKRVLANAGVDAHEISYVEMHGTGTQAGDGIEMTSVTNVFAPRHRQRRDDQPVYLGAIKANVGHAEAASGINSLAKVLMMMKHNKIPANVGIKGEMNKTFPADLKDRKVNISQKAVEWPRNGKEKRKVFLNNFSAAGGNTALLLEDGPAYEAPTATDPRGTVPVTVTARSISALKRNIANLQKYVSENPSTTLTSMSYTLTARRIQHNYRVAFPLDQIDKFSDALQAQVKESYSPVPNVPTRVAFCFTGQGSQYTGLGQKLYNDLKSFRDDIDQLDHLARVQGLPSFLEIVQGADVQTLSPVKVQLGMACIQVALARMWAAWGITPAAVIGHSLGEYAALHVAGVISASDMVLLVGRRAELLVRDCTPHTHGMLAVKGGAEAIRNTLGNKMTEIACINGPEETVLCGSGDVVGAANETLAAKGFKATKLNVPFAFHSAQVDPILEQFKKIAASVTYNKPAVPVLSPLEGDIIREAGKINPEYLARHARETVNFWTALTAGQKEKVFDEKTAWLEVGAHPVCSGMVKASIGATTTAPSLRRGEDAWKTISNSMCTLFTAGVNFNFDEFHKEFNDAQEMYTLPTYSFDNKKYWLDYHNDWTLRKGEPAQTKEVIVEKPVASASAPAVEMPAKRLSTSCQRVIAENFSGNNGSVTVQSSLADPKLYPVVCGHMVNNAALCPSSLYADMALTISDYIWKQMRPGTETPGYNVCNME

>PKSI_1_10904

MSNVLLFGDQTAEQYPLLNKIVLRKENALVITFIERCAKALREETNALPRSQRNAVPDFLTVNDLKEAYHQKGVKVPMVESALVTIAQIGHYIGYFSEHSAEQPSATNTRALGLCTGLLAAAAVVASKTVEELVLVGVEFVRLSFRSGAAVDAARTALCQTGDDNSPWSTIVTGTTEAAAKEALAKFHEEKSIPQTSHAYISAVSVMAITVSGPPTTVKRFFEESSALSKNHRVPIPVYGPYHAEHLFGETEINKIASASILEGLKQHQPVSLVHSAATGKALVAENAAELAKLVLAEMLQHPVRWDHLLEEAVSQITSKKAPAKIWAMGVSNVANSLVSALKAGGQTDVSTIDQSTWTENEPDTHGRTQNDKVAIVGMAGRFPNSADHEALWELLMKGLDVHRRIPKDRFDADTHVDPSGKGKNKSHTPFGCFIDEPGFFDPRFFNMSPREAAQTDPMGRLALVTAYEALEMSGYVPNRTPSTKLHRIGTFYGQTSDDWREINAAENVDTYFITGGVRAFAPGRINYYFKFSGPSYSVDTACSSSLAAIQLACTSLWAGDCDTACAGGLNVLTNPDIFSGLSKGQFLSKTGSCKTYDNNADGYCRGDAVGTVILKRYEDAIADKDNILGCILGAATNHSAEAVSITHPHAGAQEFLYKRVLANAGVDAHEISYVEMHGTGTQAGDGIEMTSVTNVFAPRHRQRRDDQPVYLGAIKANVGHAEAASGINSLAKVLLMMKHNKIPANVGIKGEMNKTFPADLKDRKVNISQKAVDWPRNGKEKRKVFLNNFSAAGGNTALLLEDGPAYEAPTATDPRGTVPVTVTARSISALKRNIANLQKYVSENPSTTLTSMSYTLTARRIQHNYRVAFPLDQINKFSDALQAQVKESYSPVPNAPTRVAFCFTGQGSQYTGLGQKLYNDLKSFRDDIDQLDHLARVQGLPSFLEIVQGADVQTLSPVKVQLGMACIQVALARMWAAWGITPAAVIGHSLGEYAALHVAGVISASDMVLLVGRRAELLVRDCTPHTHGMLAVKGGAEAIRNTLGSKMTEIACINGPEETVLCGSGDVVGAANETLAAKGFKATKLNVPFAFHSAQVDPILEQFKKIAASVTYNKPAVPVLSPLEGDIIREAGKINPEYLARHARETVNFWTALTAGQKEKVFDEKTAWLEVGAHPVCSGMVKASIGATTTAPSLRRGEDAWKTISNSMCTLFTAGVNFNFDEFHKEFNDAQEMYTLPTYSFDNKKYWLDYHNDWTLRKGEPAQTKEVIVEKPVASASAPAVEMPAKRLSTSCQRVIAENFSSNNGSVTVQSSLADPKLYPVVCGHMVNNAALCPSSLYADMALTISDYIWKQMRPGTETPGYNVCNME

>PKSI_1_10820

MSNVLLFGDQTAEQYPLLNKIVLRKENALVITFVERCAKALREETNALPRSQRNAVPDFLTVNDLKEAYHQKGVKVPMVESALVTIAQIGHYIGYFSEHSAEQPSATNTRALGLCTGLLAAAAVVASKTVEELVLVGVEFVRLSFRSGAAVDAARTALCQTGDDNAPWSTIVTGTTEASAKEALAKFHEEKGIPQTSHAYISAVSVMAITVSGPPTTVKRFFEESPALSKNHRVPIPVYGPYHAEHLFGETEINKIADASILEGLKQHQPVSLVHSAATGKALVAENAAELAKLVLAEMLQHPVRWDHLLEEAVSQITSKKAPAKIWAMGVSNVANSLVSALKAGGQSDVSTVDQSSWTENEPDTHGRTQNDKVAIVGMAGRFPNSADHEALWDLLMKGLDVHRRIPKDRFDADTHVDPSGKGKNKSHTPFGCFIDEPGFFDPRFFNMSPREAAQTDPMGRLALVTAYEALEMSGYVPNRTPSTKLHRIGTFYGQTSDDWREINAAENVDTYFITGGVRAFAPGRINYYFKFSGPSYSVDTACSSSLAAIQLACTSLWAGDCDTACAGGLNVLTNPDIFSGLSKGQFLSKTGSCKTYDNNADGYCRGDAVGTVILKRYEDAIADKDNILGCILGAATNHSAEAVSITHPHAGAQEFLYKRVLANAGVDAHEISYVEMHGTGTQAGDGIEMTSVTNVFAPRHRQRRDDQPVYLGAIKANVGHAEAASGINSLAKVLMMMKHNKIPANVGIKGEMNKTFPADLKDRKVNISQKAVEWPRNGKEKRKVFLNNFSAAGGNTALLLEDGPAYEAPTASDPRGTVPVTVTARSISALKRNIANLQKYVSENPSTTLTSMSYTLTARRIQHNYRVAFPLDQINKFSDALQAQVKESYSPVPNAPTRVAFCFTGQGSQYTGLGQKLYNDLKSFRDDIDQLDHLARVQGLPSFLEIVQGADVQTLSPVKVQLGMACIQVALARMWAAWGITPAAVIGHSLGEYAALHVAGVISASDMVLLVGRRAELLVRDCTPHTHGMLAVKGGAEAIRNTLGNKMTEIACINGPEETVLCGSGDVVGAANETLAAKGFKATKLNVPFAFHSAQVDPILEQFKKIAASVTYNKPAVPVLSPLEGDIIREAGKINPEYLARHARETVNFWTALTAGQKEKVFDEKTAWLEVGAHPVCSGMVKASIGATTTAPSLRRGEDAWKTISNSMCTLFTAGVNFNFDEFHKEFNDAQEMYTLPTYSFDNKKYWLDYHNDWTLRKGEPAQTKEVIVEKPVASASAPAVEMPAKRLSTSCQRVIAENFSGNNGSVTVQSSLADPKLYPVVCGHMVNNAALCPSSLYADMALTISDYIWKQMRPGTETPGYNVCNMEVPKPLIAQIPQPAEGQHIQLEANADLDSGIVKLNFRSVKPDGQKLQDHAHCIVRLEDRAAWEDEWSRYNYMVQAQMELLQHKTLNGGAHKVQRGMAYKLFKALVNYDEKYRAMAEVVLASGQTEASAVLDFPTKPEDGDFYCPPYHIDGSCHISGFIVNASDLLDSEQNVYVSHGWGAMKFSRPLTAGMKLRNYVRMQPQPNNVSKGDVYIMEGDQIVAVCEGIKFQQIPRRVLNTFLPPNKGSGPASAAKPAAAPVAAARPAPAAAPMKTAPAPAKAAPAPAPAAPKAAPKPKKAAAPKKAAGGLTAKVMKILAKETEVDEGELVDEAQFENLGVDSLLSLTISAVFREELDMDISSTLFTDYPTVGDMKKYFAQFDNGSSTSSSAEEEDSDEDSIPPTDAATPMDDLSTPASSVPSSAPSDAGKPDSPTRETLEDVGDVSLAKHIVAQEMGVDIAEVTDDADLAEMGMDSLMSLTILGELREKTGIDLPSTFLTTNPTMKDIDNALGMRPKPKAAPKPAAPKAAAPSSSKKTDMNEVSARLSALNNNTDISRYPNATSVLLQGNPKQATKKIFFLPDGSGSATSYVSIPNLGPDVCAYGLNCPFMKNPEQWQCGIEISALVYLAEIKRRQPQGPYIIGGWSAGGVIAYSVAQALLAANEGVEKLLLLDSPCPVNLAPLPARLHNFFNEIGLLGTGDPAKTPKWLLPHFSAAIRSLSDYDPKPSLRPIPTYAIWCREGVAGNPSDPRPPPAEEEDPAPMTWLLEHRTNFKDNGWAQLCGDSMKFGVMGGHHFSMMKPPHADDLGNLIREGLDWQP

>PKSI_1_10816

MSNVLLFGDQTAEQYPLLNKIVLRKENALVITFVERCAKALREETNALPRSQRNAVPDFLTVNDLKEAYHQKGVKVPMVESALVTIAQIGHYIGYFSEHSAEQPSATNTRALGLCTGLLAAAAVVASKTVEELVLVGVDFVRLSFRSGAAVDAARTALCQTGDDNAPWSTIVTGTTEASAKEALAKFHEEKGIPQTSHAYISAVSVMAITVSGPPTTVKRFFEESPALSKNHRVPIPVYGPYHAEHLFGETEINKIASDSILEGLKQHQPVSLVHSAATGKALVAENAAELAKLVLAEMLQHSVRWDHLLEEAVSQITSKKAPAKIWAMGVSNVANSLVSALKAGGQSDVSTVDQSSWTENEPDTHGRTQNDKVAIVGMAGRFPNSADHEALWDLLMKGLDVHRRIPKDRFDADTHVDPSGKGKNKSHTPFGCFIDEPGFFDPRFFNMSPREAAQTDPMGRLALVTAYEALEMSGYVPNRTPSTKLHRIGTFYGQTSDDWREINAAENVDTYFITGGVRAFAPGRINYYFKFSGPSYSVDTACSSSLAAIQLACTSLWAGDCDTACAGGLNVLTNPDIFSGLSKGQFLSKTGSCKTYDNNADGYCRGDAVGTVILKRYEDAIADKDNILGCILGAATNHSAEAVSITHPHAGAQEFLYKRVLANAGVDAHEISYVEMHGTGTQAGDGIEMTSVTNVFAPRHRQRRDDQPVYLGAIKANVGHAEAASGINSLAKVLMMMKHNKIPANVGIKGEMNKTFPADLKDRKVNISQKAVEWPRNGKEKRKVFLNNFSAAGGNTALLLEDGPAYEAPTATDPRGTVPVTVTARSISALKRNIANLQKYVSENPSTTLTSMSYTLTARRIQHNYRVAFPLDQIDKFSDALQAQVKESYSPVPNVPTRVAFCFTGQGSQYTGLGQKLYNDLKSFRDDIDQLDHLARVQGLPSFLEIVQGADVQTLSPVKVQLGMACIQVALARMWAAWGITPAAVIGHSLGEYAALHVAGVISASDMVLLVGRRAELLVRDCTPHTHGMLAVKGGAEAIRNTLGNKMTEIACINGPEETVLCGSGDVVGAANETLAAKGFKATKLNVPFAFHSAQVDPILEQFKKIAASVTYNKPAVPVLSPLEGDIIREAGKINPEYLARHARETVNFWTALTAGQKEKVFDEKTAWLEVGAHPVCSGMVKASIGATTTAPSLRRGEDAWKTISNSMCTLFTAGVNFNFDEFHKEFNDAQEMYTLPTYSFDNKKYWLDYHNDWTLRKGEPAQTKEVIVEKPVASASAPAVEMPAKRLSTSCQRVIAENFSGNNGSVTVQSSLADPKLYPVVCGHMVNNAALCPSSLYADMALTISDYIWKQMRPGTETPGYNVCNME

>PKSI_1_6663

MSNVLLFGDQTAEQYPLLNKIVLRKENALVITFVERCAKALREETNALPRSQRNAVPDFLTVNDLKEAYHQKGVKVPMVESALVTIAQIGHYIGYFSEHSAEQPSATNTRALGLCTGLLAAAAVVASKTVEELVLVGVEFVRLSFRSGAAVDAARTALCQTGDDNAPWSTIVTGTTEASAKEALAKFHEEKGIPQTSHAYISAVSVMAITVSGPPTTVKRFFEESPALSKNHRVPIPVYGPYHAEHLFGETEINKIAGASILEGLKQHQPVSLVHSAATGKALVAENAAELAKLVLAEMLQHPVRWDHLLEEAVSQITSKKAPAKIWAMGVSNVANSLVSALKAGGQSNVSTVDQSSWTENEPDTHGRTQNDKVAIVGMAGRFPNSADHEALWDLLMKGLDVHRRIPKDRFDADTHVDPSGKGKNKSHTPFGCFIDEPGFFDPRFFNMSPREAAQTDPMGRLALVTAYEALEMSGYVPNRTPSTKLHRIGTFYGQTSDDWREINAAENVDTYFITGGVRAFAPGRINYYFKFSGPSYSVDTACSSSLAAIQLACTSLWAGDCDTACAGGLNVLTNPDIFSGLSKGQFLSKTGSCKTYDNNADGYCRGDAVGTVILKRYEDAIADKDNILGCILGAATNHSAEAVSITHPHAGAQEFLYKRVLANAGVDAHEISYVEMHGTGTQAGDGIEMTSVTNVFAPRHRQRRDDQPVYLGAIKANVGHAEAASGINSLAKVLMMMKHNKIPANVGIKGEMNKTFPADLKDRKVNISQKAVEWPRNGKEKRKVFLNNFSAAGGNTALLLEDGPAYEAPTASDPRGTVPVTVTARSISALKRNIANLQKYVSENPSTTLTSMSYTLTARRIQHNYRVAFPLDQINKFSDALQAQVKESYSPVPNAPTRVAFCFTGQGSQYTGLGQKLYNDLKSFRDDIDQLDHLARVQGLPSFLEIVQGADVQTLSPVKVQLGMACIQVALARMWAAWGITPAAVIGHSLGEYAALHVAGVISASDMVLVVGRRAELLVRDCTPHTHGMLAVKGGAEAIRNTLGNKMTEIACINGPEETVLCGSGDVVGAANETLAAKGFKATKLNVPFAFHSAQVDPILEQFKKIAASVTYNKPAVPVLSPLEGDIIREAGKINPEYLARHARETVNFWTALTAGQKEKVFDEKTAWLEVGAHPVCSGMVKASIGATTTAPSLRRGEDAWKTISNSMCTLFTAGVNFNFDEFHKEFNDAQEMYTLPTYSFDNKKYWLDYHNDWTLRKGEPAQTKEVIVEKPVASASAPAVEIPAKRLSTSCQRVISENFSGNNGSVTVQSSLADPKLYPVVCGHMVNNAALCPSSLYADMALTISDYIWKQMRPGTETPGYNVCNMEVPKPLIAQIPQPAEGQHIQLEANADLDSGIVKLNFRSVKPDGQKLQDHAHCIVRLEDRAAWEDEWSRYNYMVQAQMELLQHKTLNGGAHKVQRGMAYKLFKALVNYDEKYRAMAEVVLASGQTEASAVLDFPTKPEDGDFYCPPYHIDGSCHISGFIVNASDLLDSEQNVYVSHGWGAMKFSRPLTAGMKLRNYVRMQPQPNNVSKGDVYIMEGDQIVAVCEGIKFQQIPRRVLNTFLPPNKGSGPASAAKPAAAPVAAARPAPAAAPIKTAPAPAKAAPAPAPAATKAAPKPKKAAAPKKAAGGLTAKVMKILAKETEVDEGELVDEAQFENLGVDSLLSLTISAVFREELDMDISSTLFTDYPTVGDMKKYFAQFDNGSSTSSSAEEEDSDEDSIPPTDAATPMDDLSTPASSVPSSAPSDAGKPDSPTRETLEDVGDVSLAKHIVAQEMGVDIAEVTDDADLAEMGMDSLMSLTILGELREKTGIDLPSTFLTTNPTMKDIDNALGMRPKPKAAPKPAAPKAAAPSSSKKTDMNEVSARLSALNNNTDISRYPNATSVLLQGNPKQATKKIFFLPDGSGSATSYVSIPNLGPDVCAYGLNCPFMKNPEQWQCGIEISALVYLAEIKRRQPQGPYIIGGWSAGGVIAYSVAQALLAANEGVEKLLLLDSPCPVNLAPLPARLHNFFNEIGLLGTGDPAKTPKWLLPHFSAAIRSLSDYDPKPSLRPIPTYAIWCREGVAGNPGDPRPPPAEEEDPAPMTWLLEHRTNFKDNGWAQLCGDSMKFGVMGGHHFSMMKPPHADDLGNLIREGLDWQP

>PKSI_1_8170

MSNVLLFGDQTAEQYPLLNKIVLRKENALVITFVERCAKALREETNALPRSQRNAVPDFLTVNDLKEAYHQKGVKVPMVESALVTIAQIGHYIGYFSEHSAEQPSATNTRALGLCTGLLAAAAVVASKTVEELVLVGVEFVRLSFRSGAAVDAARTALCQTGDDNAPWSTIVTGTTEASAKEALAKFHEEKGIPQTNHAYISAVSVMAITVSGPPTTVKRFFEESPALSKNHRVPIPVYGPYHAEHLFGETEINKIASASILEGLKQHQPVSLVHSAATGKALVAENAAELAKLVLAEMLQLPVRWDHLLEEAVSQITSKKAPAKIWAMGVSNVANSLVSALKAGGQTDVSTVDQGTWTENEPDTHGRTQNDKVAIVGMAGRFPNSADHEALWELLMKGLDVHRRIPKDRFDADTHVDPSGKGKNKSHTPFGCFIDEPGFFDPRFFNMSPREAAQTDPMGRLALVTAYEALEMSGYVPNRTPSTKLHRIGTFYGQTSDDWREINAAENVDTYFITGGVRAFAPGRINYYFKFSGPSYSVDTACSSSLAAIQLACTSLWAGDCDTACAGGLNVLTNPDIFSGLSKGQFLSKTGSCKTYDNNADGYCRGDAVGTVILKRYEDAIADKDNILGCILGAATNHSAEAVSITHPHAGAQEFLYKRVLANAGVDAHEISYVEMHGTGTQAGDGIEMTSVTNVFAPRHRQRRDDQPVYLGAIKANVGHAEAASGINSLAKVLLMMKHNKIPANVGIKGEMNKTFPADLKDRKVNISQKAVDWPRNGKEKRKVFLNNFSAAGGNTALLLEDGPAYEAPTATDPRGTVPVTVTARSISALKRNIANLQKYVSENPSTTLTSMSYTLTARRIQHNYRVAFPLDQINKFSDALQAQVKESYSPVPNAPTRVAFCFTGQGSQYTGLGQKLYNDLKSFRDDIDQLDHLARVQGLPSFLEIVQGADVQTLSPVKVQLGMACIQVALARMWAAWGITPAAVIGHSLGEYAALHVAGVISASDMVLLVGRRAELLVRDCTPHTHGMLAVKGGAEAIRNTLGNKMTEIACINGPEETVLCGSGDVVGAANETLAAKGFKATKLNVPFAFHSAQVDPILEQFKKIAASVTYNKPAVPVLSPLEGDIIREAGKINPEYLARHARETVNFWTALTAGQKEKVFDEKTAWLEVGAHPVCSGMVKASIGATTTAPSLRRGEDAWKTISNSMCTLFTAGVNFNFDEFHKEFNDAQEMYTLPTYSFDNKKYWLDYHNDWTLRKGEPAQTKEVIVEKPVASASAPAVEMPAKRLSTSCQRVIAENFSGNNGSVTVQSSLADPKLYPVVCGHMVNNAALCPSSLYADMALTISDYIWKQMRPGTETPGYNVCNMEVPKPLIAQIPQPAEGQHIQLEANADLDSGIVKLNFRSVKPDGQKLQDHAHCIVRLEDKAAWEDEWSRYNYMVQAQMELLQHKTLNGGAHKVQRGMAYKLFKALVNYDEKYRAMAEVVLASGQTEASAMLDFPTKPEDGDFYCPPYHIDGSCHISGFIVNASDLLDSEQNVYVSHGWGAMKFSRPLTAGMKLRNYVRMQPQPNNVSKGDVYIMEGDQIVAVCEGIKFQQIPRRVLNTFLPPNKGSGPASAAKPAAAPVAAARPAPAAAPIKTAPAPAKAAPAPAPAAPKAAPKPKKAAAPKKPAGGLTAKVMKILAKETEVDEGELVDEAQFENLGVDSLLSLTISAVFREELDMDISSTLFTDYPTVGDMKKYFAQFDNGSSTSSSAEEEDSDEDSIPPTDAATPMDDLSTPASSVPSSAPSDAGKPDSPTRETLEDVGDVSLAKHIVAQEMGVDIAEVTDDADLAEMGMDSLMSLTILGELREKTGIDLPSTFLTTNPTMKDIDNALGMRPKPKAAPKPAAPKAAAPS

>PKSI_1_3846

MSNVLLFGDQTAEQYPLLNKIVLRKENALVITFVERCAKALREETNALPRSQRNAVPDFLTVNDLKEAYHQKGVKVPMVESALVTIAQIGHYIGYFSEHSAEQPSATNTRALGLCTGLLAAAAVVASKTVEELVLVGVEFVRLSFRSGAAVDAARTALCQTGDDNAPWSTIVTGTTEASAKEALAKFHEEKGIPQTSHAYISAVSVMAITVSGPPTTVKRFFEESPALSKNHRVPIPVYGPYHAEHLFGETEINKIADASILEGLKQHQPVSLVHSAATGKALVAENAAELAKLVLAEMLQHPVRWDHLLEEAVSQITSKKAPAKIWAMGVSNVANSLVSALKAGGQSDVSTVDQSSWTENEPDTHGRTQNDKVAIVGMAGRFPNSADHEALWDLLMKGLDVHRRIPKDRFDADTHVDPSGKGKNKSHTPFGCFIDEPGFFDPRFFNMSPREAAQTDPMGRLALVTAYEALEMSGYVPNRTPSTKLHRIGTFYGQTSDDWREINAAENVDTYFITGGVRAFAPGRINYYFKFSGPSYSVDTACSSSLAAIQLACTSLWAGDCDTACAGGLNVLTNPDIFSGLSKGQFLSKTGSCKTYDNNADGYCRGDAVGTVILKRYEDAIADKDNILGCILGAATNHSAEAVSITHPHAGAQEFLYKRVLANAGVDAHEISYVEMHGTGTQAGDGIEMTSVTNVFAPRHRQRRDDQPVYLGAIKANVGHAEAASGINSLAKVLMMMKHNKIPANVGIKGEMNKTFPADLKDRKVNISQKAVEWPRNGKEKRKVFLNNFSAAGGNTALLLEDGPAYEAPTASDPRGTVPVTVTARSISALKRNIANLQKYVSENPSTTLTSMSYTLTARRIQHNYRVAFPLDQINKFSDALQAQVKESYSPVPNAPTRVAFCFTGQGSQYTGLGQKLYNDLKSFRDDIDQLDHLARVQGLPSFLEIVQGADVQTLSPVKVQLGMACIQVALARMWAAWGITPAAVIGHSLGEYAALHVAGVISASDMVLLVGRRAELLVRDCTPHTHGMLAVKGGAEAIRNTLGNKMTEIACINGPEETVLCGSGDVVGAANETLAAKGFKATKLNVPFAFHSAQVDPILEQFKKIAASVTYNKPAVPVLSPLEGDIIREAGKINPEYLARHARETVNFWTALTAGQKEKVFDEKTAWLEVGAHPVCSGMVKASIGATTTAPSLRRGEDAWKTISNSMCTLFTAGVNFNFDEFHKEFNDAQEMYTLPTYSFDNKKYWLDYHNDWTLRKGEPAQTKEVIVEKPVASASAPAVEMPAKRLSTSCQRVIAENFSGNNGSVTVQSSLADPKLYPVVCGHMVNNAALCPSSLYADMALTISDYIWKQMRPGTETPGYNVCNMEVPKPLIAQIPQPAEGQHIQLEANADLDSGIVKLNFRSVKPDGQKLQDHAHCIVRLEDRAAWEDEWSRYNYMVQAQMELLQHKTLNGGAHKVQRGMAYKLFKALVNYDEKYRAMAEVVLASGQTEASAVLDFPTKPEDGDFYCPPYHIDGSCHISGFIVNASDLLDSEQNVYVSHGWGAMKFSRPLTAGMKLRNYVRMQPQPNNVSKGDVYIMEGDQIVAVCEGIKFQQIPRRVLNTFLPPNKGSGPASAAKPAAAPVAAARPAPAAAPMKTAPAPAKAAPAPAPAAPKAAPKPKKAAAPKKAAGGLTAKVMKILAKETEVDEGELVDEAQFENLGVDSLLSLTISAVFREELDMDISSTLFTDYPTVGDMKKYFAQFDNGSSTSSSAEEEDSDEDSIPPTDAATPMDDLSTPASSVPSSAPSDAGKPDSPTRETLEDVGDVSLAKHIVAQEMGVDIAEVTDDADLAEMGMDSLMSLTILGELREKTGIDLPSTFLTTNPTMKDIDNALGMRPKPKAAPKPAAPKAAAPSSSKKTDMNEVSARLSALNNNTDISRYPNATSVLLQGNPKQATKKIFFLPDGSGSATSYVSIPNLGPDVCAYGLNCPFMKNPEQWQCGIEISALVYLAEIKRRQPQGPYIIGGWSAGGVIAYSVAQALLAANEGVEKLLLLDSPCPVNLAPLPARLHNFFNEIGLLGTGDPAKTPKWLLPHFSAAIRSLSDYDPKPSLRPIPTYAIWCREGVAGNPSDPRPPPAEEEDPAPMTWLLEHRTNFKDNGWAQLCGDSMKFGVMGGHHFSMMKPPHADDLGNLIREGLDWQP

>PKSI_1_3845

MSNVLLFGDQTAEQYQLLNKIVLRKENALVITFVERCAKALREETNALPRSQRNAVPDFLTVNDLKEAYHQKGVKVPMVESALVTIAQIGHYIGYFSEHSAEQPSATNTRALGLCTGLLAAAAVVASKTVEELVLVGVEFVRLSFRSGAAVDAARTALCQTGDDNAPWSTIVTGTTEASAKEALAKFHEEKGIPQTSHAYISAVSVMAITVSGPPTTVKRFFEESPALSKNHRVPIPVYGPYHAEHLFGETEINKIASASILEGLKQHQPVSLVHSAATGKALVAENAAELAKLVLAEMLQLPVRWDHLLEEAVSQITSKKAPAKIWAMGVSNVANSLVSALKAGGQTDVSTVDQSTWTENEPDTHGRTQNDKVAIVGMAGRFPNSADHEALWELLMKGLDVHRRIPKDRFDADTHVDPSGKGKNKSHTPFGCFIDEPGFFDPRFFNMSPREAAQTDPMGRLALVTAYEALEMSGYVPNRTPSTKLHRIGTFYGQTSDDWREINAAENVDTYFITGGVRAFAPGRINYYFKFSGPSYSVDTACSSSLAAIQLACTSLWAGDCDTACAGGLNVLTNPDIFSGLSKGQFLSKTGSCKTYDNNADGYCRGDAVGTVILKRYEDAIADKDNILGCILGAATNHSAEAVSITHPHAGAQEFLYKRVLANAGVDAHEISYVEMHGTGTQAGDGIEMTSVTNVFAPRHRQRRDDQPVYLGAIKANVGHAEAASGINSLAKVLLMMKHNKIPANVGIKGEMNKTFPADLKDRKVNISQKAVDWPRNGKEKRKVFLNNFSAAGGNTALLLEDGPAYEAPTATDPRGTVPVTVTARSISALKRNIANLQKHVSENPSTTLTSMSYTLTARRIQHNYRVAFPLDQINKFSDALQAQVKESYSPVPNAPTRVAFCFTGQGSQYTGLGQKLYNDLKSFRDDIDQLDHLARVQGLPSFLEIVQGADVQTLSPVKVQLGMACIQVALARMWAAWGITPAAVIGHSLGEYAALHVAGVISASDMVLLVGRRAELLVRDCTPHTHGMLAVKGGAEAIRNTLGNKMTEIACINGPEETVLCGSGDVVGAANETLAAKGFKATKLNVPFAFHSAQVDPILEQFKKIAASVTYNKPAVPVLSPLEGDIIREAGKINPEYLARHARETVNFWTALTAGQKEKVFDEKTAWLEVGAHPVCSGMVKASIGATTTAPSLRRGEDAWKTISNSMCTLFTAGVNFNFDEFHKEFNDAQEMYTLPTYSFDNKKYWLDYHNDWTLRKGEPAQTKEVIVEKPVASASAPAVEMPAKRLSTSCQRVIAENFSGNNGNVTVQSSLADPKLYPVVCGHMVNNAALCPSSLYADMALTISDYIWKQMRPGTETPGYNVCNMEVPKPLIAQIPQPAEGQHIQLEANADLDSGIVKLNFRSVKPDGQKLQDHAHCIVRLEDKAAWEDEWSRYNYMVQAQMELLQHKTLNGGAHKVQRGMAYKLFKALVNYDEKYRAMAEVVLASGQTEASAMLDFPTKPEDGDFYCPPYHIDGSCHISGFIVNASDLLDSEQNVYVSHGWGAMKFSRPLTAGMKLRNYVRMQPQPNNVSKGDVYIMEGDQIVAVCEGIKFQQIPRRVLNTFLPPNKGSGPASAAKPAAAPVAAARPAPAAAPIKTAPAPAKAAPAPAPAAPKAAPKPKKAAAPKKPAGGLTAKVMKILAKETEVDEGELVDEAQFENLGVDSLLSLTISAVFREELDMDISSTLFTDYPTVGDMKKYFAQFDNGSSTSSSAEEEDSDEDSIPPTDAATPMDDLSTPASSVPSSAPSDAGKPDSPTRETLEDVGDVSLAKHIVAQEMGVDIAEVTDDADLAEMGMDSLMSLTILGELREKTGIDLPSTFLTTNPTMKDIDNALGMRPKPKAAPKPAAPKAAAPSSSKKTDMNEVSARLSALNNNTDISRYPNATSVLLQGNPKQATKKIFFLPDGSGSATSYVSIPNLGPDVCAYGLNCPFMKNPEQWQCGIEISALVYLAEIKRRQPQGPYIIGGWSAGGVIAYSVAQALLAANEGVEKLLLLDSPCPVNLAPLPARLHNFFNQIGLLGTGDPAKTPKWLLPHFSAAIRSLSDYDPKPSLRPIPTYAIWCREGVAGNPGDPRPPPAEEEDPAPMTWLLEHRTNFKDNGWAQLCGDSMKFGVMGGHHFSMMKPPHADDLGNLIREGLDWQP

>PKSI_2_3845

MPDNVSFMDESQDLRHIRETSRTSHTTSILNVEGDYRNNDMPREPGECNRSTNGTVDHERMTPSADGGIPIAICGIGLRLPGGIRNDRDLYDSLYNKKDARGVIPEDRFSIDSFHSAHGKTGTIITKHGYFLQDIDLTKFDVNMFNMTPAEVERLDPHQRILLETVRETLESAGEASFRGKKVGTYVGNFTDDWLDLQNVDTVDFATYQLHGKMDFSLANRISYEYDLRGPSMTIKTACSSSALAIHEAVYSIRNGECDAAIVSGSNLNLAPRLWVGMSSQGAISPDGSSKTFDESANGYARGDGIAALFIKRLDDAVRDGNPVRAVIRSTASNADGRTPGMTMPSTEAQEALIRRAYDAANLPLSETAMVECHGTGTAVGDPMEANAVARCFGDQGMLIGSVKPNLGHSEGASAITSVVKAVLSLENRTILPNIKFHRPNPAIPWSEAKLTVPVEPLAWPKDRQERISVNSFGIGGSNVHVVLDSAASMGFRPRSLAPSKDDRPGRLLLFSGGHRASVEQSSSQHQDYVTKYPNRLSDVAYTLAKRPPVKCPGLTRSTFVFTGQGAQWLHMGKELLHESPVFAKSIGRMDSVIHSLKHAPQWTLEGIINDPENPSALTNAEISQPLCTAVQIGLVDLLKSWAIYPHAVLGHSSGEIGAAYASGVVDRAEAILLAFYRGYVCRFAQKAGGMAAVGLEKSQVIKYLQPGVCVACENSGSSVTLSGDLETLEEVLQSIRAENLNAFARKLQVGIAYHSDHMKALGGLYHQYITEHLDPKDPQVPFFSSVSGRALHSKDDFGATYWQDNLENPVLFHTAVLKSLEHTGDKQVHLEVGPHGALNGPLRQIYAETGSKARYVALQKRGANCFDTFLEGIGQLYCNGVPLQYPESADDRTLIDLPPYPWHYDHSYWSETRVMKNWRFRRQLPHDLLGLRTLDCSDAEPMWRNILRITDLPWLRDHCVGKDVVFPASGYMCMAGEAVFQETGCRDYTLREVDISTAMVLSSDHSTELLTTMKKRRLNAFLDSRWYEFLIMSYDGASWTQHCSGLVTNGPSTSHPKALLQTYDRPVSTNRWYTAMSKIGLNYGPRFTGLQNITTHVQEKKASMTIMDKQEDYESPYALHPSTLDLILQSWTVASVRGEYRRFTQLFLPTFVDEFYIGNSASKLIHLNTTAIGPDGSARGEAIGKDNDGQISFNLKGFKGSKLDNVGVDQPQEMQTIMKQQWKRDFDFADTAQLMRPAFDSTSELSLLERMFVLAAIEVHVRTSGMEGKLPHHQRYKLWIDAQIRRFGEPGYPMVEDSMELLRLDSRERQRQLCRLLEHSRKTTAHPVAEAIWRALDRIEDVFDGRIEYLDLLFNDGLMPKFYDWSNSLSDVSRLFRLLSHKKPQLKILEVGAGTGGSTARLLQYLQSDFGERQYHSYTYTDVSSGFFVQAQERFKDYEGMKYRVLDISQDPFEQGFGADEFDLICASNVLHATPRLTETLRNCRKMLRPDGILFLQELCPRQQFMGFIMGLFEGWWLGAEDGRADTPLLLPPAWDRRLRDVGFEGVEAFSFDNNPPYFMAANMTARPTVTAKSKGSITLLTFNDFLDDVAGALKQAIRAAGFEIDHCVWGEHVPLDQSLISLVDLERDKPLLQDIGDDDLRIFLDLIDAVLQTTVIWLNKPAQVSSADPNAAQMLGLARTLRAELAMHFATVEMADPTLNGMSAVVHLMCMLQRGSALPENSLDQDMEYVWAHDAMHVSRFHFQPVDEALMEMSPKFDVKTLVPLQRGMLSSLKWIGTRMLPLAEAEVQIRMSAVGMNFHDMMIAMNMFDSPLTLGSGYNSIGMEGVGYVTRKSPEVDHVQVGDRVIVIGSNSSGFATDVHRPADYCIKCPSSLTDVEAAGMSFAYMTVLWSFLDKGGLRKGQSVLIHSAAGGVGIAALHVSRWLGLEAFVTVGNEEKVRFIMNNFGLPRNRIFNSHSATFLDDVMDATAGRGVDAALSAAAGELLHNTWSCIAPGGVMLEIGKRDLINRGRLSLAPFEENRSYVGIDFSRLTIVNKPAVVRLLRQTMRLVEQGHIHPIHPTTCFDAEHAEDAFRFMQTGQHIGRIVVKIPRDTSTIPLASRPPAPEFEGQKTYLLVGGMGGLGRSVASWMVSAGARNLIFMSRSAGKSEQDQNFARELELSGCKVHCCAIDITDTDAVREALERTQASVAGVLQMAMVLRDVGIMNMDKANWDAAVAPKVQGTWNLHHALPNVDFFVMFGSNSGTLGSYGQANYAAANAFLDSFVQYRQNLGQAASVIDIGAVGDVGYVAETQVAAENMESMAGRLISEQDFLNCLQLAIARSTPTERRNKGPSTEADGYIDLKQVILLNSSILSMADPSNQIFWRKDPRMGIYRNVQRTSAESPTTDSNSLRRLVATLKADSSIADQAELAQTLARELSKQVATVLMLGEDEIDVDRSLTDVGMDSLVAIEMRNWWKQNLGVDVSVLELKDGRSILRLGELAATRLKERYSRDS

>PKSI_1_2685

MSNVLLFGDQTAEQYPLLNKIVLRKENALVITFVERCAKALREETNALPRSQRNAVPDFLTVNDLKEAYHQKGVKVPMVESALVTIAQIGHYIGYFSEHSAEQPSATNTRALGLCTGLLAAAAVVASKTVEELVLVGVEFVRLSFRSGAAVDAARTALCQTGDDNAPWSTIVTGTTEASAKEALAKFHEEKGIPQTSHAYISAVSVMAITVSGPPTTVKRFFEESPALSKNHRVPIPVYGPYHAEHLFGETEINKIASASILEGLKQHQPVSLVHSAATGKALVAENAAELAKLVLAEMLQHPVRWDHLLEEAVSQITSKKAPAKIWAMGVSNVANSLVSALKAGGQSDVSTVDQSSWTENEPDTHGRTQNDKVAIVGMAGRFPNSADHEALWDLLMKGLDVHRRIPKDRFDADTHVDPSGKGKNKSHTPFGCFIDEPGFFDPRFFNMSPREAAQTDPMGRLALVTAYEALEMSGYVPNRTPSTKLHRIGTFYGQTSDDWREINAAENVDTYFITGGVRAFAPGRINYYFKFSGPSYSVDTACSSSLAAIQLACTSLWAGDCDTACAGGLNVLTNPDIFSGLSKGQFLSKTGSCKTYDNNADGYCRGDAVGTVILKRYEDAIADKDNILGCILGAATNHSAEAVSITHPHAGAQEFLYKRVLANAGVDAHEISYVEMHGTGTQAGDGIEMTSVTNVFAPRHRQRRDDQPVYLGAIKANVGHAEAASGINSLAKVLMMMKHNKIPANVGIKGEMNKTFPADLKDRKVNISQKAVEWPRNGKEKRKVFLNNFSAAGGNTALLLEDGPAYEAPTASDPRGTVPVTVTARSISALKRNIANLQKYVSENPSTTLTSMSYTLTARRIQHNYRVAFPLDQINKFSDALQAQVKESYSPVPNAPTRVAFCFTGQGSQYTGLGQKLYNDLKSFRDDIDQLDHLARVQGLPSFLEIVQGADVQTLSPVKVQLGMACIQVALARMWAAWGITPAAVIGHSLGEYAALHVAGVISASDMVLLVGRRAELLVRDCTPHTHGMLAVKGGAEAIRNTLGNKMTEIACINGPEETVLCGSGDVVGAANETLAAKGFKATKLNVPFAFHSAQVDPILEQFKKIAASVTYNKPAVPVLSPLEGDIIREAGKINPEYLARHARETVNFWTALTAGQKEKVFDEKTAWLEVGAHPVCSGMVKASIGATTTAPSLRRGEDAWKTISNSMCTLFTAGVNFNFDEFHKEFNDAQEMYTLPTYSFDNKKYWLDYHNDWTLRKGEPAQTKEVIVEKPVASASAPAVEIPAKRLSTSCQRVISENFSGNNGSVTVQSSLADPKLYPVVCGHMVNNAALCPSSLYADMALTISDYIWKQMRPGTETPGYNVCNMEVPKPLIAQIPQPAEGQHIQLEANADLDSGIVKLNFRSVKPDGQKLQDHAHCIVRLEDRAAWEDEWSRYNYMVQAQMELLQHKTLNGGAHKVQRGMAYKLFKALVNYDEKYRAMAEVVLASGQTEASAVLDFPTKPEDGDFYCPPYHIDGSCHISGFIVNASDLLDSEQNVYVSHGWGAMKFSRPLTAGMKLRNYVRMQPQPNNVSKGDVYIMEGDQIVAVCEGIKFQQIPRRVLNTFLPPNKGSGPASAAKPAAAPVAAARPAPAAAPIKTAPAPAKAAPAPAPAAPKAAPKPKKAAAPKKAAGGLTAKVMKILAKETEVDEGELVDEAQFENLGVDSLLSLTISAVFREELDMDISSTLFTDYPTVGDMKKYFAQFDNGSSTSSSAEEEDSDEDSIPPTDAATPMDDLSTPASSVPSSAPSDAGKPDSPTRETLEDVGDVSLAKHIVAQEMGVDIAEVTDDADLAEMGMDSLMSLTILGELREKTGIDLPSTFLTTNPTMKDIDNALGMRPKPKAAPKPAAPKAAAPSSSKKTDMNEVSARLSALNNNTDISRYPNATSVLLQGNPKQATKKIFFLPDGSGSATSYVSIPNLGPDVCAYGLNCPFMKNPEQWQCGIEISALVYLAEIKRRQPQGPYIIGGWSAGGVIAYSVAQALLAANEGVEKLLLLDSPCPVNLAPLPARLHNFFNEIGLLGTGDPAKTPKWLLPHFSAAIRSLSDYDPKPSLRPIPTYAIWCREGVAGNPGDPRPPPAEEEDPAPMTWLLEHRTDFKDNGWAQLCGDSMKFGVMGGHHFSMMKPPHADDLGNLIREGLDWQP

>PKSI_1_2783

MSNVLLFGDQTAEQYPLLNKIVLRKENALVITFVERCAKALREETNALPRSQRNAVPDFLTVNDLKEAYHQKGVKVPMVESALVTIAQIGHYIGYFSEHSAEQPSATNTRALGLCTGLLAAAAVVASKTVEELVLVGVEFVRLSFRSGAAVDAARTALCQTGDDNAPWSTIVTGTTEASAKEALAKFHEEKGIPQTSHAYISAVSVMAITVSGPPTTVKRFFEESPALSKNHRVPIPVYGPYHAEHLFGETEINKIADASILEGLKQHQPVSLVHSAATGKALVAENAAELAKLVLAEMLQHPVRWDHLLEEAVSQITSKKAPAKIWAMGVSNVANSLVSALKAGGQSDVSTVDQSSWTENEPDTHGRTQNDKVAIVGMAGRFPNSADHEALWDLLMKGLDVHRRIPKDRFDADTHVDPSGKGKNKSHTPFGCFIDEPGFFDPRFFNMSPREAAQTDPMGRLALVTAYEALEMSGYVPNRTPSTKLHRIGTFYGQTSDDWREINAAENVDTYFITGGVRAFAPGRINYYFKFSGPSYSVDTACSSSLAAIQLACTSLWAGDCDTACAGGLNVLTNPDIFSGLSKGQFLSKTGSCKTYDNNADGYCRGDAVGTVILKRYEDAIADKDNILGCILGAATNHSAEAVSITHPHAGAQEFLYKRVLANAGVDAHEISYVEMHGTGTQAGDGIEMTSVTNVFAPRHRQRRDDQPVYLGAIKANVGHAEAASGINSLAKVLMMMKHNKIPANVGIKGEMNKTFPADLKDRKVNISQKAVEWPRNGKEKRKVFLNNFSAAGGNTALLLEDGPAYEAPTASDPRGTVPVTVTARSISALKRNIANLQKYVSENPSTTLTSMSYTLTARRIQHNYRVAFPLDQINKFSDALQAQVKESYSPVPNAPTRVAFCFTGQGSQYTGLGQKLYNDLKSFRDDIDQLDHLARVQGLPSFLEIVQGADVQTLSPVKVQLGMACIQVALARMWAAWGITPAAVIGHSLGEYAALHVAGVISASDMVLLVGRRAELLVRDCTPHTHGMLAVKGGAEAIRNTLGNKMTEIACINGPEETVLCGSGDVVGAANETLAAKGFKATKLNVPFAFHSAQVDPILEQFKKIAASVTYNKPAVPVLSPLEGDIIREAGKINPEYLARHARETVNFWTALTAGQKEKVFDEKTAWLEVGAHPVCSGMVKASIGATTTAPSLRRGEDAWKTISNSMCTLFTAGVNFNFDEFHKEFNDAQEMYTLPTYSFDNKKYWLDYHNDWTLRKGEPAQTKEVIVEKPVASASAPAVEMPAKRLSTSCQRVIAENFSGNNGSVTVQSSLADPKLYPVVCGHMVNNAALCPSSLYADMALTISDYIWKQMRPGTETPGYNVCNMEVPKPLIAQIPQPAEGQHIQLEANADLDSGIVKLNFRSVKPDGQKLQDHAHCIVRLEDRAAWEDEWSRYNYMVQAQMELLQHKTLNGGAHKVQRGMAYKLFKALVNYDEKYRAMAEVVLASGQTEASAVLDFPTKPEDGDFYCPPYHIDGSCHISGFIVNASDLLDSEQNVYVSHGWGAMKFSRPLTAGMKLRNYVRMQPQPNNVSKGDVYIMEGDQIVAVCEGIKFQQIPRRVLNTFLPPNKGSGPASAAKPAAAPVAAARPAPAAAPMKTAPAPAKAAPAPAPAAPKAAPKPKKAAAPKKAAGGLTAKVMKILAKETEVDEGELVDEAQFENLGVDSLLSLTISAVFREELDMDISSTLFTDYPTVGDMKKYFAQFDNGSSTSSSAEEEDSDEDSIPPTDAATPMDDLSTPASSVPSSAPSDAGKPDSPTRETLEDVGDVSLAKHIVAQEMGVDIAEVTDDADLAEMGMDSLMSLTILGELREKTGIDLPSTFLTTNPTMKDIDNALGMRPKPKAAPKPAAPKAAAPSSSKKTDMNEVSARLSALNNNTDISRYPNATSVLLQGNPKQATKKIFFLPDGSGSATSYVSIPNLGPDVCAYGLNCPFMKNPEQWQCGIEISALVYLAEIKRRQPQGPYIIGGWSAGGVIAYSVAQALLAANEGVEKLLLLDSPCPVNLAPLPARLHNFFNEIGLLGTGDPAKTPKWLLPHFSAAIRSLSDYDPKPSLRPIPTYAIWCREGVAGNPSDPRPPPAEEEDPAPMTWLLEHRTNFKDNGWAQLCGDSMKFGVMGGHHFSMMKPPHADDLGNLIREGLDWQP

>PKSI_1_2683

MSNVLLFGDQTAEQYPLLNKIVLRKENALVITFVERCAKALREETNALPRSQRNAVPDFLTVNDLKEAYHQKGVKVPMVESALVTIAQIGHYIGYFSEHSAEQPSATNTRALGLCTGLLAAAAVVASKTVEELVLVGVEFVRLSFRSGAAVDAARTALCQTGDDNAPWSTIVTGTTEASAKEALAKFHEEKGIPQTSHAYISAVSVMAITVSGPPTTVKRFFEESPALSKNHRVPIPVYGPYHAEHLFGETEINKIASASILEGLKQHQPVSLVHSAATGKALVAENAAELAKLVLAEMLQHPVRWDHLLEEAVSQITSKKAPAKIWAMGVSNVANSLVSALKAGGQSDVSTVDQSSWTENEPDTHGRTQNDKVAIVGMAGRFPNSADHEALWDLLMKGLDVHRRIPKDRFDADTHVDPSGKGKNKSHTPFGCFIDEPGFFDPRFFNMSPREAAQTDPMGRLALVTAYEALEMSGYVPNRTPSTKLHRIGTFYGQTSDDWREINAAENVDTYFITGGVRAFAPGRINYYFKFSGPSYSVDTACSSSLAAIQLACTSLWAGDCDTACAGGLNVLTNPDIFSGLSKGQFLSKTGSCKTYDNNADGYCRGDAVGTVILKRYEDAIADKDNILGCILGAATNHSAEAVSITHPHAGAQEFLYKRVLANAGVDAHEISYVEMHGTGTQAGDGIEMTSVTNVFAPRHRQRRDDQPVYLGAIKANVGHAEAASGINSLAKVLMMMKHNKIPANVGIKGEMNKTFPADLKDRKVNISQKAVEWPRNGKEKRKVFLNNFSAAGGNTALLLEDGPAYEAPTASDPRGTVPVTVTARSISALKRNIANLQKYVSENPSTTLTSMSYTLTARRIQHNYRVAFPLDQINKFSDALQAQVKESYSPVPNAPTRVAFCFTGQGSQYTGLGQKLYNDLKSFRDDIDQLDHLARVQGLPSFLEIVQGADVQTLSPVKVQLGMACIQVALARMWAAWGITPAAVIGHSLGEYAALHVAGVISASDMVLLVGRRAELLVRDCTPHTHGMLAVKGGAEAIRNTLGNKMTEIACINGPEETVLCGSGDVVGAANETLAAKGFKATKLNVPFAFHSAQVDPILEQFKKIAASVTYNKPAVPVLSPLEGDIIREAGKINPEYLARHARETVNFWTALTAGQKEKVFDEKTAWLEVGAHPVCSGMVKASIGATTTAPSLRRGEDAWKTISNSMCTLFTAGVNFNFDEFHKEFNDAQEMYTLPTYSFDNKKYWLDYHNDWTLRKGEPAQTKEVIVEKPVASASAPAVEIPAKRLSTSCQRVISENFSGNNGSVTVQSSLADPKLYPVVCGHMVNNAALCPSSLYADMALTISDYIWKQMRPGTETPGYNVCNMEVPKPLIAQIPQPAEGQHIQLEANADLDSGIVKLNFRSVKPDGQKLQDHAHCIVRLEDRAAWEDEWSRYNYMVQAQMELLQHKTLNGGAHKVQRGMAYKLFKALVNYDEKYRAMAEVVLASGQTEASAVLDFPTKPEDGDFYCPPYHIDGSCHISGFIVNASDLLDSEQNVYVSHGWGAMKFSRPLTAGMKLRNYVRMQPQPNNVSKGDVYIMEGDQIVAVCEGIKFQQIPRRVLNTFLPPNKGSGPASAAKPAAAPVAAARPAPAAAPIKTAPAPAKAAPAPAPAAPKAAPKPKKAAAPKKAAGGLTAKVMKILAKETEVDEGELVDEAQFENLGVDSLLSLTISAVFREELDMDISSTLFTDYPTVGDMKKYFAQFDNGSSTSSSAEEEDSDEDSIPPTDAATPMDDLSTPASSVPSSAPSDAGKPDSPTRETLEDVGDVSLAKHIVAQEMGVDIAEVTDDADLAEMGMDSLMSLTILGELREKTGIDLPSTFLTTNPTMKDIDNALGMRPKPKAAPKPAAPKAAAPSSSKKTDMNEVSARLSALNNNTDISRYPNATSVLLQGNPKQATKKIFFLPDGSGSATSYVSIPNLGPDVCAYGLNCPFMKNPEQWQCGIEISALVYLAEIKRRQPQGPYIIGGWSAGGVIAYSVAQALLAANEGVEKLLLLDSPCPVNLAPLPARLHNFFNEIGLLGTGDPAKTPKWLLPHFSAAIRSLSDYDPKPSLRPIPTYAIWCREGVAGNPGDPRPPPAEEEDPAPMTWLLEHRTDFKDNGWAQLCGDSMKFGVMGGHHFSMMKPPHADDLGNLIREGLDWQP

>PKSI_1_2516

MSNVLLFGDQTAEQYQLLNKIVLRKENALVITFVERCAKALREETNALPRSQRNAVPDFLTVNDLKEAYHQKGVKVPMVESALVTIAQIGHYIGYFSEHSAEQPSATNTRALGLCTGLLAAAAVVASKTVEELVLVGVEFVRLSFRSGAAVDAARTALCQTGDDNAPWSTIVTGTTEASAKEALAKFHEEKGIPQTSHAYISAVSVMAITVSGPPTTVKRFFEESPALSKNHRVPIPVYGPYHAEHLFGETEINKIASASILEGLKQHQPVSLVHSAATGKALVAENAAELAKLVLAEMLQLPVRWDHLLEEAVSQITSKKAPAKIWAMGVSNVANSLVSALKAGGQTDVSTVDQSTWTENEPDTHGRTQNDKVAIVGMAGRFPNSADHEALWELLMKGLDVHRRIPKDRFDADTHVDPSGKGKNKSHTPFGCFIDEPGFFDPRFFNMSPREAAQTDPMGRLALVTAYEALEMSGYVPNRTPSTKLHRIGTFYGQTSDDWREINAAENVDTYFITGGVRAFAPGRINYYFKFSGPSYSVDTACSSSLAAIQLACTSLWAGDCDTACAGGLNVLTNPDIFSGLSKGQFLSKTGSCKTYDNNADGYCRGDAVGTVILKRYEDAIADKDNILGCILGAATNHSAEAVSITHPHAGAQEFLYKRVLANAGVDAHEISYVEMHGTGTQAGDGIEMTSVTNVFAPRHRQRRDDQPVYLGAIKANVGHAEAASGINSLAKVLLMMKHNKIPANVGIKGEMNKTFPADLKDRKVNISQKAVDWPRNGKEKRKVFLNNFSAAGGNTALLLEDGPAYEAPTATDPRGTVPVTVTARSISALKRNIANLQKYVSENPSTTLTSMSYTLTARRIQHNYRVAFPLDQINKFSDALQAQVKESYSPVPNAPTRVAFCFTGQGSQYTGLGQKLYNDLKSFRDDIDQLDHLARVQGLPSFLEIVQGADVQTLSPVKVQLGMACIQVALGRMWAAWGITPAAVIGHSLGEYAALHVAGVISASDMVLLVGRRAELLVRDCTPHTHGMLAVKGGAEAIRNTLGNKMTEIACINGPEETVLCGSGDVVGAANEVLAAKGFKATKLNVPFAFHSAQVDPILEQFKKIAASVTYNKPSVPVLSPLEGDIIREAGKINPEYLARHARETVNFWTALTAGQKEKVFDEKTAWLEVGAHPVCSGMVKASIGATTTAPSLRRGEDAWKTISNSMCTLFTAGVNFNFDEFHKEFNDAQEMYTLPTYSFDNKKYWLDYHNDWTLRKGEPAQTKEVIVEKPVASASAPAVEMPAKRLSTSCQRVIAENFSGNNGSVTVQSSLADPKLYPVVCGHMVNNAALCPSSLYADMALTISDYIWKQMRPGTETPGYNVCNMEVPKPLIAQIPQPAEGQHIQLEANADLDSGIVKLNFRSVKPDGQKLQDHAHCIVRLEDKAAWEDEWSRYNYMVQAQMELLQHKTLNGGAHKVQRGMAYKLFKALVNYDEKYRAMAEVVLASGQTEASAMLDFPTKPEDGDFYCPPYHIDGSCHISGFIVNASDLLDSEQNVYVSHGWGAMKFSRPLTAGMKLRNYVRMQPQPNNVSKGDVYIMEGDQIVAVCEGIKFQQIPRRVLNTFLPPNKGSGPASAAKPAAAPVAAARPAPAAAPIKTSPAPAKAAPAPAPAAPKAAPKPKKAAAPKKPAGGLTAKVMKILAKETEVDEGELVDEAQFENLGVDSLLSLTISAVFREELDMDISSTLFTDYPTVGDMKKYFAQFDNGSSTSSSAEEEDSDEDSIPPTDAATPMDDLSTPASSVPSSAPSDAGKPDSPTRETLEDVGDVSLAKHIVAQEMGVDIAEVTDDADLAEMGMDSLMSLTILGELREKTGIDLPSTFLTTNPTMKDIDNALGMRPKPKAAPKPAAPKAAAPSSSKKTDMNEVSARLSALNNNTDISRYPNATSVLLQGNPKQATKKIFFLPDGSGSATSYVSIPNLGPDVCAYGLNCPFMKNPEQWQCGIEISALVYLAEIKRRQPQGPYIIGGWSAGGVIAYSVAQALLAANEGVEKLLLLDSPCPVNLAPLPARLHNFFNQIGLLGTGDPAKTPKWLLPHFSAAIRSLSDYDPKPSLRPIPTYAIWCREGVAGNPGDPRPPPAEEEDPAPMTWLLEHRTNFKDNGWAQLCGDSMKFGVMGGHHFSMMKPPHADDLGNLIREGLDWQP

>PKSI_2_Hyb_2516

MSRGTDGAALPTASTVSTQSGGPHVIDPTTKDNGPSPVPASKGPTGAHPKNGDIQSEYPSHAVAVVGMAGRFPGAKSVDALWDLLEAGKSTVEPAPMERIGLGHLPPDDPSRMWWGNFLDDVDAFDHEFFRITAREAQTWDPQQRIMLEVAYEALEDGGQFGASLPSRNRDFGCYIGAVMNNYYDNVACHKASAYATKGTSRCYISGAVSHFFGWTGPAITLDTACSSSMVAVHAACKAIIAGECSRAIAGGTNVITSPHDYRNLAAAGFLSPTGQCKPFDSDADGYCRAEGVGVVVLKSLATAIEENDRVLGVIVGSAVSQSGNVGHIAVPDTRAQVLLHRKALSIANLSPEEVSYVEAHGTGTKVGDPIEMAGLREVYCSSPRTSPLLVSSMKGNIGHTEATAGVAGLIKVLLMMNRRSIPPQASHKRLTPRIPALHPDIVAIPRHLAPWPGTARAACVASYGAAGSNAAILVREMSTRHAGGEREGTAQPKIQGEQPLFVSAATRDMLSLRCSEILLWLRKQKDYRLELSDVLFNISIRANHALPYTFCTTVSSMDGLELKLAEVAKESMVLRKIGSAKPTILVLGGQQGQYVGLSWLCYQEWQGLRRNLDQCHESLIDMGHGGLYPDIFQKAPLVDLKAFHAALFASQYSSAKAWIDSGLKVDAVVGHSFGQLAALCICGVLSLADGLKLVTGRASLIEKHWGSERGAMLSLSAPTRVVDRLLEALEAKIDYAQVACYNGPENHVVVGSEHAIKVLERHIDSRSDLRGSVVAQRLDVTLGFHSVYTEAVQPALEKLASKLEWKAPTIHLETCDTEASCAVFNHSFVVHHMRRPVHFQQAVERLSLRYPESCWISAGHAASFLRLAQKSLRGRGHHTFICLDKVFSEASGSLARAIVELWNEGQPVQYEPFERSRQHRYPHLTPPPYPFARGRHWLHFTNPTTSQPSAICPASQQPQDEDDFLSLVGGDAQGGAEFFIPAAHPRFQLLCNGHVMSGEALAPASFYLEVAARAALKLRCDIKSGQLGLRVEKLSMNAPVGNDPRNIIRLRLEQHEGPRDSWLFSMTMERSPRADTDDINPQKQVVGVVSLRSQDQTNAVAGFKDFRISDVRRRYQEIVDDESAERMQGNHVYRAFDSVVTYGEIFRRIKSIACVGTEAAGEIEWNRRCEDTHVAAATDTQFIDGFLQAAGFLVNYFNNDDFQSSLFICQKIDQIDFADAIPTDAHNLSFYSRMRMDSEDAAIADVLVLDARQQQVLFAASGLQFKKLSRVPLARSLRAANAVAPCSTDGLRKQIHTGRETASRDGSESGHRPDHRRKVLEVLANVTGTPPECVSMAGPLEDLGIDSLAAIEVLNDLRASLDLVIPLSTLMSFNSIGAFVEYVTSTVGSRDRRHRDPHDVLKDARLAKKVTSTAVGYEQVARDEENQVSVASSEPKHDMSLDVDSSAVDLDYLAAVYPSHTRLFLAYIVEAFGGIGCDLATMAAGEAMPEMQNILPRHKRLLHRFFLALVEARILIETPEGTFVRTEQHVDKARAADLHSAAMRHWPEHGSLHQLMHAVGSRTAECLTGVFDGLQIVFGDQANETALREFYQHWPLFKGATKSLAKIVLRVAGHLGRTGKIQILEVGAGTGGTTRPLLAALQAAEMPFDYHFTDVSPHLVNAARQSLDGIAGLTFGVLDVEEQPPKDLEGAYDVVLASNCIHATRDLEQSLGHLRCLLREDGIFAMVEITRRLNIFDLVFGLFDGWWRFNDNRTHCLADEHHWSSKLMAAGFQDVMWTDVGNPKAQVVRVATASPGTTPRREASKRVAKTRVCEVVYKTSSELDILADVYCPARADPAVTLPIALMIHGGSHVLFSRKDVRPAQTRTLLGMGFLPVSIDHRLCPETKLVEGPMVDVCDALDWARHTLPAIDLGDCSLKPDGGCVVAVGWSSGGQLAMSLAWTAPQRGLEPPTAILALYCPTDYEDDWWQHPIQPCGAEDVDEEYDLLDAVQPSPISSYDSICSWEPLTDVRIRQDPRCRIVLHMNWKAQTLPIIVSGLPCSSSVSDEDSSRIDWTALPQPPSREIQRCSALAQIKLGNYRTPTFLVHGTKDELIPWQQSHRTWKCLVSHGIDAKLALIKDAPHICDASKDHESAGWKAVLRGYGFLADKVRIRGHCSALEGRALPE

>PKSI_3_Hybr_2516

MAKIACPQGKPDDIAVIGLAYRFPGGADSDAKLWDLLAERKSAHSKIPVDRFNIDAFHRLGATHSDNVAADGAHLLEQDISAFDAPFFGITTEEAKVIDPQARLLLECSYEALMNAGLKLESISGSDTGCYVGCFDLDYHQMLMGDFECAPKYSGTGTAFSLLSNRLSWFYNLKGPSLSLDTACSSSLVGLHLACQSLSAGESSMAMVCGAKLLLGPHLSMWLSRLNMFSSDGKSRSFADDTAGYGRGEGIATVILKPLADALRDNDPIRAVIKGTGVNQDGHTKGITVPNPESQLDLIRSTYLAAELSFADTGYFEAHGTGTAVGDPLELAAVAQAIVESRRSEPLYVGSIKSNIGHTEGAAGLAGLIKCILMLEKGVILPNIHFDRPNKRIPFERDGIKVPTEVLPWPKDLDRRASINSFGFGGTNAHAIVESFPTPSISSLDHVAQVDASEVVAVTPNHPRLFVLSGHEPAAVEKLRQRYVEYIIQAKNSDGVDRKLDDLSYTLGCRRSRMDWSVFHVASDFEELGKKLLENTTALKRAARSPRIGFIFTGQGAQWPRMGAGLMRYGVFQESVQAADRFLSGQCHCGWSVIDELGKAKDESRIASSELAQPICTIIQVAMIDLLRCWNIRPTAVAGHSSGEIAAAYCTGAITRQSAWEIAFHRGRECARLKEVAPDLRGAMLAVGLGVEDVRPYLDAVAPDRVNIACINSPNSTTLSGDAVEIQELLTKLIANGVSARELRVVNAYHSHHMKLVADRYLDSIAHVAVQTEVVRSDVALLSSVTGTSASSSDLDPEYWVRNLVSPVLFSDAVSAMLKGTKKTFRRQQGTAEPAVDVLLEIGPHAALRGPLADILKSEAVESVAYVPTLNRGSDALESATAAAGELWSRGCPVDVTAVNNHPRQPRVLADLPSYPWDRLNKYWATSRVTHDVLHRAFPHHDILGKRLTGSDALAPAWRHFLRFSESPWVREHVVHGSIVFPGAGFLAMAIEAALQLVEKDRKLANVRMRDVHISKALVLEQEEEGAQELVTRFHRVDDRSDGTWSGWWEFSISCTKSRVEPERHISGQIMLEYRPMEPSSQPASDVVHQAQKVKYGKLVGSSADRLYRAAFYEASEAAGLAYGSQFQGVVDVARSGDGRCCWRVQLPDRKAPASESKHLIHPTTLDAIVHAMFGAMNKGSALSSAALPIAFDKVVISADMPTDAGTCLSGFTVTDVKETTREVTADVYVSSDDWARRLLQIEGLRCTELPSPEGVHNDNETQSAPLGSVAWQPDLDLLDNDGLKSYIMKNPERDQLGATDGRVAALSTRLRSAVAQILYLAAFKTPTLSILQVGGFSEGLTDTFVATIASDTNVPAVTAEILVLDSEEENIPRIRERQYDGCSVAASHWNRAGSLVSHIPKGDKFDAVLLVVPEHNDELATRVLVAQARNLLKAGALLIVLDTIQNTRANLAFYGFETAQTSSSGRLRPWQSLNVEDPDCGKAAALCRESPATISREITGAPICVLKPPHCTKETVDVISALEGILNDARLEVAFQEWPPSVAQVEGKFVISLLDTEASFLSNIDAADFEVLREVALQSRRLLWVCSGDDPHMAIALGWLRVLQNENANRVYQHLTLTRKKAAAPQNHSCAIARLALTQTREREFAEEDGNLYIPRWYHEEGLSGTLEGREMSVQLERAPLGVAKSSMPLRILHGKDAESARFVPDSPSVSRLAADEVEIEMQYIVLTDSDISPVGNTAHREGSGVVRAVGRDVTRLGLDDQVCVSYVGPLSTRVIAKEVYCQVVPPDAIMEEAACIPNTFATALRVLTDVARVKPGQTVLVQTAGTKIGRAAVLLALALDAVVYATAREKTETDKIVSLGVRLQNVFTEGDMDLPQAAKTLAGERGLDVVLRTSKNSSAPCLLPHCVAQHGTLVDVHAASEPGPDVPSCEDTISVMGVGPLLPEDPVSLQKSASRAANYLPQLSGLAASFDLFESARISDALECQQEQGTRGGVILSLDDADMVPIAPSVNRKLYLCEQATYVLAGGLGGLGQSLARLLVDHGARNLALLSRGGPNSPSAETFIKEMAEVGVAVKVLACDIGDESSMKAALDDCAGTMPPIRGVIQAATVYRDAIFDNFTFEDWQANLRAKVQGSWNLHRHLPKDIDFFVMLGSVAGLMGHVSQAGYAAGNTFQDALAHYRRSRSLPAVTIDLGPMLDVGAVNDGTVSASFSTSEATWMTEADLHAIMIMCISGEITSCNFPPQLCTGLPSGGMLQLGQHEMPEHYDRPIFALLKGLGVSAAAGKDNKVLSRRSKDFSHQLPAVTSMEEARKCVVNAVKAHLAKGLGRSADSIKSSQPLDSYGIDSLGAMGFRNWVRETMKADVSMFDVLNARSIKELAAKIVRISELIPEGLESERVTREDE

>PKSI_1_2515

MSNVLLFGDQTAEQYQLLNKIVLRKENALVITFVERCAKALREETNALPRSQRNAVPDFLTVNDLKEAYHQKGVKVPMVESALVTIAQIGHYIGYFSEHSAEQPSATNTRALGLCTGLLAAAAVVASKTVEELVLVGVEFVRLSFRSGAAVDAARTALCQTGDDNAPWSTIVTGTTEASAKEALAKFHEEKGIPQTSHAYISAVSVMAITVSGPPTTVKRFFEESPALSKNHRVPIPVYGPYHAEHLFGETEINKIASASILEGLKQHQPVSLVHSAATGKALVAENAAELAKLVLAEMLQLPVRWDHLLEEAVSQITSKKAAAKIWAMGVSNVANSLVSALKAGGQTDVSTVDQSTWTENEPDTHGRTQNDKVAIVGMAGRFPNSADHEALWELLMKGLDVHRRIPKDRFDADTHVDPSGKGKNKSHTPFGCFIDEPGFFDPRFFNMSPREAAQTDPMGRLALVTAYEALEMSGYVPNRTPSTKLHRIGTFYGQTSDDWREINAAENVDTYFITGGVRAFAPGRINYYFKFSGPSYSVDTACSSSLAAIQLACTSLWAGDCDTACAGGLNVLTNPDIFSGLSKGQFLSKTGSCKTYDNNADGYCRGDAVGTVILKRYEDAIADKDNILGCILGAATNHSAEAVSITHPHAGAQEFLYKRVLANAGVDAHEISYVEMHGTGTQAGDGIEMTSVTNVFAPRHRQRRDDQPVYLGAIKANVGHAEAASGINSLAKVLLMMKHNKIPANVGIKGEMNKTFPADLKDRKVNISQKAVDWPRNGKEKRKVFLNNFSAAGGNTALLLEDGPAYEAPTATDPRGTVPVTVTARSISALKRNIANLQKYVSENPSTTLTSMSYTLTARRIQHNYRVAFPLDQINKFSDALQAQVKESYSPVPNAPTRVAFCFTGQGSQYTGLGQKLYNDLKSFRDDIDQLDHLARVQGLPSFLEIVQGADVQTLSPVKVQLGMACIQVALARMWAAWGINPAAVIGHSLGEYAALHVAGVISASDMVLLVGRRAELLVRDCTPHTHGMLAVKGGAEAIRNTLGNKMTEIACINGPEETVLCGSGDVVGAANETLAAKGFKATKLNVPFAFHSAQVDPILEQFKKIAASVTYNKPAVPVLSPLEGDIIREAGKINPEYLARHARETVNFWTALTAGQKEKVFDEKTAWLEVGAHPVCSGMVKASIGATTTAPSLRRGEDAWKTISNSMCTLFTAGVNFNFDEFHKEFNDAQEMYTLPTYSFDNKKYWLDYHNDWTLRKGEPAQTKEVIVEKPVASASAPAVEMPAKRLSTSCQRVIAENFSGNNGNVTVQSSLADPKLYPVVCGHMVNNAALCPSSLYADMALTISDYIWKQMRPGTETPGYNVCNMEVPKPLIAQIPQPAEGQHIQLEANADLDSGIVKLNFRSVKPDGQKLQDHAHCIVRLEDKAAWEDEWSRYNYMVQAQMELLQHKTLNGGAHKVQRGMAYKLFKALVNYDEKYRAMAEVVLASGQTEASAMLDFPTKPEDGDFYCPPYHIDGSCHISGFIVNASDLLDSEQNVYVSHGWGAMKFSRPLTAGMKLRNYVRMQPQPNNVSKGDVYIMEGDQIVAVCEGIKFQQIPRRVLNTFLPPNKGSGPASAAKPAAAPVAAARPAPAAAPIKTAPAPAKAAPAPAPAAPKAASKPKKAAAPKKPAGGLTAKVMKILAKETEVDEGELVDEAQFENLGVDSLLSLTISAVFREELDMDISSTLFTDYPTVGDMKKYFAQFDNGSSTSSSAEEEDSDEDSIPPTDAATPMDDLSTPASSVPSSAPSDAGKPDSPTRETLEDVGDVSLAKHIVAQEMGVDIAEVTDDADLAEMGMDSLMSLTILGELREKTGIDLPSTFLTTNPTMKDIDNALGMRPKPKAAPKPAAPKAAAPSSSKKTDMNEVSARLSALNNNTDISRYPNATSVLLQGNPKQATKKIFFLPDGSGSATSYVSIPNLGPDVCAYGLNCPFMKNPEQWQCGIEISALVYLAEIKRRQPQGPYIIGGWSAGGVIAYSVAQALLAANEGVEKLLLLDSPCPVNLAPLPARLHNFFNQIGLLGTGDPAKTPKWLLPHFSAAIRSLSDYDPKPSLRPIPTYAIWCREGVAGNPGDPRPPPAEEEDPAPMTWLLEHRTNFKDNGWAQLCGDSMKFGVMGGHHFSMMKPPHADDLGNLIREGLDWQP

>PKS_1_561

MSNVLLFGDQTAEQYPLLNKIVLRKENALVITFIERCAKALREETNALPRSQRNVVPDFLTINDLKEAYHQKGVKVPMVESALVTIAQIGHYIGYFSEHSAEQPSATNTRALGLCTGLLAAAAVVASKTVEELVLVGVEFVRLSFRSGAAVDSARTALCQTGDDNAPWSTIVTGTTEAAAKEALAKFHEEKGIPQTSHAYISAVSVMAITVSGPPTTVKRFFEESSALSKNHRVPIPVYGPYHAEHLFGETEINKIASASILEGLKQHQPVSLVHSAATGKALVAENAAELAKLVLAEMLQHPVRWDHLLEEAVSQITSKKAPAKIWAMGVSNVANSLVSALKAGGQTDVSTVDQSTWTENEPDTHGRTQNDKVAIVGMAGRFPNSADHEALWELLMKGLDVHRRIPKDRFDADTHVDPSGKGKNKSHTPFGCFIDEPGFFDPRFFNMSPREAAQTDPMGRLALVTAYEALEMSGYVPNRTPSTKLHRIGTFYGQTSDDWREINAAENVDTYFITGGVRAFAPGRINYYFKFSGPSYSVDTACSSSLAAIQLACTSLWAGDCDTACAGGLNVLTNPDIFSGLSKGQFLSKTGSCKTYDNNADGYCRGDAVGTVILKRYEDAIADKDNILGCILGAATNHSAEAVSITHPHAGAQEFLYKRVLANAGVDAHEISYVEMHGTGTQAGDGIEMTSVTNVFAPRHRQRRDDQPVYLGAIKANVGHAEAASGINSLAKVLLMMKHNKIPANVGIKGEMNKTFPADLKDRKVNISQKAVDWPRNGKEKRKVFLNNFSAAGGNTALLLEDGPAYEAPTATDPRGTVPVTVTARSISALKRNIANLQKYVSENPSTTLTSMSYTLTARRIQHNYRVAFPLDQINKFSDALQAQVKESYSPVPNAPTRVAFCFTGQGSQYTGLGQKLYNDLKSFRDDIDQLDHLARVQGLPSFLEIVQGADVQTLSPVKVQLGMACIQVALARMWAAWGITPAAVIGHSLGEYAALHVAGVISASDMVLLVGRRAELLVRDCTPHTHGMLAVKGGAEAIRNTLGSKMTEIACINGPEETVLCGSGDVVGAANETLAAKGFKATKLNVPFAFHSAQVDPILEQFKKIAASVTYNKPAVPVLSPLEGDIIREAGKINPEYLARHARETVNFWTALTAGQKEKVFDEKTAWLEVGAHPVCSGMVKASIGATTTAPSLRRGEDAWKTISNSMCTLFTAGVNFNFDEFHKEFNDAQEMYTLPTYSFDNKKYWLDYHNDWTLRKGEPAQTKEVIVEKPVASASAPAVEMPAKRLSTSCQRVVAENFSGNNGSVTVQSSLADPKLYPVVCGHMVNNAALCPSSLYADMALTISDYIWKQMRPGTETPGYNVCNMEVPKPLIAQIPQPAEGQHIQLEANADLDSGIVKLNFRSVKPDGQKLQDHAHCIVRLEDKAAWEDEWSRYNYMVQAQMELLQHKTLNGGAHKVQRGMAYKLFKALVNYDEKYRAMAEVVLASGQTEASAELDFPTKPEDGDFYCPPYHIDGSCHISGFIVNASDLLDSEQNVYVSHGWGAMKFSRPLTAGMKLRNYVRMQPQPNNVSKGDVYIMEGDQIVAVCEGIKFQQIPRRVLNTFLPPNKGSGPASAAKPAAAPVAAARPAPAAAPIKTAPAPAKAALAPAPAAPKAAPKPKKAAAPKKAAGGLTAKVMKILAKETEVDEGELVDEAQFENLGVDSLLSLTISAVFREELDMDISSTLFTDYPTVGDMKKYFAQFDNGSSTSSSTEEDDSDEDSIPPTDAATPMDDLSTPASSVGSSAPSDAGKPDSPTRETLEDVGDVSLAKHIVAQEMGVDIAEVTDDADLAEMGMDSLMSLTILGELREKTGIDLPSTFLTTNPTMKDIDNALGMRPKPKAAPKPAAPKAAAPSSSKKTDMNEVSARLSALNNNTDISRYPNATSVLLQGNPKQATKKIFFLPDGSGSATSYVSIPNLGPDVCAYGLNCPFMKNPEQWQCGIEISALVYLAEIKRRQPQGPYIIGGWSAGGVIAYSVAQALLAANEGVEKLLLLDSPCPVNLAPLPARLHNFFNEIGLLGTGDPAKTPKWLLPHFSAAIRSLSDYDPKPSLRPIPTYAIWCREGVAGNPGDPRPPPAEEEDPAPMTWLLEHRTNFKDNGWAQLCGDSMKFGVMGGHHFSMMKPPHADDLGNLIREGLDWQP

>PKSI_1_269

MPDNVSFMDESQDLRHIRETSRTSHTTSILNVEGDYRNNDMPREPGECNRSTNGTVDHERMTPSADGGIPIAICGIGLRLPGGIRNDRDLYDSLYNKKDARGVIPEDRFSIDSFHSAHGKTGTIITKHGYFLQDIDLTKFDVNMFNMTPAEVERLDPHQRILLETVRETLESAGEASFRGKKVGTYVGNFTDDWLDLQNVDTVDFATYQLHGKMDFSLANRISYEYDLRGPSMTIKTACSSSALAIHEAVYSIRNGECDAAIVSGSNLNLAPRLWVGMSSQGAISPDGSSKTFDESANGYARGDGIAALFIKRLDDAVRDGNPVRAVIRSTASNADGRTPGMTMPSTEAQEALIRRAYDAANLPLSETAMVECHGTGTAVGDPMEANAVARCFGDQGMLIGSVKPNLGHSEGASAITSVVKAVLSLENRTILPNIKFHRPNPAIPWSEAKLTVPVEPLAWPKDRQERISVNSFGIGGSNVHVVLDSAASMGFRPRSLAPSKDDRPGRLLLFSGGHRASVEQSSSQHQDYVTKYPNRLSDVAYTLAKRPPVKCPGLTRSTFVFTGQGAQWLHMGKELLHESPVFAKSIGRMDSVIHSLKHAPQWTLEGIINDPENPSALTNAEISQPLCTAVQIGLVDLLKSWAIYPHAVLGHSSGEIGAAYASGVVDRAEAILLAFYRGYVCRFAQKAGGMAAVGLEKSQVIKYLQPGVCVACENSGSSVTLSGDLETLEEVLQSIRAENLNAFARKLQVGIAYHSDHMKALGGLYHQYITEHLDPKDPQVPFFSSVSGRALHSKDDFGATYWQDNLENPVLFHTAVLKSLEHTGDKQVHLEVGPHGALNGPLRQIYAETGSKARYVALQKRGANCFDTFLEGIGQLYCNGVPLQYPESADDRTLIDLPPYPWHYDHSYWSETRVMKNWRFRRQLPHDLLGLRTLDCSDAEPMWRNILRITDLPWLRDHCVGKDVVFPASGYMCMAGEAVFQETGCRDYTLREVDISTAMVLSSDHSTELLTTMKKRRLNAFLDSRWYEFLIMSYDGASWTQHCSGLVTNGPSTSHPKALLQTYDRPVSTNRWYTAMSKIGLNYGPRFTGLQNITTHVQEKKASMTIMDKQEDYESPYALHPSTLDLILQSWTVASVRGEYRRFTQLFLPTFVDEFYIGNSASKLIHLNTTAIGPDGSARGEAIGKDNDGQISFNLKGFKGSKLDNVGVDQPQEMQTIMKQQWKRDFDFADTAQLMRPAFDSTSELSLLERMFVLAAIEVHVRTSGMEGKLPHHQRYKLWIDAQIRRFGEPGYPMVEDSMELLRLDSRERQRQLCRLLEHSRKTTAHPVAEAIWRALDRIEDVFDGRIEYLDLLFNDGLMPKFYDWSNSLSDVSRLFRLLSHKKPQLKILEVGAGTGGSTARLLQYLQSDFGERQYHSYTYTDVSSGFFVQAQERFKDYEGMKYRVLDISQDPFEQGFGADEFDLICASNVLHATPRLTETLRNCRKMLRPDGILFLQELCPRQQFMGFIMGLFEGWWLGAEDGRADTPLLLPPAWDRRLRDVGFEGVEAFSFDNNPPYFMAANMTARPTVTAKSKGSITLLTFNDFLDDVAGALKQAIRAAGFEIDHCVWGEHVPLDQSLISLVDLERDKPLLQDIGDDDLRIFLDLIDAVLQTTVIWLNKPAQVSSADPNAAQMLGLARTLRAELAMHFATVEMADPTLNGMSAVVHLMCMLQRGSALPENSLDQDMEYVWAHDAMHVSRFHFQPVDEALMEMSPKFDVKTLVPLQRGMLSSLKWIGTRMLPLAEAEVQIRMSAVGMNFHDMMIAMNMFDSPLTLGSGYNSIGMEGVGYVTRKSPEVDHVQVGDRVIVIGSNSSGFATDVHRPADYCIKCPSSLTDVEAAGMSFAYMTVLWSFLDKGGLRKGQSVLIHSAAGGVGIAALHVSRWLGLEAFVTVGNEEKVRFIMNNFGLPRNRIFNSHSATFLDDVMDATAGRGVDAALSAAAGELLHNTWSCIAPGGVMLEIGKRDLINRGRLSLAPFEENRSYVGIDFSRLTIVNKPAVVRLLRQTMRLVEQGHIHPIHPTTCFDAEHAEDAFRFMQTGQHIGRIVVKIPRDTSTIPLASRPPAPEFEGQKTYLLVGGMGGLGRSVASWMVSAGARNLIFMSRSAGKSEQDQNFARELELSGCKVHCCAIDITDTDAVREALERTQASVAGVLQMAMVLRDVGIMNMDKANWDAAVAPKVQGTWNLHHALPNVDFFVMFGSNSGTLGSYGQANYAAANAFLDSFVQYRQNLGQAASVIDIGAVGDVGYVAETQVAAENMESMAGRLISEQDFLNCLQLAIARSTPTERRNKGPSTEADGYIDLKQVILLNSSILSMADPSNQIFWRKDPRMGIYRNVQRTSAESPTTDSNSLRRLVATLKADSSIADQAELAQTLARELSKQVATVLMLGEDEIDVDRSLTDVGMDSLVAIEMRNWWKQNLGVDVSVLELKDGRSILRLGELAATRLKERYSRDS

>PKSI_2_269

MSNVLLFGDQTAEQYPLLNKIVLRKENALVITFVERCAKALREETNALPRSQRNAVPDFLTVNDLKEAYHQKGVKVPMVESALVTIAQIGHYIGYFSEHSAEQPSATNTRALGLCTGLLAAAAVVASKTVEELVLVGVEFVRLSFRSGAAVDAARTALCQTGDDNAPWSTIVTGTTEASAKEALAKFHEEKGIPQTNHAYISAVSVMAITVSGPPTTVKRFFEESPALSKNHRVPIPVYGPYHAEHLFGETEINKIASASILEGLKQHQPVSLVHSAATGKALVAENAAELAKLVLAEMLQLPVRWDHLLEEAVSQITSKKAPAKIWAMGVSNVANSLVSALKAGGQTDVSTVDQGTWTENEPDTHGRTQNDKVAIVGMAGRFPNSADHEALWELLMKGLDVHRRIPKDRFDADTHVDPSGKGKNKSHTPFGCFIDEPGFFDPRFFNMSPREAAQTDPMGRLALVTAYEALEMSGYVPNRTPSTKLHRIGTFYGQTSDDWREINAAENVDTYFITGGVRAFAPGRINYYFKFSGPSYSVDTACSSSLAAIQLACTSLWAGDCDTACAGGLNVLTNPDIFSGLSKGQFLSKTGSCKTYDNNADGYCRGDAVGTVILKRYEDAIADKDNILGCILGAATNHSAEAVSITHPHAGAQEFLYKRVLANAGVDAHEISYVEMHGTGTQAGDGIEMTSVTNVFAPRHRQRRDDQPVYLGAIKANVGHAEAASGINSLAKVLLMMKHNKIPANVGIKGEMNKTFPADLKDRKVNISQKAVDWPRNGKEKRKVFLNNFSAAGGNTALLLEDGPAYEAPTATDPRGTVPVTVTARSISALKRNIANLQKYVSENPSTTLTSMSYTLTARRIQHNYRVAFPLDQINKFSDALQAQVKESYSPVPNAPTRVAFCFTGQGSQYTGLGQKLYNDLKSFRDDIDQLDHLARVQGLPSFLEIVQGADVQTLSPVKVQLGMACIQVALARMWAAWGITPAAVIGHSLGEYAALHVAGVISASDMVLLVGRRAELLVRDCTPHTHGMLAVKGGAEAIRNTLGNKMTEIACINGPEETVLCGSGDVVGAANETLAAKGFKATKLNVPFAFHSAQVDPILEQFKKIAASVTYNKPAVPVLSPLEGDIIREAGKINPEYLARHARETVNFWTALTAGQKEKVFDEKTAWLEVGAHPVCSGMVKASIGATTTAPSLRRGEDAWKTISNSMC

>PKSI_3_269

MSNVLLFGDQTAEQYPLLNKIVLRKENALVITFVERCAKALREETNALPRSQRNAVPDFLTVNDLKEAYHQKGVKVPMVESALVTIAQIGHYIGYFSEHSAEQPSATNTRALGLCTGLLAAAAVVASKTVEELVLVGVGFVRLSFRSGAAVDAARTALCQTGDDNAPWSTIVTGTTEASAKEALAKFHEEKGIPQTSHAYISAVSVMAITVSGPPTTVKRFFEESPALSKNHRVPIPVYGPYHAEHLFGETEINKIASDSILEGLKQHQPVSLVHSAATGKALVAENAAELAKLVLAEMLQHSVRWDHLLEEAVSQITSKKAPAKIWAMGVSNVANSLVSALKAGGQSDVSTVDQSSWTENEPDTHGRTQNDKVAIVGMAGRFPNSADHEALWDLLMKGLDVHRRIPKDRFDADTHVDPSGKGKNKSHTPFGCFIDEPGFFDPRFFNMSPREAAQTDPMGRLALVTAYEALEMSGYVPNRTPSTKLHRIGTFYGQTSDDWREINAAENVDTYFITGGVRAFAPGRINYYFKFSGPSYSVDTACSSSLAAIQLACTSLWAGDCDTACAGGLNVLTNPDIFSGLSKGQFLSKTGSCKTYDNNADGYCRGDAVGTVILKRYEDAIADKDNILGCILGAATNHSAEAVSITHPHAGAQEFLYKRVLANAGVDAHEISYVEMHGTGTQAGDGIEMTSVTNVFAPRHRQRRDDQPVYLGAIKANVGHAEAASGINSLAKVLMMMKHNKIPANVGIKGEMNKTFPADLKDRKVNISQKAVEWPRNGKEKRKVFLNNFSAAGGNTALLLEDGPAYEAPTATDPRGTVPVTVTARSISALKRNIANLQKYVSENPSTTLTSMSYTLTARRIQHNYRVAFPLDQIDKFSDALQAQVKESYSPVPNVPTRVAFCFTGQGSQYTGLGQKLYNDLKSFRDDIDQLDHLARVQGLPSFLEIVQGADVQTLSPVKVQLGMACIQVALARMWAAWGITPAAVIGHSLGEYAALHVAGVISASDMVLVVGRRAELLVRDCTPHTHGMLAVKGGAEAIRNTLGNKMTEIACINGPEETVLCGSGDVVGAANETLAAKGFKATKLNVPFAFHSAQVDPILEQFKKIAASVTYNKPAVPVLSPLEGDIIREAGKINPEYLARHARETVNFWTALTAGQKEKVFDEKTAWLEVGAHPVCSGMVKASIGATTTAPSLRRGEDAWKTISNSMCTLFTAGVNFNFDEFHKEFNDAQEMYTLPTYSFDNKKYWLDYHNDWTLRKGEPAQTKEVIVEKPVASASAPAVEMPAKRLSTSCQRVIAENFSGNNGSVTVQSSLADPKLYPVVCGHMVNNAALCPSSLYADMALTISDYIWKQMRPGTETPGYNVCNMEVPKPLIAQIPQPAEGQHIQLEANADLDSGIVKLNFRSVKPDGQKLQDHAHCIVRLEDKAAWEDEWSRYNYMVQAQMELLQHKTLNGGAHKVQRGMAYKLFKALVNYDEKYRAMAEVVLASGQTEASAMLDFPTKPEDGDFYCPPYHIDGSCHISGFIVNASDLLDSEQNVYVSHGWGAMKFSRPLTAGMKLRNYVRMQPQPNNVSKGDVYIMEGDQIVAVCEGIKFQQIPRRVLNTFLPPNKGSGPASAAKPAAAPVAAARPAPAAAPIKTAPAPAKAAPAPAPAAPKAAPKPKKAAAPKKPAGGLTAKVMKILAKETEVDEGELVDEAQFENLGVDSLLSLTISAVFREELDMDISSTLFTDYPTVGDMKKYFAQFDNGSSTSSSAEEEDSDEDSIPPTDAATPMDDLSTPASSVPSSAPSDAGKPDSPTRETLEDVGDVSLAKHIVAQEMGVDIAEVTDDADLAEM

>PKSI_1_10512

MPDNVSFMDESQDLRHIRETSRTSHTTSILNVEGDYRNNDMPREPGECNRSTNGTVDHERMTPSADGGIPIAICGIGLRLPGGIRNDRDLYDSLYNKKDARGVIPEDRFSIDSFHSAHGKTGTIITKHGYFLQDIDLTKFDVNMFNMTPAEVERLDPHQRILLETVRETLESAGEASFRGKKVGTYVGNFTDDWLDLQNVDTVDFATYQLHGKMDFSLANRISYEYDLRGPSMTIKTACSSSALAIHEAVYSIRNGECDAAIVSGSNLNLAPRLWVGMSSQGAISPDGSSKTFDESANGYARGDGIAALFIKRLDDAVRDGNPVRAVIRSTASNADGRTPGMTMPSTEAQEALIRRAYDAANLPLSETAMVECHGTGTAVGDPMEANAVARCFGDQGMLIGSVKPNLGHSEGASAITSVVKAVLSLENRTILPNIKFHRPNPAIPWSEAKLTVPVEPLAWPKDRQERISVNSFGIGGSNVHVVLDSAASMGFRPRSLAPSKDDRPGRLLLFSGGHRASVEQSSSQHQDYVTKYPNRLSDVAYTLAKRPPVKCPGLTRSTFVFTGQGAQWLHMGKELLHESPVFAKSIGRMDSVIHSLKHAPQWTLEGIINDPENPSALTNAEISQPLCTAVQIGLVDLLKSWAIYPHAVLGHSSGEIGAAYASGVVDRAEAILLAFYRGYVCRFAQKAGGMAAVGLEKSQVIKYLQPGVCVACENSGSSVTLSGDLETLEEVLQSIRAENLNAFARKLQVGIAYHSDHMKALGGLYHQYITEHLDPKDPQVPFFSSVSGRALHSKDDFGATYWQDNLENPVLFHTAVLKSLEHTGDKQVHLEVGPHGALNGPLRQIYAETGSKARYVALQKRGANCFDTFLEGIGQLYCNGVPLQYPESADDRTLIDLPPYPWHYDHSYWSETRVMKNWRFRRQLPHDLLGLRTLDCSDAEPMWRNILRITDLPWLRDHCVGKDVVFPASGYMCMAGEAVFQETGCRDYTLREVDISTAMVLSSDHSTELLTTMKKRRLNAFLDSRWYEFLIMSYDGASWTQHCSGLVTNGPSTSHPKALLQTYDRPVSTNRWYTAMSKIGLNYGPRFTGLQNITTHVQEKKASMTIMDKQEDYESPYALHPSTLDLILQSWTVASVRGEYRRFTQLFLPTFVDEFYIGNSASKLIHLNTTAIGPDGSARGEAIGKDNDGQISFNLKGFKGSKLDNVGVDQPQEMQTIMKQQWKRDFDFADTAQLMRPAFDSTSELSLLERMFVLAAIEVHVRTSGMEGKLPHHQRYKLWIDAQIRRFGEPGYPMVEDSMELLRLDSRERQRQLCRLLEHSRKTTAHPVAEAIWRALDRIEDVFDGRIEYLDLLFNDGLMPKFYDWSNSLSDVSRLFRLLSHKKPQLKILEVGAGTGGSTARLLQYLQSDFGERQYHSYTYTDVSSGFFVQAQERFKDYEGMKYRVLDISQDPFEQGFGADEFDLICASNVLHATPRLTETLRNCRKMLRPDGILFLQELCPRQQFMGFIMGLFEGWWLGAEDGRADTPLLLPPAWDRRLRDVGFEGVEAFSFDNNPPYFMAANMTARPTVTAKSKGSITLLTFNDFLDDVAGALKQAIRAAGFEIDHCVWGEHVPLDQSLISLVDLERDKPLLQDIGDDDLRIFLDLIDAVLQTTVIWLNKPAQVSSADPNAAQMLGLARTLRAELAMHFATVEMADPTLNGMSAVVHLMCMLQRGSALPENSLDQDMEYVWAHDAMHVSRFHFQPVDEALMEMSPKFDVKTLVPLQRGMLSSLKWIGTRMLPLAEAEVQIRMSAVGMNFHDMMIAMNMFDSPLTLGSGYNSIGMEGVGYVTRKSPEVDHVQVGDRVIVIGSNSSGFATDVHRPADYCIKCPSSLTDVEAAGMSFAYMTVLWSFLDKGGLRKGQSVLIHSAAGGVGIAALHVSRWLGLEAFVTVGNEEKVRFIMNNFGLPRNRIFNSHSATFLDDVMDATAGRGVDAALSAAAGELLHNTWSCIAPGGVMLEIGKRDLINRGRLSLAPFEENRSYVGIDFSRLTIVNKPAVVRLLRQTMRLVEQGHIHPIHPTTCFDAEHAEDAFRFMQTGQHIGRIVVKIPRDTSTIPLASRPPAPEFEGQKTYLLVGGMGGLGRSVASWMVSAGARNLIFMSRSAGKSEQDQNFARELELSGCKVHCCAIDITDTDAVREALERTQASVAGVLQMAMVLRDVGIMNMDKANWDAAVAPKVQGTWNLHHALPNVDFFVMFGSNSGTLGSYGQANYAAANAFLDSFVQYRQNLGQAASVIDIGAVGDVGYVAETQVAAENMESMAGRLISEQDFLNCLQLAIARSTPTERRNKGPSTEADGYIDLKQVILLNSSILSMADPSNQIFWRKDPRMGIYRNVQRTSAESPTTDSNSLRRLVATLKADSSIADQAELAQTLARELSKQVATVLMLGEDEIDVDRSLTDVGMDSLVAIEMRNWWKQNLGVDVSVLELKDGRSILRLGELAATRLKERYSRDS

>PKSI_1_156

MPDNVSFMDESQDLRHIRETSRTSHTTSILNVEGDYRNNDMPREPGECNRSTNGTVDHERMTPSADGGIPIAICGIGLRLPGGIRNDRDLYDSLYNKKDARGVIPEDRFSIDSFHSAHGKTGTIITKHGYFLQDIDLTKFDVNMFNMTPAEVERLDPHQRILLETVRETLESAGEASFRGKKVGTYVGNFTDDWLDLQNVDTVDFATYQLHGKMDFSLANRISYEYDLRGPSMTIKTACSSSALAIHEAVYSIRNGECDAAIVSGSNLNLAPRLWVGMSSQGAISPDGSSKTFDESANGYARGDGIAALFIKRLDDAVRDGNPVRAVIRSTASNADGRTPGMTMPSTEAQEALIRRAYDAANLPLSETAMVECHGTGTAVGDPMEANAVARCFGDQGMLIGSVKPNLGHSEGASAITSVVKAVLSLENRTILPNIKFHRPNPAIPWSEAKLTVPVEPLAWPKDRQERISVNSFGIGGSNVHVVLDSAASMGFRPRSLAPSKDDRPGRLLLFSGGHRASVEQSSSQHQDYVTKYPNRLSDVAYTLAKRPPVKCPGLTRSTFVFTGQGAQWLHMGKELLHESPVFAKSIGRMDSVIHSLKHAPQWTLEGIINDPENPSALTNAEISQPLCTAVQIGLVDLLKSWAIYPHAVLGHSSGEIGAAYASGVVDRAEAILLAFYRGYVCRFAQKAGGMAAVGLEKSQVIKYLQPGVCVACENSGSSVTLSGDLETLEEVLQSIRAENLNAFARKLQVGIAYHSDHMKALGGLYHQYITEHLDPKDPQVPFFSSVSGRALHSKDDFGATYWQDNLENPVLFHTAVLKSLEHTGDKQVHLEVGPHGALNGPLRQIYAETGSKARYVALQKRGANCFDTFLEGIGQLYCNGVPLQYPESADDRTLIDLPPYPWHYDHSYWSETRVMKNWRFRRQLPHDLLGLRTLDCSDAEPMWRNILRITDLPWLRDHCVGKDVVFPASGYMCMAGEAVFQETGCRDYTLREVDISTAMVLSSDHSTELLTTMKKRRLNAFLDSRWYEFLIMSYDGASWTQHCSGLVTNGPSTSHPKALLQTYDRPVSTNRWYTAMSKIGLNYGPRFTGLQNITTHVQEKKASMTIMDKQEDYESPYALHPSTLDLILQSWTVASVRGEYRRFTQLFLPTFVDEFYIGNSASKLIHLNTTAIGPDGSARGEAIGKDNDGQISFNLKGFKGSKLDNVGVDQPQEMQTIMKQQWKRDFDFADTAQLMRPAFDSTSELSLLERMFVLAAIEVHVRTSGMEGKLPHHQRYKLWIDAQIRRFGEPGYPMVEDSMELLRLDSRERQRQLCRLLEHSRKTTAHPVAEAIWRALDRIEDVFDGRIEYLDLLFNDGLMPKFYDWSNSLSDVSRLFRLLSHKKPQLKILEVGAGTGGSTARLLQYLQSDFGERQYHSYTYTDVSSGFFVQAQERFKDYEGMKYRVLDISQDPFEQGFGADEFDLICASNVLHATPRLTETLRNCRKMLRPDGILFLQELCPRQQFMGFIMGLFEGWWLGAEDGRADTPLLLPPAWDRRLRDVGFEGVEAFSFDNNPPYFMAANMTARPTVTAKSKGSITLLTFNDFLDDVAGALKQAIRAAGFEIDHCVWGEHVPLDQSLISLVDLERDKPLLQDIGDDDLRIFLDLIDAVLQTTVIWLNKPAQVSSADPNAAQMLGLARTLRAELAMHFATVEMADPTLNGMSAVVHLMCMLQRGSALPENSLDQDMEYVWAHDAMHVSRFHFQPVDEALMEMSPKFDVKTLVPLQRGMLSSLKWIGTRMLPLAEAEVQIRMSAVGMNFHDMMIAMNMFDSPLTLGSGYNSIGMEGVGYVTRKSPEVDHVQVGDRVIVIGSNSSGFATDVHRPADYCIKCPSSLTDVEAAGMSFAYMTVLWSFLDKGGLRKGQSVLIHSAAGGVGIAALHVSRWLGLEAFVTVGNEEKVRFIMNNFGLPRNRIFNSHSATFLDDVMDATAGRGVDAALSAAAGELLHNTWSCIAPGGVMLEIGKRDLINRGRLSLAPFEENRSYVGIDFSRLTIVNKPAVVRLLRQTMRLVEQGHIHPIHPTTCFDAEHAEDAFRFMQTGQHIGRIVVKIPRDTSTIPLASRPPAPEFEGQKTYLLVGGMGGLGRSVASWMVSAGARNLIFMSRSAGKSEQDQNFARELELSGCKVHCCAIDITDTDAVREALERTQASVAGVLQMAMVLRDVGIMNMDKANWDAAVAPKVQGTWNLHHALPNVDFFVMFGSNSGTLGSYGQANYAAANAFLDSFVQYRQNLGQAASVIDIGAVGDVGYVAETQVAAENMESMAGRLISEQDFLNCLQLAIARSTPTERRNKGPSTEADGYIDLKQVILLNSSILSMADPSNQIFWRKDPRMGIYRNVQRTSAESPTTDSNSLRRLVATLKADSSIADQAELAQTLARELSKQVATVLMLGEDEIDVDRSLTDVGMDSLVAIEMRNWWKQNLGVDVSVLELKDGRSILRLGELAATRLKERYSRDS

>PKSI_2_156

MSNVLLFGDQTAEQYPLLNKIVLRKENALVITFVERCAKALREETNALPRSQRNAVPDFLTVNDLKEAYHQKGVKVPMVESALVTIAQIGHYIGYFSEHSAEQPSATNTRALGLCTGLLAAAAVVASKTVEELVLVGVEFVRLSFRSGAAVDAARTALCQTGDDNAPWSTIVTGTTEASAKEALAKFHEEKGIPQTNHAYISAVSVMAITVSGPPTTVKRFFEESPALSKNHRVPIPVYGPYHAEHLFGETEINKIASASILEGLKQHQPVSLVHSAATGKALVAENAAELAKLVLAEMLQLPVRWDHLLEEAVSQITSKKAPAKIWAMGVSNVANSLVSALKAGGQTDVSTVDQGTWTENEPDTHGRTQNDKVAIVGMAGRFPNSADHEALWELLMKGLDVHRRIPKDRFDADTHVDPSGKGKNKSHTPFGCFIDEPGFFDPRFFNMSPREAAQTDPMGRLALVTAYEALEMSGYVPNRTPSTKLHRIGTFYGQTSDDWREINAAENVDTYFITGGVRAFAPGRINYYFKFSGPSYSVDTACSSSLAAIQLACTSLWAGDCDTACAGGLNVLTNPDIFSGLSKGQFLSKTGSCKTYDNNADGYCRGDAVGTVILKRYEDAIADKDNILGCILGAATNHSAEAVSITHPHAGAQEFLYKRVLANAGVDAHEISYVEMHGTGTQAGDGIEMTSVTNVFAPRHRQRRDDQPVYLGAIKANVGHAEAASGINSLAKVLLMMKHNKIPANVGIKGEMNKTFPADLKDRKVNISQKAVDWPRNGKEKRKVFLNNFSAAGGNTALLLEDGPAYEAPTATDPRGTVPVTVTARSISALKRNIANLQKYVSENPSTTLTSMSYTLTARRIQHNYRVAFPLDQINKFSDALQAQVKESYSPVPNAPTRVAFCFTGQGSQYTGLGQKLYNDLKSFRDDIDQLDHLARVQGLPSFLEIVQGADVQTLSPVKVQLGMACIQVALARMWAAWGITPAAVIGHSLGEYAALHVAGVISASDMVLLVGRRAELLVRDCTPHTHGMLAVKGGAEAIRNTLGNKMTEIACINGPEETVLCGSGDVVGAANETLAAKGFKATKLNVPFAFHSAQVDPILEQFKKIAASVTYNKPAVPVLSPLEGDIIREAGKINPEYLARHARETVNFWTALTAGQKEKVFDEKTAWLEVGAHPVCSGMVKASIGATTTAPSLRRGEDAWKTISNSMCTLFTAGVNFNFDEFHKEFNDAQEMY

>PKSI_1_241

MSNVLLFGDQTAERYQLLNKIVLRKENALVITFVERCAKALREETNALPRSQRNAVPDFLTVNDLKEAYHQKGVKVPMVESALVTIAQIGHYIGYFSEHSAEQPSATNTRALGLCTGLLAAAAVVASKTVEELVLVGVDFVRLSFRSGAAVVAARTALCQTGDDNAPWSTIVTGTTEASAKEALAKFHEEKGIPQTSHAYISAVSVMAITVSGPPTTVKRFFEESPALSKNHRVPIPVYGPYHAEHLFGETEINKIASDSILEGLKQHQPVSLVHSAATGKALVAENAAELAKLVLAEMLQHPVRWDHLLEEAVSQITSKKAPAKIWAMGVSNVANSLVSALKAGGQSDVSTVDQSSWTENEPDTHGRTQNDKVAIVGMAGRFPNSADHEALWELLMKGLDVHRRIPKDRFDADTHVDPSGKGKNKSHTPFGCFIDEPGFFDPRFFNMSPREAAQTDPMGRLALVTAYEALEMSGYVPNRTPSTKLHRIGTFYGQTSDDWREINAAENVDTYFITGGVRAFAPGRINYYFKFSGPSYSVDTACSSSLAAIQLACTSLWAGDCDTACAGGLNVLTNPDIFSGLSKGQFLSKTGSCKTYDNNADGYCRGDAVGTVILKRYEDAIADKDNILGCILGAATNHSAEAVSITHPHAGAQEFLYKRVLANAGVDAHEISYVEMHGTGTQAGDGIEMTSVTNVFAPRHRQRRDDQPVYLGAIKANVGHAEAASGINSLAKVLLMMKHNKIPANVGIKGEMNKTFPADLKDRKVNISQKAVEWPRNGKEKRKVFLNNFSAAGGNTALLLEDGPAYEAPTATDPRGTVPVTVTARSISALKRNIANLQKYVSENPSTTLTSMSYTLTARRIQHNYRVAFPLDQIDKFSDALQAQVKESYSPVPNVPTRVAFCFTGQGSQYTGLGQKLYNDLKSFRDDIDQLDHLARVQGLPSFLEIVQGADVQTLSPVKVQLGMACIQVALARMWAAWGITPAAVIGHSLGEYAALHVAGVISASDMVLLVGRRAELLVRDCTPHTHGMLAVKGGAEAIRNTLGNKMTEIACINGPEETVLCGSGDVVGAANETLAAKGFKATKLNVPFAFHSAQVDPILEQFKKIAASVTYNKPAVPVLSPLEGDIIREAGKINPEYLARHARETVNFWTALTAGQKEKVFDEKTAWLEVGAHPVCSGMVKASIGATTTAPSLRRGEDAWKTISNSMCTLFTAGVNFNFDEFHKEFNDAQEMYTLPTYSFDNKKYWLDYHNDWTLRKGEPAQTKEVIVEKPVASASAPAVEMPAKRLSTSCQRVISENFSGNNGSVTVQSSLADPKLYPVVCGHMVNNAALCPSSLYADMALTISDYIWKQMRPGTETPGYNVCNMEVPKPLIAQIPQPAEGQHIQLEANADLDSGIVKLNFRSVKPDGQKLQDHAHCIVRLEDKAAWEDEWSRYNYMVQAQMELLQHKTLNGGAHKVQRGMAYKLFKALVNYDEKYRAMAEVVLASGQTEASAVLDFPTKPEDGDFYCPPYHIDGSCHISGFIVNASDLLDSEQNVYVSHGWGAMKFSRPLTAGMKLRNYVRMQPQPNNVSKGDVYIMEGDQIVAVCEGIKFQQIPRRVLNTFLPPNKGSGPASAAKPAAAPVAAARPAPAAAPIKTAPAPAKAAPAPAPAAPKAAPKPKKAAAPKKAAGGLTAKVMKILAKETEVDEGELVDEAQFENLGVDSLLSLTISAVFREELDMDISSTLFTDYPTVGDMKKYFAQFDNGSSTSSSAEEEDSDEDSIPPTDAATPMDDLSTPASSVPSSAPSDAGKPDSPTRETLDDVGDVSLAKHIVAQEMGVDIAEVTDDADLAEMGMDSLMSLTILGELREKTGIDLPSTFLTTNPTMKDIDNALGMRPKPKAAPKPAAPKAAAPSSSKKTDMNEVSARLSALNNNTDISRYPNATSVLLQGNPKQATKKIFFLPDGSGSATSYVSIPNLGPDVCAYGLNCPFMKNPEQWQCGIEISAFVYLAEIKRRQPQGPYIIGGWSAGGVIAYSVAQALLAANEGVEKLLLLDSPCPVNLAPLPARLHNFFNQIGLLGTGDPAKTPKWLLPHFSAAIRSLSDYDPKPSLRPIPTYAIWCREGVAGNPGDPRPPPAEEEDPAPMTWLLEHRTNFKDNGWAQLCGDSMKFGVMGGHHFSMMKPPHADDLGNLIREGLDWQP

>PKSI_1_177

MSNVLLFGDQTAEQYQLLNKIVLRKENALVITFVERCAKALREETNALPRSQRNAVPDFLTVNDLKEAYHQKGVKVPMVESALVTIAQIGHYIGYFSEHSAEQPSATNTRALGLCTGLLAAAAVVASKTVEELVLVGVEFVRLSFRSGAAVDAARTALCQTGDDNAPWSTIVTGTTEASAKEALAKFHEEKGIPQTSHAYISAVSVMAITVSGPPTTVKRFFEESPALSKNHRVPIPVYGPYHAEHLFGETEINKIASASILEGLKQHQPVSLVHSAATGKALVAENAAELAKLVLAEMLQLPVRWDHLLEEAVSQITSKKAPAKIWAMGVSNVANSLVSALKAGGQTDVSTVDQSTWTENEPDTHGRTQNDKVAIVGMAGRFPNSADHEALWELLMKGLDVHRRIPKDRFDADTHVDPSGKGKNKSHTPFGCFIDEPGFFDPRFFNMSPREAAQTDPMGRLALVTAYEALEMSGYVPNRTPSTKLHRIGTFYGQTSDDWREINAAENVDTYFITGGVRAFAPGRINYYFKFSGPSYSVDTACSSSLAAIQLACTSLWAGDCDTACAGGLNVLTNPDIFSGLSKGQFLSKTGSCKTYDNNADGYCRGDAVGTVILKRYEDAIADKDNILGCILGAATNHSAEAVSITHPHAGAQEFLYKRVLANAGVDAHEISYVEMHGTGTQAGDGIEMTSVTNVFAPRHRQRRDDQPVYLGAIKANVGHAEAASGINSLAKVLLMMKHNKIPANVGIKGEMNKTFPADLKDRKVNISQKAVDWPRNGKEKRKVFLNNFSAAGGNTALLLEDGPAYEAPTATDPRGTVPVTVTARSISALKRNIANLQKYVSENPSTTLTSMSYTLTARRIQHNYRVAFPLDQINKFSDALQAQVKESYSPVPNAPTRVAFCFTGQGSQYTGLGQKLYNDLKSFRDDIDQLDHLARVQGLPSFLEIVQGADVQTLSPVKVQLGMACIQVALGRMWAAWGITPAAVIGHSLGEYAALHVAGVISASDMVLLVGRRAELLVRDCTPHTHGMLAVKGGAEAIRNTLGNKMTEIACINGPEETVLCGSGDVVGAANEVLAAKGFKATKLNVPFAFHSAQVDPILEQFKKIAASVTYNKPSVPVLSPLEGDIIREAGKINPEYLARHARETVNFWTALTAGQKEKVFDEKTAWLEVGAHPVCSGMVKASIGATTTAPSLRRGEDAWKTISNSMCTLFTAGVNFNFDEFHKEFNDAQEMYTLPTYSFDNKKYWLDYHNDWTLRKGEPAQTKEVIVEKPVASASAPAVEMPAKRLSTSCQRVIAENFSGNNGSVTVQSSLADPKLYPVVCGHMVNNAALCPSSLYADMALTISDYIWKQMRPGTETPGYNVCNMEVPKPLIAQIPQPAEGQHIQLEANADLDSGIVKLNFRSVKPDGQKLQDHAHCIVRLEDKAAWEDEWSRYNYMVQAQMELLQHKTLNGGAHKVQRGMAYKLFKALVNYDEKYRAMAEVVLASGQTEASAMLDFPTKPEDGDFYCPPYHIDGSCHISGFIVNASDLLDSEQNVYVSHGWGAMKFSRPLTAGMKLRNYVRMQPQPNNVSKGDVYIMEGDQIVAVCEGIKFQQIPRRVLNTFLPPNKGSGPASAAKPAAAPVAAARPAPAAAPIKTSPAPAKAAPAPAPAAPKAAPKPKKAAAPKKPAGGLTAKVMKILAKETEVDEGELVDEAQFENLGVDSLLSLTISAVFREELDMDISSTLFTDYPTVGDMKKYFAQFDNGSSTSSSAEEEDSDEDSIPPTDAATPMDDLSTPASSVPSSAPSDAGKPDSPTRETLEDVGDVSLAKHIVAQEMGVDIAEVTDDADLAEMGMDSLMSLTILGELREKTGIDLPSTFLTTNPTMKDIDNALGMRPKPKAAPKPAAPKAAAPSSSKKTDMNEVSARLSALNNNTDISRYPNATSVLLQGNPKQATKKIFFLPDGSGSATSYVSIPNLGPDVCAYGLNCPFMKNPEQWQCGIEISALVYLAEIKRRQPQGPYIIGGWSAGGVIAYSVAQALLAANEGVEKLLLLDSPCPVNLAPLPARLHNFFNQIGLLGTGDPAKTPKWLLPHFSAAIRSLSDYDPKPSLRPIPTYAIWCREGVAGNPGDPRPPPAEEEDPAPMTWLLEHRTNFKDNGWAQLCGDSMKFGVMGGHHFSMMKPPHADDLGNLIREGLDWQP

>PKSI_2_Hyb1_177

MSRGTDGAALPTASTVSTQSGGPHVIDPTTKDNGPSPVPASKGPTGAHPKNGDIQSEYPSHAVAVVGMAGRFPGAKSVDALWDLLEAGKSTVEPAPMERIGLGHLPPDDPSRMWWGNFLDDVDAFDHEFFRITAREAQTWDPQQRIMLEVAYEALEDGGQFGASLPSRNRDFGCYIGAVMNNYYDNVACHKASAYATKGTSRCYISGAVSHFFGWTGPAITLDTACSSSMVAVHAACKAIIAGECSRAIAGGTNVITSPHDYRNLAAAGFLSPTGQCKPFDSDADGYCRAEGVGVVVLKSLATAIEENDRVLGVIVGSAVSQSGNVGHIAVPDTRAQVLLHRKALSIANLSPEEVSYVEAHGTGTKVGDPIEMAGLREVYCSSPRTSPLLVSSMKGNIGHTEATAGVAGLIKVLLMMNRRSIPPQASHKRLTPRIPALHPDIVAIPRHLAPWPGTARAACVASYGAAGSNAAILVREMSTRHAGGEREGTAQPKIQGEQPLFVSAATRDMLSLRCSEILLWLRKQKDYRLELSDVLFNISIRANHALPYTFCTTVSSMDGLELKLAEVAKESMVLRKIGSAKPTILVLGGQQGQYVGLSWLCYQEWQGLRRNLDQCHESLIDMGHGGLYPDIFQKAPLVDLKAFHAALFASQYSSAKAWIDSGLKVDAVVGHSFGQLAALCICGVLSLADGLKLVTGRASLIEKHWGSERGAMLSLSAPTRVVDRLLEALEAKIDYAQVACYNGPENHVVVGSEHAIKVLERHIDSRSDLRGSVVAQRLDVTLGFHSVYTEAVQPALEKLASKLEWKAPTIHLETCDTEASCAVFNHSFVVHHMRRPVHFQQAVERLSLRYPESCWISAGHAASFLRLAQKSLRGRGHHTFICLDKVFSEASGSLARAIVELWNEGQPVQYEPFERSRQHRYPHLTPPPYPFARGRHWLHFTNPTTSQPSAICPASQQPQDEDDFLSLVGGDAQGGAEFFIPAAHPRFQLLCNGHVMSGEALAPASFYLEVAARAALKLRCDIKSGQLGLRVEKLSMNAPVGNDPRNIIRLRLEQHEGPRDSWLFSMTMERSPRADTDDINPQKQVVGVVSLRSQDQTNAVAGFKDFRISDVRRRYQEIVDDESAERMQGNHVYRAFDSVVTYGEIFRRIKSIACVGTEAAGEIEWNRRCEDTHVAAATDTQFIDGFLQAAGFLVNYFNNDDFQSSLFICQKIDQIDFADAIPTDAHNLSFYSRMRMDSEDAAIADVLVLDARQQQVLFAASGLQFKKLSRVPLARSLRAANAVAPCSTDGLRKQIHTGRETASRDGSESGHRPDHRRKVLEVLANVTGTPPECVSMAGPLEDLGIDSLAAIEVLNDLRASLDLVIPLSTLMSFNSIGAFVEYVTSTVGSRDRRHRDPHDVLKDARLAKKVTSTAVGYEQVARDEENQVSVASSEPKHDMSLDVDSSAVDLDYLAAVYPSHTRLFLAYIVEAFGGIGCDLATMAAGEAMPEMQNILPRHKRLLHRFFLALVEARILIETPEGTFVRTEQHVDKARAADLHSAAMRHWPEHGSLHQLMHAVGSRTAECLTGVFDGLQIVFGDQANETALREFYQHWPLFKGATKSLAKIVLRVAGHLGRTGKIQILEVGAGTGGTTRPLLAALQAAEMPFDYHFTDVSPHLVNAARQSLDGIAGLTFGVLDVEEQPPKDLEGAYDVVLASNCIHATRDLEQSLGHLRCLLREDGIFAMVEITRRLNIFDLVFGLFDGWWRFNDNRTHCLADEHHWSSKLMAAGFQDVMWTDVGNPKAQVVRVATASPGTTPRREASKRVAKTRVCEVVYKTSSELDILADVYCPARADPAVTLPIALMIHGGSHVLFSRKDVRPAQTRTLLGMGFLPVSIDHRLCPETKLVEGPMVDVCDALDWARHTLPAIDLGDCSLKPDGGCVVAVGWSSGGQLAMSLAWTAPQRGLEPPTAILALYCPTDYEDDWWQHPIQPCGAEDVDEEYDLLDAVQPSPISSYDSICSWEPLTDVRIRQDPRCRIVLHMNWKAQTLPIIVSGLPCSSSVSDEDSSRIDWTALPQPPSREIQRCSALAQIKLGNYRTPTFLVHGTKDELIPWQQSHRTWKCLVSHGIDAKLALIKDAPHICDASKDHESAGWKAVLRGYGFLADKVRIRGHCSALEGRALPE

>PKSI_3_Hybr1_177

MAKIACPQGKPDDIAVIGLAYRFPGGADSDAKLWDLLAERKSAHSKIPVDRFNIDAFHRLGATHSDNVAADGAHLLEQDISAFDAPFFGITTEEAKVIDPQARLLLECSYEALMNAGLKLESISGSDTGCYVGCFDLDYHQMLMGDFECAPKYSGTGTAFSLLSNRLSWFYNLKGPSLSLDTACSSSLVGLHLACQSLSAGESSMAMVCGAKLLLGPHLSMWLSRLNMFSSDGKSRSFADDTAGYGRGEGIATVILKPLADALRDNDPIRAVIKGTGVNQDGHTKGITVPNPESQLDLIRSTYLAAELSFADTGYFEAHGTGTAVGDPLELAAVAQAIVESRRSEPLYVGSIKSNIGHTEGAAGLAGLIKCILMLEKGVILPNIHFDRPNKRIPFERDGIKVPTEVLPWPKDLDRRASINSFGFGGTNAHAIVESFPTPSISSLDHVAQVDASEVVAVTPNHPRLFVLSGHEPAAVEKLRQRYVEYIIQAKNSDGVDRKLDDLSYTLGCRRSRMDWSVFHVASDFEELGKKLLENTTALKRAARSPRIGFIFTGQGAQWPRMGAGLMRYGVFQESVQAADRFLSGQCHCGWSVIDELGKAKDESRIASSELAQPICTIIQVAMIDLLRCWNIRPTAVAGHSSGEIAAAYCTGAITRQSAWEIAFHRGRECARLKEVAPDLRGAMLAVGLGVEDVRPYLDAVAPDRVNIACINSPNSTTLSGDAVEIQELLTKLIANGVSARELRVVNAYHSHHMKLVADRYLDSIAHVAVQTEVVRSDVALLSSVTGTSASSSDLDPEYWVRNLVSPVLFSDAVSAMLKGTKKTFRRQQGTAEPAVDVLLEIGPHAALRGPLADILKSEAVESVAYVPTLNRGSDALESATAAAGELWSRGCPVDVTAVNNHPRQPRVLADLPSYPWDRLNKYWATSRVTHDVLHRAFPHHDILGKRLTGSDALAPAWRHFLRFSESPWVREHVVHGSIVFPGAGFLAMAIEAALQLVEKDRKLANVRMRDVHISKALVLEQEEEGAQELVTRFHRVDDRSDGTWSGWWEFSISCTKSRVEPERHISGQIMLEYRPMEPSSQPASDVVHQAQKVKYGKLVGSSADRLYRAAFYEASEAAGLAYGSQFQGVVDVARSGDGRCCWRVQLPDRKAPASESKHLIHPTTLDAIVHAMFGAMNKGSALSSAALPIAFDKVVISADMPTDAGTCLSGFTVTDVKETTREVTADVYVSSDDWARRLLQIEGLRCTELPSPEGVHNDNETQSAPLGSVAWQPDLDLLDNDGLKSYIMKNPERDQLGATDGRVAALSTRLRSAVAQILYLAAFKTPTLSILQVGGFSEGLTDTFVATIASDTNVPAVTAEILVLDSEEENIPRIRERQYDGCSVAASHWNRAGSLVSHIPKGDKFDAVLLVVPEHNDELATRVLVAQARNLLKAGALLIVLDTIQNTRANLAFYGFETAQTSSSGRLRPWQSLNVEDPDCGKAAALCRESPATISREITGAPICVLKPPHCTKETVDVISALEGILNDARLEVAFQEWPPSVAQVEGKFVISLLDTEASFLSNIDAADFEVLREVALQSRRLLWVCSGDDPHMAIALGWLRVLQNENANRVYQHLTLTRKKAAAPQNHSCAIARLALTQTREREFAEEDGNLYIPRWYHEEGLSGTLEGREMSVQLERAPLGVAKSSMPLRILHGKDAESARFVPDSPSVSRLAADEVEIEMQYIVLTDSDISPVGNTAHREGSGVVRAVGRDVTRLGLDDQVCVSYVGPLSTRVIAKEVYCQVVPPDAIMEEAACIPNTFATALRVLTDVARVKPGQTVLVQTAGTKIGRAAVLLALALDAVVYATAREKTETDKIVSLGVRLQNVFTEGDMDLPQAAKTLAGERGLDVVLRTSKNSSAPCLLPHCVAQHGTLVDVHAASEPGPDVPSCEDTISVMGVGPLLPEDPVSLQKSASRAANYLPQLSGLAASFDLFESARISDALECQQEQGTRGGVILSLDDADMVPIAPSVNRKLYLCEQATYVLAGGLGGLGQSLARLLVDHGARNLALLSRGGPNSPSAETFIKEMAEVGVAVKVLACDIGDESSMKAALDDCAGTMPPIRGVIQAATVYRDAIFDNFTFEDWQANLRAKVQGSWNLHRHLPKDIDFFVMLGSVAGLMGHVSQAGYAAGNTFQDALAHYRRSRSLPAVTIDLGPMLDVGAVNDGTVSASFSTSEATWMTEADLHAIMIMCISGEITSCNFPPQLCTGLPSGGMLQLGQHEMPEHYDRPIFALLKGLGVSAAAGKDNKVLSRRSKDFSHQLPAVTSMEEARKCVVNAVKAHLAKGLGRSADSIKSSQPLDSYGIDSLGAMGFRNWVRETMKADVSMFDVLNARSIKELAAKIVRISELIPEGLESERVTREDE

>PKSI_4_Hybr2_177

MAKVERKSWRPEDVAVIGLACRFSGSASNEAKLWELLEKRESAHSKVPGERYNVEAFHQQGSTNSNNLAADGGHYLEQDVKSFDAPFFNIITKEAKAMDPQARMLLESSYEALENAGLSLESVRGSDTGCYVGCFNRDYYELLMADAEDSPEYSVTGTGFSLLANRLSWYYDLRGPSKSEDTACSSSLVALDSAYKSLLRGESKMAMVCGANLMLSPNIGLWLSKLNMLSSEGLSRSFAEGVSGYGRGEGIATVILKPLADALRDGDTIRAVINATGVNQDGHTKGITVPNSQAQSTLIESTYRRAGLDFADTGYFEAHGTGTAVGDPLEIEALERVIKHAKRTSPLHVGSIKSSIGHLEGAAGLAGLIKCILMLEKGVILPNLHFERPNRKISFENIVVPTTAVPWPEGVKRRASVNLFGYGGTNAHVIVEAFVVPCPQSRDDDLRNAQDDGLERPSPQRLFVLTGREQSTVRKMRLRYASYIQTMKDTASPSVKFDDLSYTLGKRRSRLDWAEAHVASDFKELEEKLSAPEVAATRSSNKTRLGFVFTGQGAQWPRMGLELMRYTAFRDSVEAADEYLTQKLDCSWSVIEELEKHGDESRVTSSELGQPLCTIIQVAMVDLLGSWNVRPTAVVGHSSGEIAAAYCTGVMTKQAAWQIAFHRGRECAKLKGKAPELEGSMLAVGLDVESIRPYFNNLQSGRINVACVNSPNSITISGDASEIRKLQAMLVADSVFARELPVENAYHSHHMELVAESYLHSISDVDIQHSVALSDITMVSSVTGQSIEPSELTPEYWVRNLVSPVLFADAVTAMLRGSRRRFRRGAKAEPAVDFLLELGPHATLQTPLSDIVKAQAQEDVKYASMLLRGENAVDSAMTTAGKLYCHGCPVDVTAVNDIRHECRVLVDLPAYPWNRSTKYWGVSRLMQGYLHRTHGYHSLLGARLIGSDALNPAWRHFLNLDDSPWIEEHVVHGAVVYPGAGFLSMAIEAALQLAQPGREIANVRLQNVRVLKALVIKEGEDDPEVITRFRQADSVSDEASSMRWAFEISCAKGHDERDNRATGQDEFERHATGQITLDYQPEHPYLSPLSEQIHDVRRGEYARLADTCVDTMKQDGFYEASKDVGLAYGHDFQGIATMARGPNSCCWDLRVTERSTSLPGAYESKHLIHPTTLDAIVHSLFGAMNGGKKFQNAALPVAFDSIMISPATLTTSGTKLSGFTVIREAKEREIVADIHVSSEDWSQPLVQIAGLRCTQMPSPESELHDQDARPSPVGTITYRPDIALFDENGLMNYLNERQKPESCAKVHPDLYSERLRNAVAQVVELAMFKDPGLSVLQIGGYDRGVTDSLLFTLKAESAGQALSSKIVLLDPSQETLVEIRQQYDAESSIVDALHFAVDKPLPSEVSREYGFDVVLVAIDHGLDDMSKQNVLAEAQNVLKAGGIFVVFDTLRAITERQYQHLALGEATERSPFDLALAIAKVAVVQTREGEFVEQDGSFNIPRWSYGPEMTRTIADSTVSLESDSVRLGDMAQGTPLRMLHAGHPKYAHFVTNTIQPLHLAAGQVKVELRFVDITHQDLVEPDMRALREASGVIKAVGSDVSLLRPGDNVCLSFVGHLSTSVNVDEALCQRIPPGVNMAEAACIPITLATALRALVGVAGVKPQHNVLVQAGGTKMGRAAILIASAANAVVYTTARDAEEVESLLALGISKQNIVPEGDPLLPTVTKILTGNRGWDVIVRTTKIVAETFILPECVADFGVVLDVFPSSGTGCTQETTISVMGIGSLLPEDPVLMQKTVSRIMDYLPQVSTLANSFDVFPSSAIPAALDRHGAQNQHRGVMLSFDQEDLVRVSPSATNTMKLYRDATYVMAGGLGGLGRSIARLLVDNGARNLVFLSRSGPNTTAATTMMSKMAGLGVTVKTLKCDVGDEKSVAAALDECSSMPPVRGVIQAAADIQDAIFDTYTFEQWQANLRPKVQGSWNLHCQLPEDMDFFVMLSSISGLIGHEGQAGYAAGNTFQDSLALFRHSRGLPAVTIDLGAMLDVGTIAEGSTTATFRSSDAVLMKAIDLHEIMTMCISNEINGYAIPAQVCTGLPSGGMLQVEQQEIPSYFHKPLFAALKCLGTSAVSAVNVAAPVEGVIDFAAQLTTVGSLDEADCVIANILRAHIAKAVQRAVDDIDLSQPLYSYGIDSLMAVELRAWIGEKMKADLTLFDILNAESIQALALKISKTSQLVQQEIRGMN

>PKSI_1_157

MSNVLLFGDQTAEQYPLLNKIVLRKENALVITFVERCAKALREETNALPRSQRNAVPDFLTVNDLKEAYHQKGVKVPMVESALVTIAQIGHYIGYFSEHSAEQPSATNTRALGLCTGLLAAAAVVASKTVEELVLVGVEFVRLSFRSGAAVDAARTALCQTGDDNAPWSTIVTGTTEAAAKEALAKFHEEKGIPQTSHAYISAVSVMAITVSGPPTTVKRFFEESPALSKNHRVPIPVYGPYHAEHLFGETEINKIASASILEGLKQHQPVSLVHSAATGKALVAENAAELAKLVLAEMLQHPVRWDHLLEEAVSQITSKKAPAKIWAMGVSNVANSLVSALKAGGQTDVSTVDQSTWTENEPDTHGRTQNDKVAIVGMAGRFPNSADHEALWELLMKGLDVHRRIPKDRFDADTHVDPSGKGKNKSHTPFGCFIDEPGFFDPRFFNMSPREAAQTDPMGRLALVTAYEALEMSGYVPNRTPSTKLHRIGTFYGQTSDDWREINAAENVDTYFITGGVRAFAPGRINYYFKFSGPSYSVDTACSSSLAAIQLACTSLWAGDCDTACAGGLNVLTNPDIFSGLSKGQFLSKTGSCKTYDNNADGYCRGDAVGTVILKRYEDAIADKDNILGCILGAATNHSAEAVSITHPHAGAQEFLYKRVLANAGVDAHEISYVEMHGTGTQAGDGIEMTSVTNVFAPRHRQRRDDQPVYLGAIKANVGHAEAASGINSLAKVLLMMKHNKIPANVGIKGEMNKTFPADLKDRKVNISQKAVDWPRNGKEKRKVFLNNFSAAGGNTALLLEDGPAYEAPTATDPRGTVPVTVTARSISALKRNIANLQKYVSENPSTTLTSMSYTLTARRIQHNYRVAFPLDQINKFSDALQAQVKESYSPVPNAPTRVAFCFTGQGSQYTGLGQKLYNDLKSFRDDIDQLDHLARVQGLPSFLEIVQGADVQTLSPVKVQLGMACIQVALARMWAAWGITPAAVIGHSLGEYAALHVAGVISASDMVLLVGRRAELLVRDCTPHTHGMLAVKGGAEAIRNTLGSKMTEIACINGPEETVLCGSGDVVGAANETLAAKGFKATKLNVPFAFHSAQVDPILEQFKKIAASVTYNKPAVPVLSPLEGDIIREAGKINPEYLARHARETVNFWTALTAGQKEKVFDEKTAWLEVGAHPVCSGMVKASIGATTTAPSLRRGEDAWKTISNSMCTLFTAGVNFNFDEFHKEFNDAQEMYTLPTYSFDNKKYWLDYHNDWTLRKGEPAQTKEVIVEKPVASASAPAVEMPAKRLSTSCQRVIAENFSGNNGSVTVQSSLADPKLYPVVCGHMVNNAALCPSSLYADMALTISDYIWKQMRPGTETPGYNVCNMEVPKPLIAQIPQPAEGQHIQLEANADLDSGIVKLNFRSVKPDGQKLQDHAHCIVRLEDRAAWEDEWSRYNYMVQAQMELLQHKTLNGGAHKVQRGMAYKLFKALVNYDEKYRAMAEVVLASGQTEASAVLDFPTKPEDGDFYCPPYHIDGSCHISGFIVNASDLLDSEQNVYVSHGWGAMKFSRPLTAGMKLRNYVRMQPQPNNVSKGDVYIMEGDQIVAVCEGIKFQQIPRRVLNTFLPPNKGSGPASAAKPAAAPVAAARPAPAAAPIKTAPAPAKAAPAPAPAAPKAAPKPKKAAAPKKAAGGLTAKVMKILAKETEVDEGELVDEAQFENLGVDSLLSLTISAVFREELDMDISSTLFTDYPTVGDMKKYFAQFDNGSSTSSSTEEEDSDEDSIPPTDAATPMDDLSTPASSVGSSAPSDAGKPDSPTRETLEDVGDVSLAKHIVAQEMGVDIAEVTDDADLAEMGMDSLMSLTILGELREKTGIDLPSTFLTTNPTMKDIDNALGMRPKPKAAPKPAAPKAAAPSSSKKTDMNEVSARLSALNNNTDISRYPNATSVLLQGNPKQATKKIFFLPDGSGSATSYVSIPNLGPDVCAYGLNCPFMKNPEQWQCGIEISALVYLAEIKRRQPQGPYIIGGWSAGGVIAYSVAQALLAANEGVEKLLLLDSPCPVNLAPLPARLHNFFNEIGLLGTGDPAKTPKWLLPHFSAAIRSLSDYDPKPSLRPIPTYAIWCREGVAGNPGDPRPPPAEEEDPAPMTWLLEHRTNFKDNGWAQLCGDSMKFGVMGGHHFSMMKPPHADDLGNLIREGLDWQP

>PKSI_2_157

MPDNVSFMDESQDLRHIRETSRTSHTTSILNVEGDYRNNDMPREPGECNRSTNGTVDHERMTPSADGGIPIAICGIGLRLPGGIRNDRDLYDSLYNKKDARGVIPEDRFSIDSFHSAHGKTGTIITKHGYFLQDIDLTKFDVNMFNMTPAEVERLDPHQRILLETVRETLESAGEASFRGKKVGTYVGNFTDDWLDLQNVDTVDFATYQLHGKMDFSLANRISYEYDLRGPSMTIKTACSSSALAIHEAVYSIRNGECDAAIVSGSNLNLAPRLWVGMSSQGAISPDGSSKTFDESANGYARGDGIAALFIKRLDDAVRDGNPVRAVIRSTASNADGRTPGMTMPSTEAQEALIRRAYDAANLPLSETAMVECHGTGTAVGDPMEANAVARCFGDQGMLIGSVKPNLGHSEGASAITSVVKAVLSLENRTILPNIKFHRPNPAIPWSEAKLTVPVEPLAWPKDRQERISVNSFGIGGSNVHVVLDSAASMGFRPRSLAPSKDDRPGRLLLFSGGHRASVEQSSSQHQDYVTKYPNRLSDVAYTLAKRPPVKCPGLTRSTFVFTGQGAQWLHMGKELLHESPVFAKSIGRMDSVIHSLKHAPQWTLEGIINDPENPSALTNAEISQPLCTAVQIGLVDLLKSWAIYPHAVLGHSSGEIGAAYASGVVDRAEAILLAFYRGYVCRFAQKAGGMAAVGLEKSQVIKYLQPGVCVACENSGSSVTLSGDLETLEEVLQSIRAENLNAFARKLQVGIAYHSDHMKALGGLYHQYITEHLDPKDPQVPFFSSVSGRALHSKDDFGATYWQDNLENPVLFHTAVLKSLEHTGDKQVHLEVGPHGALNGPLRQIYAETGSKARYVALQKRGANCFDTFLEGIGQLYCNGVPLQYPESADDRTLIDLPPYPWHYDHSYWSETRVMKNWRFRRQLPHDLLGLRTLDCSDAEPMWRNILRITDLPWLRDHCVGKDVVFPASGYMCMAGEAVFQETGCRDYTLREVDISTAMVLSSDHSTELLTTMKKRRLNAFLDSRWYEFLIMSYDGASWTQHCSGLVTNGPSTSHPKALLQTYDRPVSTNRWYTAMSKIGLNYGPRFTGLQNITTHVQEKKASMTIMDKQEDYESPYALHPSTLDLILQSWTVASVRGEYRRFTQLFLPTFVDEFYIGNSASKLIHLNTTAIGPDGSARGEAIGKDNDGQISFNLKGFKGSKLDNVGVDQPQEMQTIMKQQWKRDFDFADTAQLMRPAFDSTSELSLLERMFVLAAIEVHVRTSGMEGKLPHHQRYKLWIDAQIRRFGEPGYPMVEDSMELLRLDSRERQRQLCRLLEHSRKTTAHPVAEAIWRALDRIEDVFDGRIEYLDLLFNDGLMPKFYDWSNSLSDVSRLFRLLSHKKPQLKILEVGAGTGGSTARLLQYLQSDFGERQYHSYTYTDVSSGFFVQAQERFKDYEGMKYRVLDISQDPFEQGFGADEFDLICASNVLHATPRLTETLRNCRKMLRPDGILFLQELCPRQQFMGFIMGLFEGWWLGAEDGRADTPLLLPPAWDRRLRDVGFEGVEAFSFDNNPPYFMAANMTARPTVTAKSKGSITLLTFNDFLDDVAGALKQAIRAAGFEIDHCVWGEHVPLDQSLISLVDLERDKPLLQDIGDDDLRIFLDLIDAVLQTTVIWLNKPAQVSSADPNAAQMLGLARTLRAELAMHFATVEMADPTLNGMSAVVHLMCMLQRGSALPENSLDQDMEYVWAHDAMHVSRFHFQPVDEALMEMSPKFDVKTLVPLQRGMLSSLKWIGTRMLPLAEAEVQIRMSAVGMNFHDMMIAMNMFDSPLTLGSGYNSIGMEGVGYVTRKSPEVDHVQVGDRVIVIGSNSSGFATDVHRPADYCIKCPSSLTDVEAAGMSFAYMTVLWSFLDKGGLRKGQSVLIHSAAGGVGIAALHVSRWLGLEAFVTVGNEEKVRFIMNNFGLPRNRIFNSHSATFLDDVMDATAGRGVDAALSAAAGELLHNTWSCIAPGGVMLEIGKRDLINRGRLSLAPFEENRSYVGIDFSRLTIVNKPAVVRLLRQTMRLVEQGHIHPIHPTTCFDAEHAEDAFRFMQTGQHIGRIVVKIPRDTSTIPLASRPPAPEFEGQKTYLLVGGMGGLGRSVASWMVSAGARNLIFMSRSAGKSEQDQNFARELELSGCKVHCCAIDITDTDAVREALERTQASVAGVLQMAMVLRDVGIMNMDKANWDAAVAPKVQGTWNLHHALPNVDFFVMFGSNSGTLGSYGQANYAAANAFLDSFVQYRQNLGQAASVIDIGAVGDVGYVAETQVAAENMESMAGRLISEQDFLNCLQLAIARSTPTERRNKGPSTEADGYIDLKQVILLNSSILSMADPSNQIFWRKDPRMGIYRNVQRTSAESPTTDSNSLRRLVATLKADSSIADQAELAQTLARELSKQVATVLMLGEDEIDVDRSLTDVGMDSLVAIEMRNWWKQNLGVDVSVLELKDGRSILRLGELAATRLKERYSRDS

>PKSI_1_154

MSNVLLFGDQTAEQYPLLNKIVLRKENALVITFVERCAKALREETNALPRSQRNAVPDFLTVNDLKEAYHQKGVKVPMVESALVTIAQIGHYIGYFSEHSAEQPSATNTRALGLCTGLLAAAAVVASKTVEELVLVGVDFVRLSFRSGAAVDAARTALCQTGDDNAPWSTIVTGTTEASAKEALAKFHEEKGIPQTSHAYISAVSVMAITVSGPPTTVKRFFEESPALSKNHRVPIPVYGPYHAEHLFGETEINKIASDSILEGLKQHQPVSLVHSAATGKALVAENAAELAKLVLAEMLQHSVRWDHLLEEAVSQITSKKAPAKIWAMGVSNVANSLVSALKAGGQSDVSTVDQSSWTENEPDTHGRTQNDKVAIVGMAGRFPNSADHEALWDLLMKGLDVHRRIPKDRFDADTHVDPSGKGKNKSHTPFGCFIDEPGFFDPRFFNMSPREAAQTDPMGRLALVTAYEALEMSGYVPNRTPSTKLHRIGTFYGQTSDDWREINAAENVDTYFITGGVRAFAPGRINYYFKFSGPSYSVDTACSSSLAAIQLACTSLWAGDCDTACAGGLNVLTNPDIFSGLSKGQFLSKTGSCKTYDNNADGYCRGDAVGTVILKRYEDAIADKDNILGCILGAATNHSAEAVSITHPHAGAQEFLYKRVLANAGVDAHEISYVEMHGTGTQAGDGIEMTSVTNVFAPRHRQRRDDQPVYLGAIKANVGHAEAASGINSLAKVLMMMKHNKIPANVGIKGEMNKTFPADLKDRKVNISQKAVEWPRNGKEKRKVFLNNFSAAGGNTALLLEDGPAYEAPTATDPRGTVPVTVTARSISALKRNIANLQKYVSENPSTTLTSMSYTLTARRIQHNYRVAFPLDQIDKFSDALQAQVKESYSPVPNVPTRVAFCFTGQGSQYTGLGQKLYNDLKSFRDDIDQLDHLARVQGLPSFLEIVQGADVQTLSPVKVQLGMACIQVALARMWAAWGITPAAVIGHSLGEYAALHVAGVISASDMVLLVGRRAELLVRDCTPHTHGMLAVKGGAEAIRNTLGNKMTEIACINGPEETVLCGSGDVVGAANETLAAKGFKATKLNVPFAFHSAQVDPILEQFKKIAASVTYNKPAVPVLSPLEGDIIREAGKINPEYLARHARETVNFWTALTAGQKEKVFDEKTAWLEVGAHPVCSGMVKASIGATTTAPSLRRGEDAWKTISNSMCTLFTAGVNFNFDEFHKEFNDAQEMYTLPTYSFDNKKYWLDYHNDWTLRKGEPAQTKEVIVEKPVASASAPAVEMPAKRLSTSCQRVIAENFSGNNGSVTVQSSLADPKLYPVVCGHMVNNAALCPSSLYADMALTISDYIWKQMRPGTETPGYNVCNME

>PKSI_1_20

MSNVLLFGDQTAEQYPLLNKIVLRKEDALVITFIERCAKALREETNALPRSQRNAVPDFLTVNDLKEAYHQKGVKVPMVESALVTIAQIGHYIGYFSEHSAEQPSATNTRALGLCTGLLAAAAVVASKTVEELVLVGVEFVRLSFRSGAAVDAARTALCQTGDDNSPWSTIVTGTTEAAAKEALAKFHEEKSIPQTSHAYISAVSVMAITVSGPPTTVKRFFEVSSALSKNHRVPIPVYGPYHAEHLFGETEINKIASASILEGLKQHQPVSLVHSAATGKALVAENAAELAKLVLAEMLQHPVRWDHLLEEAVSQITSKKAPAKIWAMGVSNVANSLVSALKAGGQTDVSTIDQSTWTENEPDTHGRTQNDKVAIVGMAGRFPNSADHEALWELLMKGLDVHRRIPKDRFDADTHVDPSGKGKNKSHTPFGCFIDEPGFFDPRFFNMSPREAAQTDPMGRLALVTAYEALEMSGYVPNRTPSTKLHRIGTFYGQTSDDWREINAAENVDTYFITGGVRAFAPGRINYYFKFSGPSYSVDTACSSSLAAIQLACTSLWAGDCDTACAGGLNVLTNPDIFSGLSKGQFLSKTGSCKTYDNNADGYCRGDAVGTVILKRYEDAIADKDNILGCILGAATNHSAEAVSITHPHAGAQEFLYKRVLANAGVDAHEISYVEMHGTGTQAGDGIEMTSVTNVFAPRHRQRRDDQPVYLGAIKANVGHAEAASGINSLAKVLLMMKHNKIPANVGIKGEMNKTFPADLKDRKVNISQKAVDWPRNGKEKRKVFLNNFSAAGGNTALLLEDGPAYDAPTATDPRGTVPVTVTARSISALKRNIANLQKYVSENPSTTLTSMSYTLTARRIQHNYRVAFPLDQINKFSDALQAQVKESYSPVPNAPTRVAFCFTGQGSQYTGLGQKLYNDLKSFRDDIDQLDHLARVQGLPSFLEIVQGADVQTLSPVKVQLGMACIQVALARMWAAWGITPAAVIGHSLGEYAALHVAGVISASDMVLLVGRRAELLVRDCTPHTHGMLAVKGGAEAIRNTLGSKMTEIACINGPEETVLCGSGDVVGAANETLAAKGFKATKLNVPFAFHSAQVDPILEQFKKIAASVTYNKPAVPVLSPLEGDIIREAGKINPEYLARHARETVNFWTALTAGQKEKVFDEKTAWLEVGAHPVCSGMVKASIGATTTAPSLRRGEDAWKTISNSMCTLFTAGVNFNFDEFHKEFNDAQEMYTLPTYSFDNKKYWLDYHNDWTLRKGEPAQTKEVIVEKPVASASAPAVEMPAKRLSTSCQRVIAENFSSNNGSVTVQSSLADPKLYPVVCGHMVNNAALCPSSLYADMALTISDYIWKQMRPGTETPGYNVCNME

>PKSI_1_152

MSNVLLFGDQTAEQYPLLNKIVLRKENALVITFVERCAKALREETNALPRSQRNAVPDFLTVNDLKEAYHQKGVKVPMVESALVTIAQIGHYIGYFSEHSAEQPSATNTRALGLCTGLLAAAAVVASKTVEELVLVGVDFVRLSFRSGAAVDAARTALCQTGDDNAPWSTIVTGTTEASAKEALAKFHEEKGIPQTSHAYISAVSVMAITVSGPPTTVKRFFEESPALSKNHRVPIPVYGPYHAEHLFGETEINKIASDSILEGLKQHQPVSLVHSAATGKALVAENAAELAKLVLAEMLQHSVRWDHLLEEAVSQITSKKAPAKIWAMGVSNVANSLVSALKAGGQSDVSTVDQSSWTENEPDTHGRTQNDKVAIVGMAGRFPNSADHEALWDLLMKGLDVHRRIPKDRFDADTHVDPSGKGKNKSHTPFGCFIDEPGFFDPRFFNMSPREAAQTDPMGRLALVTAYEALEMSGYVPNRTPSTKLHRIGTFYGQTSDDWREINAAENVDTYFITGGVRAFAPGRINYYFKFSGPSYSVDTACSSSLAAIQLACTSLWAGDCDTACAGGLNVLTNPDIFSGLSKGQFLSKTGSCKTYDNNADGYCRGDAVGTVILKRYEDAIADKDNILGCILGAATNHSAEAVSITHPHAGAQEFLYKRVLANAGVDAHEISYVEMHGTGTQAGDGIEMTSVTNVFAPRHRQRRDDQPVYLGAIKANVGHAEAASGINSLAKVLMMMKHNKIPANVGIKGEMNKTFPADLKDRKVNISQKAVEWPRNGKEKRKVFLNNFSAAGGNTALLLEDGPAYEAPTATDPRGTVPVTVTARSISALKRNIANLQKYVSENPSTTLTSMSYTLTARRIQHNYRVAFPLDQIDKFSDALQAQVKESYSPVPNVPTRVAFCFTGQGSQYTGLGQKLYNDLKSFRDDIDQLDHLARVQGLPSFLEIVQGADVQTLSPVKVQLGMACIQVALARMWAAWGITPAAVIGHSLGEYAALHVAGVISASDMVLLVGRRAELLVRDCTPHTHGMLAVKGGAEAIRNTLGNKMTEIACINGPEETVLCGSGDVVGAANETLAAKGFKATKLNVPFAFHSAQVDPILEQFKKIAASVTYNKPAVPVLSPLEGDIIREAGKINPEYLARHARETVNFWTALTAGQKEKVFDEKTAWLEVGAHPVCSGMVKASIGATTTAPSLRRGEDAWKTISNSMCTLFTAGVNFNFDEFHKEFNDAQEMYTLPTYSFDNKKYWLDYHNDWTLRKGEPAQTKEVIVEKPVASASAPAVEMPAKRLSTSCQRVIAENFSGNNGSVTVQSSLADPKLYPVVCGHMVNNAALCPSSLYADMALTISDYIWKQMRPGTETPGYNVCNME

>PKSI_1_12

MSNVLLFGDQTAEQYPLLNKIVLRKENALVITFVERCAKALREETNALPRSQRNAVPDFLTVNDLKEAYHQKGVKVPMVESALVTIAQIGHYIGYFSEHSAEQPSATNTRALGLCTGLLAAAAVVASKTVEELVLVGVDFVRLSFRSGAAVDAARTALCQTGDDNAPWSTIVTGTTEASAKEALAKFHEEKGIPQTSHAYISAVSVMAITVSGPPTTVKRFFEESPALSKNHRVPIPVYGPYHAEHLFGETEINKIASDSILEGLKQHQPVSLVHSAATGKALVAENAAELAKLVLAEMLQHSVRWDHLLEEAVSQITSKKAPAKIWAMGVSNVANSLVSALKAGGQSDVSTVDQSSWTENEPDTHGRTQNDKVAIVGMAGRFPNSADHEALWDLLMKGLDVHRRIPKDRFDADTHVDPSGKGKNKSHTPFGCFIDEPGFFDPRFFNMSPREAAQTDPMGRLALVTAYEALEMSGYVPNRTPSTKLHRIGTFYGQTSDDWREINAAENVDTYFITGGVRAFAPGRINYYFKFSGPSYSVDTACSSSLAAIQLACTSLWAGDCDTACAGGLNVLTNPDIFSGLSKGQFLSKTGSCKTYDNNADGYCRGDAVGTVILKRYEDAIADKDNILGCILGAATNHSAEAVSITHPHAGAQEFLYKRVLANAGVDAHEISYVEMHGTGTQAGDGIEMTSVTNVFAPRHRQRRDDQPVYLGAIKANVGHAEAASGINSLAKVLMMMKHNKIPANVGIKGEMNKTFPADLKDRKVNISQKAVEWPRNGKEKRKVFLNNFSAAGGNTALLLEDGPAYEAPTATDPRGTVPVTVTARSISALKRNIANLQKYVSENPSTTLTSMSYTLTARRIQHNYRVAFPLDQIDKFSDALQAQVKESYSPVPNVPTRVAFCFTGQGSQYTGLGQKLYNDLKSFRDDIDQLDHLARVQGLPSFLEIVQGADVQTLSPVKVQLGMACIQVALARMWAAWGITPAAVIGHSLGEYAALHVAGVISASDMVLLVGRRAELLVRDCTPHTHGMLAVKGGAEAIRNTLGNKMTEIACINGPEETVLCGSGDVVGAANETLAAKGFKATKLNVPFAFHSAQVDPILEQFKKIAASVTYNKPAVPVLSPLEGDIIREAGKINPEYLARHARETVNFWTALTAGQKEKVFDEKTAWLEVGAHPVCSGMVKASIGATTTAPSLRRGEDAWKTISNSMCTLFTAGVNFNFDEFHKEFNDAQEMYTLPTYSFDNKKYWLDYHNDWTLRKGEPAQTKEVIVEKPVASASAPAVEMPAKRLSTSCQRVIAENFSGNNGSVTVQSSLADPKLYPVVCGHMVNNAALCPSSLYADMALTISDYIWKQMRPGTETPGYNVCNMEVPKPLIAQIPQPAEGQHIQLEANADLDSGIVKLNFRSVKPDGQKLQDHAHCIVRLEDRAAWEDEWSRYNYMVQAQMELLQHKTLNGGAHKVQRGMAYKLFKALVNYDEKYRAMAEVVLASGQTEASAVLDFPTKPEDGDFYCPPYHIDGSCHISGFIVNASDLLDSEQNVYVSHGWGAMKFSRPLTAGMKLRNYVRMQPQPNNVSKGDVYIMEGDQIVAVCEGIKFQQIPRRVLNTFLPPNKGSGPASAAKPAAAPVAAARPAPAAAPIKTAPAPAKAAPAPAPAAPKAAPKPKKAAAPKKAAGGLTAKVMKILAKETEVDEGELVDEAQFENLGVDSLLSLTISAVFREELDMDISSTLFTDYPTVGDMKKYFAQFDNGSSTSSSAEEEDSDEDSIPPTDAATPRDDLSTPASSVPSSAPSDAGKPDSPTRETLEDVGDVSLAKHIVAQEMGVDIAEVTDDADLAEMGMDSLMSLTILGELREKTGIDLPSTFLTTNPTMKDIDNALGMRPKPKAAPKPAAPKAAAPSSSKKTDMNEVSARLSALNNNTDISRYPNATSVLLQGNPKQATKKIFFLPDGSGSATSYVSIPNLGPDVCAYGLNCPFMKNPEQWQCGIEISALVYLAEIKRRQPQGPYIIGGWSAGGVIAYSVAQALLAANEGVEKLLLLDSPCPVNLAPLPARLHNFFNEIGLLGTGDPAKTPKWLLPHFSAAIRSLSDYDPKPSLRPIPTYAIWCREGVAGNPGDPRPPPAEEEDPAPMTWLLEHRTNFKDNGWAQLCGDSMKFGVMGGHHFSMMKPPHADDLGNLIREGLDWQP

>PKSI_1_12620

MPDNVSFMDESQDLRHIRETSRTSHTTSILNVEGDYRNNDMPREPGECNRSTNGTVDHERMTPSADGGIPIAICGIGLRLPGGIRNDRDLYDSLYNKKDARGVIPEDRFSIDSFHSAHGKTGTIITKHGYFLQDIDLTKFDVNMFNMTPAEVERLDPHQRILLETVRETLESAGEASFRGKKVGTYVGNFTDDWLDLQNVDTVDFATYQLHGKMDFSLANRISYEYDLRGPSMTIKTACSSSALAIHEAVYSIRNGECDAAIVSGSNLNLAPRLWVGMSSQGAISPDGSSKTFDESANGYARGDGIAALFIKRLDDAVRDGNPVRAVIRSTASNADGRTPGMTMPSTEAQEALIRRAYDAANLPLSETAMVECHGTGTAVGDPMEANAVARCFGDQGMLIGSVKPNLGHSEGASAITSVVKAVLSLENRTILPNIKFHRPNPAIPWSEAKLTVPVEPLAWPKDRQERISVNSFGIGGSNVHVVLDSAASMGFRPRSLAPSKDDRPGRLLLFSGGHRASVEQSSSQHQDYVTKYPNRLSDVAYTLAKRPPVKCPGLTRSTFVFTGQGAQWLHMGKELLHESPVFAKSIGRMDSVIHSLKHAPQWTLEGIINDPENPSALTNAEISQPLCTAVQIGLVDLLKSWAIYPHAVLGHSSGEIGAAYASGVVDRAEAILLAFYRGYVCRFAQKAGGMAAVGLEKSQVIKYLQPGVCVACENSGSSVTLSGDLETLEEVLQSIRAENLNAFARKLQVGIAYHSDHMKALGGLYHQYITEHLDPKDPQVPFFSSVSGRALHSKDDFGATYWQDNLENPVLFHTAVLKSLEHTGDKQVHLEVGPHGALNGPLRQIYAETGSKARYVALQKRGANCFDTFLEGIGQLYCNGVPLQYPESADDRTLIDLPPYPWHYDHSYWSETRVMKNWRFRRQLPHDLLGLRTLDCSDAEPMWRNILRITDLPWLRDHCVGKDVVFPASGYMCMAGEAVFQETGCRDYTLREVDISTAMVLSSDHSTELLTTMKKRRLNAFLDSRWYEFLIMSYDGASWTQHCSGLVTNGPSTSHPKALLQTYDRPVSTNRWYTAMSKIGLNYGPRFTGLQNITTHVQEKKASMTIMDKQEDYESPYALHPSTLDLILQSWTVASVRGEYRRFTQLFLPTFVDEFYIGNSASKLIHLNTTAIGPDGSARGEAIGKDNDGQISFNLKGFKGSKLDNVGVDQPQEMQTIMKQQWKRDFDFADTAQLMRPAFDSTSELSLLERMFVLAAIEVHVRTSGMEGKLPHHQRYKLWIDAQIRRFGEPGYPMVEDSMELLRLDSRERQRQLCRLLEHSRKTTAHPVAEAIWRALDRIEDVFDGRIEYLDLLFNDGLMPKFYDWSNSLSDVSRLFRLLSHKKPQLKILEVGAGTGGSTARLLQYLQSDFGERQYHSYTYTDVSSGFFVQAQERFKDYEGMKYRVLDISQDPFEQGFGADEFDLICASNVLHATPRLTETLRNCRKMLRPDGILFLQELCPRQQFMGFIMGLFEGWWLGAEDGRADTPLLLPPAWDRRLRDVGFEGVEAFSFDNNPPYFMAANMTARPTVTAKSKGSITLLTFNDFLDDVAGALKQAIRAAGFEIDHCVWGEHVPLDQSLISLVDLERDKPLLQDIGDDDLRIFLDLIDAVLQTTVIWLNKPAQVSSADPNAAQMLGLARTLRAELAMHFATVEMADPTLNGMSAVVHLMCMLQRGSALPENSLDQDMEYVWAHDAMHVSRFHFQPVDEALMEMSPKFDVKTLVPLQRGMLSSLKWIGTRMLPLAEAEVQIRMSAVGMNFHDMMIAMNMFDSPLTLGSGYNSIGMEGVGYVTRKSPEVDHVQVGDRVIVIGSNSSGFATDVHRPADYCIKCPSSLTDVEAAGMSFAYMTVLWSFLDKGGLRKGQSVLIHSAAGGVGIAALHVSRWLGLEAFVTVGNEEKVRFIMNNFGLPRNRIFNSHSATFLDDVMDATAGRGVDAALSAAAGELLHNTWSCIAPGGVMLEIGKRDLINRGRLSLAPFEENRSYVGIDFSRLTIVNKPAVVRLLRQTMRLVEQGHIHPIHPTTCFDAEHAEDAFRFMQTGQHIGRIVVKIPRDTSTIPLASRPPAPEFEGQKTYLLVGGMGGLGRSVASWMVSAGARNLIFMSRSAGKSEQDQNFARELELSGCKVHCCAIDITDTDAVREALERTQASVAGVLQMAMVLRDVGIMNMDKANWDAAVAPKVQGTWNLHHALPNVDFFVMFGSNSGTLGSYGQANYAAANAFLDSFVQYRQNLGQAASVIDIGAVGDVGYVAETQVAAENMESMAGRLISEQDFLNCLQLAIARSTPTERRNKGPSTEADGYIDLKQVILLNSSILSMADPSNQIFWRKDPRMGIYRNVQRTSAESPTTDSNSLRRLVATLKADSSIADQAELAQTLARELSKQVATVLMLGEDEIDVDRSLTDVGMDSLVAIEMRNWWKQNLGVDVSVLELKDGRSILRLGELAATRLKERYSRDS

>PKSI_2_Hybr_12620

MAKVERKSWRPEDVAVIGLACRFSGSASNEAKLWELLEKCESAHSKVPGERYNVEAFHQQGSTNSNNLAADGGHYLEQDVKSFDAPFFNIITKEAKAMDPQARMLLESSYEALENAGLSLESVRGSDTGCYVGCFNRDYYELLMADAEDSPEYSVTGTGFSLLANRLSWYYDLRGPSKSEDTACSSSLVALDSAYKSLLRGESKMAMVCGANLMLSPNIGLWLSKLNMLSSEGLSRSFAEGVSGYGRGEGIATVILKPLADALRDGDTIRAVINATGVNQDGHTKGITVPNSQAQSTLIESTYRRAGLDFADTGYFEAHGTGTAVGDPLEIEALERVIKHAKRTSPLHVGSIKSSIGHLEGAAGLAGLIKCILMLEKGVILPNLHFERPNRKISFENIVVPTTAVPWPEGVKRRASVNLFGYGGTNAHVIVEAFVVPCPQSRDDDLRNAQDDGLERPSPQRLFVLTGREQSTVRKMRLRYASYIQTMKDTASPSVKFDDLSYTLGKRRSRLDWAEAHVASDFKELEEKLSAPEVAATRSSNKTRLGFVFTGQGAQWPRMGLELMRYTAFRDSVEAADEYLTQKLDCSWSVIEELEKHGDESRVTSSELGQPLCTIIQVAMVDLLGSWNVRPTAVVGHSSGEIAAAYCTGVMTKQAAWQIAFHRGRECAKLKGKAPELEGSMLAVGLDVESIRPYFNNLQSGRINVACVNSPNSITISGDASEIRKLQAMLVADSVFARELPVENAYHSHHMELVAESYLHSISDVDIQHSVALSDITMVSSVTGQSIEPSELIPEYWVRNLVSPVLFADAVTAMLRGSRRRFRRGAKAEPAVDFLLELGPHATLQTPFSDIVKAQAQEDVKYASMLLRGENAVDSAMTTAGKLYCHGCPVDVTAVNDIRHECRVLVDLPAYPWNRSTKYWGVSRLMQGYLHRTHGYHSLLGARLIGSDALNPAWRHFLNLDDSPWIEEHVVHGAVVYPGAGFLSMAIEAALQLAQPGREIANVRLQNVRVLKALVIKEGEDDPEVITRFRQADSVSDEASSMRWAFEISCAKGHDERDNRATGQDEFERHATGQITLDYQPEHPYLSPLSEQIHDVRRGEYARLADTCVDTMKQDGFYEASKDVGLAYGHDFQGIATMARGPNSCCWDLRVTERSTSLPGAYESKHLIHPTTLDAIVHSLFGAMNGGKKFQNAALPVAFDSIMISPATLTTSGTKLSGFTVIREAKEREIVADIHVSSEDWSQPLVQIAGLRCTQMPSPESELHDQDARPSPVGTITYRPDIALFDENGLMNYLNERQKPESCAKVHPDLYSERLRNAVAQVVELAMFKDPGLSVLQIGGYDRGVTDSLLFTLKAESAGQALSSKIVLLDPSQETLVEIRQQYDAESSIVDALHFAVDKPLPSEVSREYGFDVVLVAIDHGLDDMSKQNVLAEAQNVLKAGGIFVVFDTLRAITERQYQHLALGEATERSPFDLALAIAKVAVVQTREGEFVEQDGSFNIPRWSYGPEMTRTIADSTVSLESDSVRLGDMAQGTPLRMLHAGHPKYAHFVTNTIQPLHLAAGQVKVELRFVDITHQDLVEPDMRALREASGVIKAVGSDVSLLRPGDNVCLSFVGHLSTSVNVDEALCQRIPPGVNMAEAACIPITLATALRALVGVAGVKPQHNVLVQAGGTKMGRAAILIASAANAVVYTTARDAEEVESLLALGISKQNIVPEGDPLLPTVTKILTGNRGWDVIVRTTKIVAETFILPECVADFGVVLDVFPSSGTGCTQETTISVMGIGSLLPEDPVLMQKTVSRIMDYLPQVSTLANSFDVFPSSAIPAALDRHGAQNQHRGVMLSFDQEDLVRVSPSATNTMKLYRDATYVMAGGLGGLGRSIARLLVDNGARNLVFLSRSGPNTTAATTMMSKMAGLGVTVKTLKCDVGDEKSVAAALDECSSMPPVRGVIQAAADIQDAIFDTYTFEQWQANLRPKVQGSWNLHCQLPEDMDFFVMLSSISGLIGHEGQAGYAAGNTFQDSLALFRHSRGLPAVTIDLGAMLDVGTIAEGSTTATFRSSDAVLMKAIDLHEIMTMCISNEINGYAIPAQVCTGLPSGGMLQVEQQEIPSYFHKPLFAALKCLGTSAVSAVNVAAPVEGVIDFAAQLTTVGSLDEADCVIANILRAHIAKAVQRAVDDIDLSQPLYSYGIDSLMAVELRAWIGEKMKADLTLFDILNAESIQALALKISKTSQLVQQEIRGMN

>PKSI_1_10511

MPDNVSFMDESQDLRHIRETSRTSHTTSILNVEGDYRNNDMPREPGECNRSTNGTVDHERMTPSADGGIPIAICGIGLRLPGGIRNDRDLYDSLYNKKDARGVIPEDRFSIDSFHSAHGKTGTIITKHGYFLQDIDLTKFDVNMFNMTPAEVERLDPHQRILLETVRETLESAGEASFRGKKVGTYVGNFTDDWLDLQNVDTVDFATYQLHGKMDFSLANRISYEYDLRGPSMTIKTACSSSALAIHEAVYSIRNGECDAAIVSGSNLNLAPRLWVGMSSQGAISPDGSSKTFDESANGYARGDGIAALFIKRLDDAVRDGNPVRAVIRSTASNADGRTPGMTMPSTEAQEALIRRAYDAANLPLSETAMVECHGTGTAVGDPMEANAVARCFGDQGMLIGSVKPNLGHSEGASAITSVVKAVLSLENRTILPNIKFHRPNPAIPWSEAKLTVPVEPLAWPKDRQERISVNSFGIGGSNVHVVLDSAASMGFRPRSLAPSKDDRPGRLLLFSGGHRASVEQSSSQHQDYVTKYPNRLSDVAYTLAKRPPVKCPGLTRSTFVFTGQGAQWLHMGKELLHESPVFAKSIGRMDSVIHSLKHAPQWTLEGIINDPENPSALTNAEISQPLCTAVQIGLVDLLKSWAIYPHAVLGHSSGEIGAAYASGVVDRAEAILLAFYRGYVCRFAQKAGGMAAVGLEKSQVIKYLQPGVCVACENSGSSVTLSGDLETLEEVLQSIRAENLNAFARKLQVGIAYHSDHMKALGGLYHQYITEHLDPKDPQVPFFSSVSGRALHSKDDFGATYWQDNLENPVLFHTAVLKSLEHTGDKQVHLEVGPHGALNGPLRQIYAETGSKARYVALQKRGANCFDTFLEGIGQLYCNGVPLQYPESADDRTLIDLPPYPWHYDHSYWSETRVMKNWRFRRQLPHDLLGLRTLDCSDAEPMWRNILRITDLPWLRDHCVGKDVVFPASGYMCMAGEAVFQETGCRDYTLREVDISTAMVLSSDHSTELLTTMKKRRLNAFLDSRWYEFLIMSYDGASWTQHCSGLVTNGPSTSHPKALLQTYDRPVSTNRWYTAMSKIGLNYGPRFTGLQNITTHVQEKKASMTIMDKQEDYESPYALHPSTLDLILQSWTVASVRGEYRRFTQLFLPTFVDEFYIGNSASKLIHLNTTAIGPDGSARGEAIGKDNDGQISFNLKGFKGSKLDNVGVDQPQEMQTIMKQQWKRDFDFADTAQLMRPAFDSTSELSLLERMFVLAAIEVHVRTSGMEGKLPHHQRYKLWIDAQIRRFGEPGYPMVEDSMELLRLDSRERQRQLCRLLEHSRKTTAHPVAEAIWRALDRIEDVFDGRIEYLDLLFNDGLMPKFYDWSNSLSDVSRLFRLLSHKKPQLKILEVGAGTGGSTARLLQYLQSDFGERQYHSYTYTDVSSGFFVQAQERFKDYEGMKYRVLDISQDPFEQGFGADEFDLICASNVLHATPRLTETLRNCRKMLRPDGILFLQELCPRQQFMGFIMGLFEGWWLGAEDGRADTPLLLPPAWDRRLRDVGFEGVEAFSFDNNPPYFMAANMTARPTVTAKSKGSITLLTFNDFLDDVAGALKQAIRAAGFEIDHCVWGEHVPLDQSLISLVDLERDKPLLQDIGDDDLRIFLDLIDAVLQTTVIWLNKPAQVSSADPNAAQMLGLARTLRAELAMHFATVEMADPTLNGMSAVVHLMCMLQRGSALPENSLDQDMEYVWAHDAMHVSRFHFQPVDEALMEMSPKFDVKTLVPLQRGMLSSLKWIGTRMLPLAEAEVQIRMSAVGMNFHDMMIAMNMFDSPLTLGSGYNSIGMEGVGYVTRKSPEVDHVQVGDRVIVIGSNSSGFATDVHRPADYCIKCPSSLTDVEAAGMSFAYMTVLWSFLDKGGLRKGQSVLIHSAAGGVGIAALHVSRWLGLEAFVTVGNEEKVRFIMNNFGLPRNRIFNSHSATFLDDVMDATAGRGVDAALSAAAGELLHNTWSCIAPGGVMLEIGKRDLINRGRLSLAPFEENRSYVGIDFSRLTIVNKPAVVRLLRQTMRLVEQGHIHPIHPTTCFDAEHAEDAFRFMQTGQHIGRIVVKIPRDTSTIPLASRPPAPEFEGQKTYLLVGGMGGLGRSVASWMVSAGARNLIFMSRSAGKSEQDQNFARELELSGCKVHCCAIDITDTDAVREALERTQASVAGVLQMAMVLRDVGIMNMDKANWDAAVAPKVQGTWNLHHALPNVDFFVMFGSNSGTLGSYGQANYAAANAFLDSFVQYRQNLGQAASVIDIGAVGDVGYVAETQVAAENMESMAGRLISEQDFLNCLQLAIARSTPTERRNKGPSTEADGYIDLKQVILLNSSILSMADPSNQIFWRKDPRMGIYRNVQRTSAESPTTDSNSLRRLVATLKADSSIADQAELAQTLARELSKQVATVLMLGEDEIDVDRSLTDVGMDSLVAIEMRNWWKQNLGVDVSVLELKDGRSILRLGELAATRLKERYSRDS

>PKSI_1_15

MSNVLLFGDQTAEQYPLLNKIVLRKENALVITFVERCAKALREETNALPRSQRYAVPDFLTVNDLKEAYHQKGVKVPMVESALVTIAQIGHYIGYFSEHSAEQPSATNTRALGLCTGLLAAAAVVASKTVEELVLVGVDFVRLSFRSGAAVDAARTALCQTGDDNAPWSTIVTGTTEASAKEALAKFHEEKGIPQTSHAYISAVSVMAITVSGPPTTVKRFFEESPALSKNHRVPIPVYGPYHAEHLFGETEINKIASDSILEGLKQHQPVSLVHSAATGKALVAENAAELAKLVLAEMLQHSVRWDHLLEEAVSQITSKKAPAKIWAMGVSNVANSLVSALKAGGQSDVSTVDQSSWTENEPDTHGRTQNDKVAIVGMAGRFPNSADHEALWDLLMKGLDVHRRIPKDRFDADTHVDPSGKGKNKSHTPFGCFIDEPGFFDPRFFNMSPREAAQTDPMGRLALVTAYEALEMSGYVPNRTPSTKLHRIGTFYGQTSDDWREINAAENVDTYFITGGVRAFAPGRINYYFKFSGPSYSVDTACSSSLAAIQLACTSLWAGDCDTACAGGLNVLTNPDIFSGLSKGQFLSKTGSCKTYDNNADGYCRGDAVGTVILKRYEDAIADKDNILGCILGAATNHSAEAVSITHPHAGAQEFLYKRVLANAGVDAHEISYVEMHGTGTQAGDGIEMTSVTNVFAPRHRQRRDDQPVYLGAIKANVGHAEAASGINSLAKVLMMMKHNKIPANVGIKGEMNKTFPADLKDRKVNISQKAVEWPRNGKEKRKVFLNNFSAAGGNTALLLEDGPAYEAPTATDPRGTVPVTVTARSISALKRNIANLQKYVSENPSTTLTSMSYTLTARRIQHNYRVAFPLDQIDKFSDALQAQVKESYSPVPNVPTRVAFCFTGQGSQYTGLGQKLYNDLKSFRDDIDQLDHLARVQGLPSFLEIVQGADVQTLSPVKVQLGMACIQVALARMWAAWGITPAAVIGHSLGEYAALHVAGVISASDMVLLVGRRAELLVRDCTPHTHGMLAVKGGAEAIRNTLGNKMTEIACINGPEETVLCGSGDVVGAANETLAAKGFKATKLNVPFAFHSAQVDPILEQFKKIAASVTYNKPAVPVLSPLEGDIIREAGKINPEYLARHARETVNFWTALTAGQKEKVFDEKTAWLEVGAHPVCSGMVKASIGATTTAPSLRRGEDAWKTISNSMCTLFTAGVNFNFDEFHKEFNDAQEMYTLPTYSFDNKKYWLDYHNDWTLRKGEPAQTKEVIVEKPVASASAPAVEMPAKRLSTSCQRVIAENFSGNNGSVTVQSSLADPKLYPVVCGHMVNNAALCPSSLYADMALTISDYIWKQMRPGTETPGYNVCNMEVPKPLIAQIPQPAEGQHIQLEANADLDSGIVKLNFRSVKPDGQKLQDHAHCIVRLEDRAAWEDEWSRYNYMVQAQMELLQHKTLNGGAHKVQRGMAYKLFKALVNYDEKYRAMAEVVLASGQTEASAVLDFPTKPEDGDFYCPPYHIDGSCHISGFIVNASDLLDSEQNVYVSHGWGAMKFSRPLTAGMKLRNYVRMQPQPNNVSKGDVYIMEGDQIVAVCEGIKFQQIPRRVLNTFLPPNKGSGPASAAKPAAAPVAAARPAPAAAPIKTAPAPAKAAPAPAPAAPKAAPKPKKAAAPKKAAGGLTAKVMKILAKETEVDEGELVDEAQFENLGVDSLLSLTISAVFREELDMDISSTLFTDYPTVGDMKKYFAQFDNGSSTSSSAEEEDSDEDSIPPTDAATPRDDLSTPASSVPSSAPSDAGKPDSPTRETLEDVGDVSLAKHIVAQEMGVDIAEVTDDADLAEMGMDSLMSLTILGELREKTGIDLPSTFLTTNPTMKDIDNALGMRPKPKAAPKPAAPKAAAPSSSKKTDMNEVSARLSALNNNTDISRYPNATSVLLQGNPKQATKKIFFLPDGSGSATSYVSIPNLGPDVCAYGLNCPFMKNPEQWQCGIEISALVYLAEIKRRQPQGPYIIGGWSAGGVIAYSVAQALLAANEGVEKLLLLDSPCPVNLAPLPARLHNFFNEIGLLGTGDPAKTPKWLLPHFSAAIRSLSDYDPKPSLRPIPTYAIWCREGVAGNPGDPRPPPAEEEDPAPMTWLLEHRTNFKDNGWAQLCGDSMKFGVMGGHHFSMMKPPHADDLGNLIREGLDWQP

>PKSI_1_153

MSNVLLFGDQTAEQYPLLNKIVLRKENALVITFVERCAKALREETNALPRSQRNAVPDFLTVNDLKEAYHQKGVKVPMVESALVTIAQIGHYIGYFSEHSAEQPSATNTRALGLCTGLLAAAAVVASKTVEELVLVGVDFVRLSFRSGAAVDAARTALCQTGDDNAPWSTIVTGTTEASAKEALAKFHEEKGIPQTSHAYISAVSVMAITVSGPPTTVKRFFEESPALSKNHRVPIPVYGPYHAEHLFGETEINKIASDSILEGLKQHQPVSLVHSAATGKALVAENAAELAKLVLAEMLQHSVRWDHLLEEAVSQITSKKAPAKIWAMGVSNVANSLVSALKAGGQSDVSTVDQSSWTENEPDTHGRTQNDKVAIVGMAGRFPNSADHEALWDLLMKGLDVHRRIPKDRFDADTHVDPSGKGKNKSHTPFGCFIDEPGFFDPRFFNMSPREAAQTDPMGRLALVTAYEALEMSGYVPNRTPSTKLHRIGTFYGQTSDDWREINAAENVDTYFITGGVRAFAPGRINYYFKFSGPSYSVDTACSSSLAAIQLACTSLWAGDCDTACAGGLNVLTNPDIFSGLSKGQFLSKTGSCKTYDNNADGYCRGDAVGTVILKRYEDAIADKDNILGCILGAATNHSAEAVSITHPHAGAQEFLYKRVLANAGVDAHEISYVEMHGTGTQAGDGIEMTSVTNVFAPRHRQRRDDQPVYLGAIKANVGHAEAASGINSLAKVLMMMKHNKIPANVGIKGEMNKTFPADLKDRKVNISQKAVEWPRNGKEKRKVFLNNFSAAGGNTALLLEDGPAYEAPTATDPRGTVPVTVTARSISALKRNIANLQKYVSENPSTTLTSMSYTLTARRIQHNYRVAFPLDQIDKFSDALQAQVKESYSPVPNVPTRVAFCFTGQGSQYTGLGQKLYNDLKSFRDDIDQLDHLARVQGLPSFLEIVQGADVQTLSPVKVQLGMACIQVALARMWAAWGITPAAVIGHSLGEYAALHVAGVISASDMVLLVGRRAELLVRDCTPHTHGMLAVKGGAEAIRNTLGNKMTEIACINGPEETVLCGSGDVVGAANETLAAKGFKATKLNVPFAFHSAQVDPILEQFKKIAASVTYNKPAVPVLSPLEGDIIREAGKINPEYLARHARETVNFWTALTAGQKEKVFDEKTAWLEVGAHPVCSGMVKASIGATTTAPSLRRGEDAWKTISNSMCTLFTAGVNFNFDEFHKEFNDAQEMYTLPTYSFDNKKYWLDYHNDWTLRKGEPAQTKEVIVEKPVASASAPAVEMPAKRLSTSCQRVIAENFSGNNGSVTVQSSLADPKLYPVVCGHMVNNAALCPSSLYADMALTISDYIWKQMRPGTETPGYNVCNME

>PKSI_1_9

MSNVLLFGDQTAEQYPLLNKIVLRKENALVITFVERCAKALREETNALPRSQRNAVPDFLTVNDLKEAYHQKGVKVPMVESALVTIAQIGHYIGYFSEHSAEQPSATNTRALGLCTGLLAAAAVVASKTVEELVLVGVEFVRLSFRSGAAVDAARTALCQTGDDNAPWSTIVTGTTEASAKEALAKFHEEKGIPQTSHAYISAVSVMAITVSGPPTTVKRFFEESPALSKNHRVPIPVYGPYHAEHLFGETEINKIASASILEGLKQHQPVSLVHSAATGKALVAENAAELAKLVLAEMLQHPVRWDHLLEEAVSQITSKKAPAKIWAMGVSNVANSLVSALKAGGQSDVSTVDQSSWTENEPDTHGRTQNDKVAIVGMAGRFPNSADHEALWDLLMKGLDVHRRIPKDRFDADTHVDPSGKGKNKSHTPFGCFIDEPGFFDPRFFNMSPREAAQTDPMGRLALVTAYEALEMSGYVPNRTPSTKLHRIGTFYGQTSDDWREINAAENVDTYFITGGVRAFAPGRINYYFKFSGPSYSVDTACSSSLAAIQLACTSLWAGDCDTACAGGLNVLTNPDIFSGLSKGQFLSKTGSCKTYDNNADGYCRGDAVGTVILKRYEDAIADKDNILGCILGAATNHSAEAVSITHPHAGAQEFLYKRVLANAGVDAHEISYVEMHGTGTQAGDGIEMTSVTNVFAPRHRQRRDDQPVYLGAIKANVGHAEAASGINSLAKVLMMMKHNKIPANVGIKGEMNKTFPADLKDRKVNISQKAVEWPRNGKEKRKVFLNNFSAAGGNTALLLEDGPAYEAPTASDPRGTVPVTVTARSISALKRNIANLQKYVSENPSTTLTSMSYTLTARRIQHNYRVAFPLDQINKFSDALQAQVKESYSPVPNAPTRVAFCFTGQGSQYTGLGQKLYNDLKSFRDDIDQLDHLARVQGLPSFLEIVQGADVQTLSPVKVQLGMACIQVALARMWAAWGITPAAVIGHSLGEYAALHVAGVISASDMVLLVGRRAELLVRDCTPHTHGMLAVKGGAEAIRNTLGNKMTEIACINGPEETVLCGSGDVVGAANETLAAKGFKATKLNVPFAFHSAQVDPILEQFKKIAASVTYNKPAVPVLSPLEGDIIREAGKINPEYLARHARETVNFWTALTAGQKEKVFDEKTAWLEVGAHPVCSGMVKASIGATTTAPSLRRGEDAWKTISNSMCTLFTAGVNFNFDEFHKEFNDAQEMYTLPTYSFDNKKYWLDYHNDWTLRKGEPAQTKEVIVEKPVASASAPAVEIPAKRLSTSCQRVISENFSGNNGSVTVQSSLADPKLYPVVCGHMVNNAALCPSTLYADMALTISDYIWKQMRPGTETPGYNVCNMEVPKPLIAQIPQPAEGQHIQLEANADLDSGIVKLNFRSVKPDGQKLQDHAHCIVRLEDRAAWEDEWSRYNYMVQAQMELLQHKTLNGGAHKVQRGMAYKLFKALVNYDEKYRAMAEVVLASGQTEASAVLDFPTKPEDGDFYCPPYHIDGSCHISGFIVNASDLLDSEQNVYVSHGWGAMKFSRPLTAGMKLRNYVRMQPQPNNVSKGDVYIMEGDQIVAVCEGIKFQQIPRRVLNTFLPPNKGSGPASAAKPAAAPVAAARPAPAAAPIKTAPAPAKAAPAPAPAAPKAAPKPKKAAAPKKAAGGLTAKVMKILAKETEVDEGELVDEAQFENLGVDSLLSLTISAVFREELDMDISSTLFTDYPTVGDMKKYFAQFDNGSSTSSSAEEEDSDEDSIPPTDAATPMDDLSTPASSVPSSAPSDAGKPDSPTRETLEDVGDVSLAKHIVAQEMGVDIAEVTDDADLAEMGMDSLMSLTILGELREKTGIDLPSTFLTTNPTMKDIDNALGMRPKPKAAPKPAAPKAAAPSSSKKTDMNEVSARLSALNNNTDISRYPNATSVLLQGNPKQATKKIFFLPDGSGSATSYVSIPNLGPDVCAYGLNCPFMKNPEQWQCGIEISALVYLAEIKRRQPQGPYIIGGWSAGGVIAYSVAQALLAANEGVEKLLLLDSPCPVNLAPLPARLHNFFNEIGLLGTGDPAKTPKWLLPHFSAAIRSLSDYDPKPSLRPIPTYAIWCREGVAGNPGDPRPPPAEEEDPAPMTWLLEHRTDFKDNGWAQLCGDSMKFGVMGGHHFSMMKPPHADDLGNLIREGLDWQP

>PKSI_1_10510

MPDNVSFMDESQDLRHIRETSRTSHTTSILNVEGDYRNNDMPREPGECNRSTNGTVDHERMTPSADGGIPIAICGIGLRLPGGIRNDRDLYDSLYNKKDARGVIPEDRFSIDSFHSAHGKTGTIITKHGYFLQDIDLTKFDVNMFNMTPAEVERLDPHQRILLETVRETLESAGEASFRGKKVGTYVGNFTDDWLDLQNVDTVDFATYQLHGKMDFSLANRISYEYDLRGPSMTIKTACSSSALAIHEAVYSIRNGECDAAIVSGSNLNLAPRLWVGMSSQGAISPDGSSKTFDESANGYARGDGIAALFIKRLDDAVRDGNPVRAVIRSTASNADGRTPGMTMPSTEAQEALIRRAYDAANLPLSETAMVECHGTGTAVGDPMEANAVARCFGDQGMLIGSVKPNLGHSEGASAITSVVKAVLSLENRTILPNIKFHRPNPAIPWSEAKLTVPVEPLAWPKDRQERISVNSFGIGGSNVHVVLDSAASMGFRPRSLAPSKDDRPGRLLLFSGGHRASVEQSSSQHQDYVTKYPNRLSDVAYTLAKRPPVKCPGLTRSTFVFTGQGAQWLHMGKELLHESPVFAKSIGRMDSVIHSLKHAPQWTLEGIINDPENPSALTNAEISQPLCTAVQIGLVDLLKSWAIYPHAVLGHSSGEIGAAYASGVVDRAEAILLAFYRGYVCRFAQKAGGMAAVGLEKSQVIKYLQPGVCVACENSGSSVTLSGDLETLEEVLQSIRAENLNAFARKLQVGIAYHSDHMKALGGLYHQYITEHLDPKDPQVPFFSSVSGRALHSKDDFGATYWQDNLENPVLFHTAVLKSLEHTGDKQVHLEVGPHGALNGPLRQIYAETGSKARYVALQKRGANCFDTFLEGIGQLYCNGVPLQYPESADDRTLIDLPPYPWHYDHSYWSETRVMKNWRFRRQLPHDLLGLRTLDCSDAEPMWRNILRITDLPWLRDHCVGKDVVFPASGYMCMAGEAVFQETGCRDYTLREVDISTAMVLSSDHSTELLTTMKKRRLNAFLDSRWYEFLIMSYDGASWTQHCSGLVTNGPSTSHPKALLQTYDRPVSTNRWYTAMSKIGLNYGPRFTGLQNITTHVQEKKASMTIMDKQEDYESPYALHPSTLDLILQSWTVASVRGEYRRFTQLFLPTFVDEFYIGNSASKLIHLNTTAIGPDGSARGEAIGKDNDGQISFNLKGFKGSKLDNVGVDQPQEMQTIMKQQWKRDFDFADTAQLMRPAFDSTSELSLLERMFVLAAIEVHVRTSGMEGKLPHHQRYKLWIDAQIRRFGEPGYPMVEDSMELLRLDSRERQRQLCRLLEHSRKTTAHPVAEAIWRALDRIEDVFDGRIEYLDLLFNDGLMPKFYDWSNSLSDVSRLFRLLSHKKPQLKILEVGAGTGGSTARLLQYLQSDFGERQYHSYTYTDVSSGFFVQAQERFKDYEGMKYRVLDISQDPFEQGFGADEFDLICASNVLHATPRLTETLRNCRKMLRPDGILFLQELCPRQQFMGFIMGLFEGWWLGAEDGRADTPLLLPPAWDRRLRDVGFEGVEAFSFDNNPPYFMAANMTARPTVTAKSKGSITLLTFNDFLDDVAGALKQAIRAAGFEIDHCVWGEHVPLDQSLISLVDLERDKPLLQDIGDDDLRIFLDLIDAVLQTTVIWLNKPAQVSSADPNAAQMLGLARTLRAELAMHFATVEMADPTLNGMSAVVHLMCMLQRGSALPENSLDQDMEYVWAHDAMHVSRFHFQPVDEALMEMSPKFDVKTLVPLQRGMLSSLKWIGTRMLPLAEAEVQIRMSAVGMNFHDMMIAMNMFDSPLTLGSGYNSIGMEGVGYVTRKSPEVDHVQVGDRVIVIGSNSSGFATDVHRPADYCIKCPSSLTDVEAAGMSFAYMTVLWSFLDKGGLRKGQSVLIHSAAGGVGIAALHVSRWLGLEAFVTVGNEEKVRFIMNNFGLPRNRIFNSHSATFLDDVMDATAGRGVDAALSAAAGELLHNTWSCIAPGGVMLEIGKRDLINRGRLSLAPFEENRSYVGIDFSRLTIVNKPAVVRLLRQTMRLVEQGHIHPIHPTTCFDAEHAEDAFRFMQTGQHIGRIVVKIPRDTSTIPLASRPPAPEFEGQKTYLLVGGMGGLGRSVASWMVSAGARNLIFMSRSAGKSEQDQNFARELELSGCKVHCCAIDITDTDAVREALERTQASVAGVLQMAMVLRDVGIMNMDKANWDAAVAPKVQGTWNLHHALPNVDFFVMFGSNSGTLGSYGQANYAAANAFLDSFVQYRQNLGQAASVIDIGAVGDVGYVAETQVAAENMESMAGRLISEQDFLNCLQLAIARSTPTERRNKGPSTEADGYIDLKQVILLNSSILSMADPSNQIFWRKDPRMGIYRNVQRTSAESPTTDSNSLRRLVATLKADSSIADQAELAQTLARELSKQVATVLMLGEDEIDVDRSLTDVGMDSLVAIEMRNWWKQNLGVDVSVLELKDGRSILRLGELAATRLKERYSRDS

>PKSI_1_10508

MPDNVSFMDESQDLRHIRETSRTSHTTSILNVEGDYRNNDMPREPGECNRSTNGTVDHERMTPSADGGIPIAICGIGLRLPGGIRNDRDLYDSLYNKKDARGVIPEDRFSIDSFHSAHGKTGTIITKHGYFLQDIDLTKFDVNMFNMTPAEVERLDPHQRILLETVRETLESAGEASFRGKKVGTYVGNFTDDWLDLQNVDTVDFATYQLHGKMDFSLANRISYEYDLRGPSMTIKTACSSSALAIHEAVYSIRNGECDAAIVSGSNLNLAPRLWVGMSSQGAISPDGSSKTFDESANGYARGDGIAALFIKRLDDAVRDGNPVRAVIRSTASNADGRTPGMTMPSTEAQEALIRRAYDAANLPLSETAMVECHGTGTAVGDPMEANAVARCFGDQGMLIGSVKPNLGHSEGASAITSVVKAVLSLENRTILPNIKFHRPNPAIPWSEAKLTVPVEPLAWPKDRQERISVNSFGIGGSNVHVVLDSAASMGFRPRSLAPSKDDRPGRLLLFSGGHRASVEQSSSQHQDYVTKYPNRLSDVAYTLAKRPPVKCPGLTRSTFVFTGQGAQWLHMGKELLHESPVFAKSIGRMDSVIHSLKHAPQWTLEGIINDPENPSALTNAEISQPLCTAVQIGLVDLLKSWAIYPHAVLGHSSGEIGAAYASGVVDRAEAILLAFYRGYVCRFAQKAGGMAAVGLEKSQVIKYLQPGVCVACENSGSSVTLSGDLETLEEVLQSIRAENLNAFARKLQVGIAYHSDHMKALGGLYHQYITEHLDPKDPQVPFFSSVSGRALHSKDDFGATYWQDNLENPVLFHTAVLKSLEHTGDKQVHLEVGPHGALNGPLRQIYAETGSKARYVALQKRGANCFDTFLEGIGQLYCNGVPLQYPESADDRTLIDLPPYPWHYDHSYWSETRVMKNWRFRRQLPHDLLGLRTLDCSDAEPMWRNILRITDLPWLRDHCVGKDVVFPASGYMCMAGEAVFQETGCRDYTLREVDISTAMVLSSDHSTELLTTMKKRRLNAFLDSRWYEFLIMSYDGASWTQHCSGLVTNGPSTSHPKALLQTYDRPVSTNRWYTAMSKIGLNYGPRFTGLQNITTHVQEKKASMTIMDKQEDYESPYALHPSTLDLILQSWTVASVRGEYRRFTQLFLPTFVDEFYIGNSASKLIHLNTTAIGPDGSARGEAIGKDNDGQISFNLKGFKGSKLDNVGVDQPQEMQTIMKQQWKRDFDFADTAQLMRPAFDSTSELSLLERMFVLAAIEVHVRTSGMEGKLPHHQRYKLWIDAQIRRFGEPGYPMVEDSMELLRLDSRERQRQLCRLLEHSRKTTAHPVAEAIWRALDRIEDVFDGRIEYLDLLFNDGLMPKFYDWSNSLSDVSRLFRLLSHKKPQLKILEVGAGTGGSTARLLQYLQSDFGERQYHSYTYTDVSSGFFVQAQERFKDYEGMKYRVLDISQDPFEQGFGADEFDLICASNVLHATPRLTETLRNCRKMLRPDGILFLQELCPRQQFMGFIMGLFEGWWLGAEDGRADTPLLLPPAWDRRLRDVGFEGVEAFSFDNNPPYFMAANMTARPTVTAKSKGSITLLTFNDFLDDVAGALKQAIRAAGFEIDHCVWGEHVPLDQSLISLVDLERDKPLLQDIGDDDLRIFLDLIDAVLQTTVIWLNKPAQVSSADPNAAQMLGLARTLRAELAMHFATVEMADPTLNGMSAVVHLMCMLQRGSALPENSLDQDMEYVWAHDAMHVSRFHFQPVDEALMEMSPKFDVKTLVPLQRGMLSSLKWIGTRMLPLAEAEVQIRMSAVGMNFHDMMIAMNMFDSPLTLGSGYNSIGMEGVGYVTRKSPEVDHVQVGDRVIVIGSNSSGFATDVHRPADYCIKCPSSLTDVEAAGMSFAYMTVLWSFLDKGGLRKGQSVLIHSAAGGVGIAALHVSRWLGLEAFVTVGNEEKVRFIMNNFGLPRNRIFNSHSATFLDDVMDATAGRGVDAALSAAAGELLHNTWSCIAPGGVMLEIGKRDLINRGRLSLAPFEENRSYVGIDFSRLTIVNKPAVVRLLRQTMRLVEQGHIHPIHPTTCFDAEHAEDAFRFMQTGQHIGRIVVKIPRDTSTIPLASRPPAPEFEGQKTYLLVGGMGGLGRSVASWMVSAGARNLIFMSRSAGKSEQDQNFARELELSGCKVHCCAIDITDTDAVREALERTQASVAGVLQMAMVLRDVGIMNMDKANWDAAVAPKVQGTWNLHHALPNVDFFVMFGSNSGTLGSYGQANYAAANAFLDSFVQYRQNLGQAASVIDIGAVGDVGYVAETQVAAENMESMAGRLISEQDFLNCLQLAIARSTPTERRNKGPSTEADGYIDLKQVILLNSSILSMADPSNQIFWRKDPRMGIYRNVQRTSAESPTTDSNSLRRLVATLKADSSIADQAELAQTLARELSKQVATVLMLGEDEIDVDRSLTDVGMDSLVAIEMRNWWKQNLGVDVSVLELKDGRSILRLGELAATRLKERYSRDS

>PKSI_1_10509

MPDNVSFMDESQDLRHIRETSRTSHTTSILNVEGDYRNNDMPREPGECNRSTNGTVDHERMTPSADGGIPIAICGIGLRLPGGIRNDRDLYDSLYNKKDARGVIPEDRFSIDSFHSAHGKTGTIITKHGYFLQDIDLTKFDVNMFNMTPAEVERLDPHQRILLETVRETLESAGEASFRGKKVGTYVGNFTDDWLDLQNVDTVDFATYQLHGKMDFSLANRISYEYDLRGPSMTIKTACSSSALAIHEAVYSIRNGECDAAIVSGSNLNLAPRLWVGMSSQGAISPDGSSKTFDESANGYARGDGIAALFIKRLDDAVRDGNPVRAVIRSTASNADGRTPGMTMPSTEAQEALIRRAYDAANLPLSETAMVECHGTGTAVGDPMEANAVARCFGDQGMLIGSVKPNLGHSEGASAITSVVKAVLSLENRTILPNIKFHRPNPAIPWSEAKLTVPVEPLAWPKDRQERISVNSFGIGGSNVHVVLDSAASMGFRPRSLAPSKDDRPGRLLLFSGGHRASVEQSSSQHQDYVTKYPNRLSDVAYTLAKRPPVKCPGLTRSTFVFTGQGAQWLHMGKELLHESPVFAKSIGRMDSVIHSLKHAPQWTLEGIINDPENPSALTNAEISQPLCTAVQIGLVDLLKSWAIYPHAVLGHSSGEIGAAYASGVVDRAEAILLAFYRGYVCRFAQKAGGMAAVGLEKSQVIKYLQPGVCVACENSGSSVTLSGDLETLEEVLQSIRAENLNAFARKLQVGIAYHSDHMKALGGLYHQYITEHLDPKDPQVPFFSSVSGRALHSKDDFGATYWQDNLENPVLFHTAVLKSLEHTGDKQVHLEVGPHGALNGPLRQIYAETGSKARYVALQKRGANCFDTFLEGIGQLYCNGVPLQYPESADDRTLIDLPPYPWHYDHSYWSETRVMKNWRFRRQLPHDLLGLRTLDCSDAEPMWRNILRITDLPWLRDHCVGKDVVFPASGYMCMAGEAVFQETGCRDYTLREVDISTAMVLSSDHSTELLTTMKKRRLNAFLDSRWYEFLIMSYDGASWTQHCSGLVTNGPSTSHPKALLQTYDRPVSTNRWYTAMSKIGLNYGPRFTGLQNITTHVQEKKASMTIMDKQEDYESPYALHPSTLDLILQSWTVASVRGEYRRFTQLFLPTFVDEFYIGNSASKLIHLNTTAIGPDGSARGEAIGKDNDGQISFNLKGFKGSKLDNVGVDQPQEMQTIMKQQWKRDFDFADTAQLMRPAFDSTSELSLLERMFVLAAIEVHVRTSGMEGKLPHHQRYKLWIDAQIRRFGEPGYPMVEDSMELLRLDSRERQRQLCRLLEHSRKTTAHPVAEAIWRALDRIEDVFDGRIEYLDLLFNDGLMPKFYDWSNSLSDVSRLFRLLSHKKPQLKILEVGAGTGGSTARLLQYLQSDFGERQYHSYTYTDVSSGFFVQAQERFKDYEGMKYRVLDISQDPFEQGFGADEFDLICASNVLHATPRLTETLRNCRKMLRPDGILFLQELCPRQQFMGFIMGLFEGWWLGAEDGRADTPLLLPPAWDRRLRDVGFEGVEAFSFDNNPPYFMAANMTARPTVTAKSKGSITLLTFNDFLDDVAGALKQAIRAAGFEIDHCVWGEHVPLDQSLISLVDLERDKPLLQDIGDDDLRIFLDLIDAVLQTTVIWLNKPAQVSSADPNAAQMLGLARTLRAELAMHFATVEMADPTLNGMSAVVHLMCMLQRGSALPENSLDQDMEYVWAHDAMHVSRFHFQPVDEALMEMSPKFDVKTLVPLQRGMLSSLKWIGTRMLPLAEAEVQIRMSAVGMNFHDMMIAMNMFDSPLTLGSGYNSIGMEGVGYVTRKSPEVDHVQVGDRVIVIGSNSSGFATDVHRPADYCIKCPSSLTDVEAAGMSFAYMTVLWSFLDKGGLRKGQSVLIHSAAGGVGIAALHVSRWLGLEAFVTVGNEEKVRFIMNNFGLPRNRIFNSHSATFLDDVMDATAGRGVDAALSAAAGELLHNTWSCIAPGGVMLEIGKRDLINRGRLSLAPFEENRSYVGIDFSRLTIVNKPAVVRLLRQTMRLVEQGHIHPIHPTTCFDAEHAEDAFRFMQTGQHIGRIVVKIPRDTSTIPLASRPPAPEFEGQKTYLLVGGMGGLGRSVASWMVSAGARNLIFMSRSAGKSEQDQNFARELELSGCKVHCCAIDITDTDAVREALERTQASVAGVLQMAMVLRDVGIMNMDKANWDAAVAPKVQGTWNLHHALPNVDFFVMFGSNSGTLGSYGQANYAAANAFLDSFVQYRQNLGQAASVIDIGAVGDVGYVAETQVAAENMESMAGRLISEQDFLNCLQLAIARSTPTERRNKGPSTEADGYIDLKQVILLNSSILSMADPSNQIFWRKDPRMGIYRNVQRTSAESPTTDSNSLRRLVATLKADSSIADQAELAQTLARELSKQVATVLMLGEDEIDVDRSLTDVGMDSLVAIEMRNWWKQNLGVDVSVLELKDGRSILRLGELAATRLKERYSRDS

>PKSI_2_10509

MSNVLLFGDQTAEQYPLLNKIVLRKENALVITFVERCAKALREETNALPRSQRNAVPDFLTVNDLKEAYHQKGVKVPMVESALVTIAQIGHYIGYFSEHSAEQPSATNTRALGLCTGLLAAAAVVASKTVEELVLVGVEFVRLSFRSGAAVDAARTALCQTGDDNAPWSTIVTGTTEAAAKEALAKFHEEKRIPQTSHAYISAVSVMAITVSGPPTTVKRFFEESPALSKNHRVPIPVYGPYHAEHLFGETEINKIASASILEGLKQHQPVSLVHSAATGKALVAENAAELAKLVLAEMLQHPVRWDHLLEEAVSQITSKKAPAKIWAMGVSNVANSLVSALKAGGQTDVSTVDQSTWTENEPDTHGRTQNDKVAIVGMAGRFPNSADHEALWELLMKGLDVHRRIPKDRFDADTHVDPSGKGKNKSHTPFGCFIDEPGFFDPRFFNMSPREAAQTDPMGRLALVTAYEALEMSGYVPNRTPSTKLHRIGTFYGQTSDDWREINAAENVDTYFITGGVRAFAPGRINYYFKFSGPSYSVDTACSSSLAAIQLACTSLWAGDCDTACAGGLNVLTNPDIFSGLSKGQFLSKTGSCKTYDNNADGYCRGDAVGTVILKRYEDAIADKDNILGCILGAATNHSAEAVSITHPHAGAQEFLYKRVLANAGVDAHEISYVEMHGTGTQAGDGIEMTSVTNVFAPRHRQRRDDQPVYLGAIKANVGHAEAASGINSLAKVLLMMKHNKIPANVGIKGEMNKTFPADLKDRKVNISQKAVDWPRNGKEKRKVFLNNFSAAGGNTALLLEDGPAYEAPTATDPRGTVPVTVTARSISALKRNIANLQKYVSENPSTTLTSMSYTLTARRIQHNYRVAFPLDQINKFSDALQAQVKESYSPVPNAPTRVAFCFTGQGSQYTGLGQKLYNDLKSFRDDIDQLDHLARVQGLPSFLEIVQGADVQTLSPVKVQLGMACIQVALARMWAAWGITPAAVIGHSLGEYAALHVAGVISASDMVLLVGRRAELLVRDCTPHTHGMLAVKGGAEAIRNTLGSKMTEIACINGPEETVLCGSGDDVGAANETLAAKGFKATKLNVPFAFHSAQVDPILEQFKKIAASVTYNKPAVPVLSPLEGDIIREAGKINPEYLARHARETVNFWTALTAGQKEKVFDEKTAWLEVGAHPVCSGMVKASIGATTTAPSLRRGEDAWKTISNSMCTLFTAGVNFNFDEFHKEFNDAQEMYTLPTYSFDNKKYWLDYHNDWTLRKGEPAQTKEVIVEKPVASASAPAVEMPAKRLSTSCQRVIAENFSGNNGSVTVQSSLADPKLYPVVCGHMVNNAALCPSSLYADMALTISDYIWKQMRPGTETPGYNVCNMEVPKPLIAQIPQPAEGQHIQLEANADLDSGIVKLNFRSVKPDGQKLQDHAHCIVRLEDRAAWEDEWSRYNYMVQAQMELLQHKTLNGGAHKVQRGMAYKLFKALVNYDEKYRAMAEVVLASGQTEASAVLDFPTKPEDGDFYCPPYHIDGSCHISGFIVNASDLLDSEQNVYVSHGWGAMKFSRPLTAGMKLRNYVRMQPQPNNVSKGDVYIMEGDQIVAVCEGIKFQQIPRRVLNTFLPPNKGSGPASAAKPAAAPVAAARPAPAAAPIKTAPAPAKAAPAPAPAAPKAAPKPKKAAAPKKAAGGLTAKVMKILAKETEVDEGELVDEAQFENLGVDSLLSLTISAVFREELDMDISSTLFTDYPTVGDMKKYFAQFDNGSSTSSSTEEEDSDEDSIPPTDAATPMDDLSTPASSVGSSAPSDAGKPESPTRETLEDVGDVSLAKHIVAQEMGVDIAEVTDDADLAEMGMDSLMSLTILGELREKTGIDLPSTFLTTNPTMKDIDNALGMRPKPKAAPKPAAPKAAAPSSSKKTDMNEVSARLSALNNNTDISRYPNATSVLLQGNPKQATKKIFFLPDGSGSATSYVSIPNLGPDVCAYGLNCPFMKNPEQWQCGIEISALVYLAEIKRRQPQGPYIIGGWSAGGVIAYSVAQALLAANEGVEKLLLLDSPCPVNLAPLPARLHNFFNEIGLLGTGDPAKTPKWLLPHFSAAIRSLSDYDPKPSLRPIPTYAIWCREGVAGNPGDPRPPPAEEEDPAPMTWLLEHRTNFKDNGWAQLCGDSMKFGVMGGHH

>PKSI_1_2785

MGYNSKSLAAHSKIPVDRFNIEAFHWLGATHPNNVAADGAHLLEQDVSAFDAPFFGITAEEAKVIDPQARMLLECSYDAFMNAGLKVESICGSDTGCYVGSFDLDYHQMLMGNFENAPRYSGTGTAFSLLSNRLSWFYNLKGPSLSLDTACSSSLVGLHLACQSLSAGESKMAMVCGAKLLLGPHLSMWLSRINMFSSDGKSRSFADDTTGYGGGEGIVTVILKPLADALRDNDPIRAVIKGTETPASSRRMKQGTGTAVGDPLELAAVAPLYVGSIKSNIGHTEGAAGLAGLIKCILMLEKGVILPNIHFDRPNKRIPFERAGIKVPTEVLPWPKNLDRRASINSFGFGGTNAHAIVESFPTPSTPSLDHAAQVDASEIVAVTPNHPRLFVISGHEPAAVDKLRQRYLEYINQAKNCDDVDCKLDDLFYTLGCRRSRMDWSVSLVASDFEELGKKLLENTTVVKRAARSPRIGFIFTGQGAQWPRMGVGLMRYGVFQESVQAADRFLSGQCHCGWSVIDELEKAKDDSRIASSELAQPICTIVQVAMIDLLRCWNIRPTAVAGHSSGEIAAAYCTGAITRQSAWEIAFHRGRECARLEEVAPDLRGAMLAVGLGVEDVRPYLDAVAPDRVNIACINSPNSTTLSGDAVEIQELLTKLIANGVSARELRVENAYHSHHMKLVADRYLDSITHVAVQTEVVRSDVALLSSVTGRLTSSSDLDPEYWVRNLVSPVLFSDAVSAMLKGTKKTFRRQQGTAEPAVDVLLEIGPHAALRGPLADILKSEAVESVAYVSMSNRGRDALESATAAAGELWSRGCPVDVTAVNNHPRQPRVLADLPSYPWDRLNKYWATSRVTHDVLHRAFPHHDLLGKRLAGSDALAPAWRHFLRFSESPWVREHVVHGSIVFSGAGFLAMAIEAALQLIEKDRKLANVRMRDVHIFKALVLEQEEEGAQELVTRFHRVDDRSDGTWSGWWKFSISCTKSHVEPERHVSGQIMLEYSPAEPSSQPASDVVHQAQKVKYGKLVGSSADRLCRAAFYEASEAAGLAYGSQFQGVVDVARSGDGRCCWRVQLPDRKAPASESKHLIHPTTLDAIVHAMFGAMNKGSAFSSAALPIAFDKVVISADMPTDAGTCLSGFTVTDVKETTREVTADVYVSSDDWARRLLQIEGLRCTELTSPEGAHNVDETQSAPLGSVAWQPDLDLLDNDGLKAYIMKSPERDQLGATEGRVAALSTRLRSAVAQVLYLAAFKTPTLSILQVGGFSENLTDTFVATIASDTNVPAVTAEMLVLDSEEENIPRIRERQHDGDKFDAVLLVVPEHNDEQATRVLVAQAQNLLKAGALLIVLDTIQNTRANVAFYGFETAQTSSSRHLRPWQSLNVEDPDCGKAAALCRESPATISRESIGVPICVLKPPHCTKETVDVISALKGILNDAGLEVAVEEWPPSVAQVRGKFVISLLDNEASFLGNIDAAGFEVLREVALQSRRLLWVCSGDDPHMAIALGWLRVLQNENANRVYQHLTLTRKKAAAPQSRSCAIARLALTQTREGEYAEEDGNLYIPRWYHEEGLSRTLEEGEVSVQLERAPLGVAKSSMPLRILHGKDTESARFVPDSPSVSRLAADEVEIEMQYVVLTDSDISPAGNTARREGSGLVRAVGRDVTRLRPADQVCVSYVGPLSTRVIAKEVYCQVVPVGAIMEEAACIPNTFATALRVLTDVARVKPGQTVLVQTAGTKIGTASVLLALALDAVVYATARDKAETDKIVSLGVSRPNVFTEGDMDLPQAVKTLAGERGLDVVLRTSKSMSAPCLLSHCVAEHGTLVDVHAASEPGQDAPSCEDTISVMGVGPLLPEDPVSLQKSASRAANYLPQLSGLAASFDLFESARISDALKCQQEQGTRGGVILSLDDADMVPIAPSVNRKLYFCEQATYVLAGGLGGLGQSLARLFVDHGARNLALLSRGGLNSPSAETFIKEMAEVGVAVKVLACDIGDDGSMKAALDDCAGTMPPIRGVIQAATVYRDAIFDNFTFEDWQANLRAKVQGSWNLHRHLPKDIDFFVMLGSVAGLMGHVSQAGYAAGNTFQDALAHYRRSRSLPAVTIDLGPMLDVGAVNDGTVSASFSTPEATWMTEADLHAIMIMCISGEITSCNFPPQLCTGLPSGGMLQLGQHELPEHYDRPFFALLKGLGVSAAAGKDNKVLRRSEDFSHQISAVTSMEEARKCVVDAVKAHLAKGLGRSANLIESSEPLHSYGIDSLSALGFRTWVRETMKADVSMFDVINERSIGELAEKIARISELTPVGLESGE

>PKSI_2_2785

MSRFTDGAALPTASTVGTQSGGPHVIDPTTEDNGPSPVPAPKGPIGAHPKNGEIRSEYPSHAVAVVGMAGRFPGAKSVDALWDLLEAGKSTVEPAPMERIGLGHLPHDDPSRMWWGNFLDDVDAFDHDFFRITAREAQTWDPQQRIMLEVAYEALEDGGQFGASLPSRNRDFGCYIGAVMNNYYDNVACHKASAYATKGTSRSYISGAVSHFFGWTGPAITLDTACSSSMVAVHAACKAIIAGECSRAIAGGTNVITSPHDYRNLAAAGFLSPTGQCKPFDSDADGYCRAEGVGVVVLESLATAIEENDRVLGVIAGSAVSQSGNVGHIAIPDTRAQVLLHRQALSIANLSPEEVSYVEAHGTGTKVGDPIEMAGLREVYCSSPRTSPLLVSSMKGNIGHTEATAGVAGLIKVLLMMNRRSIPPQASHKRLSARIPALDPDSVAIPCHLAPWPGTARAACVASYGAAGSNAAILVREMSARHASSEREGTAQPKMQGEQPLFVSAATRDTLSLRRSEILLWLRKQKDDRSQSLELSDVLFNIWIRANHALPYTLCTTVSSMDGLELKLAEVAKESMVSRKIGSAKPTILVLGGQERQYVGLSRQCYQEWQGLRRNLDQCHESLIDMGHGGLYPGIFQTAPLADLKTFHAALFASQYSSAKAWIDSGLKVDAVVGHSFGQLAALCICEVLSLADGLKLVTGRASLIEKHWGSERGAMLSLSAPTRVVDRMLEALEAKIDYAQVACYNGPESHVVVGSEHAIKVLERHIDSRSDLRGSVVAQRLDVTHGFHSVYTEAVQPALEKLATKLEWKAPTIHLETCDTEASCAVFNHSFVAHHMRRPVHFQQAVERLSLRYPESCWISAGHAASFLRLAQKSLRGRGHHSFVCLDKVFSDASGSLARAIVELWNQGQPVQYEPFERSQQHRYRHLTPPPYPFARGRHWLPFTNPTTNQASAIGPVSQQPQDEDDFLSLAGGDAQGGAEFFIPAAHPRFQLLCNGHVMSGEALAPASFYLEVAARAVLKLRCDIQSGQLDLRVEKLSMNAPVGNDPRNIIRLRLEQHEGSRDSWLFSVTIEKSPRADTDDINPQNRVVGVVSLRSRDQTNAVAGFKDFRISDVRRRYQGIVDDESAERMQGNHVYRAFDSVVTYGEVFRRIKSIACVGTEAAGEIEWNRRCGDTHVAAATDTQSIDGFMQAAGFLVNYFNNDDFQDSLFICQKIDQIDFADAIPTDAHNLSFYSRMRMDIEDAAIADVLVLDARQQQVLFAASGLQFKKLSRVPFARSLRAANAVAPCSTDGLRKQIHTGRKPASRDGSESGHRPSHRRKVLEVLANVTGTPPECVSMAGSLEDLGIDSLAAIEVLNDLRASLDMVIPLSTLMSFNSIGAFVEYVTSTVRSRERKHRGPHDVLKDARQARKTTSTAVGHEQVERDEENQVSVASSEPKHDASLDVDSSAVDLDYLAAVYPSHTRLFLAYIVEAFGRIGCDLATMAAGEAMPEMQNILPRHKRLLHRFFLALVEARILIETPEGTFVRTEQHVDKARAADLHSAAMRHWPEHGSLHQLMHAVGSRTAECLTGVFDSLQILFGDQANETALREFYQHWPLFRGATKSLAKIVLRVAGHLGRTGKIQILEVGAGTGGTTRPLLAALQVAEMPFDYHFTDVSPHLVNAARESLDGIAGLTFGVLNVEEQPPNDLEGAYDVVLASNCIHATRDLEQALGDLRCLLREDGIFALVEITRRLNIFDLVFGLLDGWWRFNDNRTHCLADEHH

>PKSI_3_Hybr_2785

MAKVERKSWRPEDVAVIGLACRFSGSASNEAKLWELLEKRESAHSKVPGERYNVEAFHQQGSTNSNNLAADGGHYLEQDVKSFDAPFFNIITKEAKAMDPQARMLLESSYEALENAGLSLESVRGSDTGCYVGCFNRDYYELLMADAEDSPEYSVTGTGFSLLANRLSWYYDLRGPSKSEDTACSSSLVALDSAYKSLLRGESKMAMVCGANLMLSPNIGLWLSKLNMLSSEGLSRSFAEGVSGYGRGEGIATVILKPLADALRDGDTIRAVINATGVNQDGHTKGITVPNSQAQSTLIESTYRRAGLDFADTGYFEAHGTGTAVGDPLEIEALERVIKHAKRTSPLHVGSIKSSIGHLEGAAGLAGLIKCILMLEKGVILPNLHFERPNRKISFENIVVPTTAVPWPEGVKRRASVNLFGYGGTNAHVIVEAFVVPCPQSRDDDLRNAQDDGLERPSPQRLFVLTGREQSTVRKMRLRYASYIQTMKDTASPSVKFDDLSYTLGKRRSRLDWAEAHVASDFKELEEKLSAPEVAATRSSNKTRLGFVFTGQGAQWPRMGLELMRYTAFRDSVEAADEYLTQKLDCSWSVIEELEKHGDESRVTSSELGQPLCTIIQVAMVDLLGSWNVRPTAVVGHSSGEIAAAYCTGVMTKQAAWQIAFHRGRECAKLKGKAPELEGSMLAVGLDVESIRPYFNNLQSGRINVACVNSPNSITISGDASEIRKLQAMLVADSVFARELPVENAYHSHHMELVAESYLHSISDVDIQHSVALSDITMVSSVTGQSIEPSELIPEYWVRNLVSPVLFADAVTAMLRGSRRRFRRGAKAEPAVDFLLELGPHATLQTPLSDIVKAQAQEDVKYASMLLRGENAVDSAMTTAGKLYCHGCPVDVTAVNDIRHECRVLVDLPAYPWNRSTKYWGVSRLMQGYLHRTHGYHSLLGARLIGSDALNPAWRHFLNLDDSPWIEEHVVHGAVVYPGAGFLSMAIEAALQLAQPGREIANVRLQNVRVLKALVIKEGEDDPEVITRFRQADSVSDEASSMRWAFEISCAKGHDERDNRATGQDEFERHATGQITLDYQPEHPYLSPLSEQIHDVRRGEYARLADTCVDTMKQDGFYEASKDVGLAYGHDFQGIATMARGPNSCCWDLRVTERSTSLPGAYESKHLIHPTTLDAIVHSLFGAMNGGKKFQNAALPVAFDSIMISPATLTTSGTKLSGFTVIREAKEREIVADIHVSSEDWSQPLVQIAGLRCTQMPSPESELHDQDARPSPVGTITYRPDIALFDENGLMNYLNERQKPESCAKVHPDLYSERLRNAVAQVVELAMFKDPGLSVLQIGGYDRGVTDSLLFTLKAESAGQALSSKIVLLDPSQETLVEIRQQYDAESSIVDALHFAVDKPLPSEVSREYGFDVVLVAIDHGLDDMSKQNVLAEAQNVLKAGGIFVVFDTLRAITERQYQHLALGEATERSPFDLALAIAKVAVVQTREGEFVEQDGSFNIPRWSYGPEMTRTIADSTVSLESDSVRLGDMAQGTPLRMLHAGHPKYAHFVTNTIQPLHLAAGQVKVELRFVDITHQDLVEPDMRALREASGVIKAVGSDVSLLRPGDNVCLSFVGHLSTSVNVDEALCQRIPPGVNMAEAACIPITLATALRALVGVAGVKPQHNVLVQAGGTKMGRAAILIASAANAVVYTTARDAEEVESLLALGISKQNIVPEGDPLLPTVTKILTGNRGWDVIVRTTKIVAETFILPECVADFGVVLDVFPSSGTGCTQETTISVMGIGSLLPEDPVLMQKTVSRIMDYLPQVSTLANSFDVFPSSAIPAALDRHGAQNQHRGVMLSFDQEDLVRVSPSATNTMKLYRDATYVMAGGLGGLGRSIARLLVDNGARNLVFLSRSGPNTTAATTMMSKMAGLGVTVKTLKCDVGDEKSVAAALDECSSMPPVRGVIQAAADIQDAIFDTYTFEQWQANLRPKVQGSWNLHCQLPEDMDFFVMLSSISGLIGHEGQAGYAAGNTFQDSLALFRHSRGLPAVTIDLGAMLDVGTIAEGSTTATFRSSDAVLMKAIDLHEIMTMCISNEINGYAIPAQVCTGLPSGGMLQVEQQEIPSYFHKPLFAALKCLGTSAVSAVNVAAPVEGVIDFAAQLTTVGSLDEADCVIANILRAHIAKAVQRAVDDIDLSQPLYSYGIDSLMAVELRAWIGEKMKADLTLFDILNAESIQALALKISKTSQLVQQEIRGMN

>PKSI_1_10304

MPDNVSFMDESQDLRHIRETSRTSHTTSILNVEGDYRNNDMPREPGECNRSTNGTVDHERMTPSADGGIPIAICGIGLRLPGGIRNDRDLYDSLYNKKDARGVIPEDRFSIDSFHSAHGKTGTIITKHGYFLQDIDLTKFDVNMFNMTPAEVERLDPHQRILLETVRETLESAGEASFRGKKVGTYVGNFTDDWLDLQNVDTVDFATYQLHGKMDFSLANRISYEYDLRGPSMTIKTACSSSALAIHEAVYSIRNGECDAAIVSGSNLNLAPRLWVGMSSQGAISPDGSSKTFDESANGYARGDGIAALFIKRLDDAVRDGNPVRAVIRSTASNADGRTPGMTMPSTEAQEALIRRAYDAANLPLSETAMVECHGTGTAVGDPMEANAVARCFGDQGMLIGSVKPNLGHSEGASAITSVVKAVLSLENRTILPNIKFHRPNPAIPWSEAKLTVPVEPLAWPKDRQERISVNSFGIGGSNVHVVLDSAASMGFRPRSLAPSKDDRPGRLLLFSGGHRASVEQSSSQHQDYVTKYPNRLSDVAYTLAKRPPVKCPGLTRSTFVFTGQGAQWLHMGKELLHESPVFAKSIGRMDSVIHSLKHAPQWTLEGIINDPENPSALTNAEISQPLCTAVQIGLVDLLKSWAIYPHAVLGHSSGEIGAAYASGVVDRAEAILLAFYRGYVCRFAQKAGGMAAVGLEKSQVIKYLQPGVCVACENSGSSVTLSGDLETLEEVLQSIRAENLNAFARKLQVGIAYHSDHMKALGGLYHQYITEHLDPKDPQVPFFSSVSGRALHSKDDFGATYWQDNLENPVLFHTAVLKSLEHTGDKQVHLEVGPHGALNGPLRQIYAETGSKARYVALQKRGANCFDTFLEGIGQLYCNGVPLQYPESADDRTLIDLPPYPWHYDHSYWSETRVMKNWRFRRQLPHDLLGLRTLDCSDAEPMWRNILRITDLPWLRDHCVGKDVVFPASGYMCMAGEAVFQETGCRDYTLREVDISTAMVLSSDHSTELLTTMKKRRLNAFLDSRWYEFLIMSYDGASWTQHCSGLVTNGPSTSHPKALLQTYDRPVSTNRWYTAMSKIGLNYGPRFTGLQNITTHVQEKKASMTIMDKQEDYESPYALHPSTLDLILQSWTVASVRGEYRRFTQLFLPTFVDEFYIGNSASKLIHLNTTAIGPDGSARGEAIGKDNDGQISFNLKGFKGSKLDNVGVDQPQEMQTIMKQQWKRDFDFADTAQLMRPAFDSTSELSLLERMFVLAAIEVHVRTSGMEGKLPHHQRYKLWIDAQIRRFGEPGYPMVEDSMELLRLDSRERQRQLCRLLEHSRKTTAHPVAEAIWRALDRIEDVFDGRIEYLDLLFNDGLMPKFYDWSNSLSDVSRLFRLLSHKKPQLKILEVGAGTGGSTARLLQYLQSDFGERQYHSYTYTDVSSGFFVQAQERFKDYEGMKYRVLDISQDPFEQGFGADEFDLICASNVLHATPRLTETLRNCRKMLRPDGILFLQELCPRQQFMGFIMGLFEGWWLGAEDGRADTPLLLPPAWDRRLRDVGFEGVEAFSFDNNPPYFMAANMTARPTVTAKSKGSITLLTFNDFLDDVAGALKQAIRAAGFEIDHCVWGEHVPLDQSLISLVDLERDKPLLQDIGDDDLRIFLDLIDAVLQTTVIWLNKPAQVSSADPNAAQMLGLARTLRAELAMHFATVEMADPTLNGMSAVVHLMCMLQRGSALPENSLDQDMEYVWAHDAMHVSRFHFQPVDEALMEMSPKFDVKTLVPLQRGMLSSLKWIGTRMLPLAEAEVQIRMSAVGMNFHDMMIAMNMFDSPLTLGSGYNSIGMEGVGYVTRKSPEVDHVQVGDRVIVIGSNSSGFATDVHRPADYCIKCPSSLTDVEAAGMSFAYMTVLWSFLDKGGLRKGQSVLIHSAAGGVGIAALHVSRWLGLEAFVTVGNEEKVRFIMNNFGLPRNRIFNSHSATFLDDVMDATAGRGVDAALSAAAGELLHNTWSCIAPGGVMLEIGKRDLINRGRLSLAPFEENRSYVGIDFSRLTIVNKPAVVRLLRQTMRLVEQGHIHPIHPTTCFDAEHAEDAFRFMQTGQHIGRIVVKIPRDTSTIPLASRPPAPEFEGQKTYLLVGGMGGLGRSVASWMVSAGARNLIFMSRSAGKSEQDQNFARELELSGCKVHCCAIDITDTDAVREALERTQASVAGVLQMAMVLRDVGIMNMDKANWDAAVAPKVQGTWNLHHALPNVDFFVMFGSNSGTLGSYGQANYAAANAFLDSFVQYRQNLGQAASVIDIGAVGDVGYVAETQVAAENMESMAGRLISEQDFLNCLQLAIARSTPTERRNKGPSTEADGYIDLKQVILLNSSILSMADPSNQIFWRKDPRMGIYRNVQRTSAESPTTDSNSLRRLVATLKADSSIADQAELAQTLARELSKQVATVLMLGEDEIDVDRSLTDVGMDSLVAIEMRNWWKQNLGVDVSVLELKDGRSILRLGELAATRLKERYSRDS
